# Bioorthogonal red and far-red fluorogenic probes for wash-free live-cell and super-resolution microscopy

**DOI:** 10.1101/2020.08.07.241687

**Authors:** Philipp Werther, Klaus Yserentant, Felix Braun, Kristin Grussmayer, Vytautas Navikas, Miao Yu, Zhibin Zhang, Michael J. Ziegler, Christoph Mayer, Antoni J. Gralak, Marvin Busch, Weijie Chi, Frank Rominger, Aleksandra Radenovic, Xiaogang Liu, Edward A. Lemke, Tiago Buckup, Dirk-Peter Herten, Richard Wombacher

## Abstract

Small-molecule fluorophores enable the observation of biomolecules in their native context with fluorescence microscopy. Specific labelling via bioorthogonal tetrazine chemistry confers minimal label size and rapid labelling kinetics. At the same time, fluorogenic tetrazine-dye conjugates exhibit efficient quenching of dyes prior to target binding. However, live-cell compatible long-wavelength fluorophores with strong fluorogenicity have been difficult to realize. Here, we report close proximity tetrazine-dye conjugates with minimal distance between tetrazine and fluorophore. Two synthetic routes give access to a series of cell permeable and impermeable dyes including highly fluorogenic far-red emitting derivatives with electron exchange as dominant excited state quenching mechanism. We demonstrate their potential for live-cell imaging in combination with unnatural amino acids, wash-free multi-colour and super-resolution STED and SOFI imaging. These dyes pave the way for advanced fluorescence imaging of biomolecules with minimal label size.

## Introduction

Over the last decades, fluorescence microscopy has become an indispensable tool to study biomolecules in their native cellular environment. A fundamental challenge is to achieve specific fluorescent labelling of biomolecules while minimizing the size of the label. This is crucial to reduce the perturbation of the biomolecule’s behaviour as well as to avoid distortions by linkage-errors in super-resolution microscopy^1^. In recent years, bioorthogonal chemistry has become an important approach to selectively introduce small organic fluorophores. Various strategies have been developed to label proteins^2,3^, nucleic acids^4,5^, sugars^6,7^ or lipids^8,9^. With the advancement of bioorthogonal chemistries, probe designs have been reported where bond formation is accompanied by fluorescence enhancement^10,11^. This so-called fluorogenic effect can substantially improve the signal-to-background ratio in fluorescence microscopy and can allow no-wash experiments^12–15^. In particular, quenched tetrazine dyes that react in inverse electron demand Diels-Alder (DA_inv_) reactions bear great potential as fluorogenic bioorthogonal labels due to fast kinetics and high biocompatibility.^16^ Previous work has shown that the quenching efficiency in tetrazine dyes determines the fluorescence turn-on of the probe upon reaction and is strongly dependent on the interchromophore distance between tetrazine and fluorophore^17–19^. Tetrazine-dye designs with short interchromophore distances, however, show only very low quenching efficiencies for far-red-shifted fluorophores compared to their blue-shifted analogues^14,20,21^.

Beyond fluorogenicity and bioconjugation functionality, fluorescent labels need to exhibit high water solubility and should not aggregate or bind unspecifically to cellular structures. Moreover, cell permeability, high fluorescence brightness and high photostability are crucial for long-term observation of intracellular targets. Rhodamines and silicon rhodamines (SiR), both xanthene-type dyes, meet these demands and are widely used in super-resolution fluorescence microscopy techniques like single-molecule localization microscopy (SMLM)^22,23^ or stimulated emission depletion microscopy (STED)^24,25^. Different structural designs for fluorogenic tetrazine conjugates have been developed for both dye classes^14,26,27^. For rhodamines, a fluorescence enhancement up to 76-fold has been reported^26^. However, so far, quenched SiR-tetrazine conjugates suffer from moderate fluorescence enhancements^14^ or do not allow live-cell applications^27^.

Here, we report a novel design concept for tetrazine-dye conjugates that enables highly efficient quenching for rhodamines and SiRs resulting in high fluorescence enhancement upon DA_inv_. We have thoroughly characterized the spectroscopic properties of the tetrazine-dyes and found evidence of Dexter exchange as the underlying fluorescence quenching mechanism. The tetrazine-dye conjugates that we name HDyes (‘Heidelberg Dyes’), exhibit excellent properties for a number of live-cell applications including the targeting of unnatural amino acids (UAA) and allow extra- and intracellular multi-colour wash-free labelling. Finally, we demonstrate that bioorthogonal labelling with HDyes enables live-cell STED as well as super-resolution optical fluctuation imaging (SOFI).

## Results and discussion

### Design and synthesis of close proximity quenched fluorogenic probes

The main challenge in the development of fluorogenic probes for bioorthogonal chemistry is to accommodate both, high fluorogenicity upon target binding and favourable properties for live-cell imaging in one molecular structure. Generally, an ideal concept enables further structural modification of the fluorophore to provide distinct molecular functionalities for special applications. In the case of fluorogenic tetrazine dyes, efficient fluorescence quenching is particularly difficult to achieve with red-shifted fluorophores^20^. To address this problem, we designed tetramethylrhodamine- (TMR) and SiR-tetrazine conjugates with minimized interchromophore distances. In this design, the tetrazine is placed in close proximity to the fluorophore by an unconjugated chemical linkage in *ortho*-position of the phenyl ring pendant to the xanthene core (Fig. 1a). We reasoned that the short and flexible oxymethyl linker would lead to a stacked conformation of fluorophore and tetrazine. Furthermore, building on literature reports we aimed to achieve distinct cell permeability properties by variation of additional phenyl ring substitutions. It has been shown that a carboxylic acid substituent in *ortho*-position results in high cell permeability owing to the formation of a transient uncharged spirolactone^12,13^. We anticipated that a carboxylic acid substituent in *para*-position lacking the possibility of spiro-lactonization yields a permanent zwitterionic, thus cell impermeable dye.

**Figure 1.**
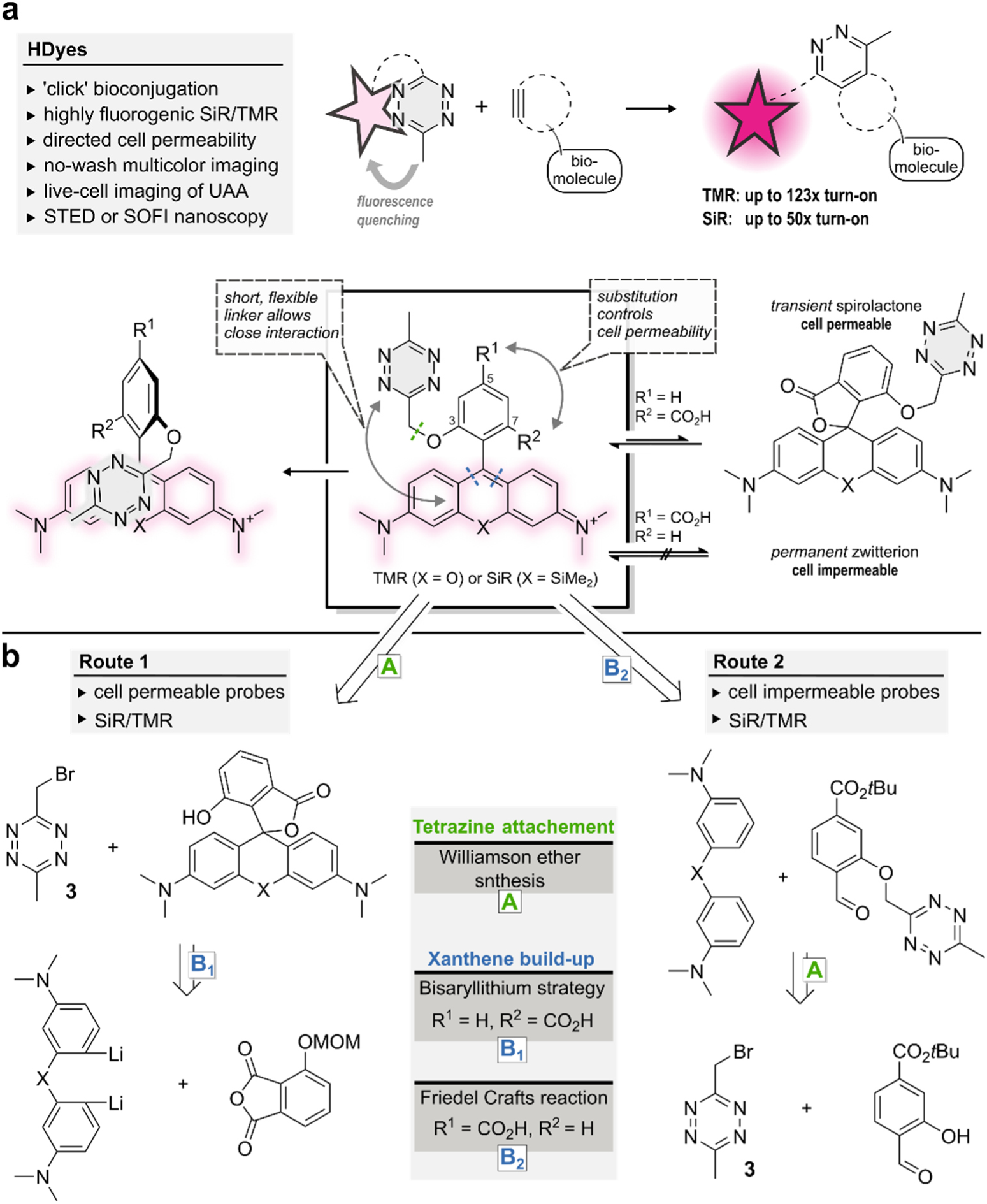
Design rationale for close proximity quenched fluorogenic probes. **a**, Tetrazine dyes can be conjugated to dienophile-modified biomolecules accompanied by an increase in fluorescence intensity. The new design allows close fluorophore tetrazine interaction leading to efficient quenching and high fluorescence turn-on. Cell permeability is controlled by varying the position of a CO_2_H-substituent. **b**, Two complementary synthetic routes provide access to fluorogenic SiR and TMR tetrazine probes with distinct cell permeability (Supplementary Schemes 1-4, Supplementary Table 2). Key compound for tetrazine derivatization in both routes is the bromomethyl tetrazine building block **3**.

To realize such cell permeable and impermeable (Si)-rhodamine-tetrazines, we developed synthetic routes using bromomethyl tetrazine building block **3** (Fig. 1b, Supplementary Schemes 1-4, Supplementary Table 2). We synthesized tetrazine benzaldehydes from **3** via Williamson ether synthesis (step A) and used them in Friedel-Crafts-reactions^14^ to create 3,5-substituted (Si)-rhodamines (route 2, step B_2_). Subsequent *tert*-butyl ester cleavage afforded cell impermeable HDyes, **HD561x** and **HD654x** (Table 1). However, corresponding cell permeable 3,7-substituted derivatives were not accessible through this route presumably due to inactivation of the benzaldehyde intermediate by hemi-acetal formation and problematic ester cleavages. Yet, we were able to obtain phenolic (Si)-rhodamines by addition of bisaryllithium intermediates to phthalic anhydrides^28^ and subsequent MOM-deprotection (route 1, step B_1_). These were reacted with **3** in Williamson ether syntheses to furnish the respective 7-CO_2_H cell permeable HDyes **HD555** and **HD653** (see Supplementary Fig. 1 for all structures).

**Table 1:** Spectral properties of HDyes. All measurements were performed in phosphate-buffered saline (PBS pH 7.4, 5 µM dye) and turn-on experiments with 15 eq BCN unless noted otherwise. ^a^ 10 eq EGFP-HaloTag-BCN. ^b^ sodium phosphate buffer pH 3.5. ^c^ 50 mM SDS in PBS pH 7.4. Properties of respective BCN-cycloadducts of the dyes listed in parentheses.

| Name | X | R<sup>1</sup> | R<sup>2</sup> | $\lambda_{\text{abs}}$<br>[nm] | $\lambda_{\text{em}}$<br>[nm] | $\epsilon$<br>[10<sup>4</sup>(Mcm)<sup>-1</sup>] | $\Phi_F$ | D<sub>0.5</sub> | turn-on | intra-<br>cellular application | extra- |
| --- | --- | --- | --- | --- | --- | --- | --- | --- | --- | --- | --- |
| **o-TzR**<br>(Ref-1) | O | H | H | 557<br>(557) | 583<br>(582) | 5.52<br>(7.41) | 0.003<br>(0.447) | - | 95 | - | - |
| **o-TzSiR** | SiMe<sub>2</sub> | H | H | 652 | 671 | 2.16 | 0.007 | - | 45 | - | - |
| **HD555**<br>(Ref-3) | O | H | CO<sub>2</sub>H | 555<br>(556) | 581<br>(583) | 5.10<br>(10.0) | 0.004<br>(0.571) | 17<br>(19) | 113/123<sup>a</sup> | + | + |
| **HD561x** | O | CO<sub>2</sub>H | H | 561 | 588 | 7.70 | 0.007 | - | 45 | - | + |
| **HD653**<br>(Ref-4) | SiMe<sub>2</sub> | H | CO<sub>2</sub>H | 653<br>(652) | 676<br>(674) | 4.86<br>(5.92) | 0.022<sup>c</sup><br>(0.477<sup>c</sup>) | 60<br>(63) | 50<sup>a</sup> | + | + |
| **HD654x** | SiMe<sub>2</sub> | CO<sub>2</sub>H | H | 654 | 673 | 12.5 | 0.007 | - | 33 | - | + |
| **sb-HD656**<br>(Ref-5) | SiMe<sub>2</sub> | H | CH<sub>2</sub>OH | 656<br>(656) | 676<br>(677) | 11.4<br>(10.4) | 0.026<br>(0.310) | - | 9.0<sup>b</sup> | + | + |

### Photophysical characterization and investigation of the quenching mechanism

Next, we studied the photophysical properties and the quenching mechanism of *ortho*-oxymethyl-linked tetrazine rhodamines. We synthesized the basic structure ***o*-TzR** (Fig. 2a, Supplementary Scheme 3) and found that it is highly quenched, with a fluorescence quantum yield (Φ_F_) of 0.3%. In addition to ***o*-TzR**, we synthesized regioisomers ***m***- and ***p*-TzR** where the oxymethyl-linked tetrazine is in *meta*- and *para*-position, respectively. Further, we reacted ***o*-TzR** with (bicyclo[6.1.0]non-4-yn-9-yl)methanol (BCN) to isolate the cycloadduct **Ref-1** and synthesized the structurally related methoxy-substituted **Ref-2**, both serving as unquenched reference dyes (Fig. 2a). We obtained the intramolecular tetrazine-xanthene distances (R) for ***o*/*m*/*p*-TzR** from geometry-optimized structures using DFT calculations (Fig. 2b). These computational results confirmed our hypothesis that ***o*-TzR** affords a stacked tetrazine-fluorophore conformation with a smaller R value than those of ***m*-TzR** and ***p*-TzR**.

**Figure 2.**
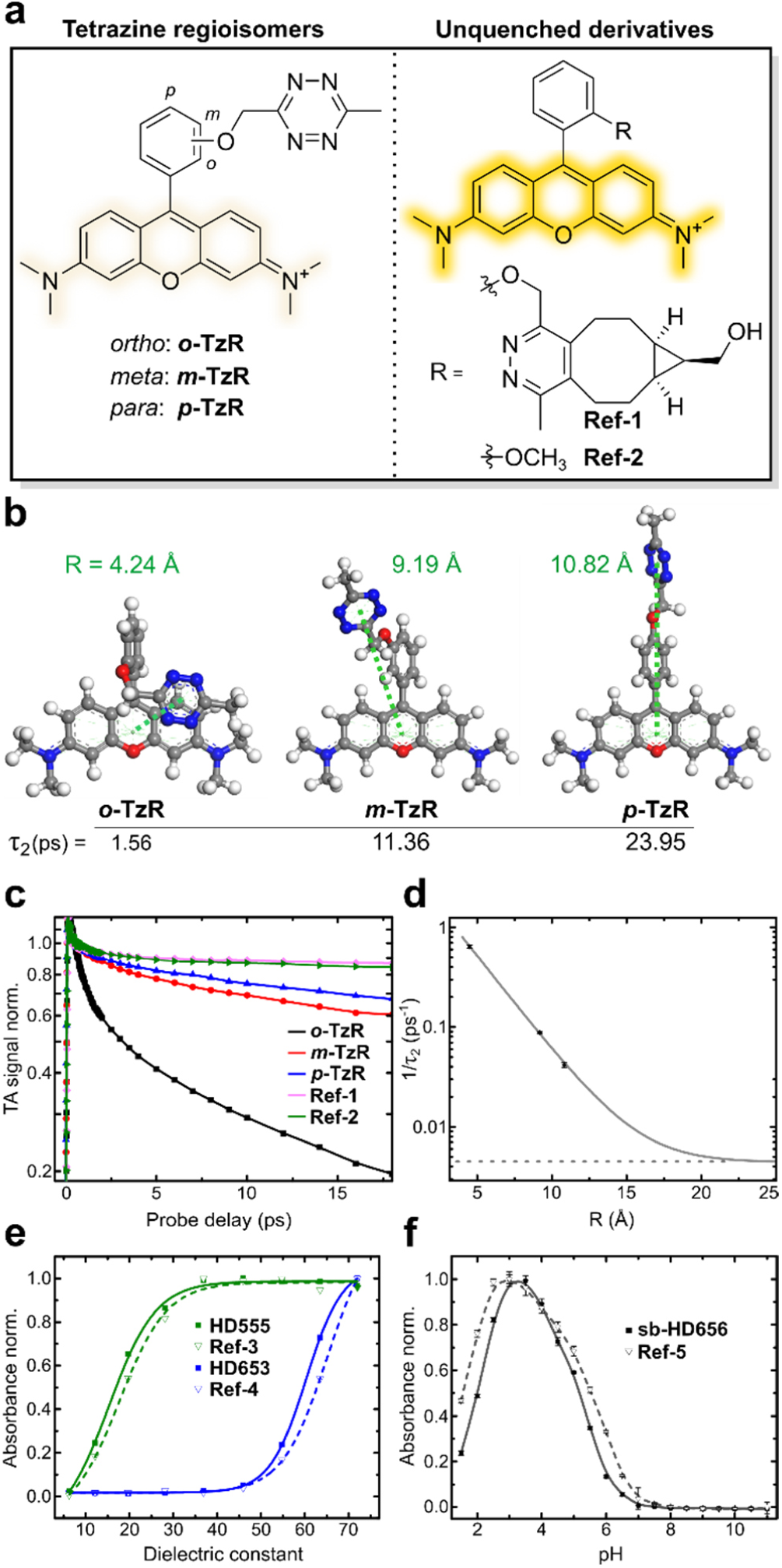
Fluorescence quenching and physicochemical properties of HDyes. **a-d**, Time-resolved spectroscopy to study fluorescence quenching: **a**, Structures of tetrazine rhodamine regioisomers and unquenched reference dyes. **b**, DFT-optimized structures of ***o*/*m*/*p*-TzR**, respective tetrazine-xanthene intramolecular distances (R) and transient absorption decay times *τ*_2_. **c**, Selected transient absorption traces at 560 nm (max. of GSB/SE). Data was normalized at a probe delay of 0.5 ps to avoid contributions of coherence spike and vibration coherence at earlier delay times. **d**, Experimental decay rates (1/*τ*_2_) at 560 nm obtained from global fitting of the transient absorption data and its dependence on R. Single exponential fitting was performed assuming a constant offset (dashed horizontal line) obtained from the average stimulated emission lifetimes of non-quenched derivatives **Ref-1** and **Ref-2** at the same detection wavelength. **e-f**, Solvatochromic spirocyclization properties: **e**, Normalized absorbance of 7-CO_2_H tetrazine dyes and respective cycloadducts (zwitterionic form) versus the dielectric constant of water/dioxane mixtures (v/v; 10/90 to 90/10). **f**, pH-dependent normalized absorbance (measured in triplicates) of 7-CH_2_OH tetrazine dye **sb-HD656** and respective cycloadduct **Ref-5**.

We then studied the distance dependence of the quenching by recording femtosecond transient absorption (TA) spectra with variable probe delay after a 20 fs excitation pulse at 575 nm. We found that all compounds show ground-state bleach (GSB) and stimulated emission (SE) bands with a maximum at 560 nm (Supplementary Fig. 12 and Supplementary Note 1). The absorbance at 560 nm in dependence of the probe delay shows the different decays of all five compounds (Fig. 2c). Reference compounds **Ref-1** and **Ref-2** lacking a quenching group showed a minor signal recovery within 1.3 ps followed by a very slow decay compatible with their emission characteristics. In contrast, the TA signal recovery was strongly accelerated for tetrazine regioisomers ***o*/*m*/*p*-TzR**: While in case of ***m*/*p*-TzR** the TA signal decayed almost to half of its initial amplitude after 16 ps, it was quenched to below 20% of the initial amplitude for ***o*-TzR** (Supplementary Fig. 12). Accordingly, ***m***- and ***p*-TzR** exhibited substantially lower turn-on in DA_inv_ of 6.9- and 8.5-fold, respectively (Supplementary Fig. 3, Supplementary Table 1). For a quantitative assessment of the TA data, we performed global target analysis with a sequential model to the full data set (Supplementary Fig. 14 and Supplementary Note 2). The fast recovery with a time constant *τ*_2_ of 1.56(±0.06) ps for ***o*-TzR** confirmed an efficient quenching compared to ***m*-TzR** and ***p*-TzR** with *τ*_2_ of 11.36(±0.30) ps and 23.95(±1.30) ps, respectively. **Ref-1** and **Ref-2**, lacking the quenching group, yielded time constants of 258(±10) ps and 196(±3) ps (Supplementary Table 4).

**Figure 3:**
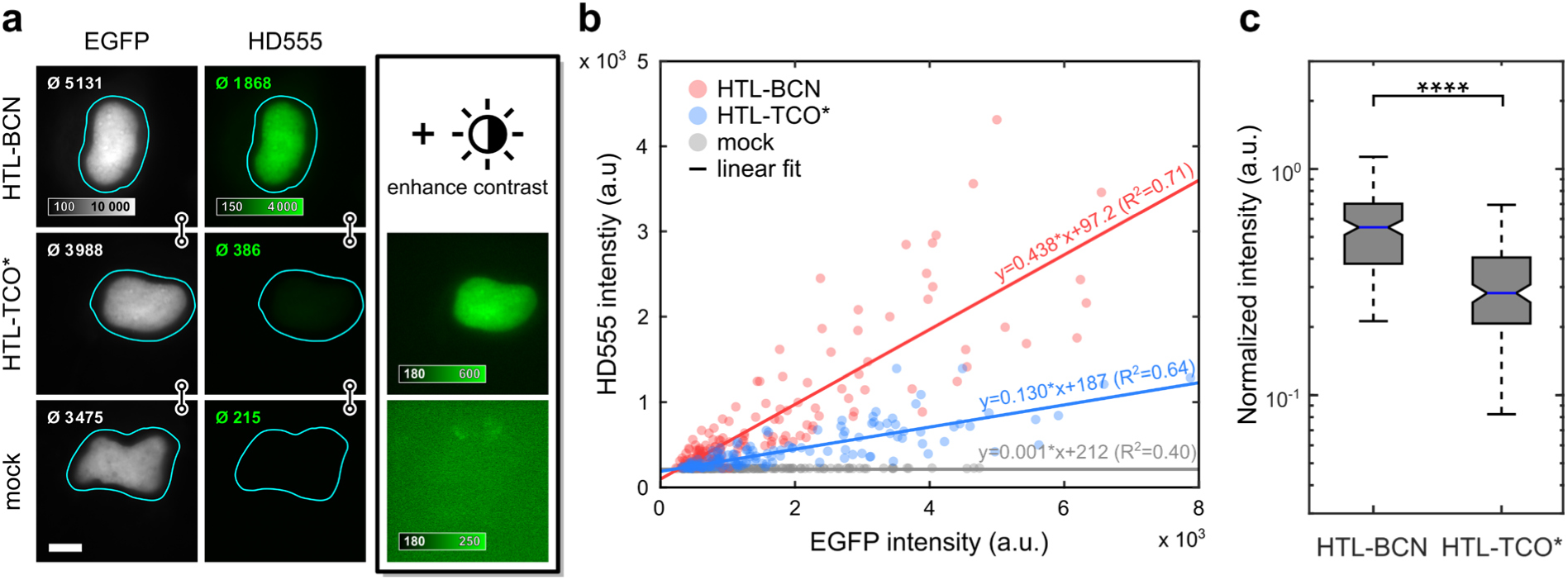
In cellulo fluorescence intensity quantification of HD555. COS-7 cells expressing H2B-EGFP-HaloTag labelled with **HD555** via HTL-BCN or HTL-TCO* and a mock control omitting any HaloTag linker. **a**, Epifluorescence microscopy images of cells with comparable expression level in EGFP and **HD555** channels with indicated mean intensity within segmented area. **HD555** channel is shown at identical display settings (indicated by linked images) and with enhanced contrast. **b**, Mean nucleus intensities of 151 (HTL-BCN), 145 (HTL-TCO*) and 147 (mock) cells pooled from three independent experiments in EGFP and **HD555** channel. **c**, Box plot of **HD555** intensities normalized to EGFP are significantly higher for HTL-BCN compared to HTL-TCO* (Wilcoxon rank-sum test, p < 10^−23^).

The interchromophore distance dependence of the decay rate *τ*_2_^-1^ (Fig. 2d) for ***o*/*m*/*p*-TzR** could be well described with an exponential fit using an offset from the unquenched reference dyes **Ref-1** and **Ref-2**. These findings rule out resonance energy transfer (RET) (*τ*_2_^−1^ ∝ *R*^−6^) and point towards an electron transfer-based quenching mechanism, like Dexter-type electron exchange or photo-induced electron transfer (both *τ*_2_^−1^ ∝ *e*^−*R*^)^29^. Additionally, the spectral features of the TA signals of ***o*/*m*/*p*-TzR** serve to further differentiate the mechanism (see Supplementary Note 2). Essentially, for ***o*-TzR**, GSB, SE and excited state absorption (ESA) signals are recovered simultaneously, which we interpret as an indication for a solely Dexter-type electron exchange mechanism. In contrast, for ***m*/*p*-TzR** the GSB/ESA recovery is delayed compared to the SE recovery. Therefore, we conclude that at these higher interchromophore distances another quenching mechanism begins to contribute.

In TA measurements, we observed that fluorescence quenching occurs from an *ortho*-tetrazine substituent with high efficiency, yielding a low Φ_F_ and a fast decay time for ***o*-TzR**. This led to a high fluorescence turn-on of 95-fold in kinetic measurements in reaction with BCN (Table 1, Supplementary Fig. 3). Importantly, the red-shifted analogue ***o*-TzSiR** exhibited comparable fluorogenicity with a Φ_F_ of 0.7% and 45-fold turn-on. Likewise, CO_2_H-substituted HDyes showed turn-ons of 123-, 45-, 50- and 33-fold for **HD555, HD561x, HD653** and **HD654x**, respectively (Table 1, Supplementary Fig. 4-6).

**Figure 4:**
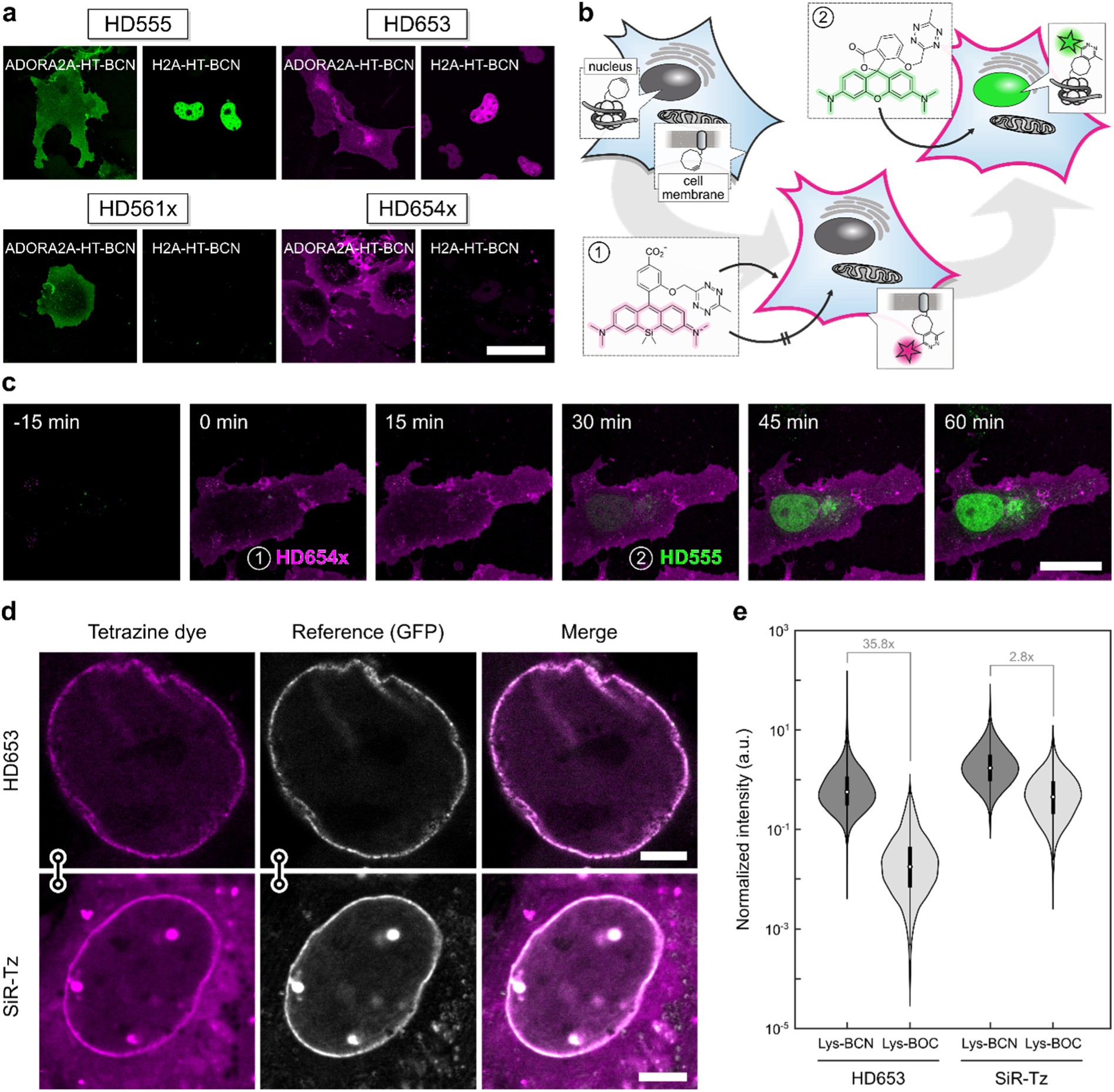
Wash-free multi-colour and unnatural amino acid imaging in live cells. **a**, Wash-free labelling of intra- and extracellular targets in COS-7 cells. Cells were transiently transfected with ADORA2A-HaloTag (plasma membrane) or H2A-HaloTag (nucleus), loaded with HTL-BCN and incubated with HDyes at 1 µM for 30 min. Scale bar 50 µm. For reference and controls see Supplementary Fig. 17-20. **b**, Dual colour labelling with a single chemistry. Intra- and extracellular proteins H2A and ADORA2A are targeted via HaloTag and HTL-BCN and then incubated with cell impermeable **HD654x** for 30 min followed by addition of **HD555. c**, Time-lapse confocal microscopy shows labelling of intra- and extracellular targets with spectrally distinct fluorophores can be achieved without any washing. Scale bar 20 µm. For individual colour channels and reference staining see (Supplementary Fig. 23). **d**, Unnatural amino acid labelling of pEGFP^N149TAG^-Nup153 with Lys-BCN in COS-7 cells. Live-cell confocal microscopy after incubation with tetrazine dyes **HD653** and **SiR-Tz** at 500 nM for 30 min and mere buffer replacement. Scale bar 5 µm. For controls see Supplementary Fig. 25. **e**, Comparison of labelling specificity with flow cytometry analysis. pEGFP^N149TAG^-Nup153 COS-7 cells were loaded with either Lys-BCN for specific or Lys-BOC for unspecific labelling. Intensity of **HD653** and **SiR-Tz** was normalized to EGFP and signal-to-background ratios calculated from population medians.

With regards to live-cell labelling of intracellular targets, we examined the solvatochromic spirocyclization behaviour that determines the cell permeability of TMR and SiR dyes. As mentioned above, 3,7-substituted **HD555** and **HD653** can transiently form an uncharged and non-chromophoric spirolactone from the zwitterionic and chromophoric open form. Therefore, we studied the solvent polarity dependent spirolactone-zwitterion equilibrium of the 7-CO_2_H dyes. We measured the absorbance of **HD555** and **HD653** along with their respective cycloadducts **Ref-3** and **Ref-4** in dioxane-water mixtures of varying composition (Fig. 2e, Supplementary Fig. 7) to determine D_0.5_, the dielectric constant at half-maximum absorption^25,30^ (Table 1). While **HD555** (D_0.5_ = 17) resided predominantly in the zwitterionic form even in low polarity media, **HD653** (D_0.5_ = 60) had a distinct propensity to adopt to the colourless spirolactone which is in good accordance with reports for other TMR and SiR fluorophores^30,31^. Of note, **Ref-3** (D_0.5_ = 19) and **Ref-4** (D_0.5_ = 63) had slightly higher propensities to adopt to lactone forms than their respective tetrazine parent dyes. The formation of a non-chromophoric, spirocyclic species is also a central feature of self-blinking hydroxymethyl-substituted SiR dyes. It was shown that the reversible transition from a fluorescent zwitterion to a non-fluorescent spiroether results in fluorophore blinking that enables stochastic switching-based super-resolution microscopy such as SMLM^32^ or SOFI^33^. Recently, we showed that the combination of fluorogenic labelling with self-blinking (sb) dyes decreases background localizations in SMLM^34^ and environment-sensitive sb-tetrazine probes were used for long-term SMLM^35^. To complement our set of HDyes with a self-blinking SiR, we synthesized **sb-HD656** from 7-hydroxy-phthalide following route 1 (Supplementary Scheme 4). We evaluated the pH-dependence of the spiroether-zwitterion equilibrium of **sb-HD656** and its cycloadduct **Ref-5** (Fig. 1f, Supplementary Fig. 8). In both cases, we observed low absorbance at physiological pH, which indicates the predominance of the closed, non-fluorescent spiroether isomer (pK_cycl_ 5.2 and 5.8 for **sb-HD656** and **Ref-5**, respectively). The on-fraction of ∼5% at physiological pH opens the possibility of SOFI, which does not rely on single emitter localization and is therefore compatible with a wide range of illumination and blinking conditions^36^.

### Quantification of fluorescence intensity in live cells

Having confirmed generally favourable spectral and physicochemical properties of the dyes, we next evaluated their suitability for bioconjugation. After successful in vitro labelling of BCN-modified EGFP (Supplementary Fig. 11), we tested if HDyes are suited for live-cell labelling. While we used BCN as dienophile for in vitro characterization due to its formation of a single product, we tested both, BCN and *trans*-cyclooctene (TCO) as dienophiles in live cells. TCO derivatives have been reported to possess superior kinetics over BCN^37–39^ and it is further known that the choice of the DA_inv_-dienophile can influence the fluorescence turn-on of tetrazine dyes^26,40,41^. We therefore compared the brightness of cell permeable **HD555** upon conjugation with a BCN- or (E)-cyclooct-2-en-1-ol (TCO*)-modified intracellular target.

First, COS-7 cells transiently expressing a histone fusion protein H2B-HaloTag-EGFP were loaded with respective HaloTag ligand-dienophile (HTL-dienophile), a bifunctional linker, to provide specific binding sites for fluorogenic labelling with **HD555**. With HTL-BCN, we observed specific staining with excellent contrast after dye incubation at 1 µM over 2 h and mere buffer replacement (Fig. 3a, top row). Second, we tested TCO* as DA_inv_-dienophile under comparable conditions. Here, we observed specific signal, but substantially lower contrast (Fig. 3a, middle row). Accordingly, mean intensities (see inset in nuclei images) of the nucleus were significantly lower for HTL-TCO* compared to HTL-BCN. A mock control omitting HTL-dienophiles (Fig. 3a, bottom row) showed no unspecific staining from unreacted dye with intensity values at background level. For a quantitative evaluation of fluorescence intensities with HTL-BCN or HTL-TCO*, we compared mean **HD555** intensities against EGFP intensity, which served as expression control (Fig. 3b). For both dienophiles, we observed a linear correlation of **HD555** and EGFP signal with considerable spread (Supplementary Fig. 16) and a 3.4-fold higher intensity obtained using HTL-BCN compared to HTL-TCO*. Considering the observed spread, we additionally compared **HD555** intensities normalized to EGFP, confirming the significantly reduced in cellulo emission intensity obtained with TCO* (Fig. 3c). This was in good accordance with the different emission intensities obtained after reaction of **HD555** with BCN and TCO* in vitro (Supplementary Fig. 15). Due to the higher obtainable fluorescence brightness, we employed BCN as DA_inv_-dienophile for cellular labelling in all following experiments.

### Intra- and extracellular wash-free protein imaging in living cells

Cell permeability of synthetic small-molecule fluorophores is a crucial factor in live-cell imaging. Hence, after having identified a suitable DA_inv_-dienophile for live-cell conditions, we next evaluated the cell permeability of the newly developed tetrazine dye series, including the potentially impermeable 3,5-substituted derivatives. This was tested using HTL-BCN conjugated to HaloTag fusion proteins with intra- or extracellular localization. In transiently transfected COS-7 cells all four HDyes showed specific extracellular labelling of adenosine A_2A_ receptor (ADORA2A) via HaloTag-BCN. Incubation with 1 µM dye for 30 min without further washing yielded uniform membrane staining with negligible background and excellent overlap with a reference label (Fig. 4a, Supplementary Fig. 17-20). To address an intracellular target, we incubated COS-7 cells transiently expressing histone H2A-HaloTag loaded with HTL-BCN for 30 min with 1 µM dye. Imaging under wash-free conditions showed that only **HD555** and **HD653**, dyes capable of forming transient spirolactones, were able to pass through the plasma membrane and specifically label the nuclear target. Using **HD654x** and **HD561x** under identical labelling and imaging conditions showed no nuclear signal (Fig. 4a, Supplementary Fig. 17-20). Omission of HTL-BCN yielded negligible signal for all four dyes (Supplementary Fig. 17-20). In contrast to these zwitterionic and net neutral dyes, positively charged dyes ***o*-TzR** and ***o*-TzSiR** showed significant unspecific mitochondrial accumulation (Supplementary Fig. 21, 22).

We then used complementary dye pairs with distinct cell permeability and absorption/emission wavelengths to perform dual-target labelling for two-colour microscopy. Frist, we labelled an extracellular target at saturating concentrations with cell impermeable **HD654x**, followed by labelling of an intracellular target with cell permeable **HD555** (Fig. 4b). This concept allows to exploit the high efficiency of labelling via DA_inv_ with fluorogenic tetrazines on two targets in parallel, which overcomes the need for orthogonal labelling reactions. To demonstrate this, we loaded COS-7 cells co-expressing ADORA2A-HaloTag and H2A-HaloTag with HTL-BCN. After incubation with **HD654x** at 1 µM for 30 min, we added **HD555** to yield a concentration of 1 µM without additional washing steps (Fig. 4c). The crucial factor is a complete saturation of extracellular BCN-modified proteins with the impermeable dye to avoid cross reactivity of the second dye. This is achievable due to high reaction rates of the DA_inv_ reaction and elevated dye concentrations facilitated by effective quenching of unreacted dyes. Accordingly, unspecific background and cross-labelling of remaining extracellular target with the secondly added dye was not observed (for individual colour channel images see Supplementary Fig. 23). After a total of 60 min and without any washing we observed high contrast specific target labelling as indicated by reference chase staining with HTL-Rhodamine110 (Supplementary Fig. 23).

Spectrally interchanged labelling using **HD561x** for extra- and **HD653** for intracellular labelling yielded the expected similar results (Supplementary Fig. 24). Therefore, our dyes enable live-cell two-colour microscopy via specific dual targeting under wash-free conditions with full spectral flexibility.

Encouraged by these results for intra- and extracellular labelling using protein tags, we next moved on to apply HDyes for live-cell protein labelling using UAAs. The labelling of UAA-modified proteins with small-molecule fluorophores via DA_inv_ results in a minimal size modification, which reduces perturbation of the native protein and linkage errors in super-resolved images.^42,43^ Due to low cellular autofluorescence in the far-red spectral region, we anticipated the highest contrast from SiR dyes. We therefore used **HD653** for site-specific labelling of the nuclear pore complex via the Amber (TAG) mutant nucleoporin 153 construct (EGFP^N149TAG^–Nup153) and performed a side-by-side comparison with commercial **SiR-Tz** (for structure, see Supplementary Fig. 9). To account for an additional fluorogenic effect due to environmental influence on the zwitterion-spirolactone equilibrium, we determined fluorescence turn-on values with a purified BCN-tagged protein of 55-fold for **HD653** and 3.1-fold for **SiR-Tz** (Supplementary Fig. 6). COS-7 cells were transiently transfected in presence of the respective genetic code expansion machinery with the Nup153 construct and in the presence of Lys-BCN, washed and then labelled with **HD653** or **SiR-Tz** (see methods). Confocal microscopy (Fig. 4d) showed that **HD653** yields comparable fluorescent signal but lower background than **SiR-Tz**. Such background can be attributed to residual Lys-BCN in the cell and/or unspecific labelling by the dye. We then used nonreactive Lys-BOC to further assess the labelling specificity and observed unspecific background staining from **SiR-Tz** in contrast to barely detectable background from **HD653** (Supplementary Fig. 25). To quantify the labelling specificity, we performed flow cytometry analysis of specific (Lys-BCN) and unspecific (Lys-BOC) labelling of the Nup153 construct with **HD653** and **SiR-Tz**, respectively (Fig. 4e and Supplementary Fig. 26). Due to efficient quenching, the signal-to-background ratio of **HD653** (35.8x) was substantially higher compared to **SiR-Tz** (2.8x). This shows that the high fluorogenicity of **HD653** allows for UAA live-cell labelling with minimal washing and enables high signal-to-background ratio for fluorescence imaging applications.

### STED and SOFI super-resolution microscopy

With improving spatial resolution of imaging techniques, the size of fluorescent labels becomes increasingly important. Having shown that specific labelling of UAA with improved contrast could be achieved with our dyes, we evaluated their use in super-resolution microscopy. In this context we used **HD653** in combination with a minimal label size UAA. Based on previous reports using structurally akin SiRs^12,24,31^ we evaluated the performance of **HD653** in STED microscopy. To test its applicability, we labelled the Amber mutant vimentin^N116TAG^-mOrange construct with Lys-BCN and **HD653** in live COS-7 cells. STED imaging using a 775 nm continuous wave laser for depletion resolved vimentin structures and clearly showed enhanced resolution compared to confocal microscopy (Fig. 5a). Of note, after 50 cycles of STED imaging, bleaching for **HD653**-labelled vimentin was lower compared to commercial SiR-Tz (Supplementary Fig. 27).

**Figure 5:**
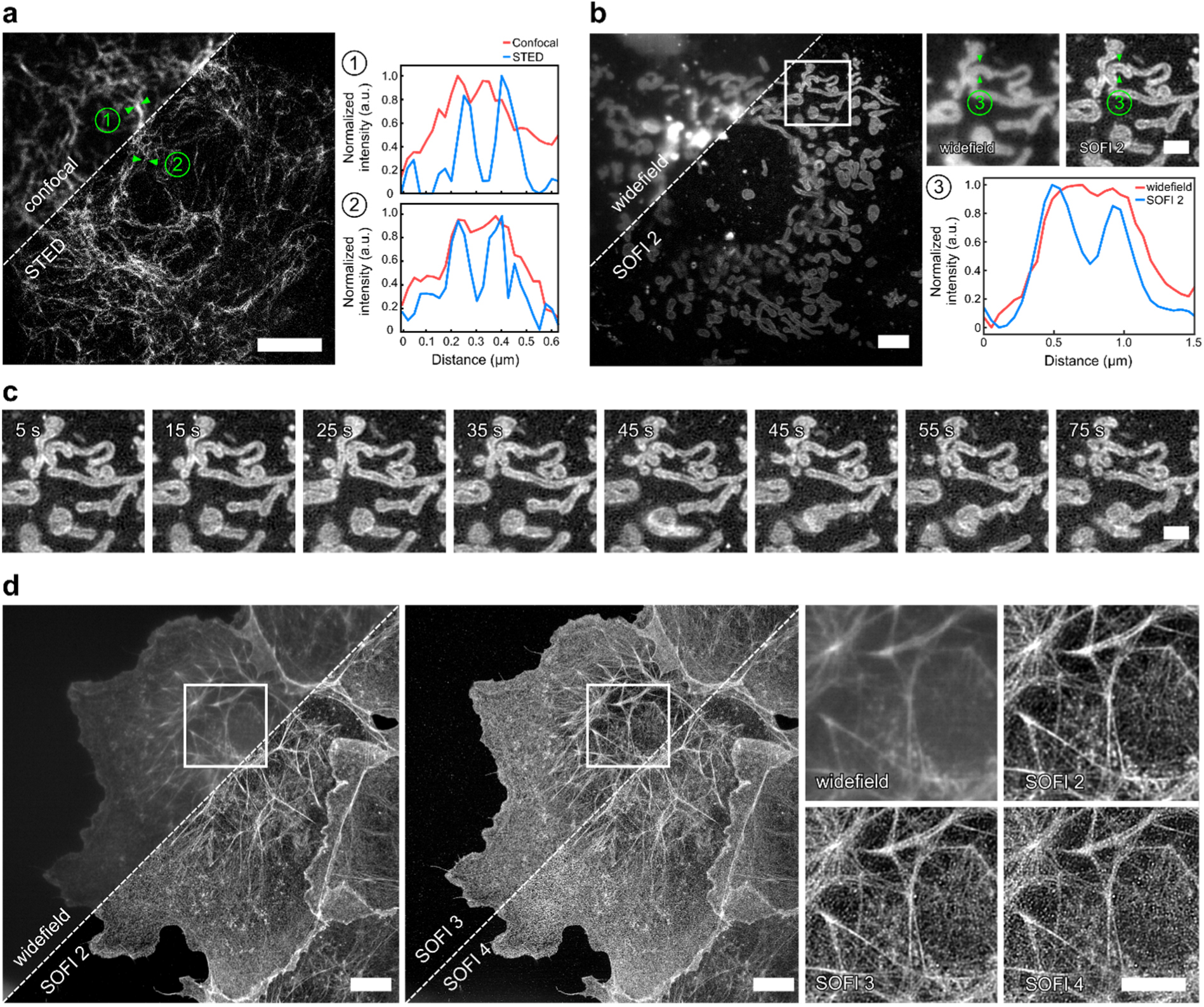
Super-resolution microscopy with HDyes. **a**, Live-cell confocal and STED microscopy of COS-7 cells expressing pVimentin^N116TAG→BCN^ labelled with **HD653** with intensity line-profiles showing the resolution improvement. Scale bar 5 µm. **b**, Live-cell widefield (temporal average of series) and second order SOFI of mitochondria in COS-7 cells transiently expressing TOMM20-mCherry-HaloTag labelled with **sb-HD656** via HTL-BCN. Scale bar 5 µm. The close-ups correspond to the ROI indicated in the overview image. The cross-sectional profile of the averaged widefield image and second order SOFI shows the improved image contrast and resolution. Scale bar 2 µm. **c**, Time-course of second order SOFI corresponding to the ROI indicated in (b) showing the movement of mitochondria with a time resolution of 10 s (see Supplementary Fig. 28 for the full sequence). Image acquisition: 500 frames per time point, 20 ms exposure time, 635 nm laser 140 W/cm^2^. The images are representative of >5 cells from two independent experiments. **d**, High-order SOFI imaging of **sb-HD656** labelled f-actin in fixed COS-7 cells showing SOFI analysis up to 4^th^ cumulant order. Image acquisition: 20,000 frames, 50 ms exposure time, 635 nm laser 275 W/cm^2^. The images are representative of 6 cells from two independent experiments. Scale bars 10 µm.

Another approach for super-resolution microscopy uses stochastic temporal emission fluctuations of single fluorophores. Based on our own previous work^34^ we used a self-blinking fluorogenic tetrazine-SiR for minimally invasive live-cell super-resolution microscopy. While SMLM can achieve resolutions down to a few nanometres^44^, it requires sparse signals to allow for localization of individual emitters. In contrast, SOFI exploits spatially and temporally correlated intensity fluctuations for resolution improvement without the need of isolated emitters^45^. This is especially beneficial for live-cell microscopy of highly abundant targets at low excitation intensities. We used the self-blinking derivative **sb-HD656** in combination with SOFI for sub-diffraction imaging in living and fixed cells. First, we performed live-cell imaging of mitochondrial import receptor protein TOMM20 labelled via HaloTag and HTL-BCN with **sb-HD656** at very low illumination intensities in the near infrared (∼140 W/cm^2^ at 635 nm). Due to low phototoxicity it was possible to perform time-lapse imaging of the rearrangement of the mitochondrial network in COS-7 cells (Fig. 5c, for longer time course see Supplementary Fig. 28). Second order cumulant analysis required only a few hundred frames and resolved the mitochondrial outer membrane (see close-up and line profiles in Fig. 5b). In contrast to SMLM, SOFI was also compatible with higher active emitter densities as encountered with highly abundant histone protein H2B in nuclei of live COS-7 cells (Supplementary Fig. 29).

In order to demonstrate higher-order SOFI imaging enabled by the high photostability of HDyes, **sb-HD656** was used to stain the cytoskeletal actin network. We exploited the versatility offered by DA_inv_ tetrazine labelling and used a small-molecule labelling approach with phalloidin, a natural toxin with a high affinity to f-actin. To this end, we targeted f-actin with phalloidin-BCN in fixed COS-7 cells and subsequently labelled it with **sb-HD656**. The pH of the imaging buffer was chosen for a reduced on/off ratio for high-order SOFI analysis (Supplementary Fig. 30). **sb-HD656** was found to be photostable at relatively low illumination intensities (275 W/cm^2^), thus allowing the acquisition of up to 20,000 frames with moderately low photobleaching (Supplementary Fig. 31). SOFI analysis was performed up to 4^th^ cumulant order while maintaining sufficiently high signal to noise ratio (Fig. 5d). Compared to widefield imaging, 2^nd^ order SOFI provided a significant contrast enhancement, while higher order (3^rd^ - 4^th^) SOFI images allowed to resolve individual actin filaments that are not visible in widefield images.

## Conclusion

We developed a series of red and far-red fluorogenic tetrazine dyes (HDyes) for live-cell bioorthogonal labelling of intra- and extracellular targets. The design rationale with tetrazines flexibly linked to the fluorophore at minimal distance lead to efficient fluorescence quenching by Dexter exchange. HDyes exhibited high fluorogenicity and were well-suited for live-cell imaging under wash-free conditions as demonstrated in two-colour labelling experiments and STED super-resolution imaging of UAA-labelled target proteins. The modular synthetic route provided a self-blinking HDye, which was successfully exploited in long-term live-cell SOFI imaging. We expect the HDyes to leverage applications of bioorthogonal chemistry in the field of super-resolution microscopy. Their excellent photophysical properties, improved signal to background ratios and small labelling size are ideal prerequisites for the direct labelling of UAA-modified proteins, which has great potential of reducing the linkage error. We expect the herein presented tetrazine-xanthene scaffold to serve as general motif in the development of numerous other fluorogenic probes in the future.

## Data availability

Crystallographic data has been deposited at the Cambridge Crystallographic Data Centre (CCDC) under the CCDC deposition number 2021104. The data can be obtained free of charge via https://www.ccdc.cam.ac.uk/structures. Syntheses, compound characterization data, details of the computational and time-resolved spectroscopy studies as well as of imaging experiments, are provided in the Supplementary Information. All data supporting the findings of this work are available within the paper and the Supplementary Information. Additional materials and information are available from the corresponding authors on request.

## Acknowledgements

RW and EAL acknowledge funding from the Deutsche Forschungsgemeinschaft DFG (SPP1623, WO 1888/1-2) and DPH from the Federal Ministry of Education and Research (BMBF/VDI; MorphiQuant3D and Switch-Click-Microscopy) and the DFG (HE4559/5-1, HE4559/6-1). EAL also thanks ERC SMPFv2.0 for funding. AR, KG and VN thank the EPFL BioImaging & Optics Core Facility (EPFL-BIOP) for access to STED microscope and Horizon 2020 research and innovation program of the European Union via grant 686271/SEFRI 16.0047 for funding. XL acknowledges funding from Singapore University of Technology and Design (T1SRCI17126). ZZ acknowledges funding from National Natural Science Foundation of China (Grant No. 61675057) and China Scholarship Council. MJZ acknowledges a fellowship by the Carl-Zeiss-Stiftung. We gratefully acknowledge access to the Nikon Imaging Center at Heidelberg University. We thank Daniel Wolf for experimental help and Prof. Dr. Andres Jäschke for constant support.

## Author contributions

PW and RW designed the strategy and fluorophore structures. PW, CM, AJG and MB performed chemical syntheses under supervision of RW. KY, FB and MJZ performed cell experiments, confocal microscopy and data analysis under supervision of RW and DPH. PW, FB, ZZ, and TB characterized fluorophores, performed TA measurements, and analyzed data under supervision of TB. MY performed UAA-labelling and STED microscopy under supervision of EAL. KG and VN performed SOFI experiments under supervision of AR. WC and XL performed DFT calculations. FR solved the crystal structure. PW, KY and FB wrote the manuscript with the help of DPH and RW. All authors reviewed and approved the final version of the manuscript.

