## Supplementary Information for "Bioorthogonal red and far-red fluorogenic probes for wash-free live-cell and super-resolution microscopy"

| SUPPL. FIGURE | | PAGE |
|---|---|---|
| 1 | Overview of fluorogenic tetrazine dyes and unquenched derivatives | 3 |
| 2 | Normalized absorption and fluorescence emission spectra | 4 |
| 3 | $DA_{inv}$ turn-on experiments of tetrazine regioisomers | 6 |
| 4 | $DA_{inv}$ turn-on experiments of HD555 and HD561x | 7 |
| 5 | $DA_{inv}$ turn-on experiments of Si-rhodamine/rhosamine tetrazines | 8 |
| 6 | $DA_{inv}$ turn-on experiments of (Si)-rhodamine tetrazines with protein-bound BCN. | 9 |
| 7 | Dependence of spirolactonization on solvent dielectric constants | 10 |
| 8 | Spiroether–zwitterion equilibria of sb-HD656 and respective cycloadduct Ref-5 | 11 |
| 9 | Structures of miscellaneous compounds | 44 |
| 10 | X-ray solid-state structure of HD555 | 45 |
| 11 | In vitro labeling of EGF-HaloTag-BCN with HDyes | 49 |
| 12 | Selected transient spectra at specific delays | 50 |
| 13 | Selected transient absorption traces at 560 nm and 630 nm | 51 |
| 14 | Species associated difference spectra (SADS) | 52 |
| 15 | Emission intensity achieved with dienophiles BCN and TCO* | 55 |
| 16 | In cellulo fluorescence intensity quantification | 55 |
| 17 | Live-cell no-wash labeling of intra- and extracellular targets with HD555 | 58 |
| 18 | Live-cell no-wash labeling of intra- and extracellular targets with HD654x | 58 |
| 19 | Live-cell no-wash labeling of intra- and extracellular targets with HD561x | 59 |
| 20 | Live-cell no-wash labeling of intra- and extracellular targets with HD653 | 59 |
| 21 | Live-cell no-wash labeling of intra- and extracellular targets with o-TzR | 60 |
| 22 | Live-cell no-wash labeling of intra- and extracellular targets with o-TzSiR | 60 |
| 23 | Dual target no-wash labeling | 61 |
| 24 | Dual target no-wash labeling with HD561x and HD653 | 62 |
| 25 | Unspecific background with unnatural amino acid labeling | 65 |
| 26 | Flow cytometry analysis of HD653 and SiR-Tz labeling specificity | 66 |
| 27 | Signal loss in STED imaging | 67 |
| 28 | Time-course live-cell SOFI on mitochondria | 71 |
| 29 | SOFI of histone H2B | 71 |
| 30 | Kymograph indicating signal fluctuation over time | 72 |
| 31 | Photostability of sb-HD656 in SOFI of live and fixed cells | 73 |

| SUPPL. TABLE | | PAGE |
|---|---|---|
| 1 | Photophysical data of tetrazine regioisomers and related unquenched derivatives. | 5 |
| 2 | Optimization of reaction conditions for the synthesis of OMOM-substituted (Si)rhodamines. | 15 |
| 3 | Crystal data and structure refinement for HD555 | 46 |
| 4 | Time constants obtained via global target analysis. | 52 |

| SUPPL. SCHEME | | PAGE |
|---|---|---|
| 1 | Synthesis of tetrazine precursors | 12 |
| 2 | Synthesis of other precursors | 13 |
| 3 | Friedel-Crafts strategy for tetrazine-substituted (Si)rhosamines | 13 |
| 4 | Synthesis of cell permeable HDyes | 14 |

### 1 General information

Chemicals were purchased from Sigma-Aldrich Chemie GmbH, TCI Deutschland GmbH, abcr GmbH, Acros Organics, Alfa Aesar, Fluorochem Ltd., SiChem GmbH, Bachem Holding AG and were used as received. NMR solvents were purchased from euriso-top SAS. Polygram Sil G/UV254 TLC plates from Macherey-Nagel GmbH & Co KG and Sigma Aldrich Chemie GmbH were used for thin layer chromatography. Normal phase column chromatography was performed using silica gel from Fluka with a pore size of 60 Å and a particle size range of 40-63 µm. Solvents were p.a. quality.

HPLC analytics and semi-preparative purifications were conducted on an Agilent 1100 series HPLC system. Phenomenex Luna 3 µm and 5 µm C18 reversed-phase columns were used for these purposes (Solvent A: $H_2O$ containing 0.1% TFA; Solvent B: MeCN containing 0.1% TFA). Collected HPLC fractions were dried by lyophilization. Dye samples for photophysical measurements and imaging experiments were prepared from HPLC-purified (20-80% Solvent B / Solvent A) material (1 mM or 10 mM stock solutions in anhydrous DMSO) and stored at -20 °C.

Mass spectrometry was performed on a Bruker microTOF-QII (for ESI-MS) or a JEOL AccuTOF GCx (for EI-MS) mass spectrometer.

NMR spectra were recorded on a Varian Mercury Plus 300 MHz spectrometer or a Varian 500 MHz NMR System. Peak shifts were reported relative to solvent peaks according to Fulmer *et al.*[1] The amount of substance for dye samples ≤ 2 mg was determined from $^1H$ NMR in $CD_3CN$ or $CD_3OD$ using solutions of 2,2,2-trifluoroethanol (TFE, 20 mM) in the respective deuterated solvents as a reference. TFE was removed by lyophilization before DMSO stock solutions from the dried samples were prepared.

Fluorescence measurements were performed on a JASCO Spectrofluorometer FP-6500 or a Varian Cary Eclipse. Absorption spectra were recorded on a Varian Cary 100 Bio UV-Visible or a Varian Cary 500 Scan. The pH-dependent spirocyclization of sb-HD656 was studied using a Safire 2 multimode microplate reader in fluorescence and absorption measurements. In-gel fluorescence was scanned with a Typhoon FLA 9500 fluorescence scanner (GE Healthcare Life Sciences). Fluorescence lifetime measurements were measured with a FluoTime 100 spectrometer using a TCSPC TimeHarp 200 and a pulsed diode PLS 500 (all PicoQuant).

X-ray crystallographic measurements were executed with a STOE Stadivari instrument at 200 K.

Protein purification was performed on a FPLC-system (BioLogic DuoFlow, Bio-Rad) at 4 °C.

### 2 Photophysical characterization

##### Extinction coefficients and fluorescence quantum yields

Extinction coefficients and quantum yields were determined in PBS buffer (pH = 7.4) or in PBS containing 50 mM SDS in the case of **HD653** to achieve maximum amount of the quinoid form. Extinction coefficients were determined by measuring the absorbance at four different concentrations (1-10 µM). Quantum yields were determined as described by Brouwer[2] relative to cresyl violet in ethanol ($\Phi_f$ = 0.56[2], $\lambda_{ex}$ = 580 nm) for silicon rhodamine/rhosamine dyes or sulforhodamine 101 in ethanol ($\Phi_f$ = 0.95[3], $\lambda_{ex}$ = 520 nm) for rhodamine/rhosamine dyes.

##### Fluorescence lifetimes

Emission decay curves were analyzed with software FluoFit 4.1 from PicoQuant using an instrument response function measured using a dispersion of silica nanoparticles (LUDOX HS-40). Fluorescence kinetics were best described with a monoexponential or biexponential decay. In the latter case an amplitude weighted average of decay times is given.

##### Kinetic measurements

For time-course fluorescence measurements, stock solutions of (1*R*,8*S*,9s)-bicyclo[6.1.0]non-4-yn-9-ylmethanol (BCN) in DMSO or EGFP-HaloTag-BCN in PBS and the respective dyes in DMSO were used. Starting fluorescence intensity values were measured with a fluorimeter from 1 to 5 µM dye solutions in PBS at 37 °C. Then BCN was added (10 to 50 equiv, final DMSO content ≤ 0.6%), solutions mixed thoroughly, and measurement were started immediately. Fluorescence intensities were read out every 10 s until plateaus were reached. Temperature was kept constant at 37 °C. Dividing mean end and start fluorescence intensity values from triplicate time course measurements yielded the fluorescence enhancements (‘turn-on’).

#### UV VIS Spectra in water/dioxane mixtures

2.5 µM dye solutions were prepared in various water/1,4-dioxane mixtures (10-90%, v/v), containing 0.01% $NEt_3$,[4] and UV VIS spectra were recorded at 20 °C. Dielectric constants for the water/1,4-dioxane solvent system were taken from the literature[5], and plotted against $\lambda_{abs,max}$.

#### pH Dependent spirocyclization of sb-HD656

10 µM dye solutions were measured in sodium phosphate buffer (12 mM $Na_2HPO_4$, 88 mM $NaH_2PO_4$ adjusted to respective pH with 5 M HCl or 5 M NaOH) and measured in triplicates at 20 °C.

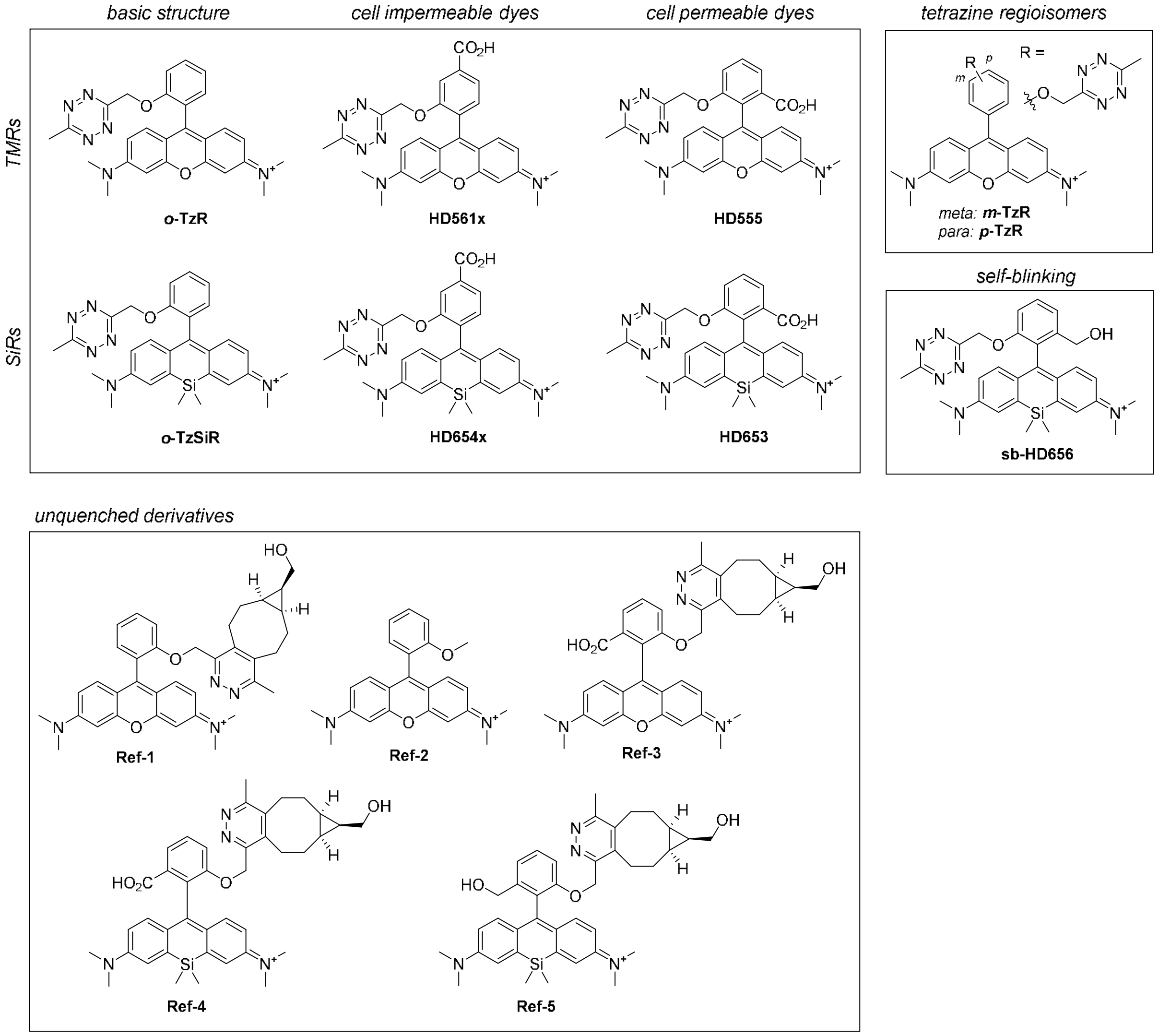

**Supplementary Figure 1. Overview of fluorogenic tetrazine dyes and unquenched derivatives.**

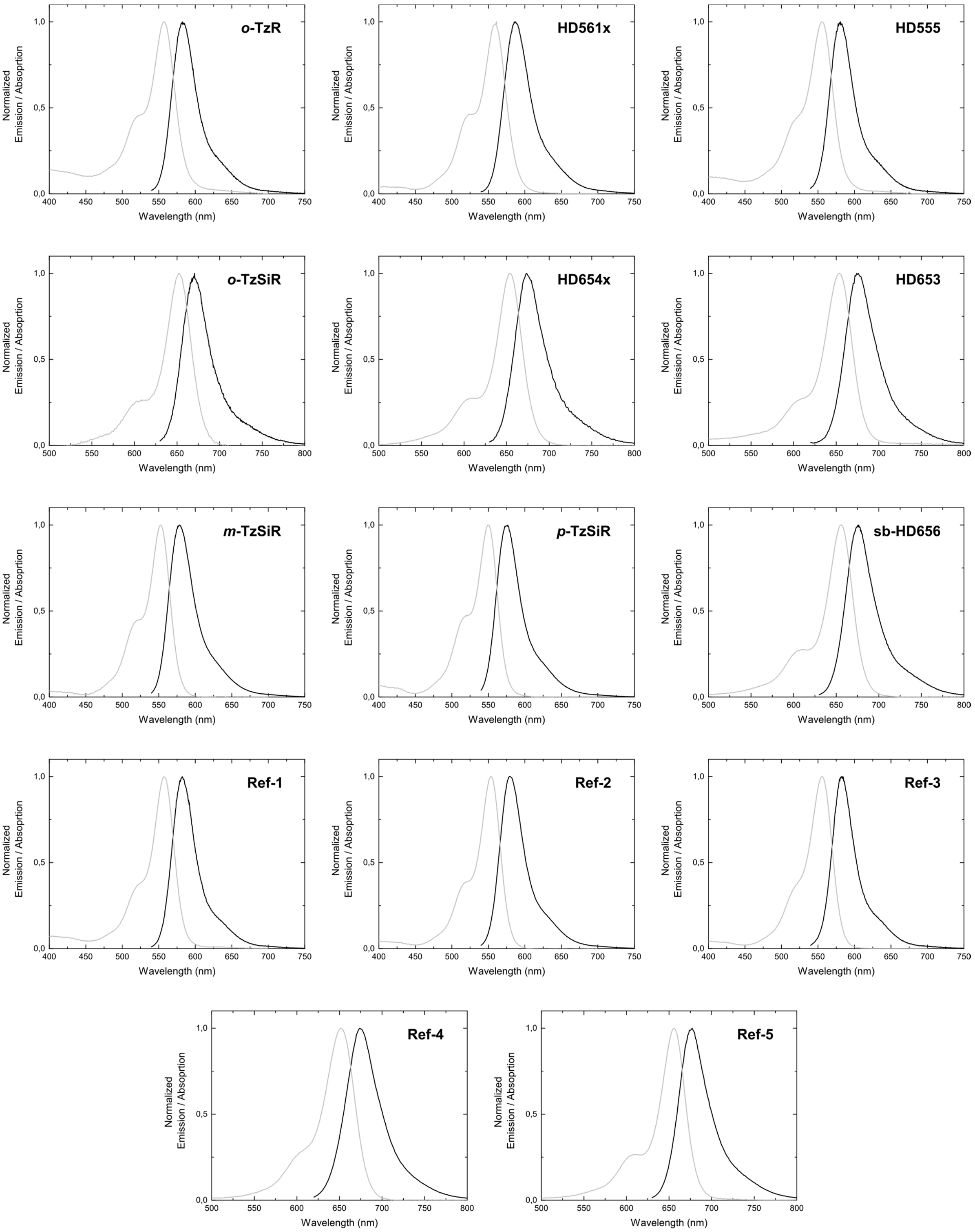

**Supplementary Figure 2. Normalized absorption and fluorescence emission spectra.**

**Supplementary Table 1. Photophysical data of tetrazine regioisomers and related unquenched derivatives.** $\tau_2$: decay time determined in transient absorption experiments. $\tau_F$: fluorescence lifetime. turn-on: 5 µM dye (in PBS) reacted with 15 eq BCN (***o*-TzR**) or 50 eq BCN (***m*-TzR**, ***p*-TzR**). $\Phi_F$: fluorescence quantum yield. [a] slow component of recovery of the TA signal was fitted over 500 ps.

| | $\lambda_{abs}$ [nm] | $\lambda_{em}$ [nm] | ε [$10^4$ (M cm)$^{-1}$] | $\tau_2$ (ps) | $\tau_F$ (ns) | turn-on | $\Phi_F$ |
|---|---|---|---|---|---|---|---|
| ***o*-TzR** | 557 | 583 | 5.52 | 1.56 | - | 95 | 0.003 |
| ***m*-TzR** | 553 | 578 | 4.87 | 11.36 | - | 6.9 | 0.026 |
| ***p*-TzR** | 550 | 577 | 6.42 | 23.95 | - | 8.5 | 0.016 |
| **Ref-1** | 557 | 582 | 7.41 | 257.0[a] | 2.42 | - | 0.447 |
| **Ref-2** | 554 | 580 | 7.42 | 196.1[a] | 2.03 | - | 0.393 |

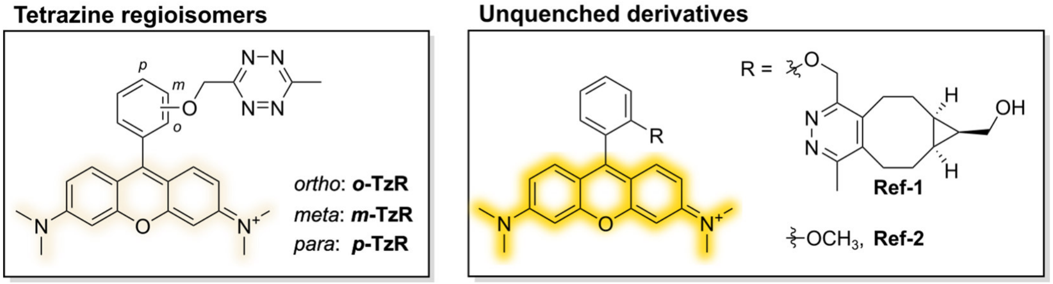

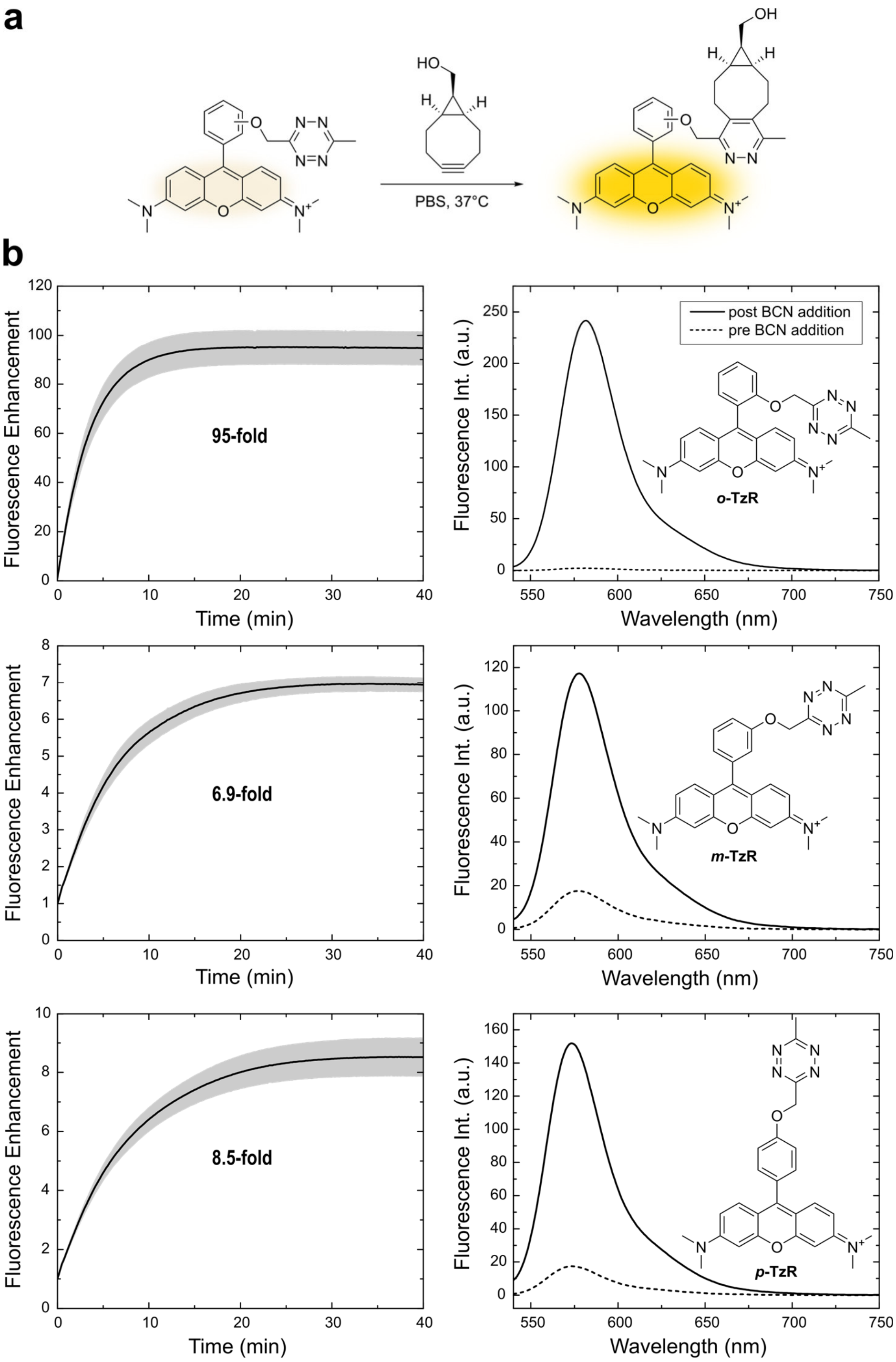

**Supplementary Figure 3. $DA_{inv}$ turn-on experiments of tetrazine regioisomers.** 5 µM dye in PBS, 15 eq BCN (***o*-TzR**) or 50 eq BCN (***m*-TzR**, ***p*-TzR**).

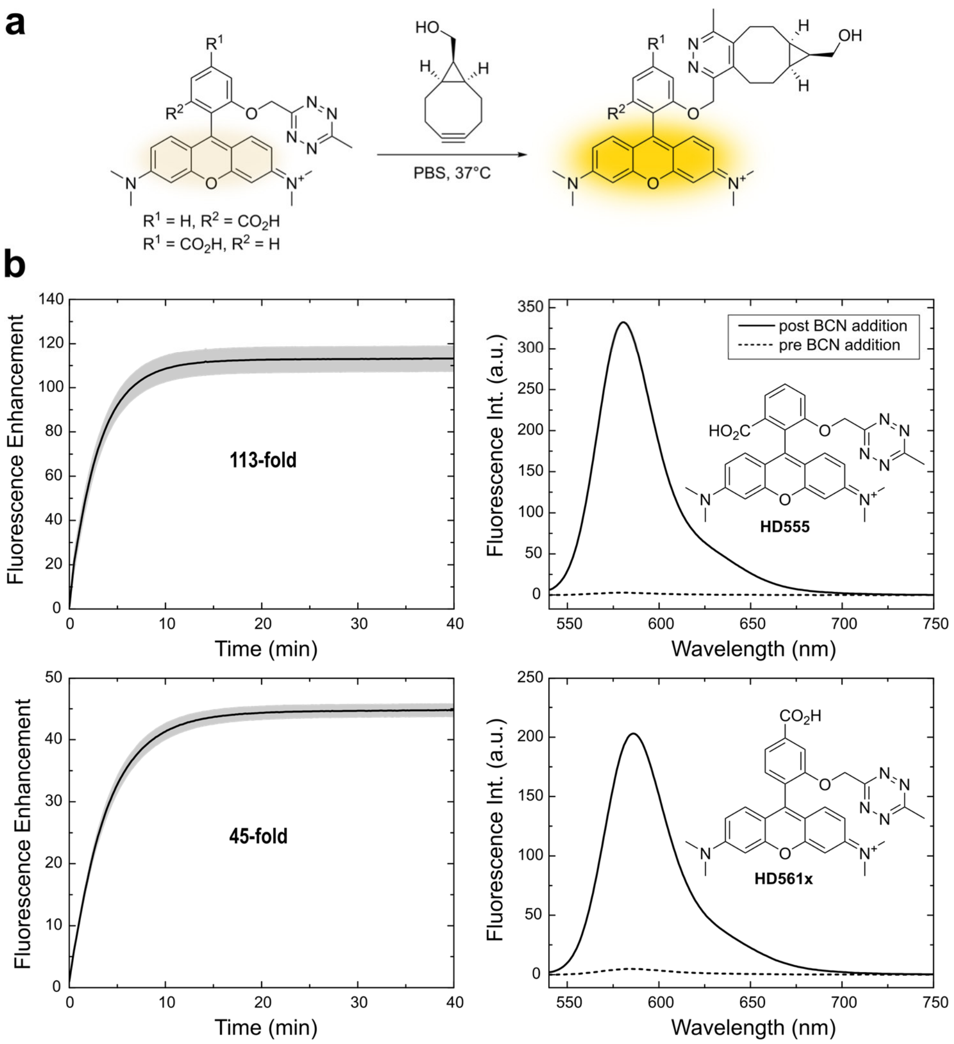

**Supplementary Figure 4. DA$_{inv}$ turn-on experiments of HD555 and HD561x.** 5 µM dye in PBS, 15 eq BCN.

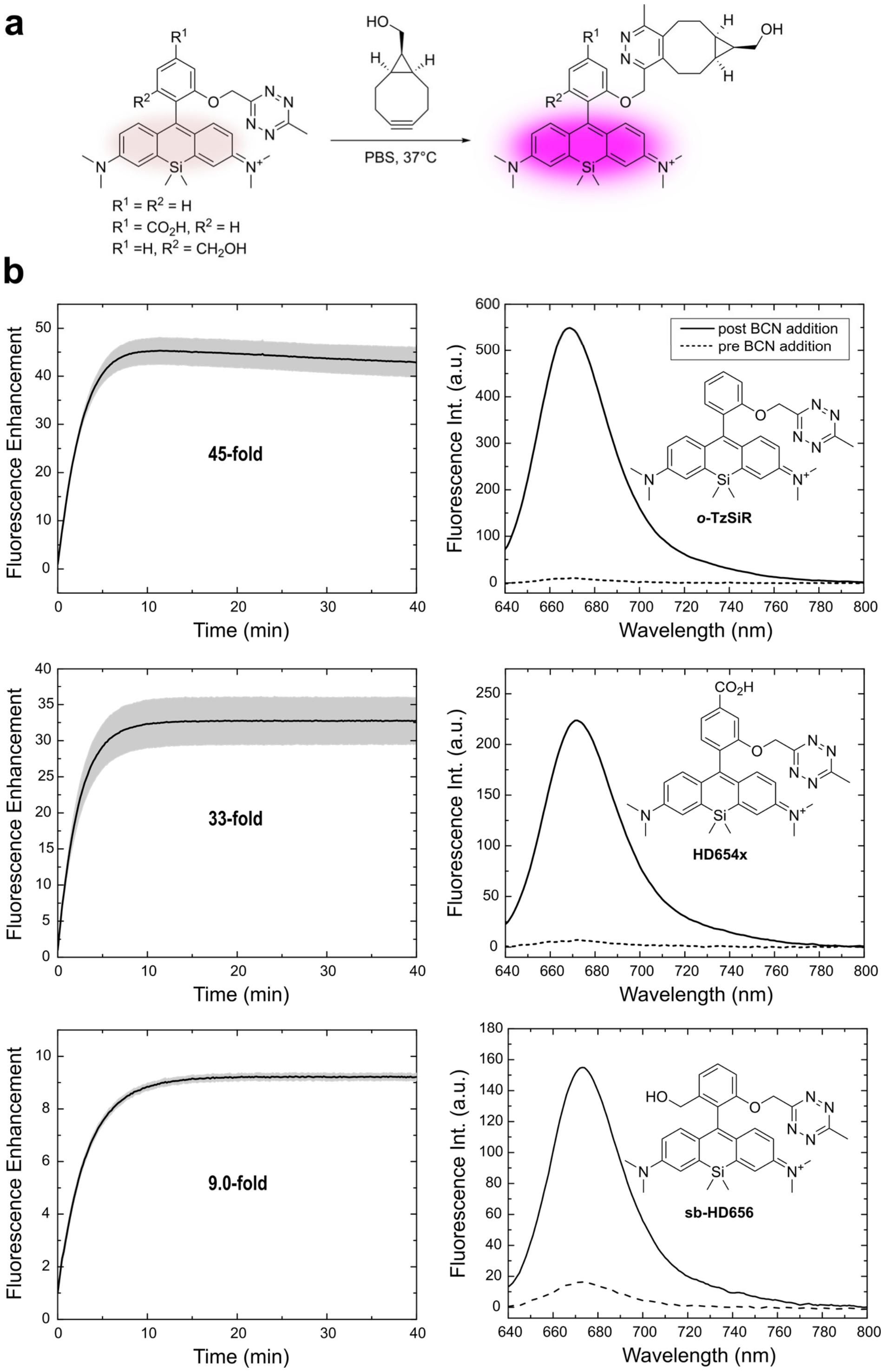

**Supplementary Figure 5. $DA_{inv}$ turn-on experiments of Si-rhodamine/rhosamine tetrazines.** 5 µM dye in PBS (***o*-TzSiR**, **HD654x**) or aqueous phosphate buffer pH 3.5 (**sb-HD656**), 15 eq BCN.

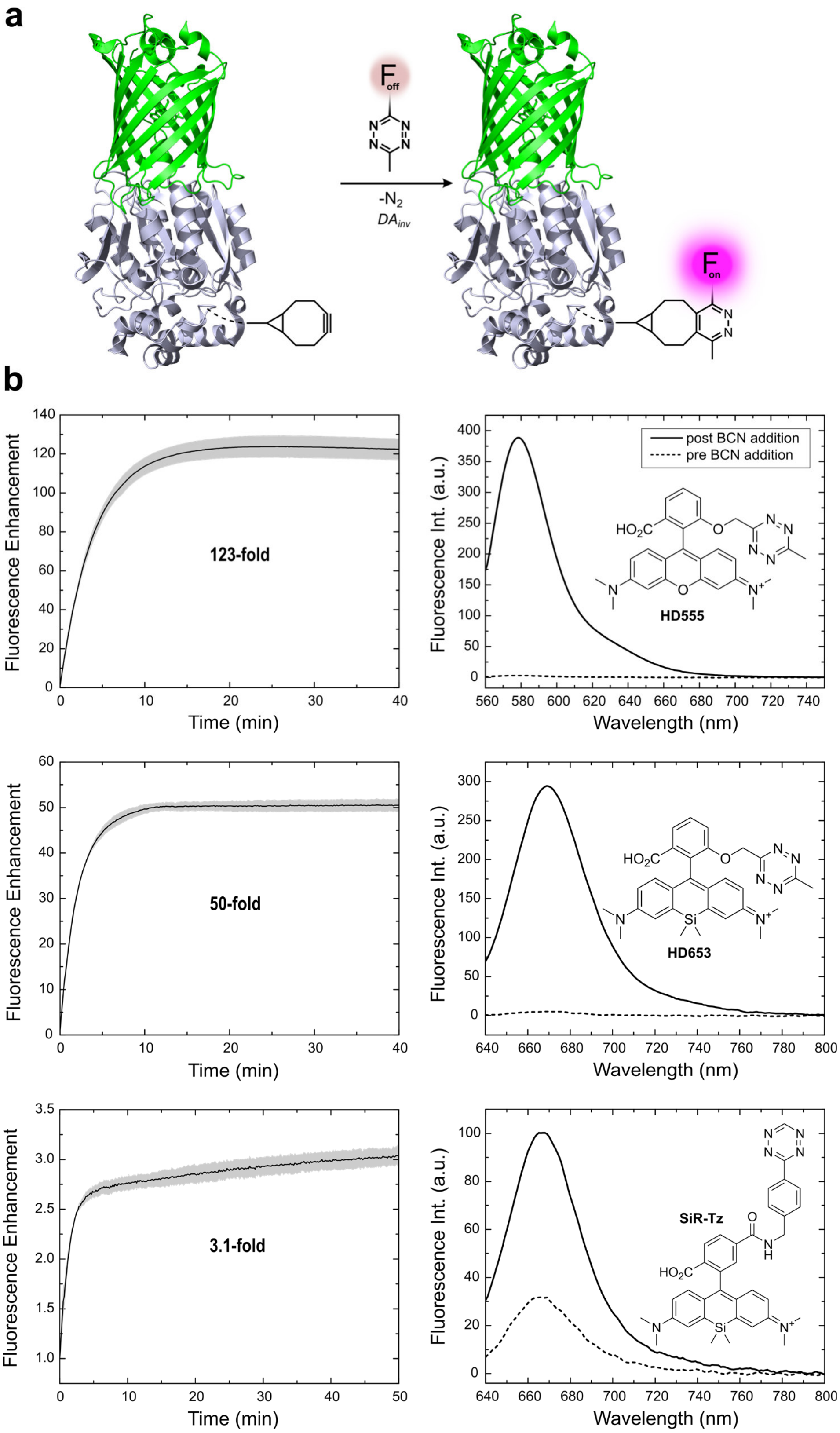

**Supplementary Figure 6. DA$_{inv}$ turn-on experiments of (Si)-rhodamine tetrazines with protein-bound BCN.** (a) Schematic presentation of the reaction (based on PDBs 2Y0G and 5Y2Y). (b) DA$_{inv}$ turn-on experiments (1 µM dye in PBS, 10 eq EGFP-HaloTag-BCN) of HDyes and commercially available **SiR-Tz**.

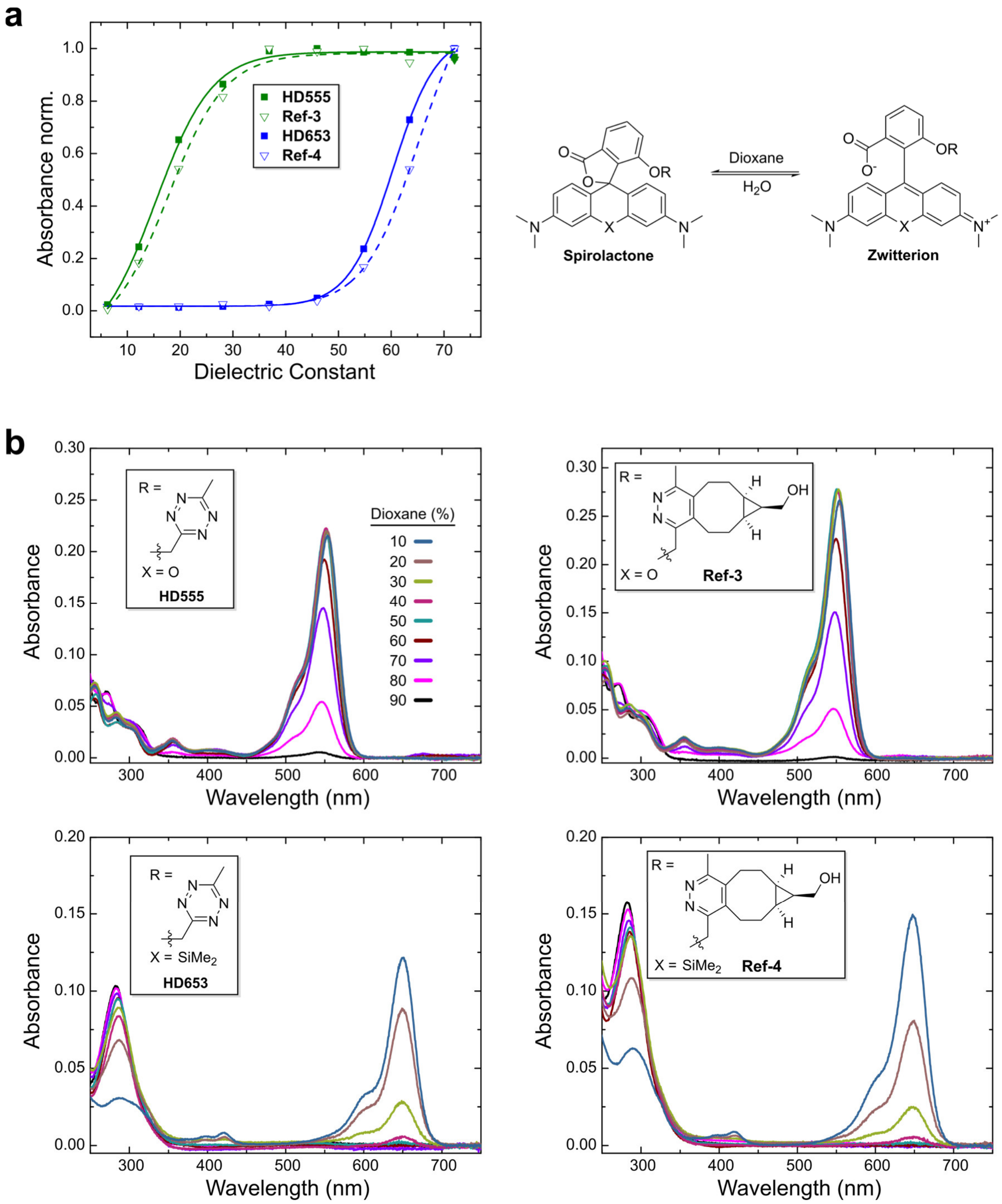

**Supplementary Figure 7. Dependence of spirolactonization on solvent dielectric constants.** (a) Normalized absorbance of tetrazine dyes and respective cycloadducts at $\lambda_{abs,\ max}$ versus the dielectric constant of water/1,4-dioxane mixtures (v/v; 10/90 to 90/10). (b) UV VIS spectra of tetrazine dyes and respective cycloadducts (2.5 µM) in water/1,4-dioxane mixtures.

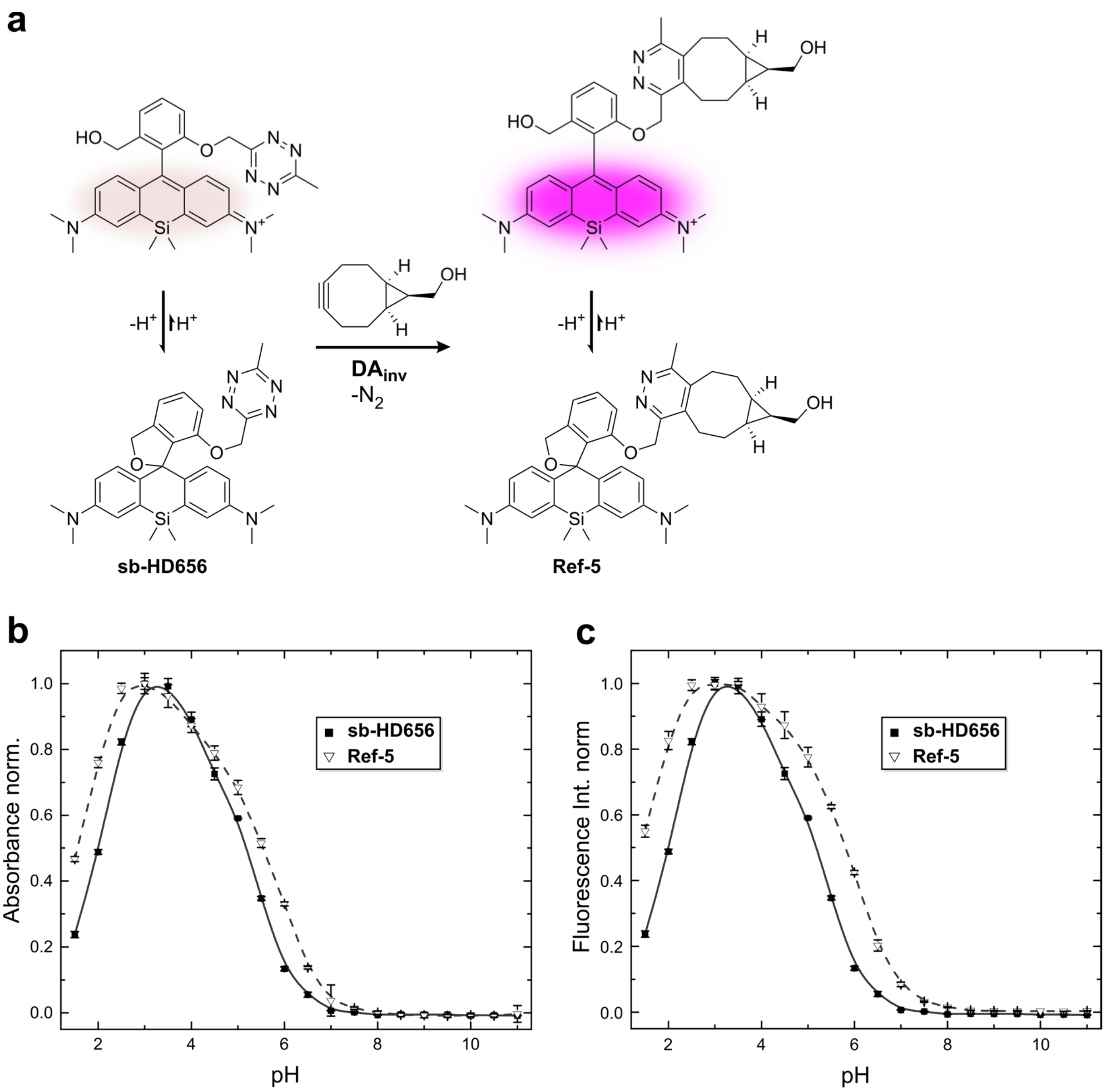

**Supplementary Figure 8. Spiroether–zwitterion equilibria of sb-HD656 and respective cycloadduct, Ref-5.** (a) Molecular mechanism of fluorescence on/off switching and turn-on. (b/c) pH-dependence of absorbance/fluorescence of 10 µM dye solutions in aqueous buffer.

### 3 Chemical synthesis

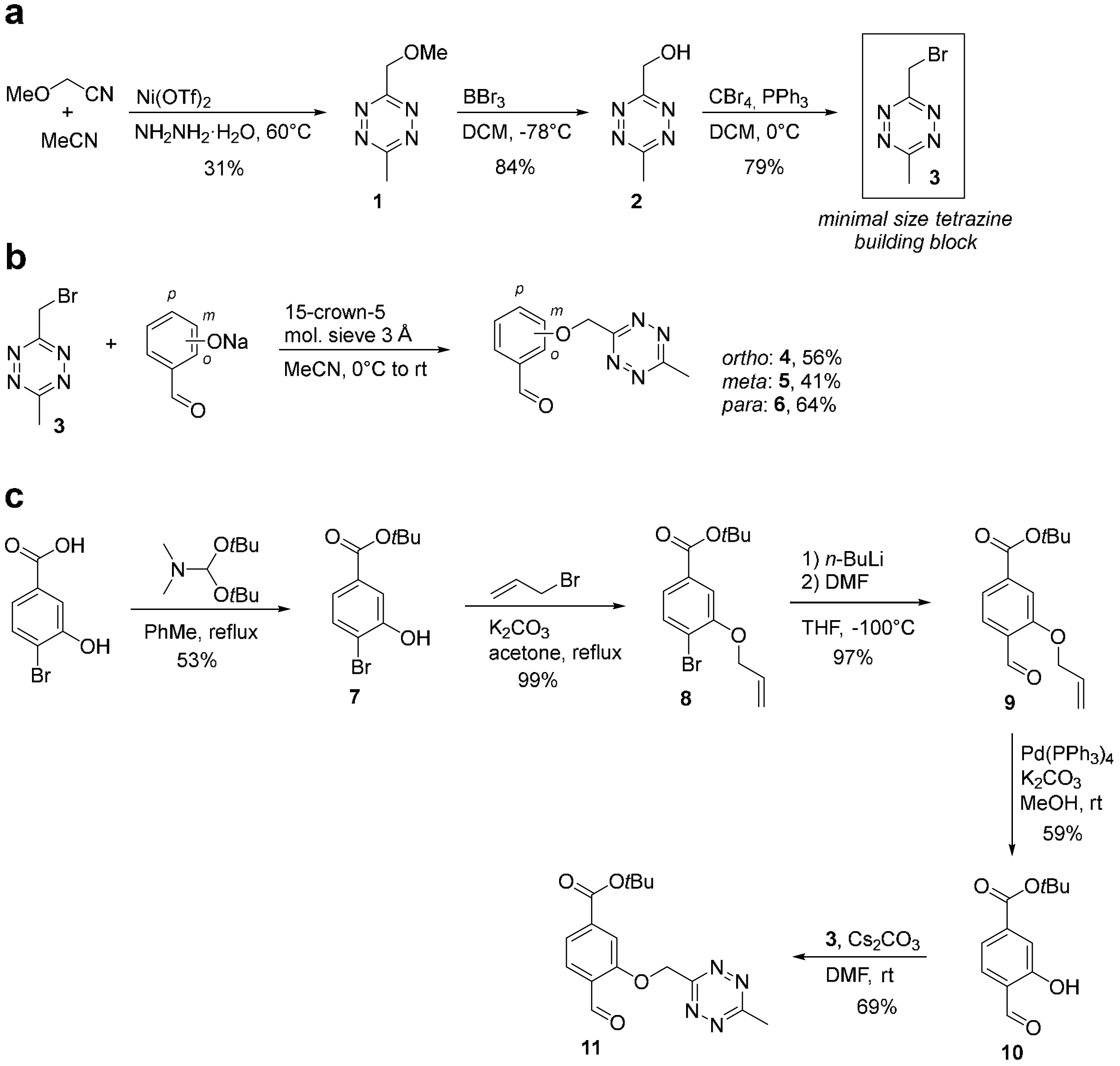

**Supplementary Scheme 1. Synthesis of tetrazine precursors.** (a) Minimal size building block bromomethyl tetrazine **3**. (b/c) Tetrazinyl benzaldehydes for subsequent Friedel-Crafts reactions.

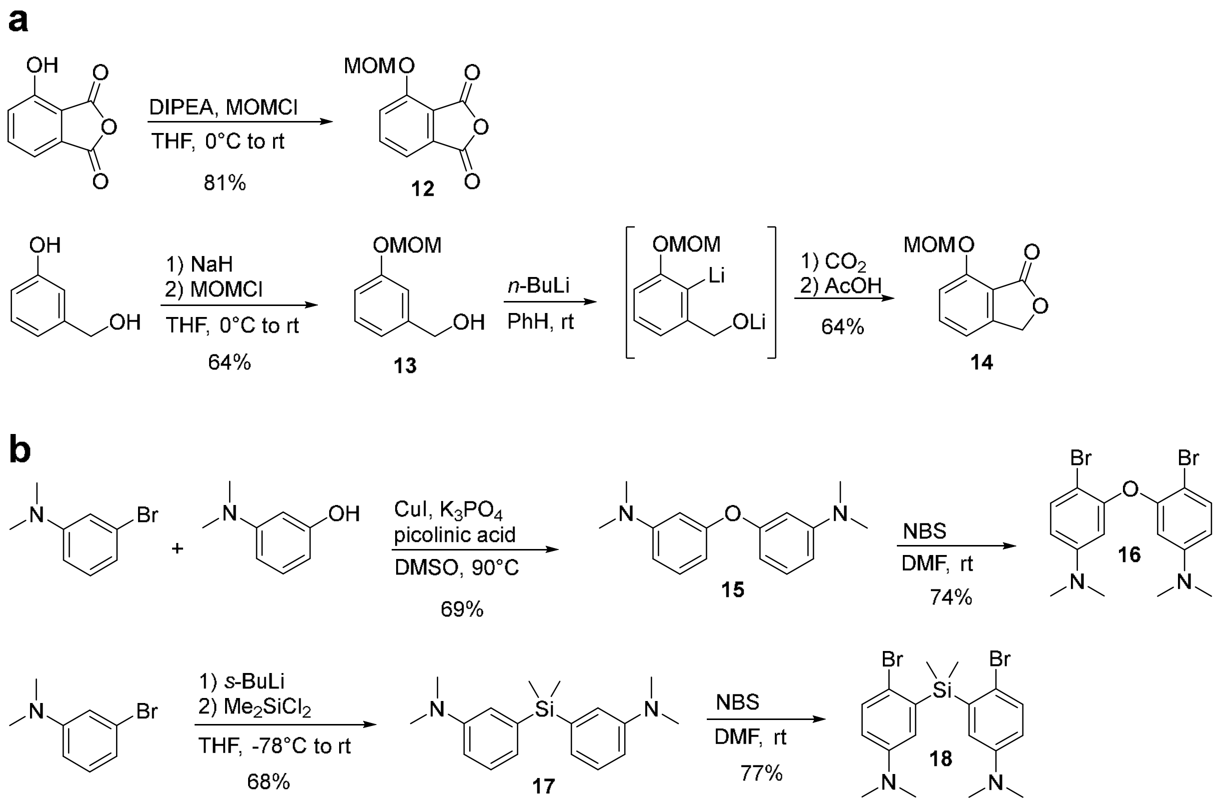

**Supplementary Scheme 2. Synthesis of other precursors.** (a) Electrophiles **12** and **14** for the reaction with bisaryllithium intermediates. (b) Friedel-Crafts nucleophiles **15**, **17**. Bromoarylether **16**/ bromoarylsilane **18** for lithium halogen exchange.

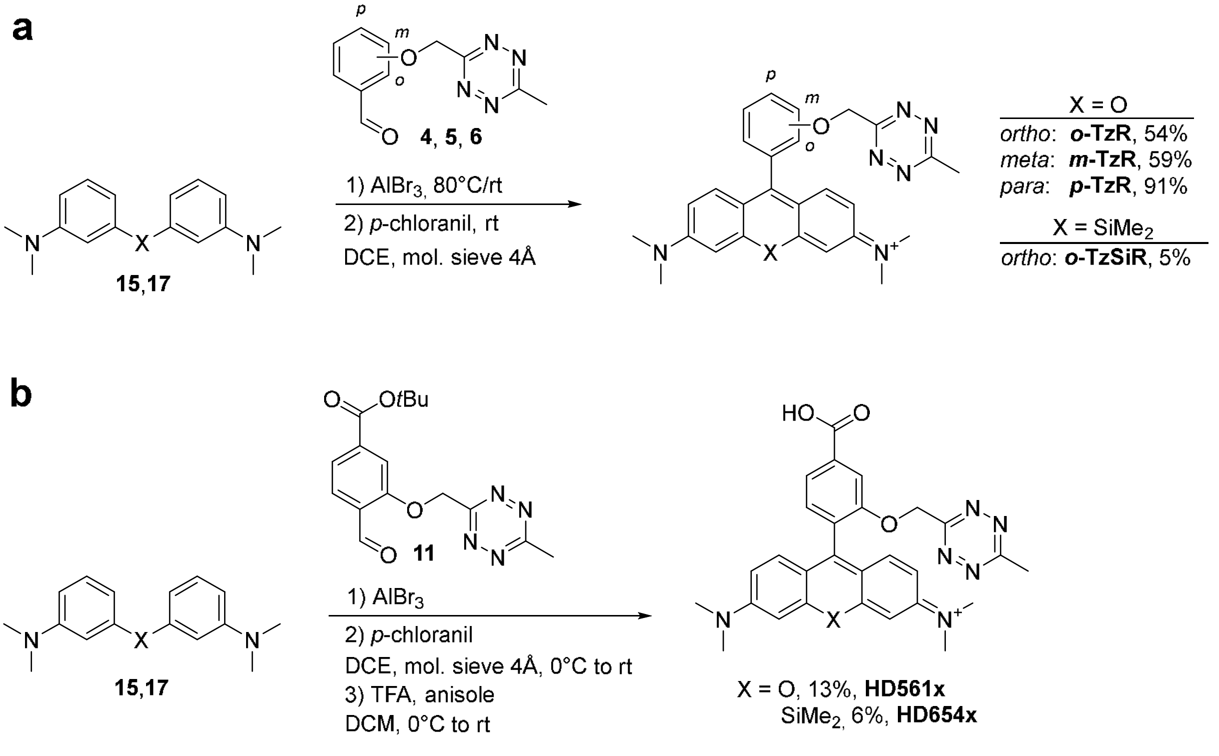

**Supplementary Scheme 3. Friedel-Crafts strategy for tetrazine-substituted (Si)rhosamines.** (a) Without additional substitutent. (b) Cell impermeable HDyes (**HD561x**, **HD654x**; extracellular application).

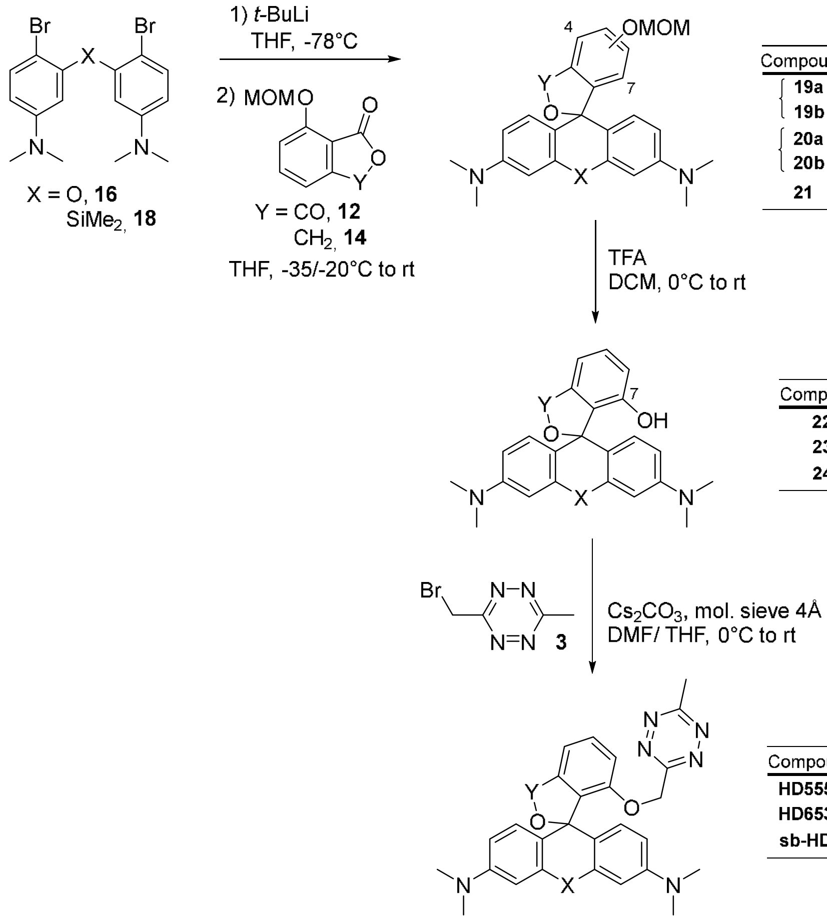

| Compound | X | Y | Pos. OMOM | Yield (%) | |
|---|---|---|---|---|---|
| **19a** | O | CO | 7 | 6 | 34 |
| **19b** | O | CO | 4 | 28 | |
| **20a** | $SiMe_2$ | CO | 7 | 13 | 42 |
| **20b** | $SiMe_2$ | CO | 4 | 29 | |
| **21** | $SiMe_2$ | $CH_2$ | 7 | 53 | |

| Compound | X | Y | Yield (%) |
|---|---|---|---|
| **22** | O | CO | 61 |
| **23** | $SiMe_2$ | CO | 75 |
| **24** | $SiMe_2$ | $CH_2$ | 85 |

| Compound | X | Y | Yield (%) |
|---|---|---|---|
| **HD555** | O | CO | 52 |
| **HD653** | $SiMe_2$ | CO | 43 |
| **sb-HD656** | $SiMe_2$ | $CH_2$ | 7 |

**Supplementary Scheme 4. Synthesis of cell permeable HDyes.** Addition of bisaryllithium ethers to phthalic anhydrides[6]/ phthalides and subsequent Williamson ether synthesis afford tetrazine-substituted (Si)rhodamines (**HD555**, **HD653**, **sb-HD656**; intra- and extracellular application). Curly brackets denote isolated regioisomers from single reaction mixtures.

**Supplementary Table 2. Optimization of reaction conditions for the synthesis of OMOM-substituted (Si)rhodamines.** The reaction of organolithium or Grignard reagents with phthalic anhydrides is based on a recent literature procedure[6]. Phthalic anhydride **12** or phthalide **14** addition over a period of 30 to 45 min. [a] determined by $^{1}$H NMR of the crude residue.

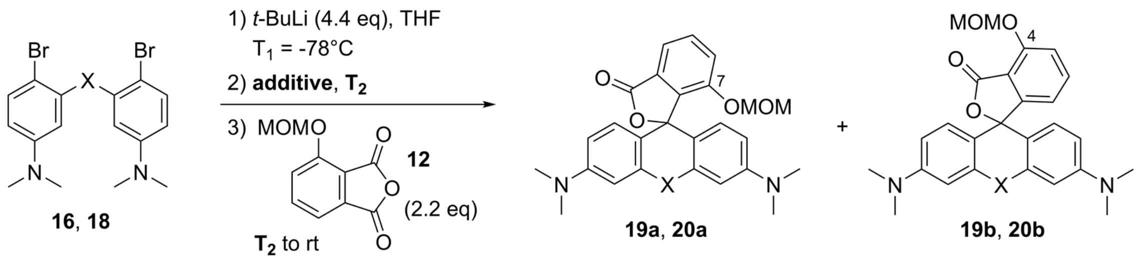

| X | scale [mmol] | additive | $T_2$ (°C) | product ratio[a] (7 : 4)-isomer | yield (%) (7 / 4)-isomer |
|---|---|---|---|---|---|
| O | 0.12 | $MgBr_2 \cdot OEt_2$ | -10 | - | <1 / <1 |
| O | 0.15 | - | -20 | 1 : 10 | <1 / 21 |
| O | 0.15 | - | -35 | 1 : 5 | 9 / 37 |
| O | 1.2 | - | -35 | 1 : 6 | 6 / 28 |
| $SiMe_2$ | 0.11 | $MgBr_2 \cdot OEt_2$ | -10 | - | 3 / 10 |
| $SiMe_2$ | 0.11 | - | -20 | 1 : 5 | 10 / 45 |
| $SiMe_2$ | 0.11 | - | 0 | 1 : 4 | 5 / 11 |
| $SiMe_2$ | 1.1 | - | -20 | 1 : 2.7 | 13 / 29 |

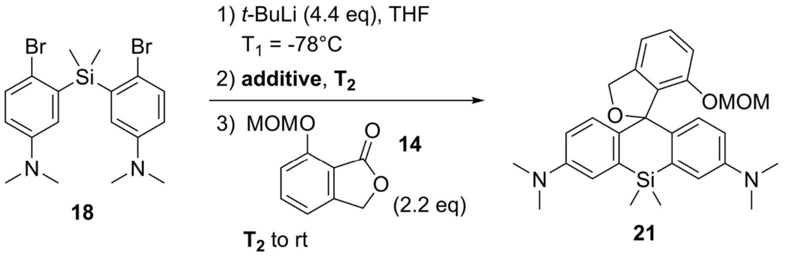

| scale [mmol] | additive | $T_2$ (°C) | yield (%) |
|---|---|---|---|
| 0.11 | $MgBr_2 \cdot OEt_2$ | -10 | <1 |
| 0.11 | - | -20 | 46 |
| 0.16 | - | -20 | 53 |

#### 3.1 Tetrazine derivatives and their precursors

**3-(Methoxymethyl)-6-methyl-1,2,4,5-tetrazine (1)**

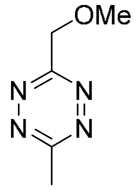

This compound was synthesized following a modified published procedure.[7]

Methoxyacetonitrile (3.82 g, 53.7 mmol) and MeCN (22.06 g, 537 mmol) were dissolved in hydrazine monohydrate (134.5 g, 2.69 mol) and $Ni(OTf)_2$ (9.59 g, 26.9 mmol) was added. The solution was stirred at 60 °C for 18 h and the color of the solution turned from violet to black over time. The solution was then cooled to rt and extracted with DCM (15 x 100 mL). The combined organic layers were washed with brine (300 mL), dried over $Na_2SO_4$ and carefully concentrated under reduced pressure. The concentrated solution was stirred at rt for 4 h while oxygen was bubbled through. Subsequently, it was adsorbed on silica and purification via flash column chromatography ($SiO_2$, *n*-pentane/$Et_2O$, 10:1 to 5:1) gave the title product as a red liquid (2.3 g, 16.6 mmol, 31 %).

$^1H$ NMR (300 MHz, $CDCl_3$) δ 5.04 (s, 2H), 3.60 (s, 3H), 3.09 (s, 3H). $^{13}C$ NMR (75 MHz, $CDCl_3$) δ 168.7, 166.3, 72.3, 59.8, 21.4. HRMS ($ESI^+$) m/z 163.0590 calcd for $[C_5H_8N_4NaO]^+$ ($M+Na^+$), 163.0591 found.

<u>Take note</u>: Title product is volatile. One can consider using e.g. EtOAc/cyclohexane as eluent instead of *n*-pentane/diethyl ether. It is of advantage to concentrate the eluents from chromatography as far as possible, and filtrate over a small silica plug after diluting with *n*-pentane. The adsorbed pink tetrazine can then be released from the silica filter with $Et_2O$ and, after concentration under vacuum, isolated without significant loss.

**(6-Methyl-1,2,4,5-tetrazin-3-yl)methanol (2)**

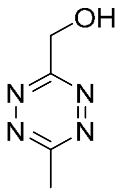

This compound was synthesized following a modified published procedure.[7]

**1** (50 mg, 357 µmol) was dissolved in DCM (5 mL) and cooled to -78 °C. $BBr_3$ (1.42 mL, 1.42 mmol) was added dropwise as 1 M solution in DCM. The solution was stirred at -78 °C for 1 h and then warmed to rt over 30 min. The solution turned from red to orange. Methanol (5 mL) was added dropwise and the reaction mixture turned from a yellow emulsion to a red solution. Then water (1 mL) was added and the aqueous layer was extracted with DCM (5 x 3 mL). Combined organic layers were dried over $Na_2SO_4$ and carefully concentrated under reduced pressure. Purification via column chromatography ($SiO_2$, 5:1 *n*-pentane/$Et_2O$) gave the title product as red crystals (78 mg, 300 µmol, 84 %).

$^1$H NMR (300 MHz, $CDCl_3$) δ 5.27 (d, $J$ = 6.3 Hz, 2H), 3.11 (s*, 3H), 3.01 (t, $J$ = 6.2 Hz, 1H). $^{13}$C NMR (75 MHz, $CDCl_3$) δ 169.0, 167.5, 62.8, 21.4. HRMS (ESI$^+$) m/z 149.0434 calcd for $[C_4H_6N_4NaO]^+$ (M+Na$^+$), 149.0430 found.

(* $^7J$ coupling to C$H_2$ can be observed, compare data from Baalmann *et al.*: $^1$H NMR (300 MHz, $CDCl_3$) δ 5.27 (d, $J$ = 5.9 Hz, 2H), 3.11 (t, $J$ = 0.6 Hz, 3H), 3.03 (t, $J$ = 6.3 Hz, 1H))[7]

**3-(Bromomethyl)-6-methyl-1,2,4,5-tetrazine (3)**

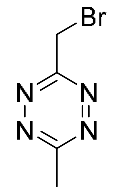

$PPh_3$ (749 mg, 2.85 mmol) was dissolved in DCM (5 mL) and cooled to 0 °C. $CBr_4$ (631 mg, 1.90 mmol) was diluted in DCM (1 mL) and added dropwise and the mixture was stirred for 30 min. Then **2** (120 mg, 952 µmol) was dissolved in DCM (1 mL) and added dropwise. The ice bath was removed after complete addition and the reaction mixture was stirred for 2 h at rt. Subsequently it was adsorbed on silica and purification by flash chromatography ($SiO_2$, 0-10% cyclohexane/EtOAc) yielded the title product as a red liquid (143 mg, 0.757 mmol, 79%).

$^1H$ NMR (300 MHz, $CDCl_3$) δ 4.94 (s, 2H), 3.11 (s, 3H). $^{13}C$ NMR (126 MHz, $CDCl_3$) δ 168.0, 167.5, 27.6, 21.4. HRMS ($EI^+$) m/z 187.9689 calcd for $[C_4H_5BrN_4]^+$ ($M^+$), 187.9689 found.

Take note: Title product is volatile. It is of advantage to concentrate the eluents from chromatography as far as possible, and filtrate over a small silica plug after diluting with *n*-pentane. The adsorbed pink tetrazine can then be released from the silica pad with $Et_2O$ and further concentrated to afford the neat product without substantial loss.

**General Procedure (1) for the preparation of tetrazinyl benzaldehydes 4-6**

This procedure for **4** is representative: 2-Hydroxybenzaldehyde (47 mg, 0.38 mmol), 15-crown-5 (69 µL, 0.35 mmol) were dissolved in anhydrous acetonitrile (13 mL) at rt in a baked-out Schlenk flask equipped with 3 Å mol. sieves. Then NaH (60% dispersion in mineral oil, 14 mg, 0.35 mmol) was added and the mixture was stirred until hydrogen evolution had ceased. In a second flask, **3** (72 mg, 0.38 mmol) was dissolved in anhydrous acetonitrile (2.5 mL) and cooled to 0 °C. Subsequently the sodium phenolate mixture was added to second flask over a period of 10 min. The reaction mixture was stirred for 17 h at rt. Then sat. aq. $NH_4Cl$ solution (15 mL) and DCM (15 mL) were added the aq. phase was extracted with DCM (5x10 mL). The combined organic layers were dried over $Na_2SO_4$ and concentrated under reduced pressure. The crude residue was purified by flash chromatography ($SiO_2$, 10-30% EtOAc / cyclohexane) affording **4** as pink crystals (45 mg, 56%).

**2-((6-Methyl-1,2,4,5-tetrazin-3-yl)methoxy)benzaldehyde (4)**

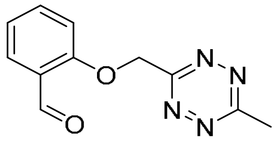

The title compound (45 mg, 56%, red crystals) was prepared according to general procedure (1) from **3** (72 mg, 0.38 mmol) and 2-hydroxybenzaldehyde (47 mg, 0.38 mmol).

$^1H$ NMR (500 MHz, $CDCl_3$) δ 10.55 (d, *J* = 0.8 Hz, 1H), 7.89 (dd, *J* = 7.7, 1.8 Hz, 1H), 7.57 (ddd, *J* = 8.4, 7.3, 1.8 Hz, 1H), 7.17 (dd, *J* = 8.4, 0.9 Hz, 1H), 7.14 – 7.10 (m, 1H), 5.79 (s, 2H), 3.13 (s, 3H). $^{13}C$ NMR (126 MHz, $CDCl_3$) δ 189.4, 169.1, 165.3, 160.2, 136.0, 129.0, 125.8, 122.3, 113.1, 68.3, 21.5. HRMS ($ESI^+$) m/z 253.0696 calcd for $[C_{11}H_{10}N_4NaO_2]^+$ $(M+Na)^+$, 253.0702 found.

**3-((6-Methyl-1,2,4,5-tetrazin-3-yl)methoxy)benzaldehyde (5)**

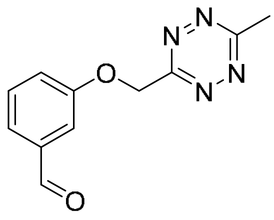

The title compound (31 mg, 41%, red solid) was prepared according to general procedure (1) from **3** (69 mg, 0.37 mmol) and 3-hydroxybenzaldehyde (45 mg, 0.37 mmol).

$^{1}H$ NMR (300 MHz, $CDCl_3$) δ 9.99 (s, 1H), 7.58 – 7.46 (m, 3H), 7.34 (ddd, *J* = 7.8, 2.7, 1.6 Hz, 1H), 5.72 (s, 2H), 3.12 (s, 3H). $^{13}C$ NMR (75 MHz, $CDCl_3$) δ 191.8, 169.0, 165.4, 158.7, 138.1, 130.6, 124.7, 122.3, 113.5, 68.0, 21.5. HRMS ($ESI^+$) m/z 253.0696 calcd for $[C_{11}H_{10}N_4NaO_2]^+$ $(M+Na)^+$, 253.0714 found.

**4-((6-Methyl-1,2,4,5-tetrazin-3-yl)methoxy)benzaldehyde (6)**

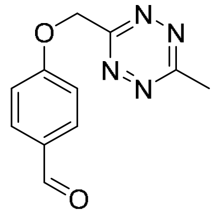

The title compound (50 mg, 64%, red crystals) was prepared according to general procedure (1) from **3** (71 mg, 0.38 mmol) and 4-hydroxybenzaldehyde (46 mg, 0.38 mmol).

$^{1}H$ NMR (300 MHz, $CDCl_3$) δ 9.91 (s, 1H), 7.87 (d, *J* = 8.8 Hz, 2H), 7.17 (d, *J* = 8.7 Hz, 2H), 5.74 (s, 2H), 3.13 (s, 3H). $^{13}C$ NMR (75 MHz, $CDCl_3$) δ 190.8, 169.1, 165.2, 162.8, 132.2, 131.1, 115.4, 67.9, 21.5. HRMS ($ESI^+$) m/z 253.0696 calcd for $[C_{11}H_{10}N_4NaO_2]^+$ $(M+Na)^+$, 253.0698 found.

**tert-Butyl 4-bromo-3-hydroxybenzoate (7)**

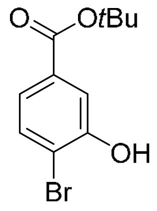

4-Bromo-3-hydroxybenzoic acid (2.17 g, 10.0 mmol) was suspended in anhydrous toluene (60 mL) under an argon atmosphere and heated to 80 °C. Then *N*,*N*-dimethylformamide di-*tert*-butyl acetal (4.80 mL, 20.0 mmol) was added dropwise over 15 min. After stirring for 45 min, the mixture was cooled down to rt, diluted with DCM (10 mL) and washed with sat. aq. $NaHCO_3$ (2x10 mL) and brine (10 mL). The organic layer was dried over $Na_2SO_4$ and cocentrated under reduced pressure. Subsequently, the crude residue was purified by flash chromatography ($SiO_2$, DCM) to afford the disubstituted side product *tert*-butyl 4-bromo-3-(*tert*-butoxy)benzoate as transparent oil (157 mg, 4.8%) as well as the title product as a white solid (1.46 g, 54%).

$^1$H NMR (500 MHz, $CDCl_3$) δ 7.63 (d, *J* = 2.0 Hz, 1H), 7.51 (d, *J* = 8.4 Hz, 1H), 7.42 (dd, *J* = 8.3, 2.0 Hz, 1H), 5.79 (s, 1H), 1.58 (s, 9H). $^{13}$C NMR (126 MHz, $CDCl_3$) δ 165.0, 152.4, 133.3, 132.1, 122.7, 117.1, 115.1, 81.8, 28.3. HRMS (ESI$^+$) m/z 294.9940 calcd for $[C_{11}H_{13}BrNaO_3]^+$ (M+Na$^+$), 294.9936 found.

**tert-Butyl 3-(allyloxy)-4-bromobenzoate (8)**

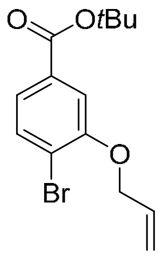

Allyl bromide (1.85 mL, 21.4 mmol) and $K_2CO_3$ (2.96 g, 21.4 mmol) were added in sequence to a stirred solution of **7** (1.95 g, 7.14 mmol) in anhydrous acetone (70 mL) at rt. The solution was refluxed at 65 °C for 3 h. After cooling down to rt, the mixture was filtered and concentrated under reduced pressure. Purification by flash chromatography ($SiO_2$, DCM) afforded the title product as colorless oil (2.22 g, 99%).

$^1$H NMR (500 MHz, $CDCl_3$) δ 7.58 (d, *J* = 8.2 Hz, 1H), 7.51 (d, *J* = 1.8 Hz, 1H), 7.44 (dd, *J* = 8.2, 1.8 Hz, 1H), 6.08 (ddt, *J* = 17.3, 10.4, 5.1 Hz, 1H), 5.51 (dd, *J* = 17.3, 1.6 Hz, 1H), 5.34 (dd, *J* =

10.6, 1.4 Hz, 1H), 4.68 – 4.65 (m, 2H), 1.59 (s, 9H). $^{13}C$ NMR (75 MHz, $CDCl_3$) δ 165.2, 154.9, 133.2, 132.5, 132.4, 123.0, 118.2, 117.5, 114.0, 81.7, 69.9, 28.3. HRMS ($ESI^+$) m/z 335.0253 calcd for $[C_{11}H_{13}BrNaO_3]^+$ ($M+Na^+$), 335.0265 found.

***tert*-Butyl 3-(allyloxy)-4-formylbenzoate (9)**

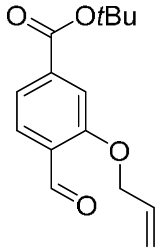

**8** (1.19 g, 3.80 mmol) was dissolved in anhydrous THF (25 mL) under argon and cooled down to -100 °C. Subsequently *n*-butyllithium (2.5 M in hexanes, 1.75 mL, 4.37 mmol) was added dropwise and the mixture was stirred for 30 min. Then dimethylformamide (1.46 mL, 19.0 mmol) was added dropwise and after 30 min of stirring, the reaction mixture was allowed to heat up to -40 °C and sat. aq. $NH_4Cl$ (10 mL) as well as water (10 mL) were added. After warming to rt, phases were separated and the aq. phase was extracted with EtOAc (3x40 mL). The combined organic layers were washed with aq. LiCl (5%, w/v, 2x40 mL), brine (40 mL) and dried over $MgSO_4$. Removal of volatiles under reduced pressure afforded the title product as off-white solid (0.969 g, 97%). The compound was used in the next step without further purification.

$^{1}H$ NMR (500 MHz, $CDCl_3$) δ 10.56 (s, 1H), 7.88 – 7.84 (m, 1H), 7.64 – 7.59 (m, 2H), 6.09 (ddt, *J* = 17.3, 10.5, 5.2 Hz, 1H), 5.48 (dd, *J* = 17.2, 1.5 Hz, 1H), 5.37 (dd, *J* = 10.6, 1.4 Hz, 1H), 4.75 – 4.70 (m, 2H), 1.61 (s, 9H). $^{13}C$ NMR (75 MHz, $CDCl_3$) δ 189.6, 164.8, 160.6, 138.5, 132.2, 128.4, 127.6, 121.7, 118.6, 114.0, 82.2, 69.6, 28.2. HRMS ($ESI^+$) m/z 285.1097 calcd for $[C_{15}H_{18}NaO_4]^+$ ($M+Na^+$), 285.1098 found.

***tert*-Butyl 4-formyl-3-hydroxybenzoate (10)**

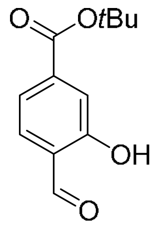

Tetrakis(triphenylphosphine)palladium(0) (17.2 mg, 14.9 µmol) was added to a solution of **9** (390 mg, 1.49 mmol) in anhydrous and degassed methanol (15 mL) at rt. After 5 min, $K_2CO_3$ (616 mg, 4.46 mmol) was added and the mixture was stirred for 2 h. Then it was filtered and concentrated under reduced pressure. The residue was taken up in DCM (20 mL) and sat. aq. $NH_4Cl$ (10 mL). Subsequently, the aq. phase was extracted with DCM (3x10 mL), the combined organic phases were washed with brine (20 mL) and dried over $Na_2SO_4$. The crude mixture was concentrated under reduced pressure and purified by flash chromatography ($SiO_2$, 10:1 cyclohexane, EtOAc) to afford the title product as off-white solid (196 mg, 59%).

$^{1}H$ NMR (300 MHz, $CDCl_3$) δ 10.94 (s, 1H), 9.98 (s, 1H), 7.63 – 7.58 (m, 3H), 1.60 (s, 9H). $^{13}C$ NMR (75 MHz, $CDCl_3$) δ 196.6, 164.3, 161.3, 139.5, 133.6, 122.7, 120.5, 119.0, 82.3, 28.2. HRMS (ESI$^+$) m/z 245.0784 calcd for $[C_{12}H_{14}NaO_4]^+$ (M+Na$^+$), 245.0789 found.

***tert*-Butyl 4-formyl-3-((6-methyl-1,2,4,5-tetrazin-3-yl)methoxy)benzoate (11)**

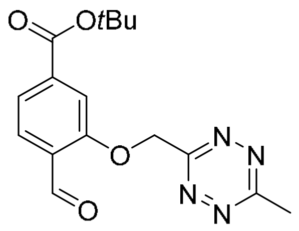

**10** (136 mg, 0.614 mmol) and **3** (122 mg, 0.644 mmol) were dissolved in anhydrous THF (6 mL) under argon and cooled to 0 °C. After the addition of $Cs_2CO_3$ (300 mg, 0.920 mmol), the ice bath was removed and the mixture was stirred for 4 h at rt. Subsequently, sat. aq. $NH_4Cl$ (6 mL), water (2 mL) and DCM (10 mL) were added. The aq. phase was extracted with DCM (2x10 mL). The combined organic phases were washed with brine (10 mL), dried over $Na_2SO_4$ and the solvent was removed under reduced pressure. Purification of the crude residue by flash chromatography ($SiO_2$, 5-15% EtOAc / cyclohexane) gave the title product as pink solid (139 mg, 69%).

$^{1}H$ NMR (300 MHz, $CDCl_3$) δ 10.56 (d, *J* = 0.8 Hz, 1H), 7.90 (d, *J* = 7.9 Hz, 1H), 7.79 (d, *J* = 1.3 Hz, 1H), 7.69 (ddd, *J* = 8.0, 1.3, 0.8 Hz, 1H), 5.83 (s, 2H), 3.13 (s, 3H), 1.60 (s, 9H). $^{13}C$ NMR (75 MHz, $CDCl_3$) δ 189.1, 169.2, 165.1, 164.4, 159.9, 138.6, 128.8, 128.0, 123.0, 113.9, 82.5, 68.4, 28.2, 21.5. HRMS (ESI$^+$) m/z 353.1220 calcd for $[C_{16}H_{18}N_4NaO_4]^+$ (M+Na$^+$), 353.1217 found.

#### 3.2 Dye precursors

**4-(Methoxymethoxy)isobenzofuran-1,3-dione (12)**

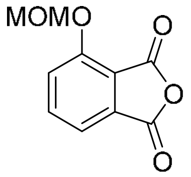

3-Hydroxyphthalic anhydride (0.50 g, 3.05 mmol) was dissolved in anhydrous THF (30 mL) under argon, cooled to 0 °C and *N*,*N*-diisopropylethylamine (0.484 mL, 3.20 mmol) was added. After 5 min, chloromethyl methyl ether[a] (0.278 mL, 3.66 mmol) was added dropwise and the solution was stirred for 48 h while allowing to heat up to rt. Subsequently, the mixture was adsorbed on silica and purified by flash chromatography ($SiO_2$, 20-40% EtOAc / cyclohexane) to afford the title product as white solid (513 mg, 81%).

(Note: Can also be performed in 24 mmol scale with 77% yield.)

$^{1}H$ NMR (300 MHz, $CDCl_3$) δ 7.81 (dd, *J* = 8.4, 7.4 Hz, 1H), 7.63 – 7.56 (m, 2H), 5.42 (s, 2H), 3.54 (s, 3H). $^{13}C$ NMR (75 MHz, $CDCl_3$) δ 162.8, 160.7, 156.1, 138.3, 133.2, 122.4, 118.6, 117.9, 95.1, 57.1. HRMS (ESI$^+$) m/z 231.0264 calcd for $[C_{10}H_8NaO_5]^+$ (M+Na)$^+$, 231.0273 found.

[a] Caution: Chloromethyl methyl ether is a known human carcinogen; handle with extreme care and work in a well-ventilated fume hood.

**(3-(Methoxymethoxy)phenyl)methanol (13)**

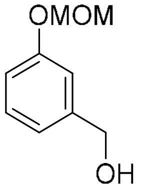

This compound was synthesized following a modified published procedure.[8]

A mixture of NaH (60% mineral oil dispersion, 0.799 g, 20.0 mmol) in anhydrous THF (60 mL) was cooled to 0 °C under an argon atmosphere. Then it was slowly treated with a solution of 3-hydroxybenzyl alcohol (2.48 g, 20.0 mmol) in THF (10 mL) and stirred for 1 h at 0 °C, before the ice bath was removed and the mixture stirred for 30 min at rt. Subsequently, chloromethyl methyl ether[a] (1.67 mL, 22.0 mmol) was added dropwise at 0 °C and the solution was stirred for 1 h while allowing to heat up to rt. Then ice-cooled aq. sat. $NH_4Cl$ (40 mL) and the resulting mixture was extracted with diethyl ether (3x50 mL). The combined organic layers were washed with aq. NaOH (40 mL, 1 M), water (40 mL), brine (40 mL) and dried over $MgSO_4$. The solvent was removed under reduced pressure and purification by flash chromatography ($SiO_2$, 40-50% EtOAc / cyclohexane) afforded the title product as colorless oil (2.14 g, 64%).

$^1$H NMR (300 MHz, $CDCl_3$) δ 7.28 (t, $J$ = 7.8 Hz, 1H), 7.09 – 6.93 (m, 3H), 5.19 (s, 2H), 4.68 (s, 2H), 3.48 (s, 3H). $^{13}$C NMR (75 MHz, $CDCl_3$) δ 157.6, 142.7, 129.8, 120.5, 115.7, 114.8, 94.5, 65.4, 56.2. HRMS (ESI$^+$) m/z 191.0679 calcd for $[C_9H_{12}NaO_3]^+$ (M+Na$^+$), 191.0676 found.

**7-(Methoxymethoxy)isobenzofuran-1(3*H*)-one (14)**

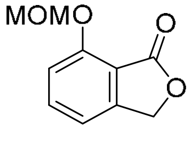

This compound was synthesized following a modified published procedure.[8]

**13** (1.01 g, 6.00 mmol) was dissolved in anhydrous benzene (50 mL) under argon in a dried Schlenk flask. After the dropwise addition of *n*-BuLi (6.00 mL, 15.0 mmol, 2.5 M in hexanes) at rt, the mixture was stirred for 2 h. Gaseous carbon dioxide was generated from dry ice in an additional two-necked flask and bubbled through the *n*-BuLi-treated solution at 0 °C. A wash bottle filled with sulfuric acid was interposed between the gas inlet and the flask filled with dry ice. After 1 h, the ice bath and the gas inlet were removed. Subsequently, chunks of dry ice were added to the mixture and it was stirred for 18 h. Volatiles were then removed under reduced pressure and the solid residue was dissolved in glacial acetic acid (40 mL) and stirred for 2 h at rt. The acid was removed under reduced pressure and the remaining yellow residue dissolved in diethyl ether (60 mL) and stirred for 30 min at rt. The newly-formed white precipitate was filtered off and the filtrate was washed with aq. $Na_2CO_3$ (20 mL, 5% w/v), brine (20 mL), dried over $Na_2SO_4$ and concentrated under reduced pressure. Purification of the residue by flash chromatography ($SiO_2$, 0-10% EtOAc/DCM) gave the title product as yellow crystals (0.732 g, 62%).

$^1H$ NMR (300 MHz, $CDCl_3$) δ 7.59 (t, *J* = 7.9 Hz, 1H), 7.20 (d, *J* = 8.3 Hz, 1H), 7.07 (d, *J* = 7.5 Hz, 1H), 5.39 (s, 2H), 5.24 (s, 2H), 3.54 (s, 3H). $^{13}C$ NMR (75 MHz, $CDCl_3$) δ 169.0, 156.5, 149.3, 136.2, 115.0, 114.8, 114.4, 94.9, 68.8, 56.7. HRMS ($ESI^+$) m/z 217.0471 calcd for $[C_{10}H_{10}NaO_4]^+$ ($M+Na^+$), 217.0481 found.

**3,3'-Oxybis(*N*,*N*-dimethylaniline) (15)**

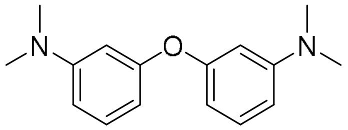

This compound was synthesized following a modified published procedure.[9,10]

3-Bromo-*N*,*N*-dimethylaniline (10.0 g, 50.0 mmol) was added to a mixture of 3-dimethylaminophenol (8.23 g, 60.0 mmol), CuI (925 mg, 5.00 mmol), anhydrous $K_3PO_4$ (21.2 g, 100 mmol) and picolinic acid (1.23 g, 10.0 mmol) in anhydrous DMSO (95 mL) under argon. The mixture was stirred at 90 °C for 40 h. After cooling down to rt, it was filtered and washed with EtOAc (0.25 L). The filtrate was further diluted with EtOAc (0.5 L) and washed with water (2 x 0.75 L). The combined aq. phases were diluted with brine and, adjusted to pH 9 with sat. aq. $Na_2CO_3$ and extracted with EtOAc (3x0.5 L). The combined organic layers were washed with brine (3 x 0.2 L), dried over $Na_2SO_4$ and concentrated under reduced pressure. Purification by flash chromatography ($SiO_2$, 0-10% EtOAc / cyclohexane) afforded the title product as white crystals (8.89 g, 69%).

$^1H$ NMR (300 MHz, $CDCl_3$) δ 7.19 – 7.10 (m, 2H), 6.51 – 6.42 (m, 4H), 6.35 (ddd, *J* = 8.0, 2.1, 1.0 Hz, 2H), 2.93 (s, 12H). $^{13}C$ NMR (75 MHz, $CDCl_3$) δ 158.5, 152.2, 129.9, 107.5, 106.9, 103.5, 40.7. HRMS ($ESI^+$) m/z 257.1648 calcd for $[C_{16}H_{21}N_2O]^+$ ($M+H^+$), 257.1665 found.

**3,3'-Oxybis(4-bromo-*N*,*N*-dimethylaniline) (16)**

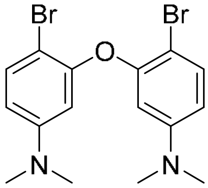

This compound was synthesized following a modified published procedure.[6]

**15** (0.200 g, 0.78 mmol) was dissolved in dimethylformamide (10 mL) at rt and *N*-bromosuccinimide (0.278 g, 1.56 mmol) was added in portions over 2 min. The reaction was stirred for 72 h at rt. After the addition of water, the mixture was extracted with EtOAc, the organic layers dried over $Na_2SO_4$ and the solvent removed under reduced pressure. The crude residue was purified by flash chromatography ($SiO_2$, 0-30% EtOAc / cyclohexane) to afford the title product as off-white solid (240 mg, 74%).

$^1H$ NMR (500 MHz, $CDCl_3$) δ 7.39 (d, *J* = 8.9 Hz, 2H), 6.37 (dd, *J* = 8.9, 2.9 Hz, 2H), 6.22 (d, *J* = 2.9 Hz, 2H), 2.86 (s, 12H). $^{13}C$ NMR (126 MHz, $CDCl_3$) δ 153.8, 151.2, 133.5, 109.3, 104.0, 100.0, 40.6. HRMS ($ESI^+$) m/z 412.9859 calcd for $[C_{16}H_{19}Br_2N_2O]^+$ ($M+H^+$), 412.9864 found.

**3,3'-(Dimethylsilanediyl)bis(*N*,*N*-dimethylaniline) (17)**

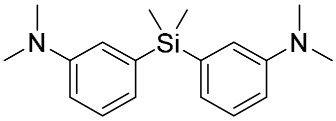

This compound was synthesized following a modified published procedure.[9]

3-Bromo-*N*,*N*-dimethylaniline (3.66 mL, 25.0 mmol) was dissolved in anhydrous THF (50 mL) under argon and cooled to -78°C. Then *s*-BuLi (18.7 mL, 26.2 mmol, 1.4 M in cyclohexane) was added dropwise and the mixture was stirred for 1 h. A solution of dichlorodimethylsilane (1.83 mL, 15.0 mmol) in anhydrous THF (8 mL) was prepared and added dropwise. The mixture was subsequently stirred for 18 h while allowing to heat up to rt. After the addition of sat. aq. $NH_4Cl$ (20 mL) and water (30 mL), the mixture was extracted with EtOAc (3x50 mL), the combined organic layers were dried over $Na_2SO_4$ and concentrated under reduced pressure. The crude residue was purified by flash chromatography (5% EtOAc / cyclohexane) to afford the title product as colorless oil (2.53 g, 68%).

$^1H$ NMR (300 MHz, $CDCl_3$) δ 7.26 (dd, *J* = 8.3, 7.1 Hz, 2H), 7.00 – 6.90 (m, 4H), 6.79 (dd, *J* = 8.2, 2.3 Hz, 2H), 2.95 (s, 12H), 0.56 (s, 6H). $^{13}C$ NMR (75 MHz, $CDCl_3$) δ 150.0, 139.1, 128.6, 122.9,

118.5, 113.7, 40.8, -2.0. HRMS (ESI+) m/z 299.1938 calcd for $[C_{18}H_{27}N_2Si]^+$ ($M+H^+$), 299.1962 found.

**3,3'-(Dimethylsilanediyl)bis(4-bromo-*N*,*N*-dimethylaniline) (18)**

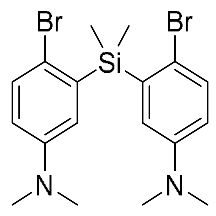

This compound was synthesized following a modified published procedure.[6]

**17** (2.59 g, 8.69 mmol) was dissolved in DMF (50 mL) at rt and *N*-bromosuccinimide (3.09 g, 17.4 mmol) was added in portions over 10 min. The reaction mixture was stirred for 72 h while shielding the flask from light. After subsequent concentration under reduced pressure, the residue was extracted with EtOAc. The combined organic phases were washed with water, dried over $MgSO_4$ and concentrated under reduced pressure. Purification by flash chromatography ($SiO_2$, 0-15% diethylether / cyclohexane) afforded the title product as white solid (3.04 g, 77%).

$^1$H NMR (300 MHz, $CDCl_3$) δ 7.34 (d, *J* = 8.7 Hz, 2H), 6.83 (d, *J* = 3.2 Hz, 2H), 6.59 (dd, *J* = 8.8, 3.2 Hz, 2H), 2.87 (s, 12H), 0.75 (s, 6H). $^{13}$C NMR (75 MHz, $CDCl_3$) δ 149.0, 138.9, 133.1, 121.9, 117.0, 115.4, 40.7, -0.8. HRMS (ESI+) m/z 455.0148 calcd for $[C_{18}H_{25}Br_2N_2Si]^+$ ($M+H^+$), 455.0151 found.

#### 3.3 Cell permeable probes

**3',6'-Bis(dimethylamino)-[7 or 4]-(methoxymethoxy)-3*H*-spiro[isobenzofuran-1,9'-xanthen]-3-one (19a, 19b)**

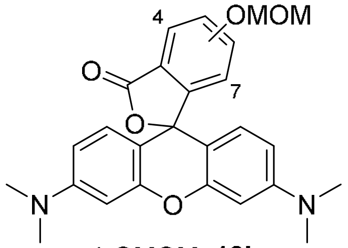

4-OMOM, **19b**
7-OMOM, **19a**

A procedure from the literature was adapted for the synthesis of this compound.[6]

**16** (0.5 g, 1.21 mmol) was dissolved in anhydrous and degassed THF (30 mL) under argon and cooled to -78 °C. *tert*-Butyllithium (3.12 mL, 5.31 mmol, 1.7 M in pentane) was added dropwise and the mixture was stirred for 1 h. Next, it was warmed to -35 °C and a solution of **12** (553 mg, 2.66 mmol) in anhydrous and degassed THF (30 mL) was added over 45 min using a syringe pump. The reaction was stirred for 18 h while allowing to heat to rt. Then sat. aq. $NH_4Cl$ (25 mL) and water (25 mL) were added and the mixture was extracted with DCM (3x30 mL). The combined organic layers were dried over $Na_2SO_4$ and concentrated under reduced pressure. The crude residue was purified by flash chromatography ($SiO_2$, 0-9% 2 M $NH_3$ in MeOH / DCM) affording the 4-OMOM isomer **19b** as pale pink crystals (150 mg, 28%) and the 7-OMOM isomer **19a** as purple solid (34.0 mg, 6.3%).

**19a**, *Spirolactone*: $^1H$ NMR (500 MHz, $CDCl_3$) δ 7.64 (d, *J* = 7.6 Hz, 1H), 7.51 (t, *J* = 7.8 Hz, 1H), 7.21 (d, *J* = 8.1 Hz, 1H), 6.72 (d, *J* = 8.8 Hz, 2H), 6.48 (d, *J* = 2.5 Hz, 2H), 6.38 (dd, *J* = 8.8, 2.6 Hz, 2H), 4.86 (s, 2H), 2.96 (s, 12H), 2.87 (s, 3H). $^{13}C$ NMR (126 MHz, $CDCl_3$) δ 170.0, 152.8, 152.1, 151.3, 141.2, 131.3, 129.1, 128.0, 119.5, 117.8, 108.6, 106.7, 98.9, 93.3, 84.1*, 55.7, 40.5. (* weak signal, confirmed via gHBMC) *Quinoid*: $^1H$ NMR (500 MHz, $CD_3OD$) δ 7.71 (dd, *J* = 7.8, 1.0 Hz, 1H), 7.60 (t, *J* = 8.0 Hz, 1H), 7.40 (dd, *J* = 8.3, 1.0 Hz, 1H), 7.28 (d, *J* = 9.4 Hz, 2H), 6.98 (dd, *J* = 9.4, 2.5 Hz, 2H), 6.88 (d, *J* = 2.5 Hz, 2H), 5.00 (s, 2H), 3.26 (s, 12H), 3.08 (s, 3H). HRMS ($ESI^+$) m/z 447.1914 calcd for $[C_{26}H_{27}N_2O_5]^+$ ($M+H^+$), 447.1919 found.

**19b**, $^1H$ NMR (500 MHz, $CDCl_3$) δ 7.52 (t, *J* = 7.9 Hz, 1H), 7.23 (d, *J* = 8.2 Hz, 1H), 6.74 (d, *J* = 7.5 Hz, 1H), 6.69 (d, *J* = 8.8 Hz, 2H), 6.47 (d, *J* = 2.6 Hz, 2H), 6.40 (dd, *J* = 8.8, 2.6 Hz, 2H), 5.46 (s,

2H), 3.62 (s, 3H), 2.97 (s, 12H). $^{13}C$ NMR (126 MHz, $CDCl_3$) δ 167.7, 156.5, 155.5, 152.9, 152.1, 136.5, 128.8, 117.2, 115.8, 114.9, 108.7, 107.2, 98.7, 95.3, 83.3, 56.9, 40.4. HRMS ($ESI^+$) m/z 447.1914 calcd for $[C_{26}H_{27}N_2O_5]^+$ ($M+H^+$), 447.1915 found.

**2-(6-(Dimethylamino)-3-(dimethyliminio)-3*H*-xanthen-9-yl)-3-hydroxybenzoate (22)**

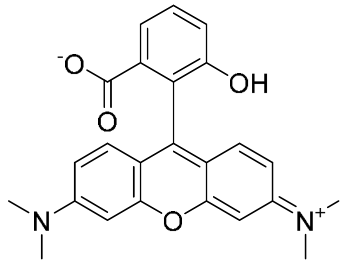

A solution of **19a** (27.3 mg, 61.1 µmol) in DCM (4 mL) was cooled to 0 °C and TFA (0.4 mL, 6.11 mmol) was added dropwise. The ice bath was removed and the mixture was stirred for 18 h at rt. After coevaporation with toluene (2 mL) under reduced pressure, the crude residue was purified by flash chromatography ($SiO_2$, 0-8% 2 M $NH_3$ in MeOH / DCM) to afford the title product as pink solid with metallic shine (15.1 mg, 61%).

$^1H$ NMR (300 MHz, $CD_3OD$) δ 7.81 (dd, *J* = 7.8, 0.9 Hz, 1H), 7.60 (t, *J* = 8.0 Hz, 1H), 7.28 (dd, *J* = 8.2, 0.9 Hz, 1H), 7.25 (d, *J* = 9.4 Hz, 2H), 7.04 (dd, *J* = 9.4, 2.4 Hz, 2H), 6.91 (d, *J* = 2.4 Hz, 2H), 3.28 (s, 12H). $^{13}C$ NMR (75 MHz, $CD_3OD$) δ 168.3, 160.8, 159.1, 158.9, 156.4, 133.4, 132.4, 132.0, 123.3, 121.5, 121.1, 115.5, 115.2, 97.2, 40.9. (TFA was added to the NMR sample prior measuring) HRMS ($ESI^+$) m/z 403.1652 calcd for $[C_{24}H_{23}N_2O_4]^+$ ($M+H^+$), 403.1669 found.

**HD555**

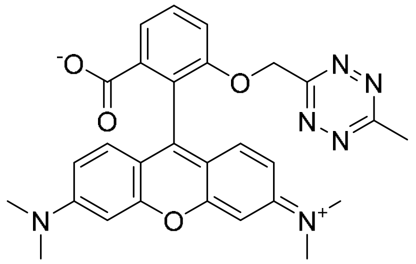

**22** (7.10 mg, 17.4 µmol) was dissolved in anhydrous DMF (0.4 mL) in a Schlenk tube charged with 4 Å mol. sieve under argon and at rt. $Cs_2CO_3$ (17.2 mg, 52.9 µmol) was added and it was stirred for 15 min. The marginally pink coloration of the solution indicated predominant presence of the spirolactone isomer. Separately, **3** (10.0 mg, 52.9 µmol) was dissolved in anhydrous DMF (0.2 mL) and added to the cooled (0 °C) first solution. While allowing to heat to rt, it was subsequently stirred for 1 h. Next, sat. aq. $NH_4Cl$ (1 mL), water (0.5 mL) and brine (3 mL) were added in sequence and the mixture was extracted with DCM (4x2 mL). The combined organic phases were dried over $Na_2SO_4$ and concentrated under reduced pressure. Purification of the crude residue by flash chromatography ($SiO_2$, 0-12% MeOH / DCM) afforded the title product as pink solid (4.7 mg, 52%).

$^1$H NMR (500 MHz, $CD_3OD$) δ 7.98 (dd, *J* = 7.7, 0.9 Hz, 1H), 7.77 (t, *J* = 8.1 Hz, 1H), 7.71 (d, *J* = 7.8 Hz, 1H), 7.20 (d, *J* = 9.5 Hz, 2H), 7.01 (dd, *J* = 9.5, 2.5 Hz, 2H), 6.88 (d, *J* = 2.5 Hz, 1H), 5.65 (s, 2H), 3.29 (s, 12H), 2.93 (s, 3H). $^{13}$C NMR (126 MHz, $CD_3OD$) δ 169.9, 167.7, 166.2, 159.3, 159.0, 158.9, 156.9, 133.7, 132.8, 132.0, 125.5, 124.3, 119.0, 115.4, 115.3, 97.2, 69.6, 40.9, 21.2 (TFA was added to the NMR sample prior measuring). HRMS (ESI$^+$) m/z 533.1908 calcd for $[C_{28}H_{26}N_6NaO_4]^+$ (M+Na$^+$), 511.1926 found.

**3,7-Bis(dimethylamino)-[7' or 4']-(methoxymethoxy)-5,5-dimethyl-3'*H*,5*H*-spiro[dibenzo-[b,e]siline-10,1'-isobenzofuran]-3'-one (20a, 20b)**

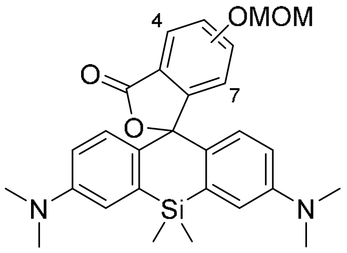

4-OMOM, **20b**
7-OMOM, **20a**

A procedure from the literature was adapted for the synthesis of this compound.[6]

**18** (0.5 g, 1.10 mmol) was dissolved in anhydrous and degassed THF (25 mL) under argon and cooled to -78 °C. *tert*-Butyllithium (2.84 mL, 4.82 mmol, 1.7 M in pentane) was added dropwise and the mixture was stirred for 1 h. Next, it was warmed to -20 °C and a solution of **12** (502 mg, 2.41 mmol) in anhydrous and degassed THF (25 mL) was added over 40 min using a syringe pump. The reaction was stirred for 18 h while allowing to heat to rt. Then sat. aq. $NH_4Cl$ (60 mL) and water (30 mL) were added and the mixture was extracted with EtOAc (4x40 mL). The combined organic layers were washed with brine (50 mL), dried over $MgSO_4$ and concentrated under reduced pressure. The crude residue was purified by flash chromatography ($SiO_2$, 0-40% EtOAc / cyclohexane; const. 20% DCM) affording the 4-OMOM isomer **20b** as off-white solid (156 mg, 29%) and the 7-OMOM isomer **20a** as yellow solid (67 mg, 13%).

**20a:** $^1$H NMR (500 MHz, $CDCl_3$) δ 7.64 (d, *J* = 7.2 Hz, 1H), 7.56 (t, *J* = 7.8 Hz, 1H), 7.38 (d, *J* = 8.0 Hz, 1H), 6.97 (d, *J* = 2.8 Hz, 2H), 6.74 (d, *J* = 8.8 Hz, 2H), 6.53 (dd, *J* = 8.9, 2.8 Hz, 2H), 5.02 (s, 2H), 3.07 (s, 3H), 2.95 (s, 12H), 0.61 (s, 3H), 0.61 (s, 3H). $^{13}$C NMR (126 MHz, $CDCl_3$) δ 170.1, 152.1, 149.5, 141.3, 137.6, 131.2, 131.1, 130.1, 128.1, 118.7, 118.5, 116.7, 113.2, 93.3, 91.9, 56.2, 40.5, 0.9, -2.1. HRMS (ESI$^+$) m/z 489.2204 calcd for $[C_{28}H_{33}N_2O_4Si]^+$ (M+H$^+$), 489.2209 found.

**20b:** $^1$H NMR (500 MHz, $CD_3CN$) δ 7.56 (dd, *J* = 7.9 Hz, 1H), 7.16 (d, *J* = 8.2 Hz, 1H), 7.02 (d, *J* = 2.8 Hz, 2H), 6.82 (d, *J* = 8.9 Hz, 2H), 6.70 (d, *J* = 7.6 Hz, 1H), 6.66 (dd, *J* = 9.0, 2.9 Hz, 2H), 5.39 (s, 2H), 3.52 (s, 3H), 2.94 (s, 12H), 0.61 (s, 3H), 0.52 (s, 3H). $^{13}$C NMR (126 MHz, $CD_3CN$) δ 169.0, 159.2, 156.9, 150.6, 137.2, 136.7, 132.5, 132.5, 128.9, 117.7, 117.5, 115.3, 114.9, 95.9, 90.3, 57.0, 40.5, 0.1, -0.9. HRMS (ESI$^+$) m/z 489.2204 calcd for $[C_{28}H_{33}N_2O_4Si]^+$ (M+H$^+$), 489.2187 found.

**3,7-Bis(dimethylamino)-7'-hydroxy-5,5-dimethyl-3'*H*,5*H*-spiro[dibenzo[b,e]siline-10,1'-isobenzofuran]-3'-one (23)**

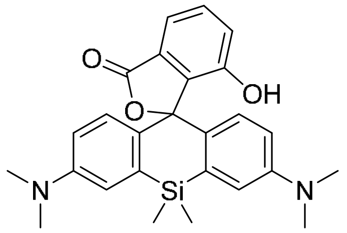

A solution of **20a** (28.0 mg, 57.3 µmol) in DCM (2 mL) was cooled to 0 °C and TFA (0.38 mL, 5.73 mmol) was added dropwise. The ice bath was removed and the mixture was stirred for 18 h at rt. After coevaporation with toluene (3 mL) under reduced pressure, the crude residue was subjected to flash chromatography ($SiO_2$, 0-6% MeOH / DCM, const. 10% EtOAc, const. 1% AcOH) to afford the purified product as blue solid (zwitterionic; poor solubility in apolar solvents and poor NMR spectra). It was further dissolved in DCM (3 mL) and washed with sat. aq. $NaHCO_3$ (1.5 mL), then with brine (1.5 mL) and dried over $Na_2SO_4$. After removal of the solvent under reduced pressure, the title product was afforded as pale green solid (19.0 mg, 75%).

*Spirolactone:* $^1$H NMR (500 MHz, $CDCl_3$) δ 7.62 (d, *J* = 7.3 Hz, 1H), 7.57 (t, *J* = 7.7 Hz, 1H), 7.25 (d, *J* = 7.8 Hz, 1H), 7.03 (d, *J* = 2.8 Hz, 2H), 6.78 (d, *J* = 8.8 Hz, 2H), 6.53 (dd, *J* = 8.8, 2.8 Hz, 2H), 2.99 (s, 12H), 0.66 (s, 3H), 0.63 (s, 3H). $^{13}$C NMR (126 MHz, $CDCl_3$) δ 169.8, 151.3, 149.9, 139.3,

137.4, 131.6, 130.1, 129.6, 127.7, 121.2, 118.4, 117.5, 113.1, 90.7, 40.4, 1.1, -2.9. _Quinoid (TFA added to NMR sample)_: $^{1}$H NMR (500 MHz, $CD_3OD$) δ 7.64 (d, *J* = 8.2 Hz, 1H), 7.52 (t, *J* = 7.9 Hz, 1H), 7.27 (d, *J* = 2.7 Hz, 2H), 7.18 (d, *J* = 8.0 Hz, 1H), 7.06 (d, *J* = 9.3 Hz, 2H), 6.76 (dd, *J* = 9.3, 2.7 Hz, 2H), 3.21 (s, 12H), 0.64 (s, 3H), 0.56 (s, 3H). HRMS (ESI$^+$) m/z 445.1942 calcd for $[C_{26}H_{29}N_2O_3Si]^+$ (M+H$^+$), 445.1931 found.

**HD653**

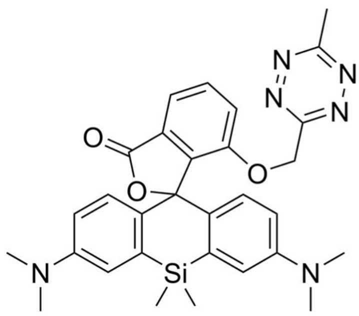

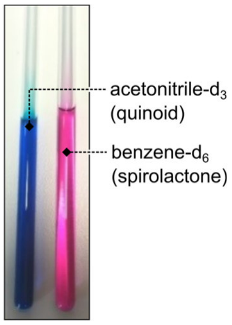

**23** (16.1 mg, 36.2 µmol) was dissolved in anhydrous THF (0.4 mL) in a Schlenk tube charged with 4 Å mol. sieve under argon and at rt. The marginally blue coloration of the solution indicated predominant presence of the spirolactone isomer. Separately, **3** (20.5 mg, 109 µmol) was dissolved in anhydrous THF (0.3 mL) and added to the first solution. The resulting mixture was cooled to 0 °C and $Cs_2CO_3$ (35.4 mg, 109 µmol) was added. While allowing to heat to rt, it was subsequently stirred for 1 h. Next, sat. aq. $NH_4Cl$ (2 mL), water (1 mL) and DCM (2 mL) were added in sequence and the aq. phase was extracted with DCM (4x2 mL). The combined organic phases were washed with brine (4 mL), dried over $Na_2SO_4$ and concentrated under reduced pressure. Purification of the crude residue by flash chromatography ($SiO_2$, 0-30% EtOAc / cyclohexane, const. 20% DCM) afforded the title product as pink solid (8.7 mg, 43%).

_Spirolactone:_ $^{1}$H NMR (500 MHz, $C_6D_6$) δ 7.70 (d, *J* = 7.6 Hz, 1H), 7.07 (t, *J* = 7.9 Hz, 1H), 7.03 (d, *J* = 8.8 Hz, 2H), 6.98 (d, *J* = 2.7 Hz, 2H), 6.89 (d, *J* = 8.1 Hz, 1H), 6.39 (dd, *J* = 8.8, 2.8 Hz, 2H), 4.92 (s, 2H), 2.52 (s, 12H), 2.30 (s, 3H), 0.71 (s, 3H), 0.48 (s, 3H). $^{13}$C NMR (126 MHz, $C_6D_6$) δ 169.1, 168.3, 165.0, 153.9, 149.8, 141.4, 138.2, 132.4, 131.7, 131.3, 128.4, 119.1, 117.4, 117.1, 113.8, 92.0, 68.1, 40.1, 20.7, 1.1, -2.7. _Quinoid (TFA added to NMR sample):_ $^{1}$H NMR (500 MHz, $CD_3CN$) δ 7.74 – 7.65 (m, 2H), 7.57 (d, *J* = 8.0 Hz, 1H), 7.18 (d, *J* = 2.4 Hz, 2H), 6.83 (d, *J* = 9.1 Hz, 2H), 6.65 (dd, *J* = 9.1, 2.5 Hz, 2H), 5.55 (s, 2H), 3.09 (s, 12H), 2.93 (s, 3H), 0.53 (s, 3H), 0.52 (s, 3H). HRMS (ESI$^+$) m/z 553.2378 calcd for $[C_{30}H_{33}N_6O_3Si]^+$ (M+H$^+$), 553.2364 found.

**7'-(Methoxymethoxy)-*N*$^3$,*N*$^3$,*N*$^7$,*N*$^7$,5,5-hexamethyl-3'*H*,5*H*-spiro[dibenzo[b,e]siline-10,1'-isobenzofuran]-3,7-diamine (21)**

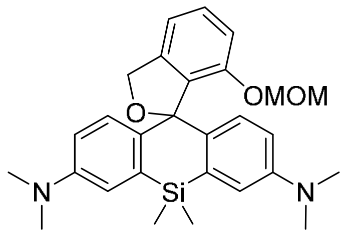

A procedure from the literature was adapted for the synthesis of this compound.[6]

**18** (75 mg, 164 µmol) was dissolved in anhydrous and degassed THF (4 mL) under argon and cooled to -78 °C. *tert*-Butyllithium (0.425 mL, 723 µmol, 1.7 M in pentane) was added dropwise and the mixture was stirred for 45 min. Next, it was warmed to -20 °C and a solution of **14** (70.2 mg, 362 µmol) in anhydrous and degassed THF (4 mL) was added over 30 min using a syringe pump. The reaction was stirred for 18 h while allowing to heat to rt. Then sat. aq. $NH_4Cl$ (3 mL) and water (3 mL) were added and the mixture was extracted with EtOAc (3x4 mL). The combined organic layers were washed with brine (5 mL), dried over $MgSO_4$ and concentrated under reduced pressure. The residue was dissolved in MeOH (3 mL) and glacial acetic acid was added (30 µL). It was then again concentrated under reduced pressure and purification by flash chromatography ($SiO_2$, 1. DCM, 2. 10-20% EtOAc / DCM, 3. 0-8% MeOH / DCM; const. 1 % AcOH over all steps) afforded the title product as colorless solid (41 mg, 53%).

$^1$H NMR (500 MHz, $CDCl_3$) δ 7.35 (t, *J* = 7.8 Hz, 1H), 7.03 – 6.95 (m, 4H), 6.88 (d, *J* = 8.8 Hz, 2H), 6.61 (d, *J* = 7.4 Hz, 2H), 5.12 (s, 2H), 4.92 (s, 2H), 3.01 (s, 3H), 2.94 (s, 12H), 0.57 (s, 3H), 0.55 (s, 3H). $^{13}$C NMR (75 MHz, $CDCl_3$) δ 152.3, 148.7, 143.2, 139.1, 136.1, 133.6, 129.8, 128.6, 116.9, 114.3, 113.9, 112.5, 93.1, 92.6, 72.2, 55.9, 40.9, 1.0, -1.9. HRMS (ESI$^+$) m/z 475.2411 calcd for $[C_{28}H_{35}N_2O_3Si]^+$ (M+Na$^+$), 475.2409 found.

**3,7-Bis(dimethylamino)-5,5-dimethyl-3'*H*,5*H*-spiro[dibenzo[b,e]siline-10,1'-isobenzofuran]-7'-ol (24)**

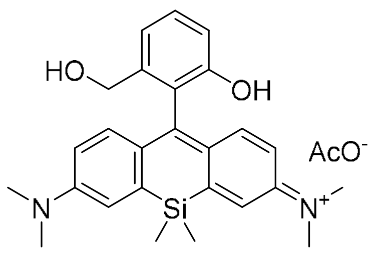

A solution of **21** (25.0 mg, 52.7 µmol) in DCM (1.25 mL) was cooled to 0 °C and TFA (0.25 mL, 3.78 mmol) was added dropwise. The ice bath was removed and the mixture was stirred for 18 h at rt. After coevaporation with toluene under reduced pressure, the crude residue was purified by flash chromatography ($SiO_2$, 0-10% MeOH / DCM, const. 1% AcOH) to afford the title product as blue solid (22 mg, 85%).

_Quinoid:_ $^{1}$H NMR (500 MHz, $CD_3OD$) δ 7.41 (t, *J* = 7.9 Hz, 1H), 7.32 (d, *J* = 2.1 Hz, 2H), 7.25 (d, *J* = 9.6 Hz, 2H), 7.20 (d, *J* = 7.7 Hz, 1H), 6.90 (d, *J* = 8.1 Hz, 1H), 6.76 (dd, *J* = 9.6, 2.1 Hz, 2H), 4.23 (s, 2H), 3.33 (s, 12H), 0.62 (s, 3H), 0.57 (s, 3H). $^{13}$C NMR (126 MHz, $CD_3OD$) δ 168.8, 155.9, 155.2, 149.4, 142.1, 141.9, 131.1, 129.1, 125.4, 121.7, 118.9, 115.1, 115.0*, 62.4, 40.8, -0.8, -1.6 (* peak derived from gHSQC, overlapping signal in $^{13}$C NMR). _Spiroether:_ $^{1}$H NMR (300 MHz, $CDCl_3$) δ 7.41 – 7.32 (m, 1H), 7.06 – 7.01 (m, 2H), 6.96 – 6.82 (m, 1H), 6.60 (d, *J* = 9.3 Hz, 2H), 5.01 (s, 2H), 2.96 (s, 12H), 0.63 (s, 3H), 0.56 (s, 3H). HRMS (ESI$^+$) m/z 431.2149 calcd for $[C_{26}H_{31}N_2O_2Si]^+$ (M+H$^+$), 431.2150 found.

**sb-HD656**

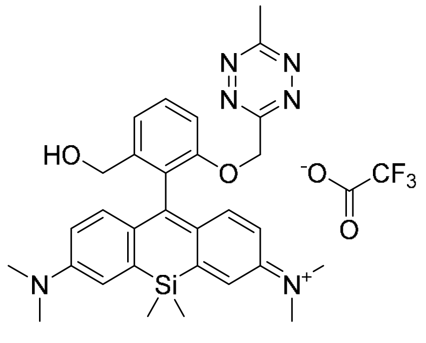

**24** (9.52 mg, 19.4 µmol) was dissolved in anhydrous THF (0.26 mL) in a Schlenk tube charged with 4 Å mol. sieve under argon and at rt. The marginally blue coloration of the solution indicated predominant presence of the spiroether isomer. Separately, **3** (16.5 mg, 87.3 µmol) was dissolved in anhydrous THF (0.2 mL) and added to the first solution. The resulting mixture was cooled to 0 °C and $Cs_2CO_3$ (25.3 mg, 77.6 µmol) was added. While allowing to heat to rt, it was subsequently stirred for 18 h. Next, sat. aq. $NH_4Cl$ (2 mL), water (2 mL) and DCM (2 mL) were added in sequence and the aq. phase was extracted with DCM (3x2 mL). The combined organic phases were washed with brine (3 mL), dried over $MgSO_4$ and concentrated under reduced pressure. The crude residue was directly purified by HPLC (30-90% solvent B/ solvent A) to afford the title product as blue solid (0.86 mg, 6.8%).

$^1$H NMR (500 MHz, $CD_3OD$) δ 7.58 (t, *J* = 8.2 Hz, 1H), 7.40 (d, *J* = 7.7 Hz, 1H), 7.30 (d, *J* = 8.4 Hz, 1H), 7.28 (d, *J* = 2.7 Hz, 2H), 7.14 (d, *J* = 9.5 Hz, 2H), 6.71 (dd, *J* = 9.6, 2.8 Hz, 2H), 5.57 (s, 2H), 4.25 (s, 2H), 3.33 (s, 12H), 2.93 (s, 3H), 0.57 (s, 3H), 0.51 (s, 3H). $^{13}$C NMR (126 MHz, $CD_3OD$) δ 169.8, 167.2*, 166.6, 156.3, 155.8, 149.2, 142.4, 141.9, 131.4, 128.9, 127.9, 121.7, 121.5, 115.2, 113.4, 69.7, 62.1, 40.9, 21.2, -1.3, -1.4 (*peak derived from gHMBC). HRMS (ESI$^+$) m/z 539.2585 calcd for $[C_{30}H_{35}N_6O_2Si]^+$ (M+H$^+$), 539.2610 found.

#### 3.4 Cell impermeable probes

**HD561x**

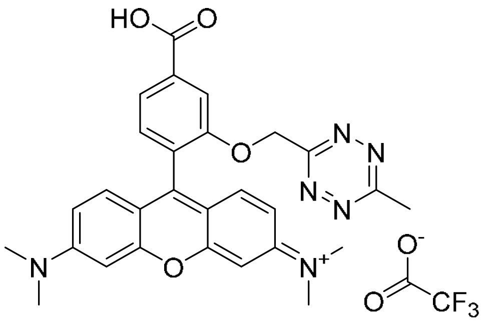

In a Schlenk tube charged with 4 Å mol. sieves, **11** (38.3 mg, 166 µmol) and **15** (29.7 mg, 116 µmol) were dissolved in anhydrous 1,2-dichloroethane (0.6 mL) under argon at rt. The solution was cooled to 0 °C and $AlBr_3$ (30.9 mg, 116 µmol) was added. The ice bath was removed and the reaction mixture was stirred for 3 h at rt. Subsequently, *p*-chloranil (28.5 mg, 116 µmol) was added and it was stirred for 30 min. Next, the mixture was filtered over a small silica pad (1-10% MeOH / DCM) and the filtrate was concentrated. The residue was dissolved in DCM (1 mL), cooled to 0 °C and anisole (16.8 µL, 154 µmol) was added. After the dropwise addition of trifluoroacetic acid (275 µL, 4.24 mmol), the mixture was allowed to heat to rt and stirred for 5 h. Then toluene (2 mL) was added and the solution was concentrated under reduced pressure. Purification of the crude residue by flash chromatography ($SiO_2$, 0-10% MeOH / DCM, const. 0.1% TFA) gave the title product as purple solid (9.5 mg, 13%).

$^1H$ NMR (500 MHz, $CD_3OD$) δ 8.06 (s, 1H), 7.95 (d, *J* = 7.7 Hz, 1H), 7.46 (d, *J* = 7.7 Hz, 1H), 7.33 (d, *J* = 9.5 Hz, 2H), 7.07 (dd, *J* = 9.5, 2.5 Hz, 2H), 6.91 (d, *J* = 2.5 Hz, 2H), 5.75 (s, 2H), 3.31 (s, 12H), 2.95 (s, 3H). $^{13}C$ NMR (126 MHz, $CD_3OD$) δ 170.1, 168.4, 166.3, 159.1, 159.0, 156.8, 155.9, 135.8, 132.7, 132.4, 127.4, 124.3, 115.6, 115.5, 114.8, 97.4, 69.3, 41.0, 21.2. HRMS ($ESI^+$) m/z 511.2088 calcd for $[C_{28}H_{27}N_6O_4]^+$ ($M^+$), 511.2095 found.

**HD654x**

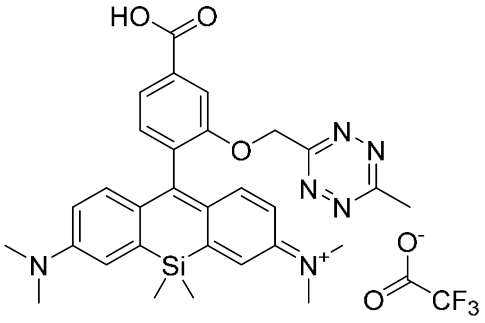

In a Schlenk tube charged with 4 Å mol. sieves, **11** (61.2 mg, 185 µmol) and **17** (55.3 mg, 185 µmol) were dissolved in anhydrous 21,2-dichloroethane (1.2 mL) under argon at rt. The solution was cooled to 0 °C and $AlBr_3$ (49.4 mg, 185 µmol) was added. The ice bath was removed and the reaction mixture was stirred for 2.5 h at rt. Subsequently, *p*-chloranil (91.1 mg, 371 µmol) was added and it was stirred for 3.5 h. Next, water (3 mL) and DCM (4 mL) were added and the mixture was filtered over a small pad of wool. The filtrate was collected and the organic phase was washed with sat. aq. $NH_4Cl$ (3 mL), dried over $Na_2SO_4$ and concentrated. The residue was dissolved in DCM (0.5 mL), cooled to 0 °C and anisole (16.1 µL, 116 µmol) was added. After the dropwise addition of trifluoroacetic acid (250 µL, 3.24 mmol), the mixture was allowed to heat to rt and stirred for 2 h. Then toluene (1 mL) was added and the solution was concentrated under reduced pressure. Purification of the crude residue by flash chromatography ($SiO_2$, 0-10% MeOH / DCM, const. 0.1% TFA) gave the title product as blue solid (6.8 mg, 5.5%).

$^1H$ NMR (500 MHz, $CD_3OD$) δ 7.98 (d, *J* = 1.4 Hz, 1H), 7.88 (dd, *J* = 7.8, 1.4 Hz, 1H), 7.34 – 7.27 (m, 3H), 7.14 (d, *J* = 9.7 Hz, 2H), 6.75 (dd, *J* = 9.6, 2.9 Hz, 2H), 5.70 (s, 2H), 3.34 (s, 12H), 2.95 (s, 3H), 0.58 (s, 3H), 0.54 (s, 3H). $^{13}C$ NMR (126 MHz, $CD_3OD$) δ 169.9, 168.7, 166.9, 166.4, 156.8, 155.7, 149.3, 142.4, 134.8, 134.4, 132.1, 128.6, 124.1, 122.0, 115.4, 115.1, 69.5, 40.9,

21.2, -1.2, -1.3. (TFA was added to the NMR sample prior measuring.) HRMS ($ESI^+$) m/z 553.2378 calcd for $[C_{30}H_{33}N_6O_3Si]^+$ ($M^+$), 553.2395 found.

#### 3.5 Various tetrazine (Si)rhosamines

**General procedure (2) the preparation of tetrazine (Si)rhosamines**

This procedure for ***m*-TzR** is representative:

In a Schlenk tube charged with 4 Å mol. sieves, **5** (15.0 mg, 65.2 µmol) and **15** (16.7 mg, 65.2 µmol) were dissolved in anhydrous 1,2-dichloroethane (0.65 mL) under argon at rt. Next, $AlBr_3$ was added dropwise (72 µL, 72 µmol, 1 M in dibromomethane) and the reaction mixture was stirred for 3 h at 80 °C. After cooling down to rt, *p*-chloranil (16.0 mg, 65.2 µmol) was added and it was stirred for 17 h. Subsequently, methanol (0.5 mL) was added, the mixture was filtered over a small silica pad and the filtrate was concentrated under reduced pressure. Purification of the crude residue by flash chromatography ($SiO_2$, 0-10% methanol / DCM) afforded ***m*-TzR** as a purple solid with green pearlescent shine (21 mg, 59%).

***o*-TzR**

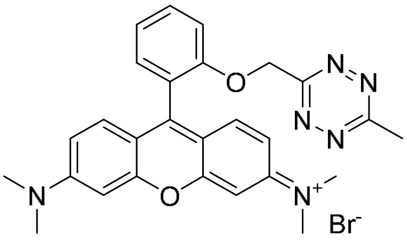

The title compound (22 mg, 54%, purple solid with green pearlescent glance) was prepared according to general procedure (2) from **4** (17.0 mg, 73.8 µmol) and **15** (18.9 mg, 73.8 µmol).

$^1H$ NMR (500 MHz, $CD_3CN$) δ 7.65 (ddd, *J* = 8.7, 6.1, 2.9 Hz, 1H), 7.44 (d, *J* = 8.4 Hz, 1H), 7.31 – 7.28 (m, 2H), 7.26 (d, *J* = 9.5 Hz, 2H), 6.95 (dd, *J* = 9.5, 2.3 Hz, 2H), 6.79 (d, *J* = 2.3 Hz, 2H), 5.60 (s, 2H), 3.25 (s, 12H), 2.92 (s, 3H). $^{13}C$ NMR (126 MHz, $CD_3CN$) δ 169.6, 165.9, 158.6, 158.3, 156.6, 156.1, 132.9, 132.5, 131.7, 123.1, 122.5, 115.2, 115.1, 114.7, 97.1, 69.5, 41.4, 21.5. HRMS ($ESI^+$) m/z 467.2190 calcd for $[C_{27}H_{27}N_6O_2]^+$ ($M^+$), 467.2185 found.

***m*-TzR**

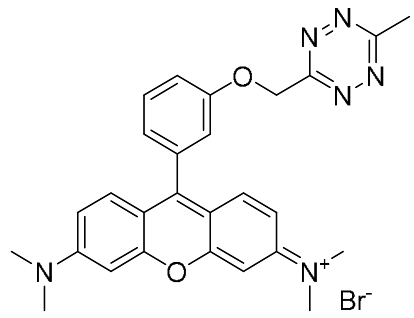

The title compound (21 mg, 59%, purple solid with green pearlescent glance) was prepared according to general procedure (2) from **5** (15.0 mg, 65.2 µmol) and **15** (16.7 mg, 65.2 µmol).

$^{1}$H NMR (500 MHz, $CD_3CN$): δ 7.60 (dd, *J* = 8.4, 7.5 Hz, 1H), 7.36 (ddd, *J* = 8.5, 2.7, 0.9 Hz, 1H), 7.35 (d, *J* = 9.5 Hz, 2H), 7.15 (dd, *J* = 2.7, 1.5 Hz, 1H), 7.08 (dt, *J* = 7.5, 1.2 Hz, 1H), 7.00 (dd, *J* = 9.5, 2.5 Hz, 2H), 6.83 (d, *J* = 2.5 Hz, 2H), 5.73 (s, 2H), 3.26 (s, 12H), 3.02 (s, 3H). $^{13}$C NMR (126 MHz, $CDCl_3$) δ 169.0, 165.3, 158.3, 157.9, 157.4, 157.2, 133.5, 131.9, 130.6, 123.2, 116.6, 116.6, 114.7, 113.5, 97.1, 68.2, 41.4, 21.5. HRMS (ESI$^+$) m/z 467.2190 calcd for $[C_{27}H_{27}N_6O_2]^+$ ($M^+$), 467.2209 found.

***p*-TzR**

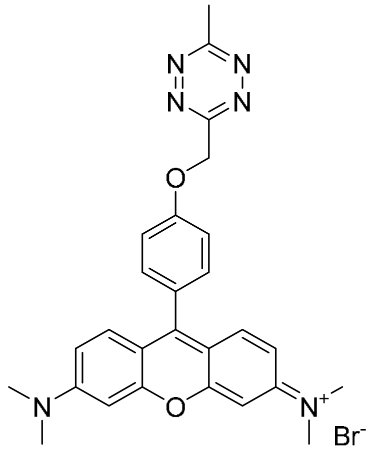

The title compound (32 mg, 91%, purple solid) was prepared according to general procedure (2) from **6** (15.0 mg, 65.2 µmol) and **15** (16.7 mg, 65.2 µmol).

$^{1}$H NMR (500 MHz, $CD_3CN$) δ 7.43 (d, *J* = 8.8 Hz, 2H), 7.42 (d, *J* = 9.5 Hz, 2H), 7.35 (d, *J* = 8.7 Hz, 2H), 7.02 (dd, *J* = 9.5, 2.6 Hz, 2H), 6.84 (d, *J* = 2.5 Hz, 2H), 5.80 (s, 2H), 3.26 (s, 12H), 3.05 (s, 3H). $^{13}$C NMR (126 MHz, $CD_3CN$) δ 169.9, 166.3, 160.7, 158.9, 158.8, 158.3, 132.6, 132.5, 126.2,

116.3, 115.2, 114.5, 97.2, 69.0, 41.3, 21.6. HRMS (ESI$^+$) m/z 467.2190 calcd for $[C_{27}H_{27}N_6O_2]^+$ ($M^+$), 467.2204 found.

***o*-TzSiR**

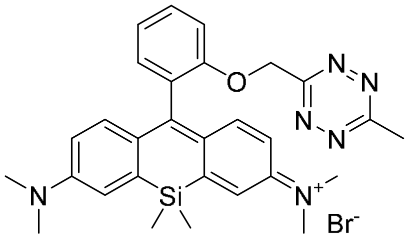

In a Schlenk tube charged with 4 Å mol. sieves, **4** (19.8 mg, 86.0 µmol) and **17** (25.7 mg, 86.0 µmol) were dissolved in anhydrous 1,2-dichloroethane (0.6 mL) under argon at rt. Next, $AlBr_3$ was added (22.9 mg, 86 µmol) and the reaction mixture was stirred for 2 h at rt. Subsequently, *p*-chloranil (21.1 mg, 86 µmol) was added and it was stirred for 3 h. Subsequently, water (2 mL) was added, the mixture was filtered over a small wool pad and the filtrate was concentrated under reduced pressure. Purification of the crude residue by flash chromatography ($SiO_2$, 1. 0-20% EtOAc / DCM, 2. 0-5% MeOH / DCM) afforded the title product as a blue solid (2.3 mg, 4.5%).

$^1$H NMR (500 MHz, $CD_3CN$) δ 7.56 (ddd, *J* = 8.4, 7.5, 1.7 Hz, 1H), 7.36 (d, *J* = 8.4 Hz, 1H), 7.23 (d, *J* = 2.9 Hz, 2H), 7.21 (dd, *J* = 7.5, 0.7 Hz, 1H), 7.15 (dd, *J* = 7.5, 1.8 Hz, 1H), 7.09 (d, *J* = 9.6 Hz, 2H), 6.65 (dd, *J* = 9.6, 2.9 Hz, 2H), 5.56 (s, 2H), 3.27 (s, 12H), 2.91 (s, 3H), 0.55 (s, 3H), 0.52 (s, 3H). $^{13}$C NMR (126 MHz, $CD_3OD$) δ 169.8, 168.7, 166.6, 156.7, 155.7, 149.4, 142.8, 131.8, 131.7, 130.1, 129.1, 122.8, 121.8, 114.9, 114.9, 69.5, 40.8, 21.2, -1.2, -1.3. HRMS (ESI$^+$) m/z 509.2480 calcd for $[C_{29}H_{33}N_6OSi]^+$ ($M^+$), 509.2483 found.

#### 3.6 Non-quenched fluorophores

**General procedure (3) the preparation of $DA_{inv}$ cycloadducts**

This procedure for **Ref-1** is representative:

***o*-TzR** (2.0 mg, 3.7 µmol) was dissolved in in PBS/MeCN (7:3 (v/v), 1 mL) at rt. Then a solution of ((1R,8S,9s)-bicyclo[6.1.0]non-4-yn-9-yl)methanol **BCN** (2.1 mg, 14.0 µmol) in PBS/MeCN (1:1 (v/v), 0.4 mL) was added. The reaction was incubated at 25 °C for 2 h in a thermoshaker at 750 rpm and subsequently purified by HPLC (30-90% solvent B/ solvent A).

**Ref-1**

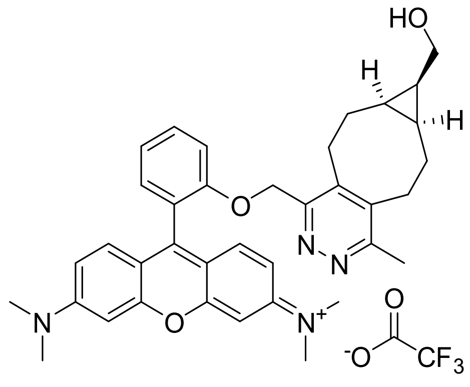

The title compound was prepared according to general procedure (3) from ***o*-TzR** (2.0 mg, 3.7 µmol) and ((1R,8S,9s)-bicyclo[6.1.0]non-4-yn-9-yl)methanol **BCN** (2.1 mg, 14.0 µmol).

HRMS (ESI$^+$) m/z 589.3173 calcd for $[C_{37}H_{41}N_4O_3]^+$ ($M^+$), 589.3221 found.

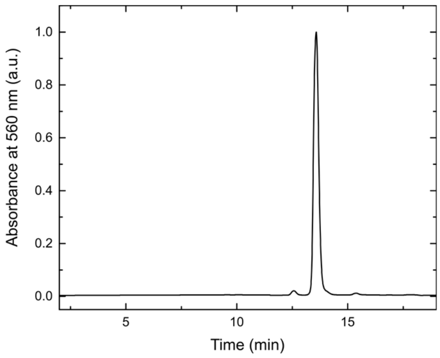

HPLC chromatogram of **Ref-1**.

**Ref-3**

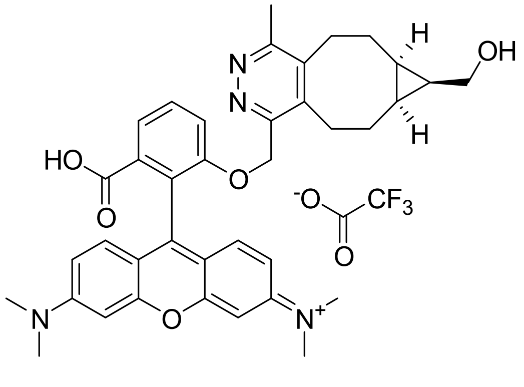

The title compound was prepared according to general procedure (3) from **HD555** (1.0 mg, 1.6 µmol) and ((1R,8S,9s)-bicyclo[6.1.0]non-4-yn-9-yl)methanol **BCN** (1.20 mg, 8.01 µmol).

HRMS (ESI$^+$) m/z 633.3071 calcd for $[C_{38}H_{41}N_4O_5]^+$ ($M^+$), 633.3079 found.

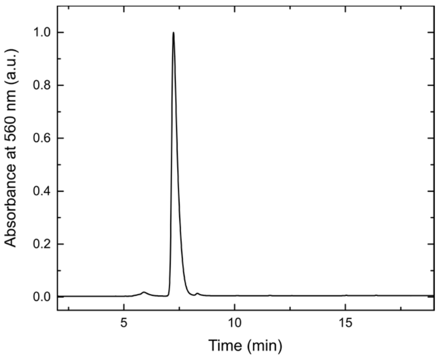

HPLC chromatogram of **Ref-3**.

**Ref-4**

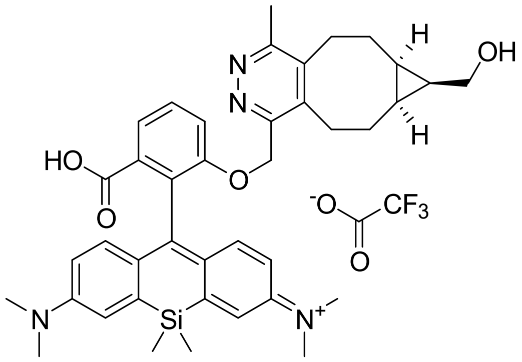

The title compound was prepared according to general procedure (3) from **HD653** (1.0 mg, 1.8 µmol) and ((1R,8S,9s)-bicyclo[6.1.0]non-4-yn-9-yl)methanol **BCN** (1.36 mg, 9.05 µmol).

HRMS (ESI$^+$) m/z 675.3361 calcd for $[C_{40}H_{47}N_4O_4Si]^+$ ($M^+$), 675.3369 found.

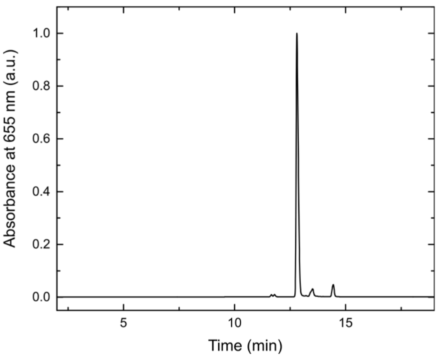

HPLC chromatogram of **Ref-4**.

**Ref-5**

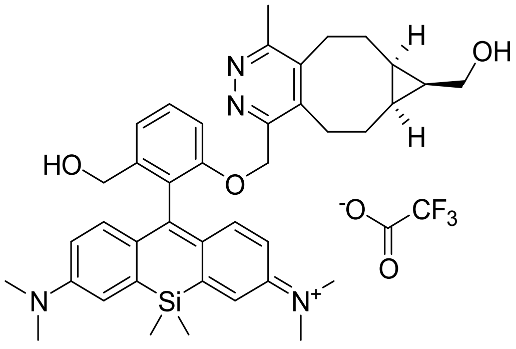

The title compound was prepared according to general procedure (3) from **sb-HD656** (44 µL, 10 mM in DMSO, 0.44 µmol) and ((1R,8S,9s)-bicyclo[6.1.0]non-4-yn-9-yl)methanol **BCN** (22 µL, 0.1 M in DMSO, 2.20 µmol).

HRMS (ESI$^+$) m/z 661.3568 calcd for $[C_{40}H_{49}N_4O_3Si]^+$ ($M^+$), 661.3586 found.

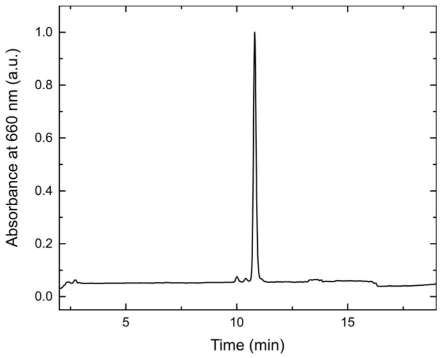

HPLC chromatogram of **Ref-5**.

**Ref-2**

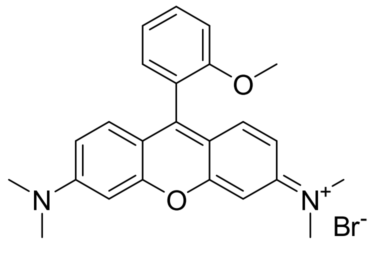

The title compound (54.0 mg, 81%, red solid) was prepared according to general procedure (2) from 2-methoxybenzaldehyde (22.2 µL, 184 µmol) and **15** (47.1 mg, 184 µmol).

$^1$H NMR (300 MHz, $CD_3OD$) δ 7.69 (ddd, *J* = 8.9, 7.0, 2.2 Hz, 1H), 7.39 – 7.20 (m, 5H), 7.11 (dd, *J* = 9.5, 2.5 Hz, 2H), 6.98 (d, *J* = 2.5 Hz, 2H), 3.77 (s, 3H), 3.34 (s, 12H). $^{13}$C NMR (75 MHz, $CD_3OD$) δ 159.2, 159.0, 158.2, 158.0, 133.2, 132.8, 131.7, 122.0, 121.9, 115.4, 112.9, 97.4, 56.3, 40.9. HRMS (ESI$^+$) m/z 373.1911 calcd for $[C_{24}H_{25}N_2O_2]^+$ ($M^+$), 373.1917 found.

#### 3.7 Other Compounds

**Phalloidin-BCN**

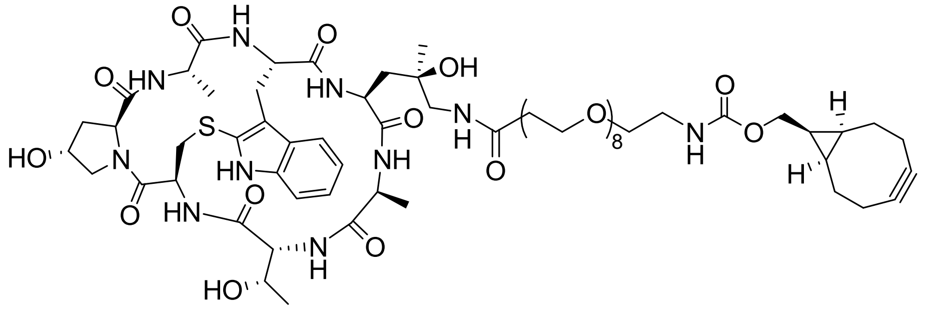

Amino-phalloidin (0.20 mg, 0.25 µmol; Bachem Holding AG, Bubendorf, Switzerland; cat. number: H-7634) was dissolved in anhydrous DMF (50 µL) and added to a solution of BCN-PEG8-NHS (1.81 mg, 2.54 µmol; SiChem GmbH, Bremen, Germany; cat. number: SC-8108) in DMF (170 µL) at rt. After the addition of DIPEA (0.58 µL, 3.39 µmol), the mixture was incubated for 90 min in a thermoshaker at 600 rpm and subsequently purified by HPLC (20-90% MeCN/$H_2O$, const. 0.1% formic acid). Phalloidin-BCN was afforded as white solid, and after photometric determination of the amount of substance[11], a 1 mM stock solution in anhydrous DMSO was prepared.

HRMS (ESI$^+$) m/z 1409.6521 calcd for $[C_{65}H_{98}N_{10}NaO_{21}S]^+$ (M+Na$^+$), 1409.6613 found.

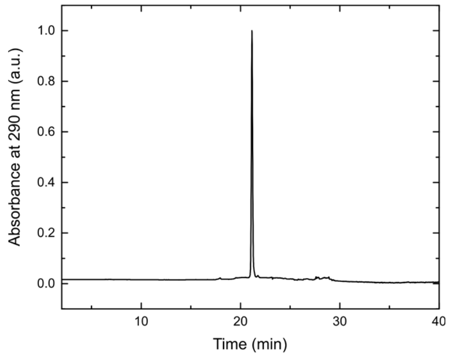

HPLC chromatogram of **Phalloidin-BCN**.

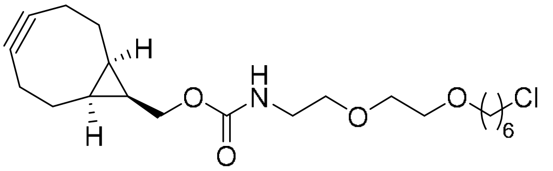

**HTL-BCN**

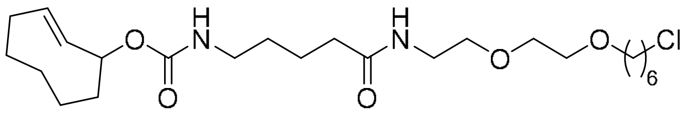

**HTL-TCO***

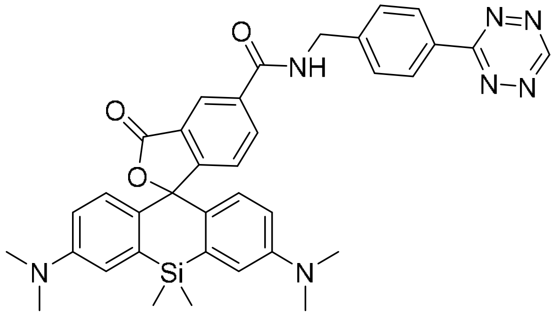

**SiR-Tz**

**Supplementary Figure 9.** Structures of miscellaneous compounds.

**HTL-BCN**[12] and **HTL-TCO***[13] were prepared according to literature procedures. **SiR-Tz** is commercially available and can be purchased from Spirochrome Ltd., Stein am Rhein, Switzerland (cat. number: SC008).

### 4 X-ray crystallography

Pink brick crystals were grown by vapor diffusion of *n*-pentane into a solution of **HD555** (~2 mg) in MeOH (50 µL) and DCM (150 µL) at rt.

Crystal data and structure refinement is compiled in Supplementary Table 3, the x-ray solid-state molecular structure including 50% probability ellipsoids is depicted in Supplementary Figure 10. X-ray crystallographic measurements were executed on a STOE Stadivari instrument. 0.5 deg omega-scans were performed with CCD area detector, covering the asymmetric unit in reciprocal space with a mean redundancy of 5.10 and a completeness of 99.7%. This lead to a resolution of 0.95 Å (19445 reflections measured, 3724 unique (R(int) = 0.0400), 2337 observed ($I > 2\sigma(I)$)). Intensities were corrected for Lorentz and polarization effects and an empirical scaling and absorption correction was applied using X-Area LANA 1.70.0.0 (STOE, 2017) based on the Laue symmetry of the reciprocal space ($\mu = 1.44 mm^{-1}$, $T_{min} = 0.63$, $T_{max} = 1.45$) The structure was solved with SHELXT-2014[14] and refined against $F^2$ with a full-matrix least-squares algorithm using the SHELXL-2018/3

(Sheldrick, 2018) software[15]. 385 parameters were refined and hydrogen atoms were treated using appropriate riding models (except H38 at O38, which was not considered at all) Goodness of fit was 2.69 for observed reflections, final residual values R1(F) = 0.154, wR($F^2$) = 0.380 for observed reflections, residual electron density -0.36 to 0.87 eÅ$^{-3}$.

CCDC 2021104 contains the supplementary crystallographic data for this paper. The data can be obtained free of charge from the Cambridge Crystallographic Data Centre via www.ccdc.cam.ac.uk/structures.

The structure in Supplementary Figure 10 was visualized with Mercury 2020.1[16] and POV-Ray 3.7[17].

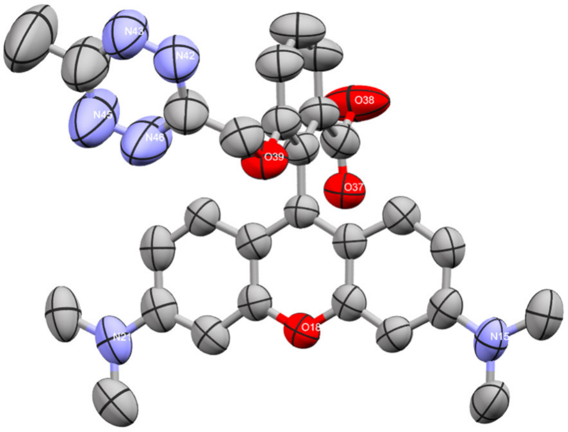

**Supplementary Figure 10. X-ray solid-state structure of HD555**. Anisotropic displacement parameters set to 50% probability. Hydrogen atoms omitted for clarity.

**Supplementary Table 3.** Crystal data and structure refinement for **HD555**.

| | **HD555** |
|---|---|
| Empirical formula | $C_{28}H_{26}ClN_6O_4$ |
| T / K | 200(2) |
| X radiation, λ / Å | $CuK_{\alpha}$, 1.54178 |
| Crystal system | Monoclinic |
| Space group | C2/c |
| Z | 8 |
| a / Å | 28.071(2) |
| b / Å | 15.7283(8) |
| C / Å | 13.9652(11) |
| α / ° | 90 |
| β / ° | 97.823(6) |
| γ / ° | 90 |
| V / $Å^3$ | 6108.4(8) |
| ρ / g $cm^{-3}$ | 1.19 |
| μ / $mm^{-1}$ | 1.44 |
| Crystal size / $mm^3$ | 0.170 x 0.075 x 0.049 |
| σ range / ° | 3.2 to 54.2 |
| Index ranges | -29≤h≤29, -16≤k≤15, -14≤l≤11 |
| Reflections collected | 19445 |
| Independent reflections | 3724 ($R_{int}$ = 0.0400) |
| Observed reflections | 2337 (I > 2σ(I)) |
| Absorption correction | Semi-empirical from equivalents |
| Max. and min. transmission | 1.45, 0.63 |
| Refinement method | Full-matrix least-squares on $F^2$ |
| Data / restraints / parameters | 3724 / 325 / 385 |
| GoF on $F^2$ | 2.69 |
| R1, wR2 (I > 2σ(I)) | 0.154, 0.380 |
| Largest diff. peak, hole / e$Å^{-3}$ | 0.87, -0.36 |

### 5 In vitro protein labeling

##### *Preparation of EGFP-HaloTag purified protein*

A gylcerolstock of E. coli BL21(DE3) pLys (Promega GmbH) transformed with pEGFP-HaloTag[18] was added to lysogeny broth (LB) medium (30 mL) supplemented with kanamycin (50 µg/mL kanamycin) and incubated at 37 °C overnight. After inoculation of an expression culture in LB medium (1 L) with the overnight culture, protein expression was induced with isopropyl β-D-1-thiogalactopyranoside (IPTG, 0.5 mM) at $OD_{600}$ = 0.6 and carried out for 3.5 h. The culture was centrifuged (4000 rcf) at 4 °C, cells were resuspended in 10 mL Buffer A (50 mM Tris-HCl pH 7.5, 300 mM NaCl, 5 mM $MgSO_4$, 5 mM 2-mercaptoethanol, 5 % glycerol, 5 mM imidazole, cOmplete EDTA-free protease inhibitor [Roche]), and centrifuged (4000 rcf) at 4 °C. The supernatant was discardet and the pellet was stored at -80°C. The pellet was resuspended in 10 mL Buffer A and sonicated (3 x 2 min with 20 s break). The suspension was centrifuged at 37500 rcf at 4 °C for 30 min, the supernatant filtered through a 0.45 µm filter unit and subsequently loaded onto a HisTrap HP Ni-NTA affinity column (1 mL, GE Healthcare). The column was equilibrated with buffer A for 60 min and the protein was subsequently eluted with a gradient to 100% elution buffer (50 mM Tris-HCl pH 7.5, 300 mM NaCl, 5 mM $MgSO_4$, 5 mM 2-mercaptoethanol, 5% glycerol, 500 mM imidazole, cOmplete EDTA-free protease inhibitor (Roche)) over 60 min. Fractions were then analyzed by SDS-PAGE (10% (w/v) acrylamide). Combined protein fractions were concentrated using Amicon Ultra centrifugal filter devices (15 mL, 10 kDa cutoff, Merck) and further subjected to size exclusion chromatography (HiPrep 26/60 Sephacryl S-300R column (GE Healthcare), isocratic flow, Buffer B: 50 mM Tris-HCl (pH 7.5), 300 mM NaCl, cOmplete EDTA-free protease inhibitor (Roche)). Fractions were analyzed by SDS-PAGE (10% (w/v) acrylamide), pooled and concentrated using Amicon Ultra centrifugal filter devices (15 mL, 10 kDa cutoff, Merck). They were subsequently diluted with glycerol (1/1, v/v) snap-frozen at -77 °C and stored at -80 °C.

##### *Preparation of EGFP-HaloTag-BCN*

An aliquot purified EGFP-HaloTag protein was subjected to buffer exchange to PBS using an Amicon Ultra centrifugal filter device (0.5 mL, 10 kDa cutoff, Merck). The aliquot was further diluted with PBS to 1 µM and a 5-fold molar excess of a HTL-BCN DMSO stock solution was added (25µM). The mixture was incubated at 25 °C for 70 min in a thermoshaker at 600 rpm and subsequently diluted with PBS (1 mL) and concentrated with an Amicon Ultra centrifugal filter device (15 mL, 10 kDa cutoff, Merck). This step was repeated once with PBS (2 mL). The flowthrough was discarded and the sample transferred to another Amicon Ultra centrifugal filter device (0.5 mL, 10 kDa cutoff, Merck). After dilution with PBS (0.4 mL) and concentration the protein was recovered and the concentration of EGFP-HaloTag-BCN was calculated from the absorbance at 280 nm.

##### *Labeling of EGFP-HaloTag-BCN with tetrazine dyes and gel analysis*

A 10-fold molar excess of EGFP-HaloTag-BCN was added to dye solutions in PBS (1 µM dye, from a 0.2 mM DMSO stock solution) and the fluorescence increase was measured over time using a fluorimeter. For negative controls, DMSO was added instead of dye or EGFP-HaloTag was added instead of EGFP-HaloTag-BCN. After 45 min incubation at 37 °C, 20 equivalents of BCN were added to quench the reactions and they were incubated at 25 °C for 5 min. The samples were then treated with non-reducing SDS-sample buffer at 95 °C for 5 min and and loaded onto a SDS gel (10% (w/v) acrylamide). After SDS-PAGE was performed, the gels were rinsed briefly with water and scanned with a Typhoon FLA 9500 fluorescence scanner (GE Healthcare Life Sciences) with 635 nm laser excitation and ≥ 665 nm fluorescence detection (LPR filter) or 532 nm laser excitation and ≥ 575 nm fluorescence detection (LPG filter) depending on the applied fluorophore. In case of **sb-HD656**, the gel was treated with PBS (pH 3.5 or pH 7.0) for 3 h at rt before in-gel fluorescence scans. Finally, gels were stained with Coomassie Brilliant Blue.[19]

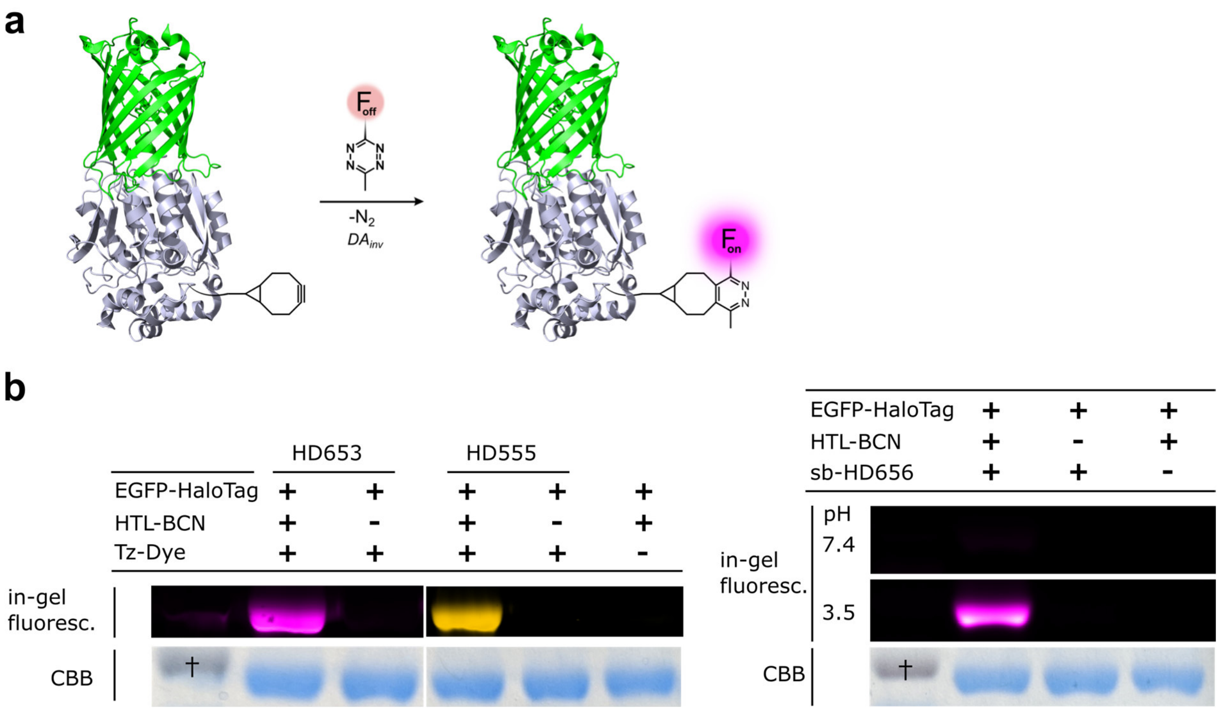

**Supplementary Figure 11. In vitro labeling of EGF-HaloTag-BCN with HDyes.** a: Schematic presentation of the labeling reaction (based on PDBs 2Y0G and 5Y2Y). b: In-gel fluorescence shows selective labeling of BCN-functionalized protein and confirms the pH-sensitivity of the protein-bound **sb-HD656**. †: ~55 kDa. CBB = Coomassie Brilliant Blue.

### 6 Ultrafast laser spectroscopy

##### Transient absorption measurements

Transient absorption (TA) was measured for all compounds using a femtosecond ultrashort pump pulse and white-light probe. The pump pulse was generated in a home-made noncollinear optical parametric amplifier and was spectrally centered at 575 nm for all samples. Typical pump pulses had a duration of 20 fs and an energy of 100 nJ with a repetition rate of 1 kHz. The white light probe continuum (from 450 to 750 nm) was generated in a 1-mm-thick sapphire substrate. Typical pump-probe cross-correlations of about 50 fs were characterized by fitting of the coherent spike measured in a buffer solution under similar experimental conditions. In order to avoid contributions from rotational dynamics, all TA measurements were performed in magic angle (54.7°) between the pump and probe polarizations. All samples were dissolved in a buffer solution (20% MeOH/PBS) with an absorption maximum of 0.3-0.33 OD (measured in 0.45 mm) and circulated in a flow cell to avoid accumulation effects during TA measurements.

#### Supplementary Note 1

Supplementary Figure 12 shows selected TA spectra for the initial 16 ps probe delay. The decay of the stimulated emission (SE) and ground-state bleach (GSB) for ***o*-TzR** is clearly observed in the initial picoseconds. An isosbestic point at about 508 nm between the GSB and the excited state dynamics (ESA) band in the initial 2 ps points to a picture where the excited state S1 decays mainly via emission back to the ground state recovering the GSB with the simultaneous decay of the ESA. This quenching of the excited state takes longer for the other two compounds (***m*/*p*-TzR**) and is absent for the reference compounds. See also Supplementary Figure 13 for selected traces at 560 nm and 630 nm for each compound.

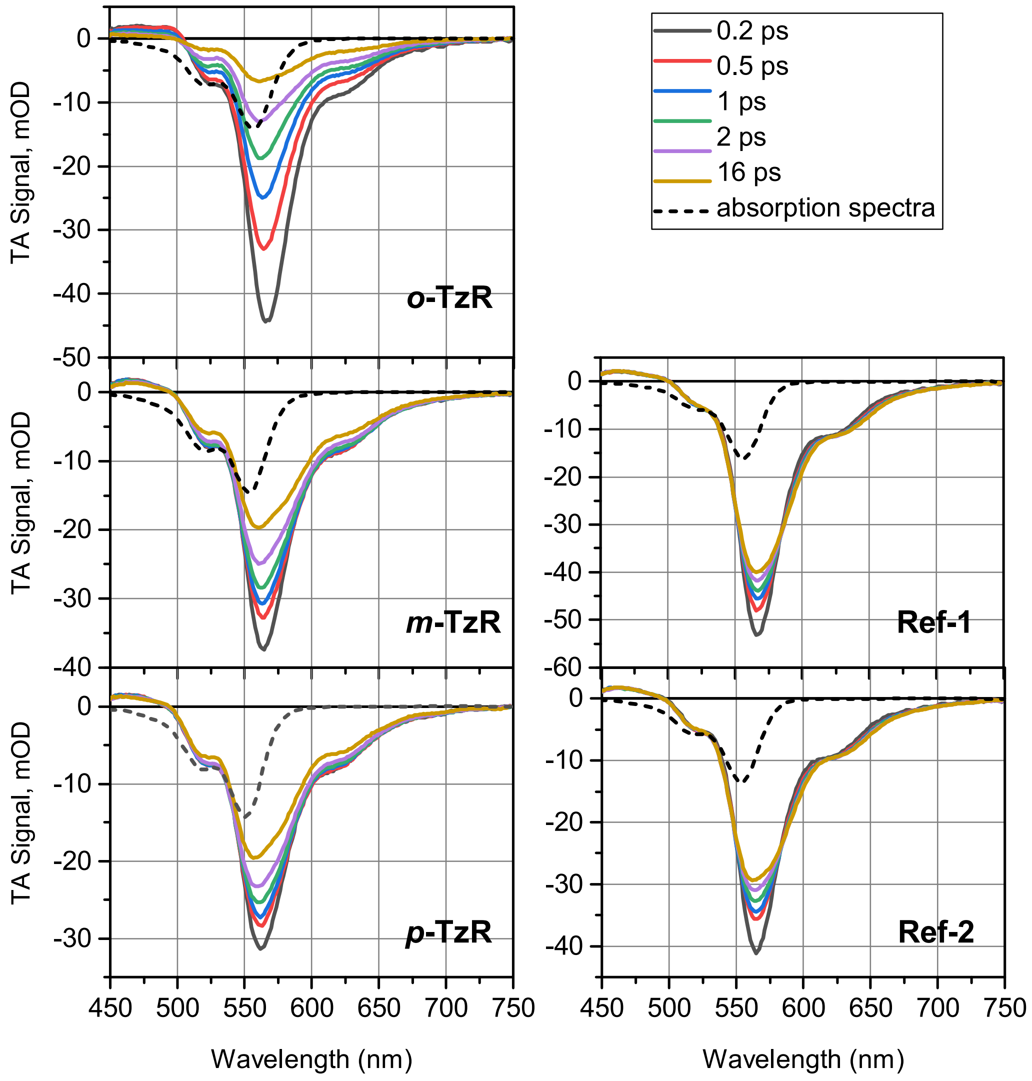

**Supplementary Figure 12. Selected transient spectra at specific delays.** Ground state absorption (dashed line) was rescaled to match the ground-state bleach signal at 0.2 ps around 525-530 nm.

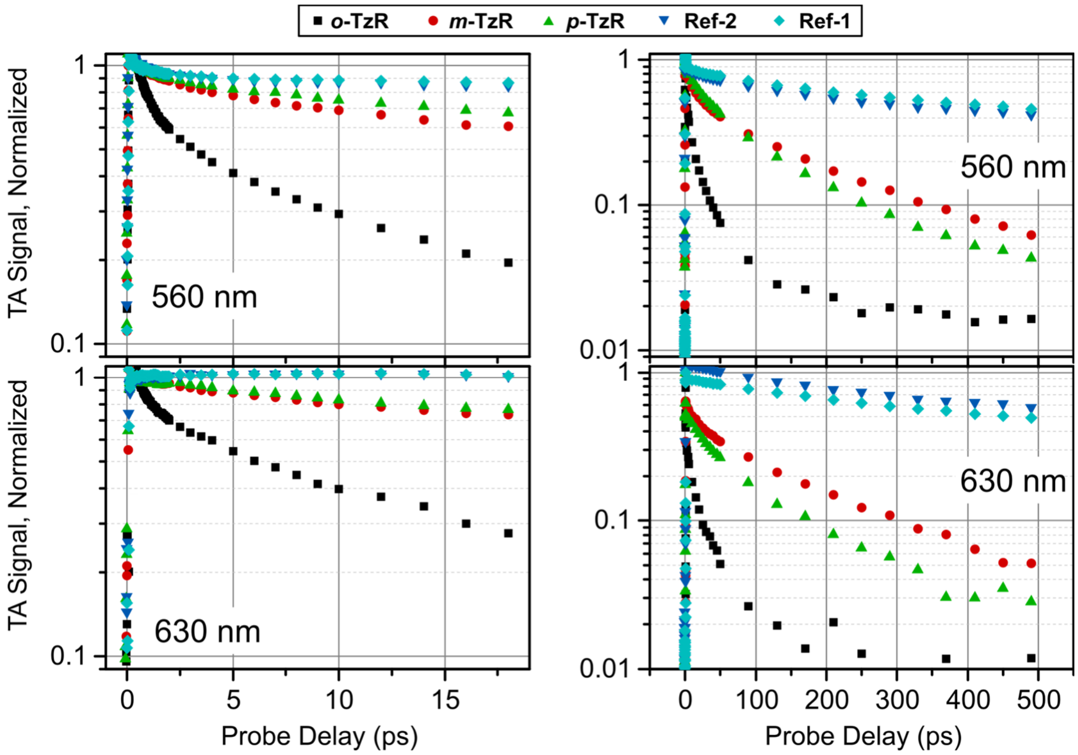

**Supplementary Figure 13. Selected transient absorption traces at 560 nm and 630 nm.** (Left Column) Short-time dynamics up to 18 ps. (Right column) Long-time dynamics up to 500 ps. Traces at 560 nm contain mainly ground-state bleach and stimulated emission. Traces at 630 nm contains stimulated emission only. Signal was normalized at 0.5 ps to improve comparison.

#### Supplementary note 2

Global target analysis applied with a sequential model to the full data set up to 500 ps requires three time constants and an offset. Supplementary Figure 14 shows the Species Associated Difference Spectra (SADS) for ***o*-TzR** and ***m*-TzR** exemplary. The first time constant is below one picosecond (Supplementary Table 4) and is related to the fast recovery of the signal at about 560 nm in all compounds. It is assigned to an evolution of the excited state near the Franck Condon region. This leads mainly to a fast recovery of the stimulated emission (also observed in the reference compounds) and almost no recovery in the ESA region. The respective SADS in the ***o*-TzR** compound shows additionally a pure SE recovery for red-shifted wavelengths above 560 nm, and decay of the ESA signal below 500 nm. This is not observed for ***m/p*-TzR**, where the SE contributions at 630 nm for the first and second SADS are the same. The third SADS is similar for all three compounds, with a recovery of the ESA, GSB and SE contributions. Here it is interesting to note that all three SADS for ***m/p*-TzR** form an isosbestic point at about 508 nm, while ***o*-TzR** displays an isosbestic point only for the two initial SADS. This an indication that the SE quenching mechanism may involve more than one quenching channel. This is more clearly by comparing the time constants associated to the second and third SADS in Supplementary Table 4. While $(\tau_2)^{-1}$ decreases exponentially with the separation between rhodamine and tetrazine (see Fig. 2d), $(\tau_3)^{-1}$ shows a non-trivial increase with the separation.

**Supplementary Table 4. Time constants obtained via global target analysis (sequential model).** Two datasets were analyzed independently: The initial 20 ps and up to 500 ps.

| | Time constants (ps) | ***o*-TzR** | ***m*-TzR** | ***p*-TzR** | **Ref-1** | **Ref-2** |
|---|---|---|---|---|---|---|
| Initial 20 ps | $\tau_1$ | 0.48(±0.01) | 0.39(±0.01) | 0.60(±0.02) | 1.33(±0.02) | 1.24(±0.02) |
| | $\tau_2$ | 5.54(±0.10) | 7.88(±0.02) | 16.64(±1.1) | - | - |
| | $\tau_3$ | offset | offset | offset | offset | offset |
| Up to 500 ps | $\tau_1$ | 0.21(±0.01) | 0.50(±0.02) | 0.76(±0.03) | 1.87(±0.05) | 1.65(±0.05) |
| | $\tau_2$ | 1.56(±0.06) | 11.36(±0.30) | 23.95(±1.30) | 258(±0.10) | 196(±0.03) |
| | $\tau_3$ | 23.50(±0.30) +offset | 142(±3) + offset | 116(±3) +offset | offset | offset |

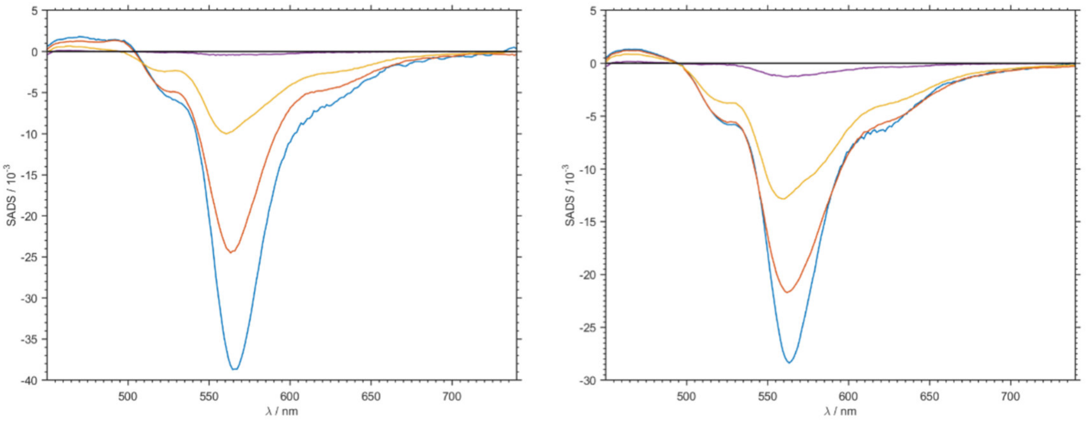

**Supplementary Figure 14. Species associated difference spectra (SADS) of *o*-TzR (left) and *m*-TzR (right).** First species (blue), second species (orange), third species (yellow) and residual (purple).

### 7 Computational methods

All DFT and TD-DFT calculations were carried out using Gaussian 16A.[20] Optimization of geometric structures were carried out at M062X/DEF2svp level.[21] Solvation effects (in water) were taken into account using SMD model.[22] Frequency calculations were performed to confirm that we obtained stable structures without imaginary vibration frequencies.

### 8 In cellulo reaction studies

**Molecular cloning of H2B-EGFP-HaloTag**

H2B-EGFP-HaloTag was cloned from H2B-EGFP (Addgene plasmid #11680)[23] and TOMM20-mCherry-HaloTag[24]. Plasmid was linearized by PCR using flanked primers (H2B-EGFP-fw: 5’-TAAAGCGGCCGCGACTCTAG-3’; H2B-EGFP-rv: 5’-CTTGTACAGCTCGTCCATGC-3’). PCR reaction mix of backbone fragments were digested with DPN1 (addition of 10 µL CutSmart Buffer + 1 µL DPN1 to 50 µL PCR mix, incubation at 37 °C for 1 h) to remove template vector. HaloTag fragment was amplified with appropriate Gibson overlaps using following primers: HALO-fw: 5’-TGGACGAGCTGTACAAGGAAATGAAGGTTTCTCTCGAGCC-3’ and HALO-rv: 5’-GTCGCGGCC-GCTTTAACCGGAAATCTCCAGAGTAGAC-3’. Backbone and HALO fragment were purified with preparative agarose gel, extracted with QIAquick Gel Extraction Kit (Qiagen) and assembled in a 1:1 molar ratio by Gibson assembly. Sequencing was performed at Sequence Laboratories Göttingen (Seqlab, Germany) using standard primer (EGFP-for).

**Cell culture**

COS-7 cells (obtained from ATCC) were cultured in full growth medium at 37 °C and under humidified conditions. The growth medium consisted of Dulbecco’s modified eagle medium (DMEM) supplemented with 10% fetal calf serum, 1 mM L-glutamine, 1 mM sodium pyruvate (all Gibco, United States). Every 2-3 days, cultures were subcultured into fresh petri dishes by enzymatically detaching cells using Trypsin-like enzyme (TrypLE, Gibco). Cultures were kept for 20-30 passages.

**Sample preparation**

For epi-fluorescence imaging, 8-well LabTek chambered coverslips (Thermo Fisher, United States) were prepared by cleaning with 0.1 M hydrofluoric acid followed by extensive washing using phosphate buffered saline (PBS, Sigma Aldrich, Germany) and DMEM growth medium. COS-7 wild-type cells were seeded into pre-cleaned LabTek dishes 48 h prior to imaging. Transient transfections with H2B-GFP-HaloTag was performed 24 h post seeding and 20-24 h prior to staining using TransIT-X2 (Mirus Bio, United States) according to instructions by the manufacturer.

Before imaging, transfected cells were loaded with a HaloTag linkers (HTL-BCN or HTL-TCO* at 10 µM) in starvation medium (DMEM FluoBrite supplemented with 1 mM L-glutamine and 1 mM sodium pyruvate, all Gibco) for 30 min at 37 °C. For mock sample any HTL-linker was omitted. Cells were washed four times (1 min, 15 min, 30 min, 15 min) with full growth medium and incubated with **HD555** at 1 µm in starvation medium for 120 min. Prior to imaging labeling solution was replaced with L-15 buffer.

**Automated epi-fluorescence microscopy**

Imaging was performed on an automated inverted fluorescence microscope. A Nikon Ti-E microscope body was equipped with a fibre-coupled multi-wavelength laser light source (iChrome MLE-LFA, Toptica, with 405, 488 and 640 nm diode and 561 nm DPPS lasers). The laser beam was reflected by a quad-band dichroic filter (Di01-R405/488/561/635, Semrock) into the sample through an oil immersion objective (CFI Apo TIRF 100x Oil, 1.49 NA, Nikon, combined with a 1.5x tube lens). Emission light was collected by the objective and directed through a notch filter (TIRF-Quad filter set 405/488/561/640, AHF Analysentechnik), a motorized filter wheel (FW102C, Thorlabs) equipped with bandpass filters (525/25 ET, 605/70 ET or 685/70, AHF Analysentechnik) and an image splitter device (Optosplit III, Cairn) onto an emCCD camera (iXon Ultra 897, Andor). The laser source was synchronized with the camera TTL output with a microcontroller board (Arduino Uno) routing the signal to the four individual laser channels. Samples were mounted on a motorized XY-stage (Marzhäuser) and focused with a motorized nosepiece equipped with a auto-focus system (PFS, Nikon). The microscope was controlled by open-source microscopy software Micro-Manager (version 1.4)[25] and operated through the graphical user interface and custom beanshell scripts. Two-color imaging in rhodamine channel (20 ms exposure, 1.1 mW laser power, emGain off) and GFP channel (20 ms exposure, 0.15 mW laser power, emGain off) was performed automatically after manual selection of cell positions from an overview image in the GFP channel. Recorded image sequences were averaged over 5 frames for analysis. For image analysis in Fiji (version 2.0.0, ImageJ version 1.52)[26] using custom macro scripts a nucleus segmentation was performed in the GFP channel by applying a median filtering with 10 px kernel size and a triangle thresholding. Mean intensities were measured in both fluorescence channels inside the segmented area. A total of ~150 cells per condition was pooled from three experiments prepared and imaged independently.

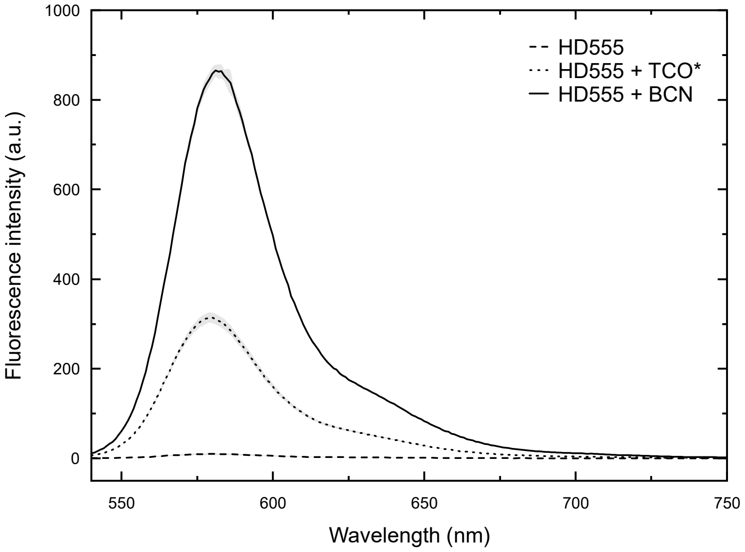

**Supplementary Figure 15. Emission intensity achieved with dienophiles BCN and TCO*.** Fluorescence emission spectra of **HD555** (5 µM in PBS, $\lambda_{ex}$ = 520 nm) before (dashed line) and after reaction with 20 eq of TCO* (dotted line) or BCN (solid line) averaged from three independent samples, standard deviation indicated with gray shaded bands.

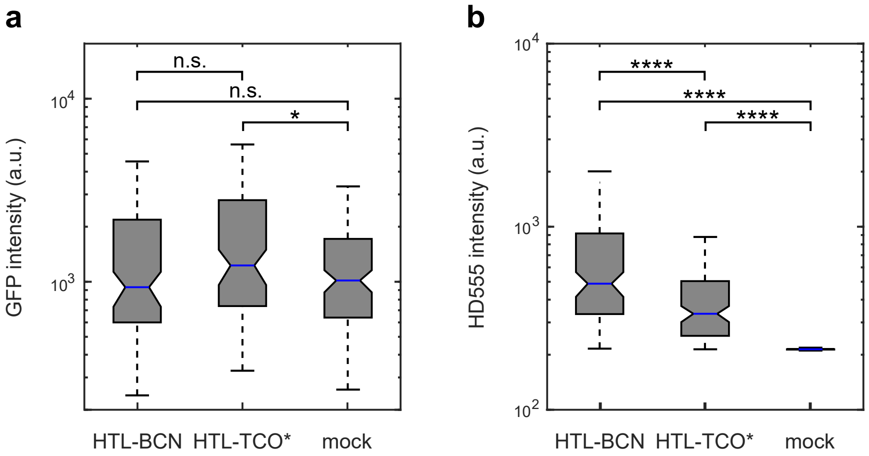

**Supplementary Figure 16. In cellulo fluorescence intensity quantification.** COS-7 cells expressing H2A-GFP-HaloTag labeled with **HD555** *via* HTL-BCN and HTL-TCO*, respectively. For a mock control, any HaloTag ligand was omitted. Intensity of nucleus area in GFP channel (a) and **HD555** channel (b) of 151 (HTL-BCN), 145 (HTL-TCO*) and 147 (mock) cells pooled from three independent measurements. Box plots: blue horizontal line: median, notches: 95 %-confidence of the median, box limits: 1st and 3rd quartile, whiskers: 1.5-fold interquartile range. Asterisks indicate significance from Wilkoxon rank sum test ($p > 0.05$ (n.s.), $p < 0.05$ (*), $p < 0.01$ (**), $p < 0.001$ (***) or $p < 0.0001$ (****)).

### 9 Confocal fluorescence microscopy

**Cell culture**

COS-7 cells (obtained from ATCC) were cultured in full growth medium at 37 °C and under humidified conditions. The growth medium consisted of Dulbecco's modified eagle medium (DMEM) supplemented with 10% fetal calf serum, 1 mM L-glutamine, 1 mM sodium pyruvate (all Gibco, United States). Every 2-3 days, cultures were subcultured into fresh petri dishes by enzymatically detaching cells using Trypsin-like enzyme (TrypLE, Gibco). Cultures were kept for 20-30 passages.

**Sample preparation**

For confocal imaging, 8-well LabTek chambered coverslips (Thermo Fisher, United States) were prepared by cleaning with 0.1 M hydrofluoric acid followed by extensive washing using phosphate buffered saline (PBS, Sigma Aldrich, Germany) and DMEM growth medium. COS-7 wild-type cells were seeded into pre-cleaned LabTek dishes 48 h prior to imaging. Transient transfections were performed 24 h post seeding and 20-24 h prior to staining using TransIT-X2 (Mirus Bio, United States) according to instructions by the manufacturer. Cloning of expression plasmid H2A-HaloTag is described in Hauke *et al.*[24]; HaloTag-A2A-adenosine receptor was a gift from Catherine Berlot (Addgene plasmid #66995; http://n2t.net/addgene:66995; RRID:Addgene_66995).

Before imaging, transfected cells were loaded with a HaloTag ligand-bicyclononyne linker (HTL-BCN) by incubation with HTL-BCN at 10 µM in full growth medium for at least 30 min at 37 °C. Cells were then washed twice with starvation medium (DMEM supplemented with 1 mM L-glutamine and 1 mM sodium pyruvate). For straightforward identification of transfected cells, samples were additionally labeled with 50 nM of a cell permeable, fluorescent HaloTag ligand (HTL-R110d, Promega, Germany) for 30 min at 37 °C. Following HTL-R110d labeling, cells were washed repeatedly with starvation medium at 37 °C to remove unbound HTL-BCN and HTL-R110d. Cells were transferred to the microscope immediately after washing.

Tetrazine-conjugated dyes were diluted from DMSO-dissolved stocks to 2x final concentration in starvation medium directly before addition to samples. Diluted dyes were added to samples during imaging by addition of one volume dye solution to one volume of medium on cells resulting in final dye concentrations of 1-5 µM. Samples were kept on microscope for less than 2 h for all experiments.

**Confocal fluorescence microscopy**

Imaging was performed on a Nikon A1R confocal laser scanning microscope operated by the Nikon Imaging Center at Heidelberg University. The microscope was equipped with a 60x Apo lambdaS NA 1.4 oil immersion objective, a galvanometric scanner and a perfect focus system (all Nikon, Japan). Samples were kept at 37 °C and 5% $CO_2$ during imaging using a stage-top incubation chamber (Tokai Hit, Shizuoka, Japan). Samples were illuminated with 483, 558 and 632 nm solid state lasers (Nikon). Emitted light was separated from excitation light using a 405/488/561/640 nm quadband dichroic mirror and further filtered using bandpass 515/30, 595/50 and 700/75 nm bandpass filters for each spectral channel. GaAsP detectors were used in the 483 and 558 nm excitation channels and a PMT detector was used for the 632 nm excitation channel. Images were acquired with a pixel size of 110 nm, with 2.5 µs pixel dwell time, 2x line averaging and a pinhole size of 34.5 µm. Multi-channel images were acquired in the line-interleaved mode. Z-stack were acquired at 0.5-1.0 µm spacing with 3-5 steps per position. Nikon Elements was used to control image acquisition and all connected devices.

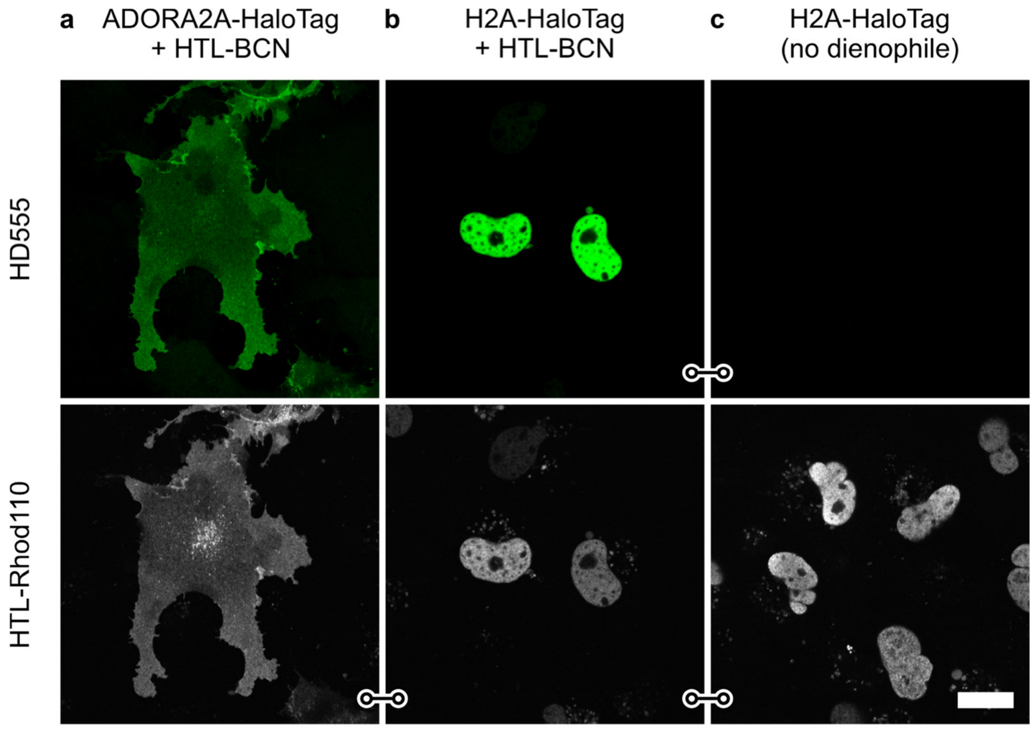

**Supplementary Figure 17. Live-cell no-wash labeling of intra- and extracellular targets with HD555.** COS-7 cells expressing membrane receptor fusion construct ADORA2A-HaloTag (a) or nuclear histone construct H2A-HaloTag (b,c) loaded with HTL-BCN and incubated with **HD555** at 1 µM for 30 min (top, green). Reference staining with chase labeling using HTL-Rhodamine 110 (bottom, gray). For evaluation of unspecific staining H2A-HaloTag expressing cells were labeled under omission of HTL-BCN (c) and imaged under identical conditions. Identical image and display for linked images. Scale bar 20 µm.

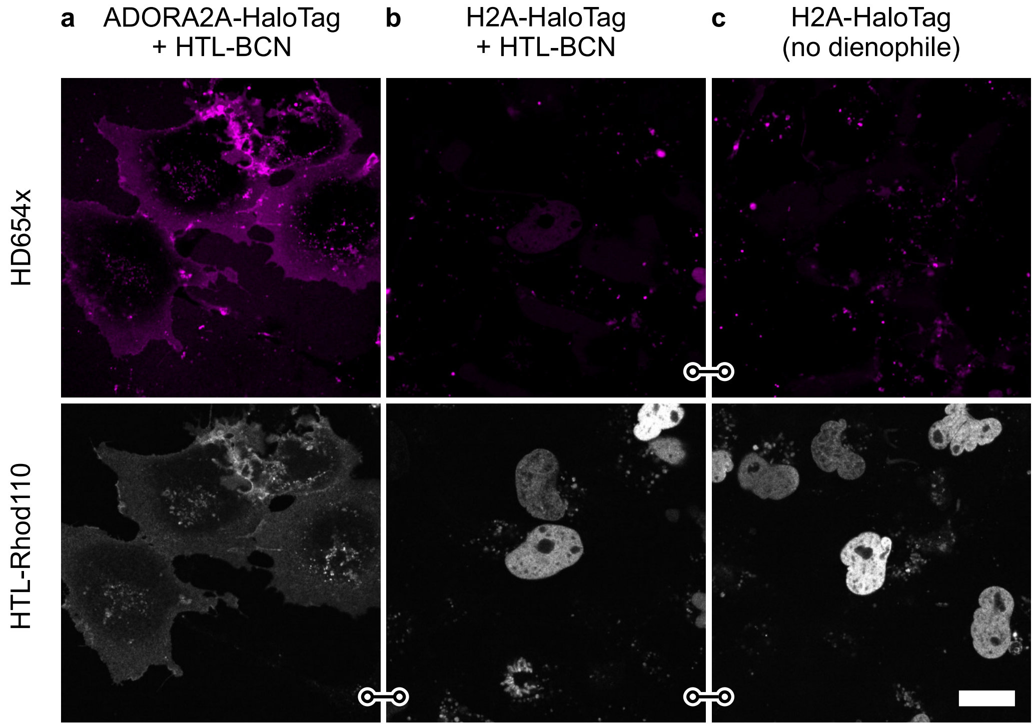

**Supplementary Figure 18. Live-cell no-wash labeling of intra- and extracellular targets with HD654x.** COS-7 cells expressing membrane receptor fusion construct ADORA2A-HaloTag (a) or nuclear histone construct H2A-HaloTag (b,c) loaded with HTL-BCN and incubated with **HD654x** at 1 µM for 30 min (top, magenta). Reference staining with chase labeling using HTL-Rhodamine 110 (bottom, gray). For evaluation of unspecific staining H2A-HaloTag expressing cells were labeled under omission of HTL-BCN (c) and imaged under identical conditions. Identical image and display for linked images. Scale bar 20 µm.

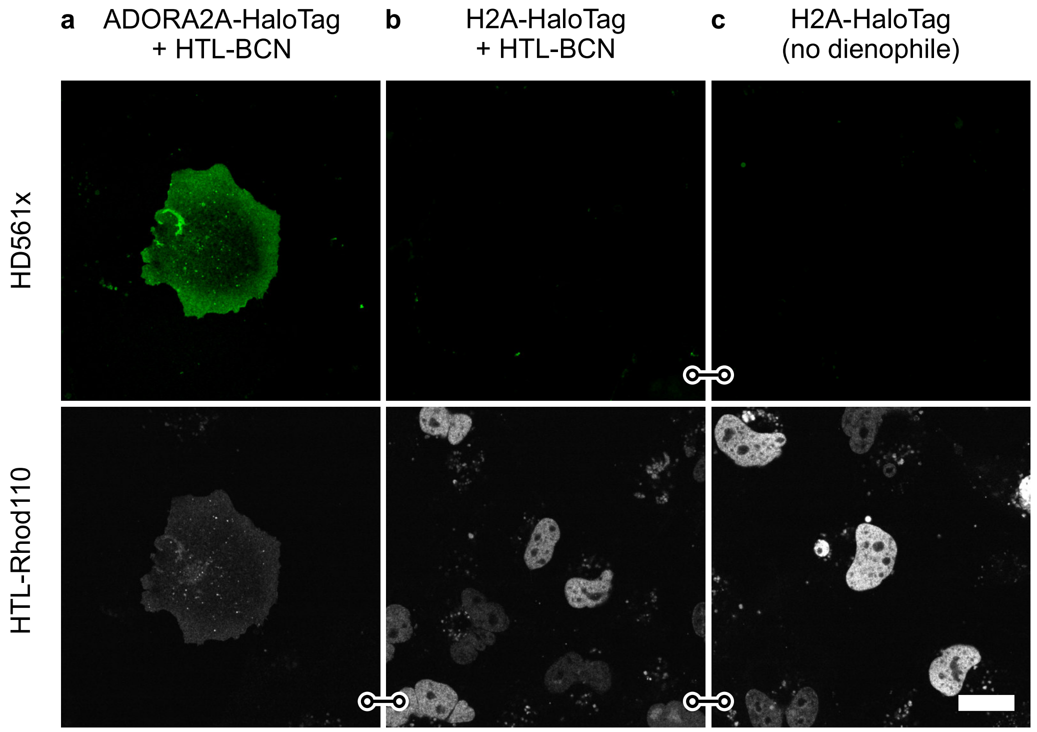

**Supplementary Figure 19. Live-cell no-wash labeling of intra- and extracellular targets with HD561x.** COS-7 cells expressing membrane receptor fusion construct ADORA2A-HaloTag (a) or nuclear histone construct H2A-HaloTag (b,c) loaded with HTL-BCN and incubated with **HD561x** at 1 µM for 30 min (top, green). Reference staining with chase labeling using HTL-Rhodamine 110 (bottom, gray). For evaluation of unspecific staining H2A-HaloTag expressing cells were labeled under omission of HTL-BCN (c) and imaged under identical conditions. Identical image and display for linked images. Scale bar 20 µm.

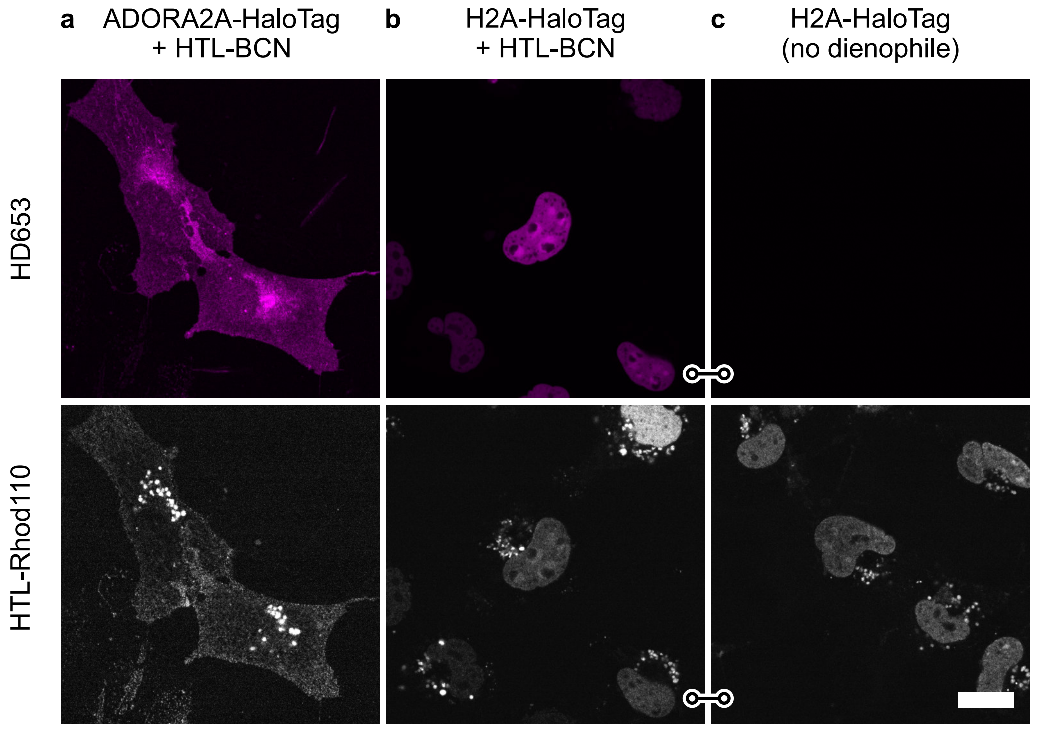

**Supplementary Figure 20. Live-cell no-wash labeling of intra- and extracellular targets with HD653.** COS-7 cells expressing membrane receptor fusion construct ADORA2A-HaloTag (a) or nuclear histone construct H2A-HaloTag (b,c) loaded with HTL-BCN and incubated with **HD653** at 1 µM for 30 min (top, magenta). Reference staining with chase labeling using HTL-Rhodamine 110 (bottom, gray). For evaluation of unspecific staining H2A-HaloTag expressing cells were labeled under omission of HTL-BCN (c) and imaged under identical conditions. Identical image and display for linked images. Scale bar 20 µm.

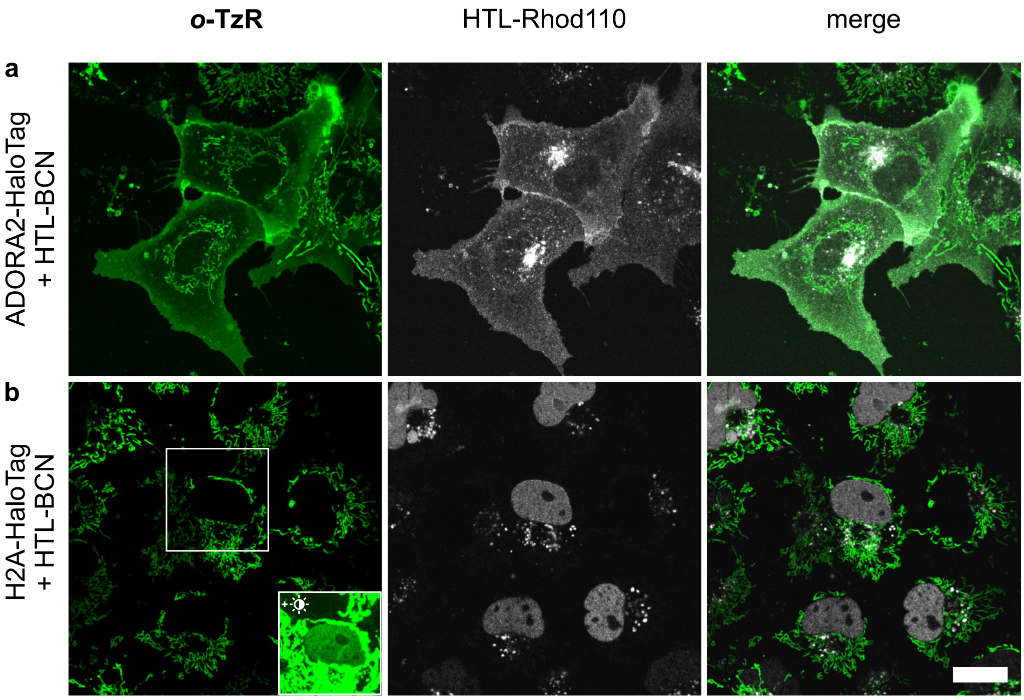

**Supplementary Figure 21. Live-cell no-wash labeling of intra- and extracellular targets with *o*-TzR.** COS-7 cells expressing membrane receptor fusion construct ADORA2A-HaloTag (a) or nuclear histone construct H2A-HaloTag (b) loaded with HTL-BCN and incubated with ***o*-TzR** at 1 µM for 30 min (green). Reference staining with chase labeling using HTL-Rhodamine 110 (gray). Unspecific mitochondrial accumulation of the positively charged dye can dominate the images, specific staining can be observed at increased contrast settings (inset). Scale bar 20 µm.

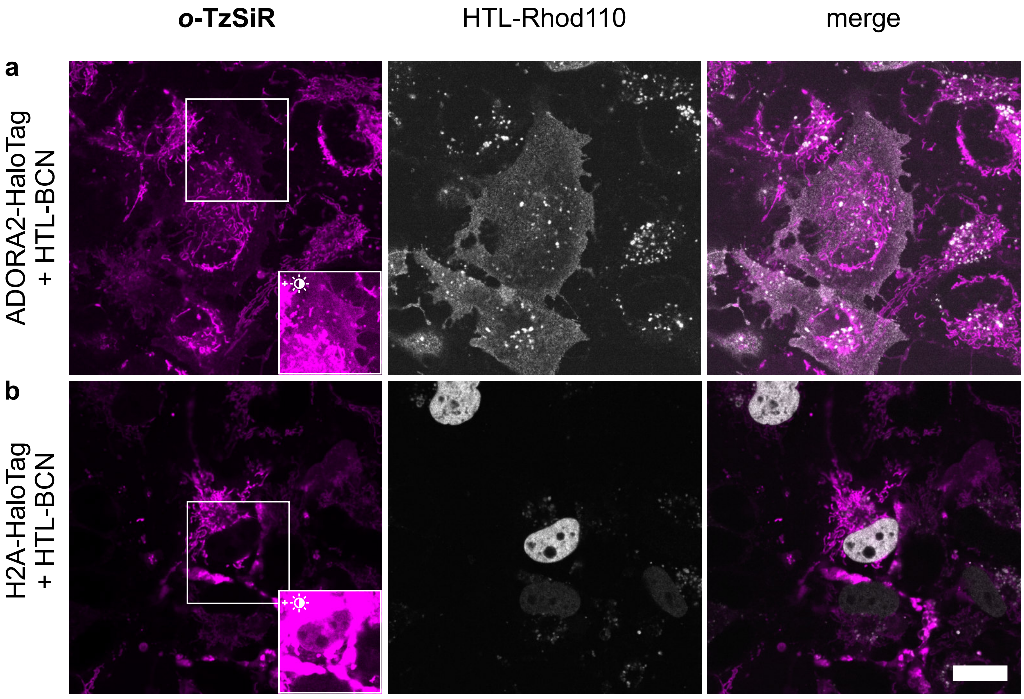

**Supplementary Figure 22. Live-cell no-wash labeling of intra- and extracellular targets with *o*-TzSiR.** COS-7 cells expressing membrane receptor fusion construct ADORA2A-HaloTag (a) or nuclear histone construct H2A-HaloTag (b) loaded with HTL-BCN and incubated with ***o*-TzSiR** at 1 µM for 30 min (magenta). Reference staining with chase labeling using HTL-Rhodamine 110 (gray). Unspecific mitochondrial accumulation of the positively charged dye can dominate the images, specific staining can be observed at increased contrast settings (insets). Scale bar 20 µm.

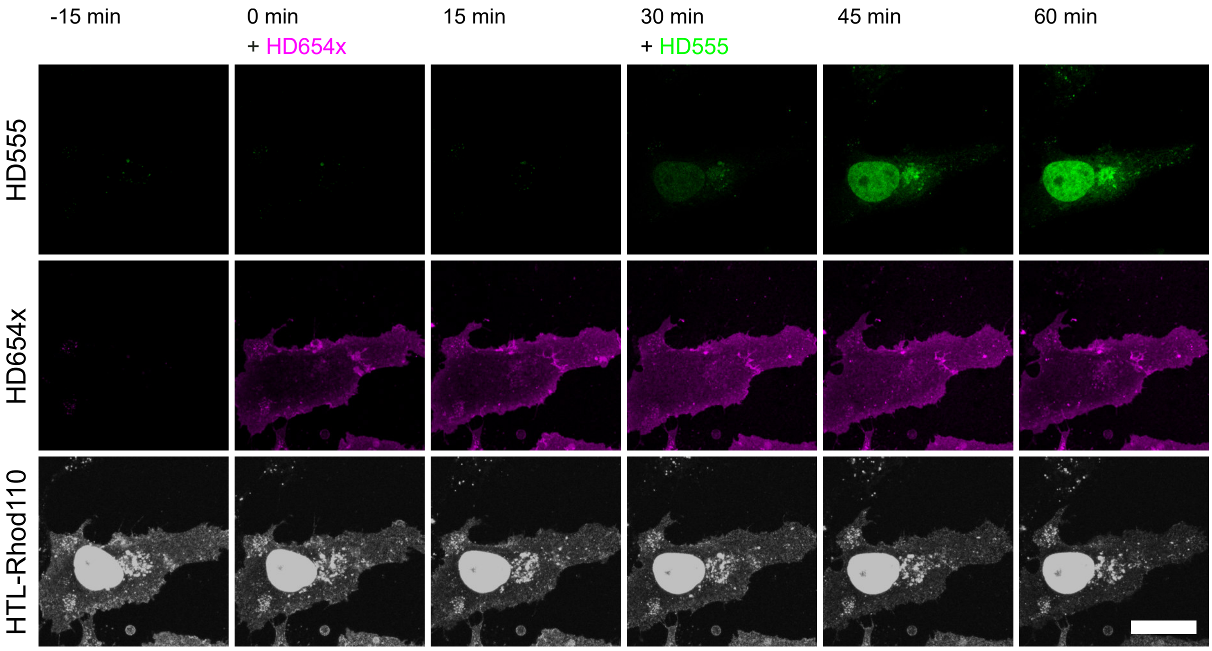

**Supplementary Figure 23. Dual target no-wash labeling.** COS-7 cells coexpressing H2A-HaloTag and ADORA2-HaloTag loaded with HTL-BCN were imaged during dual target labeling with **HD555** (green) and **HD654x** (magenta). Both structures were stained for reference with chase labeling using HTL-Rhod110 (gray). Cell impermeable dye **HD555** was added to yield a final concentration of 1 µM and incubated for 30 min when **HD654x** was added to yield a concentration of 1 µM without any buffer replacement or washing. The individual color channels show no cross labeling proving effective cell impermeability of **HD654x** and effective saturation of extracellular targets within 30 min. Scale bar 20 µm.

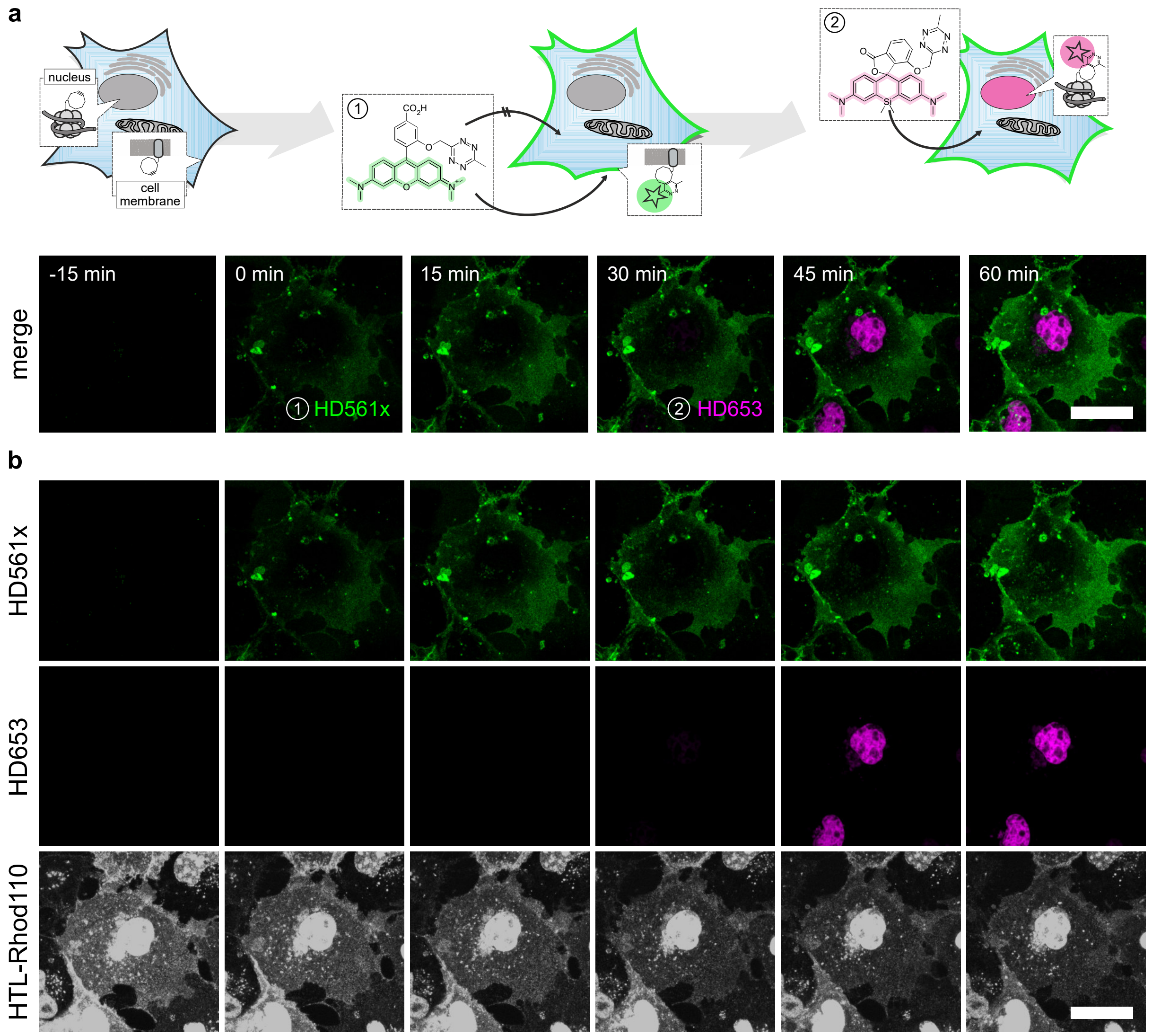

**Supplementary Figure 24. Dual target no-wash labeling with HD561x and HD653.** COS-7 cells coexpressing H2A-HaloTag and ADORA2-HaloTag loaded with HTL-BCN were imaged during dual target labeling with **HD561x** (green) and **HD653** (magenta). Cell impermeable dye **HD561x** was added to yield a final concentration of 1 µM and incubated for 30 min when **HD653** was added to yield a concentration of 1 µM without any buffer replacement or washing. a) Labeling strategy and two-color merge for intra-/extracellular labeling where targets and spectral features are interchanged compared to **HD555** and **HD654x**. b) The individual color channels show no cross labeling proving effective cell impermeability of **HD561x** and effective saturation of both targets within 30 min. Both structures were stained for reference with chase labeling using HTL-Rhod110 (gray). Scale bars 20 µm.

### 10 Unnatural amino acid labeling and STED-microscopy

**Cell culture**

COS-7 cells (Sigma 87021302) were maintained in Dulbecco's modified Eagle's medium (Life Technologies 41965-039) supplemented with 1% penicillin-streptomycin (Sigma P0781), 1% L-Glutamine (Sigma G7513), 1% sodium pyruvate (Life Technologies 11360), and 9% FBS (Sigma F7524). Cells were cultured at 37°C in a 5% $CO_2$ atmosphere and passaged every 2–3 days up to 15–20 passages. Cells were seeded 15-20 h prior to transfection at a density resulting in 70-80% confluency at the time of transfection. For flow cytometry, cells were cultured and labeled on 24-well plates (Greiner Bio-One). For stimulated emission depletion (STED) imaging, four-well chambered LabTek #1.0 borosilicate coverglasses (ThermoFisher) were used.

**Constructs, cloning and mutagenesis**

For vimentin, mutagenesis was performed on a pVimentin–PSmOrange plasmid (Addgene plasmid #31922), and the TAG was introduced at position N116, generating the pVimentin$^{N116TAG}$–PSmOrange construct.[27] For Nup153, a pGFP-Nup153 plasmid was first constructed by cloning a codon-optimized Nup153 cDNA into a pEGFP backbone. The TAG codon was then introduced at position N149 of the GFP gene, generating the pGFP$^{N149TAG}$–Nup153 construct. For the expression of the Amber suppression system in mammalian cells, we used the pcDNA3.1-tRNAPyl/NES-PylRS$^{AF}$ plasmid, which contains the tRNA under control of a human U6 promoter and the synthetase under a CMV promoter.

**Transfection and labeling**

All transfections were performed with JetPrime reagent (PeqLab) according to the manufacturer's recommendations. For Amber suppression system experiments cells were transfected at a ratio of a 1:1 with a POITAG vector and tRNAPyl/NESPylRSAF vectors. After 4-6 h, the medium was exchanged to a fresh one containing 50 µM BOC (BOC-L-Lys-OH) or BCN (endo BCN-L-Lysine). After overnight incubation, the medium was exchanged to DMEM without serum and incubate for 1-2 h at 37 °C. For labeling, 500 nM **HD653** or **SiR-Tz** working solutions were prepared using DMEM without serum and immediately added to the cells.

After incubation for 30 min at 37 °C, the medium was exchanged to 1x PBS. The cells were directly imaged without additional washing steps.

**UAA imaging**

Confocal and STED images were acquired on a Leica SP8 STED 3X microscope using 488 nm (for eGFP), 594 nm (for mOrange) and 633 nm (for **HD653** or **SiR-Tz**) laser lines for excitation, and 775 nm STED laser at 75% maximum output power and a delay of -400 for depletion. Emission light was collected with HyD detectors at 500-550 nm and 640-750 nm for pGFP$^{N149TAG}$–Nup153, and 600-630 nm and 640-750 nm for pVimentin$^{N116TAG}$–PSmOrange, respectively. The images were taken using a Leica HC PL APO 100×/1.40 oil immersion objective. Sequential acquisition between lines was used to avoid crosstalk. Time-gated fluorescence detection was used for STED to further improve lateral resolution. For STED and confocal imaging of pVimentinN116TAG–PSmOrange structures, the pixel size was 37.9 nm and pixel dwell time was 1.2 µs, with 16x line averaging.

**UAA flow cytometry**

The transfected and freshly labeled COS-7 cells were harvested, resuspended in 1x PBS and passed through 100 µm nylon mesh. Data acquisition was performed in an LSRFortessa SORP Cell Analyzer (BD). At least 40.000 cells per condition were analyzed. Cells were first gated by cell type (using FSC-A x SSC-A parameters) and then by single cell (FSC-A x SSC-W). Lastly, fluorescence was acquired in the 488-530/30 channel for GFP signal and in the 640-730/45 channel for **HD653**/**SiR-Tz**. Acquired data was analyzed using custom-written Matlab (MathWorks) code. Cells were gated based on FSC-A and SSC-A signal. The GFP-expressing population was then selected for further analysis by thresholding the FSC-A corrected GFP signal. Signal to background ratios were calculated as $(I_{BCN} - I_{BOC})/I_{BOC}$; intensities were obtained by cell-wise correction for GFP expression strength according to $I = Counts_{Dye} / Counts_{GFP}$.

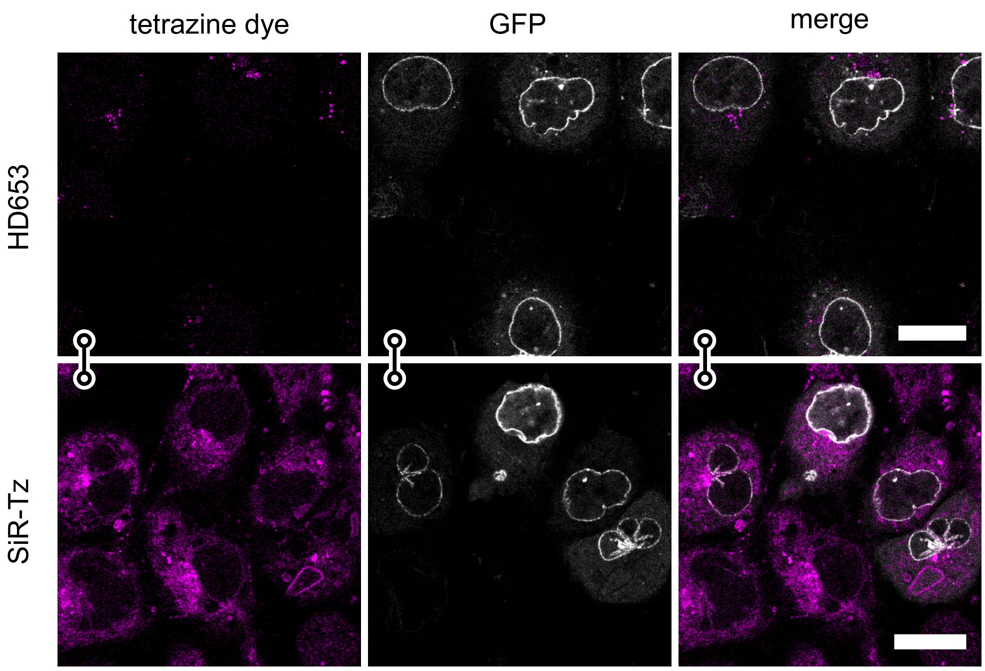

**Supplementary Figure 25. Unspecific background with unnatural amino acid labeling.** Confocal images of live COS-7 cells expressing pEGFP$^{N149TAG}$-Nup153 in the presence of BOC-L-Lys-OH incubated with **HD653** and **SiR-Tz**. Scale bar: 20 µm.

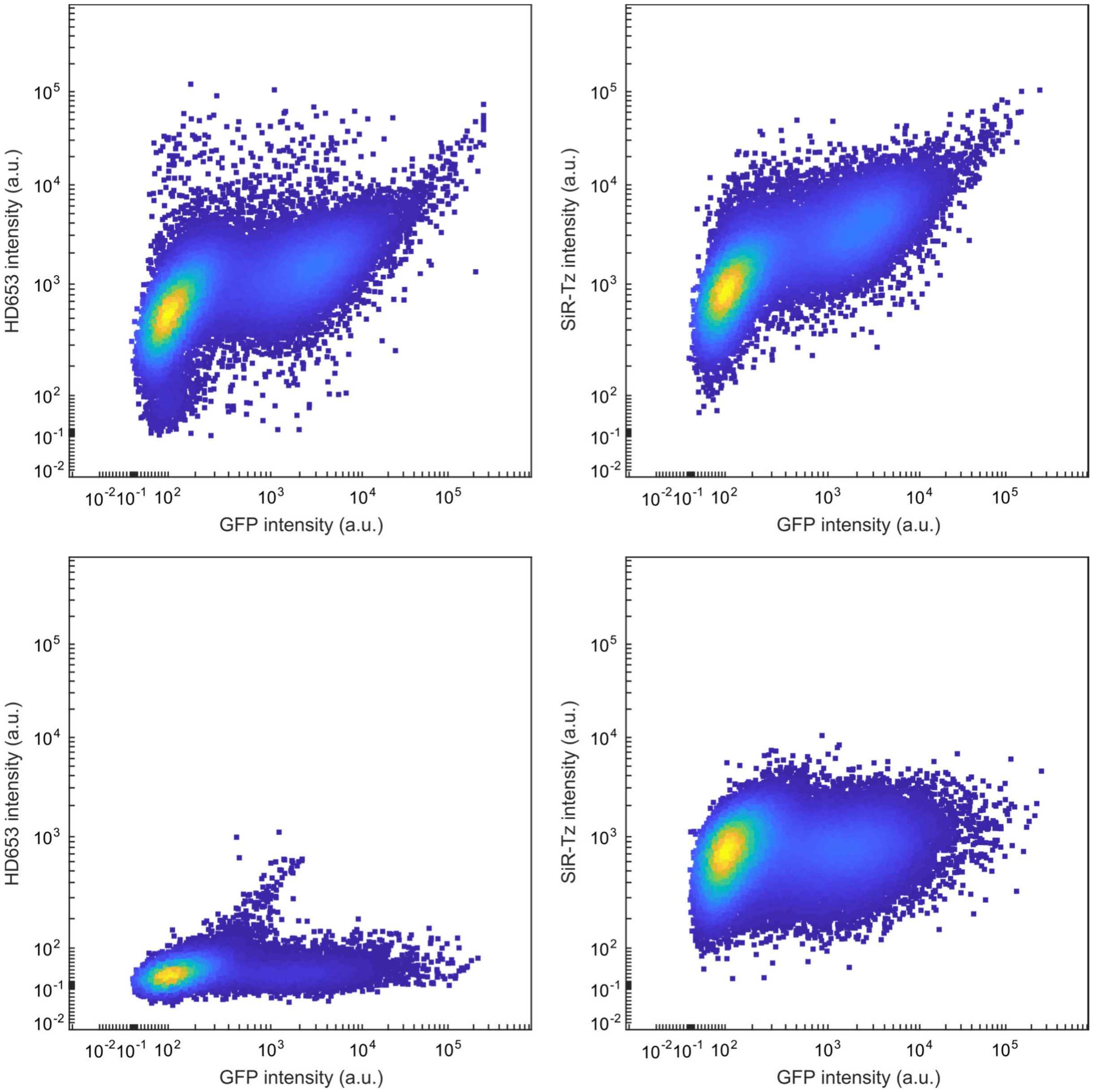

**Supplementary Figure 26. Flow cytometry analysis of HD653 and SiR-Tz labeling specificity.** COS-7 cells were transiently transfected with pGFPN149TAG–Nup153, loaded with endo-BCN-L-Lysine (top) or BOC-L-Lys-OH (bottom) and labeled with **HD653** (left) or SiR-Tz (right). At least 40,000 events per condition were recorded and gated to select for single cells (see methods).

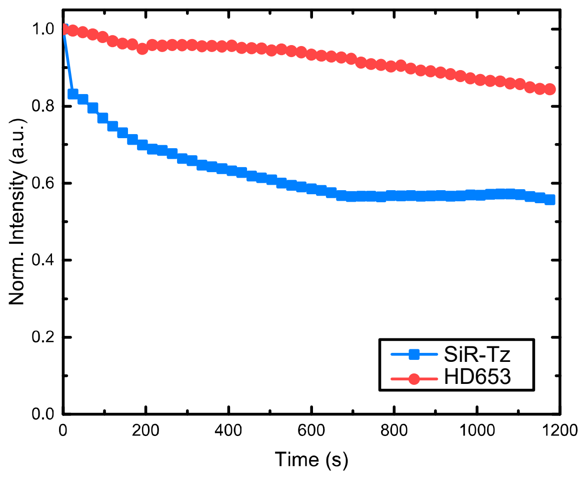

**Supplementary Figure 27. Signal loss in STED imaging.** Normalized average intensity in sequential STED images of live COS-7 cells expressing vimentin$^{N116TAG}$-mOrange construct with Lys-BCN as UAA labeled with **HD653** and **SiR-Tz**.

### 11 Super-resolution optical fluctuation imaging

**Cell culture**

COS-7 cells were cultured at 37 °C and 5 % $CO_2$ using DMEM high glucose without phenol red (4.5 g $l^{-1}$ glucose) supplemented with 4 mM L-gluthamine, 10 % fetal bovine serum and 1×penicillin-streptomycin (all gibco®, Thermo Fisher Scientific). Cells were seeded in Lab-tek® II chambered cover slides (nunc) or on 25 mm high-precision No. 1.5 borosilicate coverslips (Marienfeld) in 12 well plates (Thermo Fisher Scientific) 1-2 days before fixation or transfection in DMEM high glucose w/o phenol red.

**Filamentous actin staining**

Cells were washed once with growth medium w/o phenol red and incubated for 90 s in a buffer consisting of 80 mM PIPES, 7 mM $MgCl_2$, 1 mM EDTA, 150 mM NaCl and 5 mM D-glucose at a pH adjusted to 6.8 using KOH with 0.3% (v/v) Triton X-100 (AppliChem) and 0.25% (v/v) high-grade glutaraldehyde (Electron Microscopy Sciences). After 90 s solution was exchanged to pre-warmed 4% paraformaldehyde dissolved in PBS (pH = 7.4) and samples were incubated for 10 min at room temperature to ensure complete fixation. Afterwards, samples were washed thrice for 5 min with PBS on orbital shaker and kept for 5 min with a freshly prepared 10mM $NaBH_4$ solution in PBS on an orbital shaker in order to reduce background fluorescence. This step was followed by one quick wash with PBS, and two washes of 10 min with PBS on an orbital shaker. Samples then were additionally permeabilized with 0.1% (v/v) Triton X-100 in PBS (pH = 7.4) on an orbital shaker followed by an additional wash with PBS and then labeled with 250 nM phalloidin-BCN in labeling buffer consisting of 20 mM Tris-HCl, 150 mM NaCl, 2% BSA, 0.2% TX-100 (pH = 7.4) for 1 h at room temperature. Finally, samples were washed 3x for 5 min on an orbital shaker with a labelling buffer without BSA and subsequently incubated with 250 nM of **sb-HD656** in the labeling buffer without BSA for 1 h at 37 °C. A final wash with PBS (pH = 8) was done thrice on an orbital shaker and samples were imaged immediately after in the same buffer. 50 ms exposure time was used under 275 W/$cm^2$ 635 nm laser excitation and 8000 frames were typically acquired.

#### SOFI processing

The long image sequence for high-order super-resolution optical fluctuation imaging (SOFI) of actin was split into batches that were independently processed and averaged to mitigate the influence of photobleaching on SOFI. For all other data, the provided number of frames were directly analyzed. After cumulant calculation, the images were deconvolved and linearized similarly as described before.[28] For second order processing, the brightness of the images was linearized by taking the square root of the SOFI intensities.

#### Histone H2B and TOMM20 transfection and labeling

About 7000-15000 cells per well were seeded in Lab-tek® II chambered cover slides (nunc) and transfected with H2B-eGFP-HaloTag or TOMM20-mCherry-HaloTag (225 ng per well) using Lipofectamine 3000 according to manufacturer's instructions. About 20-24 h after transfection, cells were incubated with 10 µM HTL-BCN linker in starvation medium (no serum) for 30 minutes at 37 °C, followed by two washes, 2x15 min washes and 1x30 min wash all in full growth medium at 37 °C. Subsequently cells were labeled with 1-5 µM **sb-HD656** in starvation medium (no serum) at 37 °C for 30-60 minutes, again followed by two washes, 2x15 min washes and 1x30 min wash all in full growth medium at 37 °C. The medium was exchanged by HBSS w/ $MgCl_2$ and $CaCl_2$ and cells were immediately imaged at RT for up to 30 min or cells were fixed. Fixation was performed using 4% PFA in PBS pH 7.4 for 15 min, followed by 3x washes for 5 min in PBS. Samples were kept at 4 °C until imaging. 20 ms exposure time was used under 275 $W/cm^2$ 635 nm laser excitation and 1000 frames were analyzed with subsequent brightness linearization and deconvolution.

#### Microscope setup

Data was acquired on a home-built widefield microscope. Briefly, four illumination laser lines, i.e. 200 mW 405 nm laser (MLL-III-405-200mW), a 350 mW 561 nm laser (Gem561, Laser Quantum) and a 200 mW 488 nm laser (iBEAM-SMART-488-S-HP, Toptica Photonics) are collimated, expanded and combined with a flat fielded 1 W 635 nm laser (SD-635-HS-1W, Roithner Lasertechnik) using dichroic filters and focused in the back focal plane of the objective (Nikon SR Plan Apo IR 60× 1.27 NA WI). The beam of 635 nm laser is flat-fielded using a multimode fiber and a speckle reducer similarly as described previously.[29] The fluorescence light is filtered using a combination of a dichroic mirror, a multiband emission filter and a bandpass filter (Quad Line Beamsplitter R405/488/561/635 flat and Quad Line Laser Rejectionband ZET405/488/561/640, 685/70 ET Bandpass, all AHF Analysetechnik) and subsequently focused on an sCMOS camera (ORCA Flash 4.0, Hamamatsu; back projected pixel size of 108 nm). The sample position is controlled in X and Y by a Scan-plus IM 120x80 (Marzheuser) and in Z by a Nano-Z piezo nanopositioner (Mad City Labs). Z-stabilization was achieved with a PID controller using a 780 nm laser diode reflecting from the coverslip-sample interface. Custom written LabView Z-stabilization software was used for long imaging stacks (>4000 frames). All acquisitions were performed using Micromanager.[25]

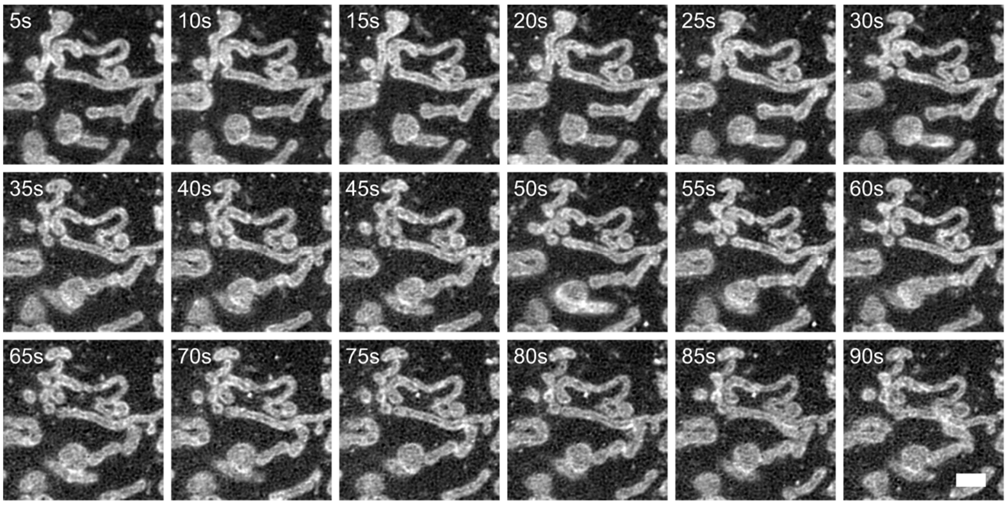

**Supplementary Figure 28. Time-course live-cell SOFI on mitochondria.** COS-7 cells transiently expressing TOMM20-mCherry-HaloTag were incubated with HTL-BCN and labeled with **sb-HD656**. Live-cell second order SOFI images corresponding to Figure 5c. Scale bar: 2 µM.

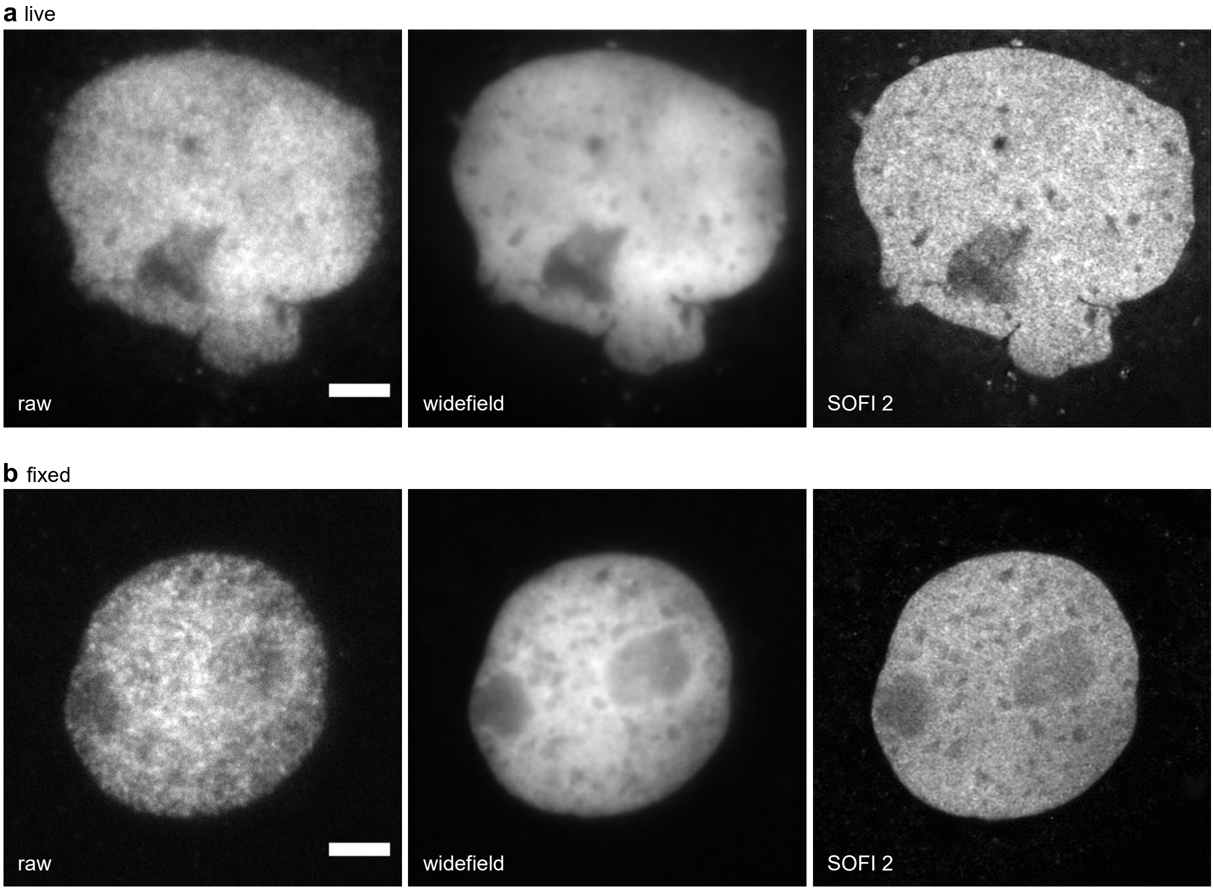

**Supplementary Figure 29. SOFI of histone H2B.** COS-7 cells transiently expressing H2B-eGFP-HaloTag were incubated with HTL-BCN and labeled with **sb-HD656**. First frame of the time series, widefield (average of the time series) and second order SOFI in (a) live and (b) fixed cells. Image acquisition: 1000 frames, 20 ms exposure time, 635 nm laser 275 W/cm$^2$. Second order SOFI can be performed even at high fluorophore densities, allowing analysis of cells with a large range of expression levels. The images are representative of 6 fixed cells from one experiment and of 14 live cells from two independent experiments. Scale bar: 5 µM.

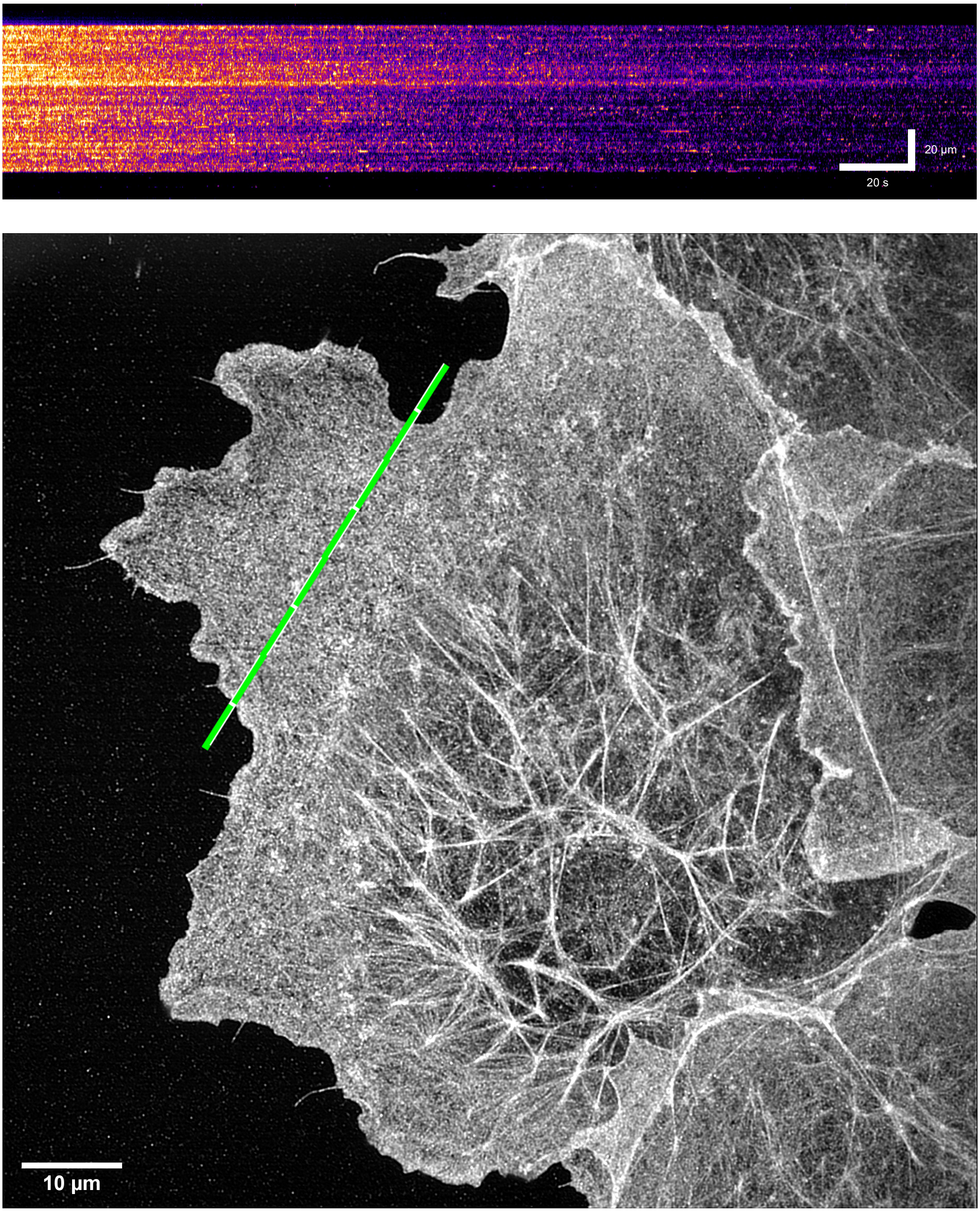

**Supplementary Figure 30. Kymograph indicating signal fluctuation over time.** Kymograph displayed was obtained from the time sequence of the first 5000 frames of phalloidin-**sb-HD656** labeled actin in COS-7 cells (same data as Figure 5d). The sequence was resliced by taking a line profile along the time axis, thus giving the information about the signal intensity over time. This qualitative measurement shows both the bleaching effect (which is shown in Supplementary Figure 31) and the temporal signal fluctuation in each pixel due to dye blinking. Cross section area is indicated in the image below.

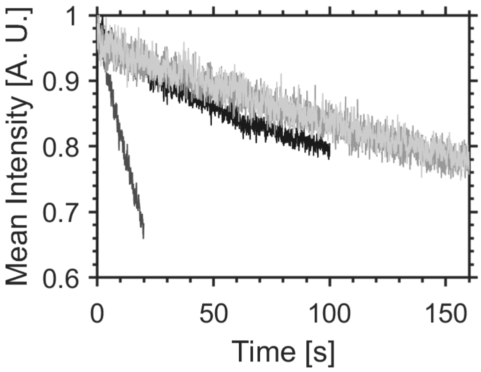

**Supplementary Figure 31. Photostability of sb-HD656 in SOFI of live and fixed cells.** Average intensity over time from a 50 x 50 pixel area of (part of) the data underlying the mitochondria images in Figure 5b,c (black), the histone H2B images in Supplementary Figure 29 a (dark grey) and b (grey) and the actin images in Figure 5d (light grey).

### 12 NMR spectra

$^{1}H$ NMR of **3** ($CDCl_3$)

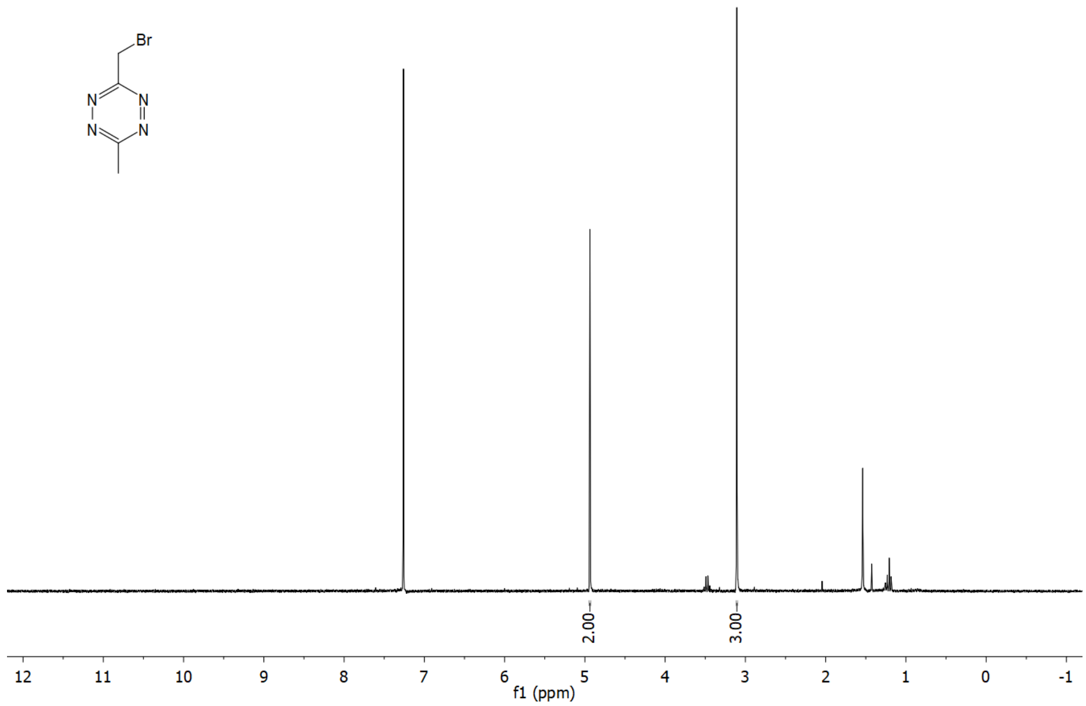

$^{13}C$ NMR (APT) of **3** ($CDCl_3$)

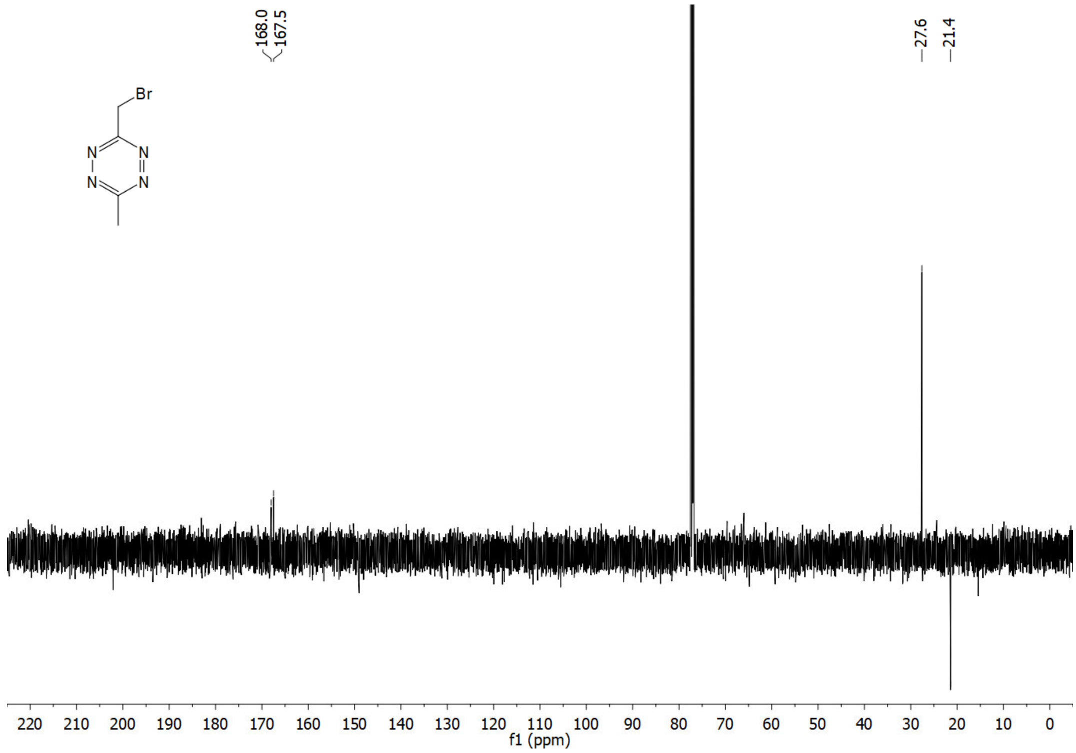

[1]H NMR of **4** ($CDCl_3$)

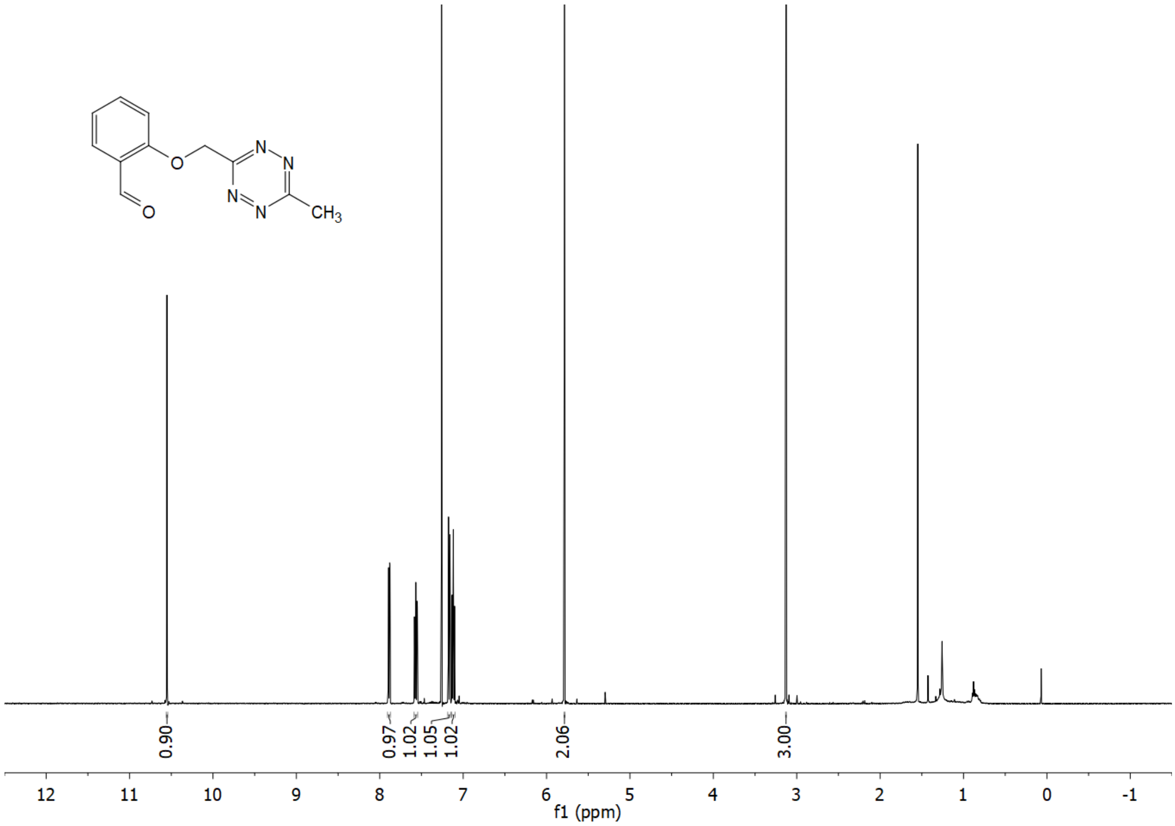

[13]C NMR (APT) of **4** ($CDCl_3$)

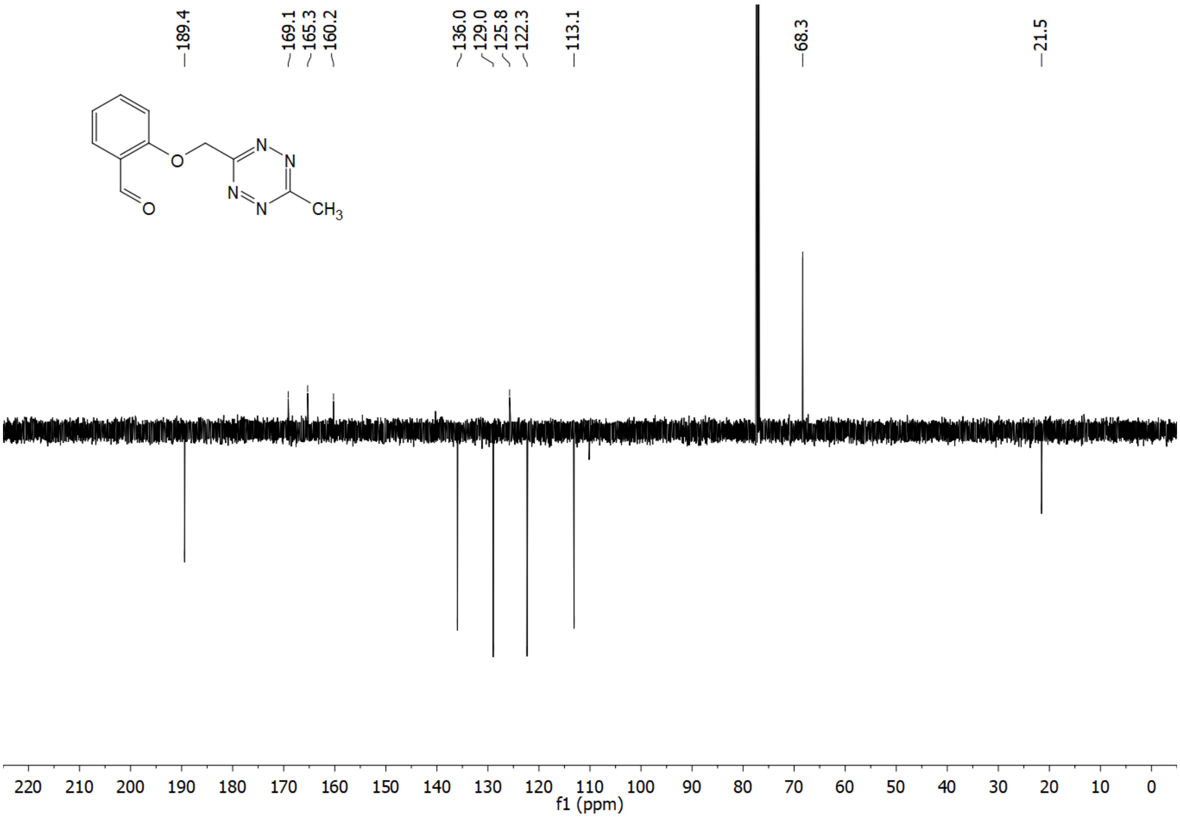

$^1H$ NMR of **5** ($CDCl_3$)

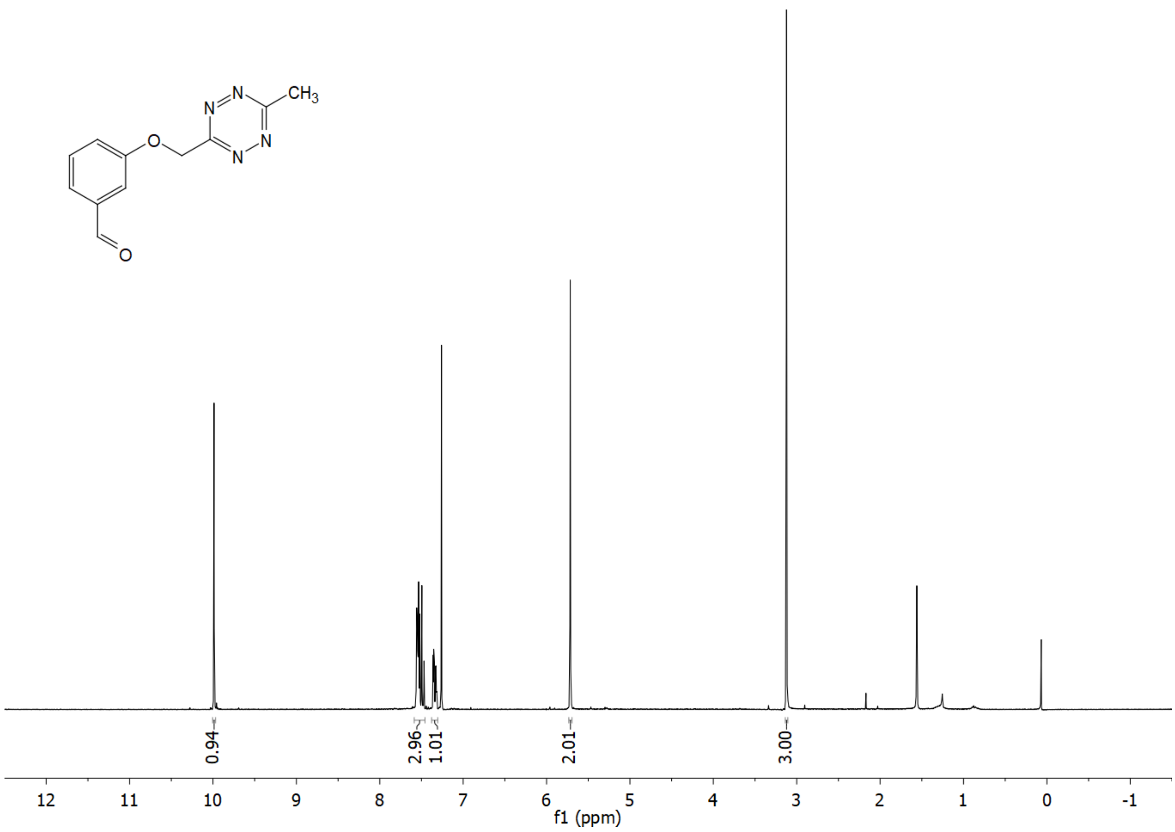

$^{13}C$ NMR (APT) of **5** ($CDCl_3$)

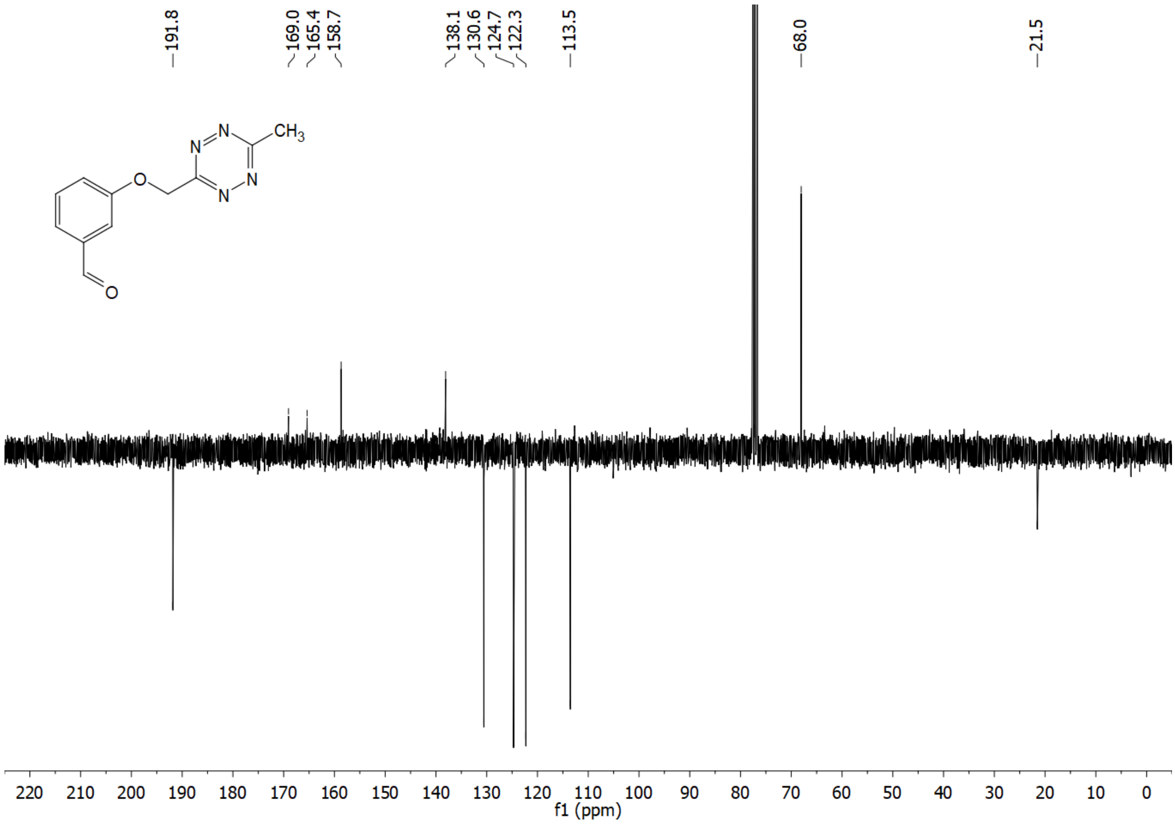

$^1H$ NMR of **6** ($CDCl_3$)

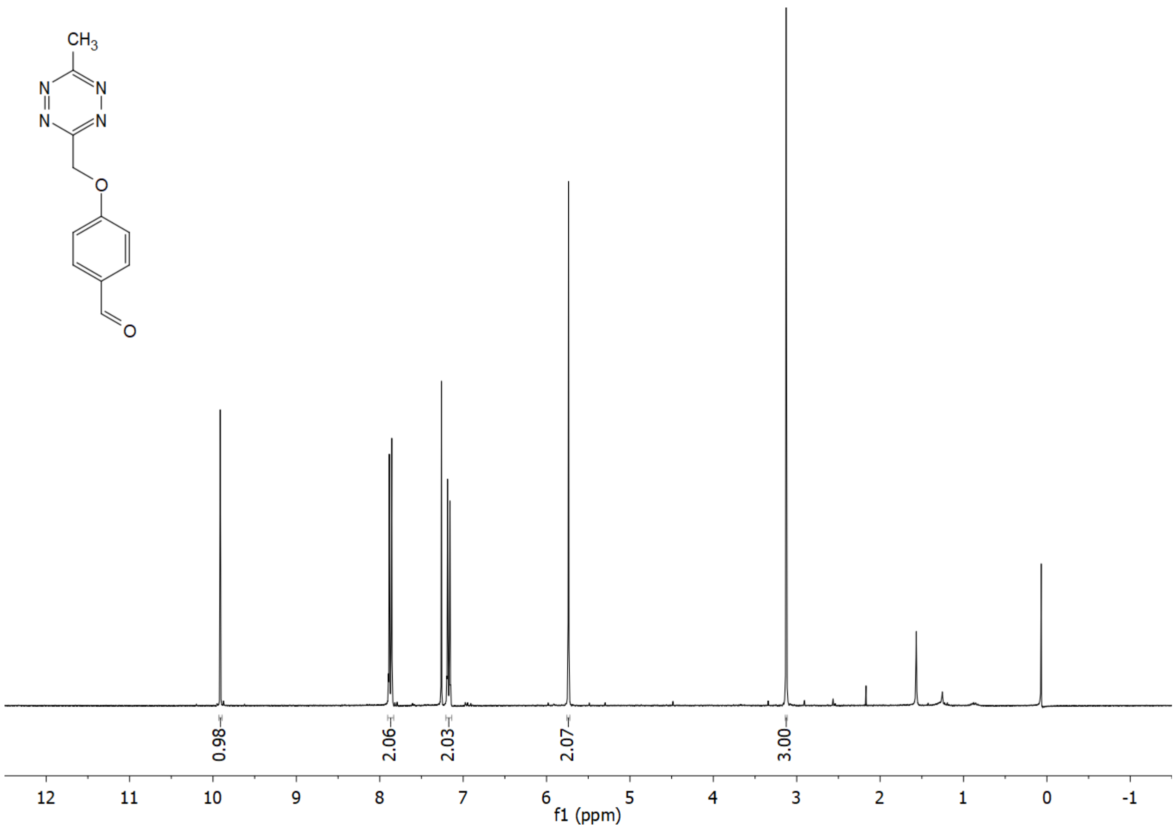

$^{13}C$ NMR (APT) of **6** ($CDCl_3$)

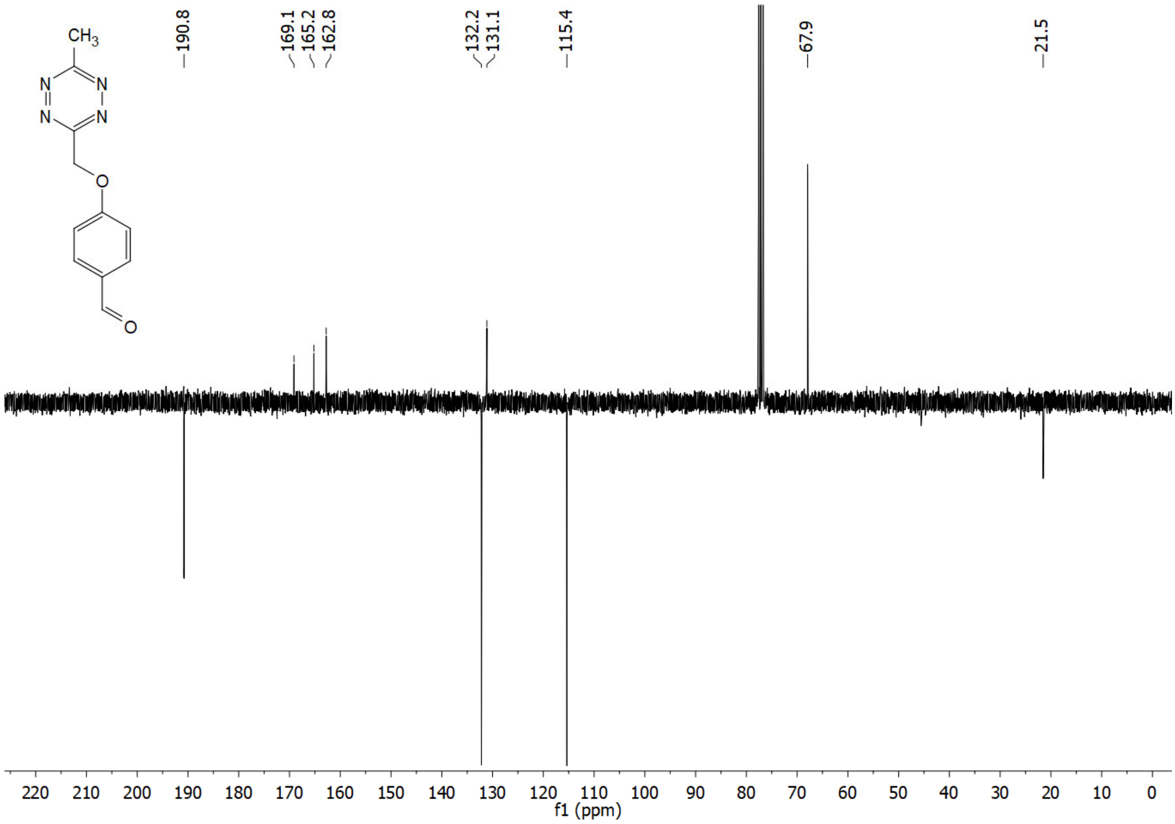

[1]H NMR of **7** ($CDCl_3$)

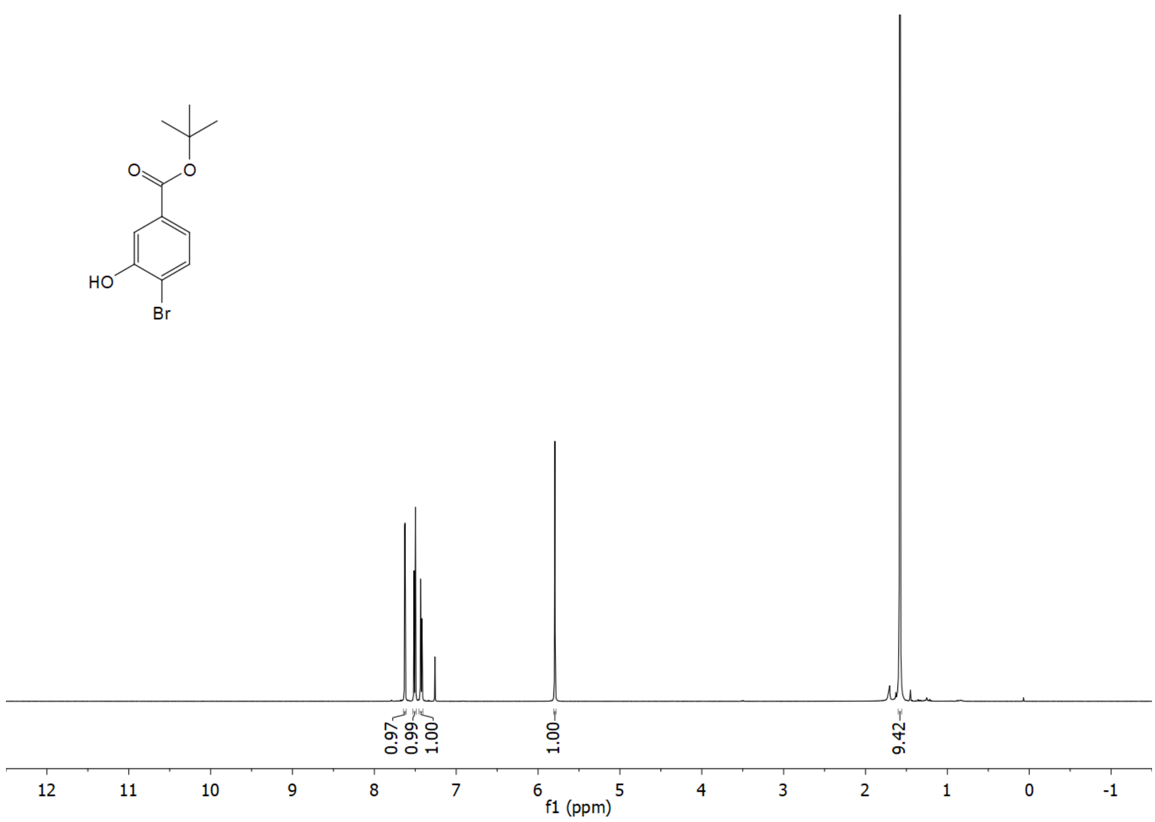

[13]C NMR (APT) of **7** ($CDCl_3$)

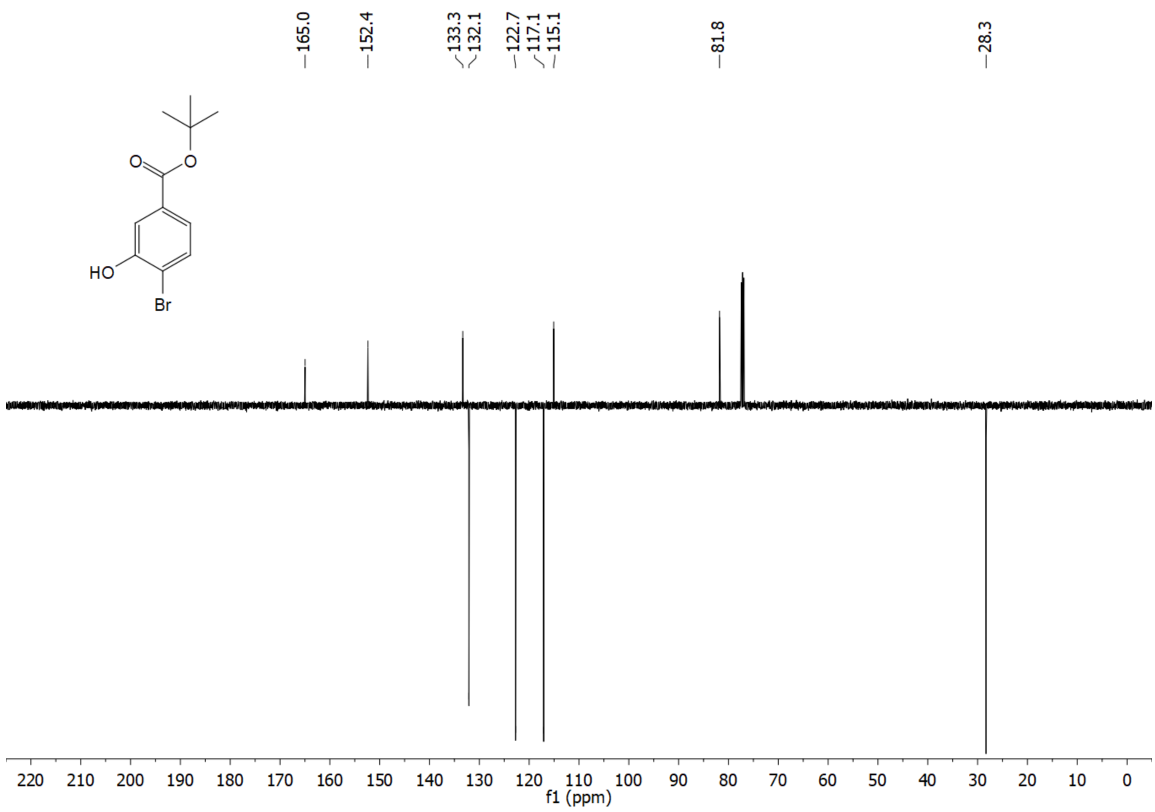

$^{1}H$ NMR of **8** ($CDCl_3$)

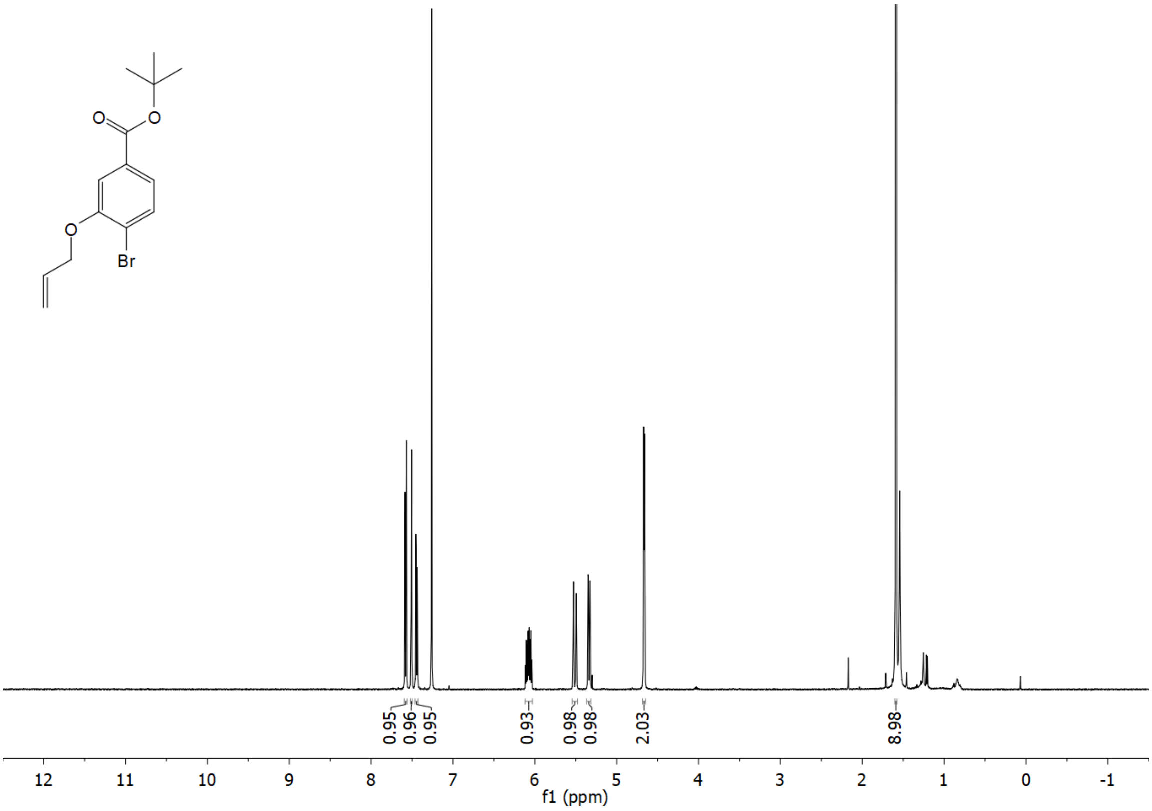

$^{13}C$ NMR (APT) of **8** ($CDCl_3$)

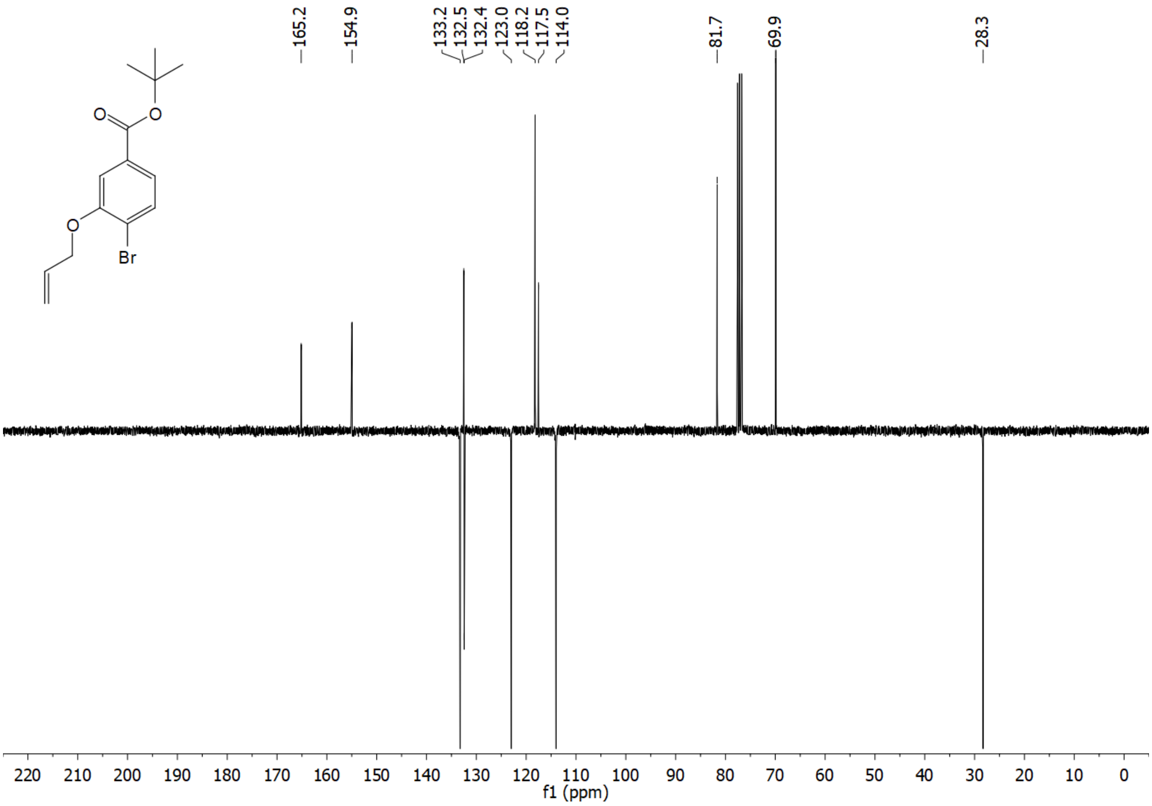

$^{1}$H NMR of **9** ($CDCl_3$)

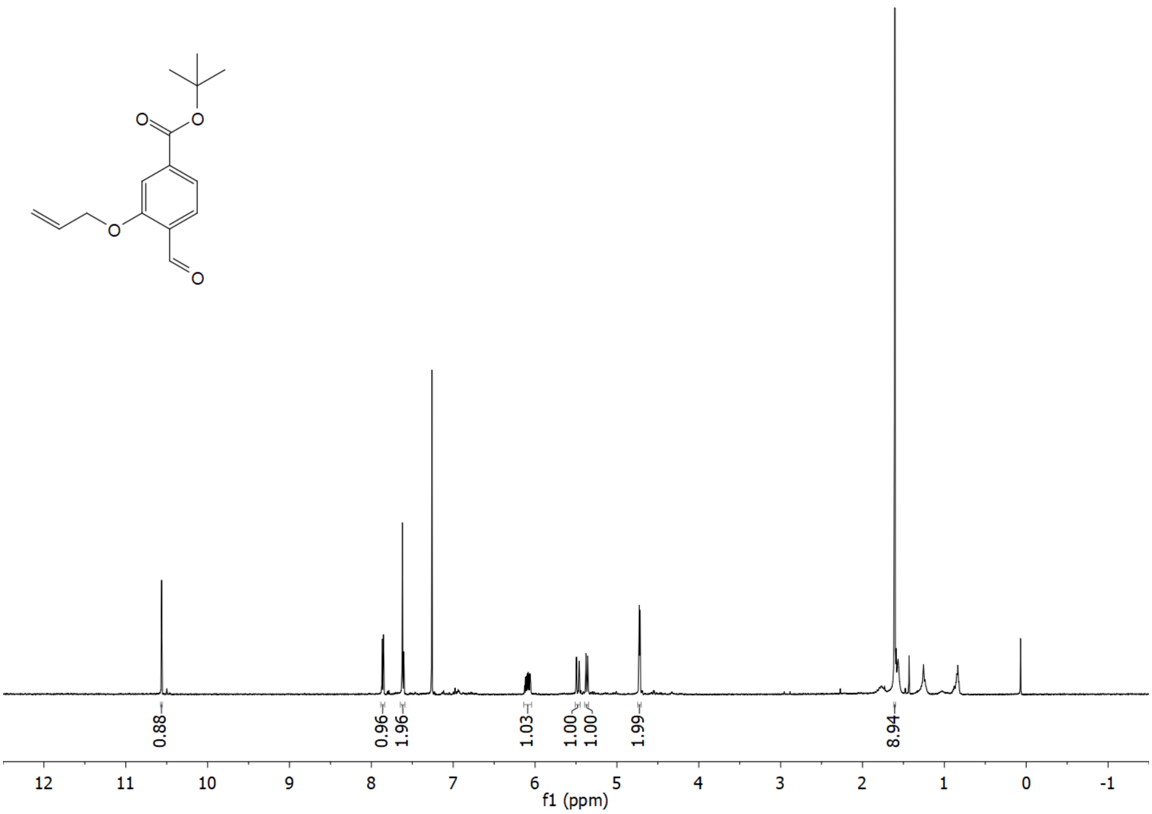

$^{13}$C NMR (APT) of **9** ($CDCl_3$)

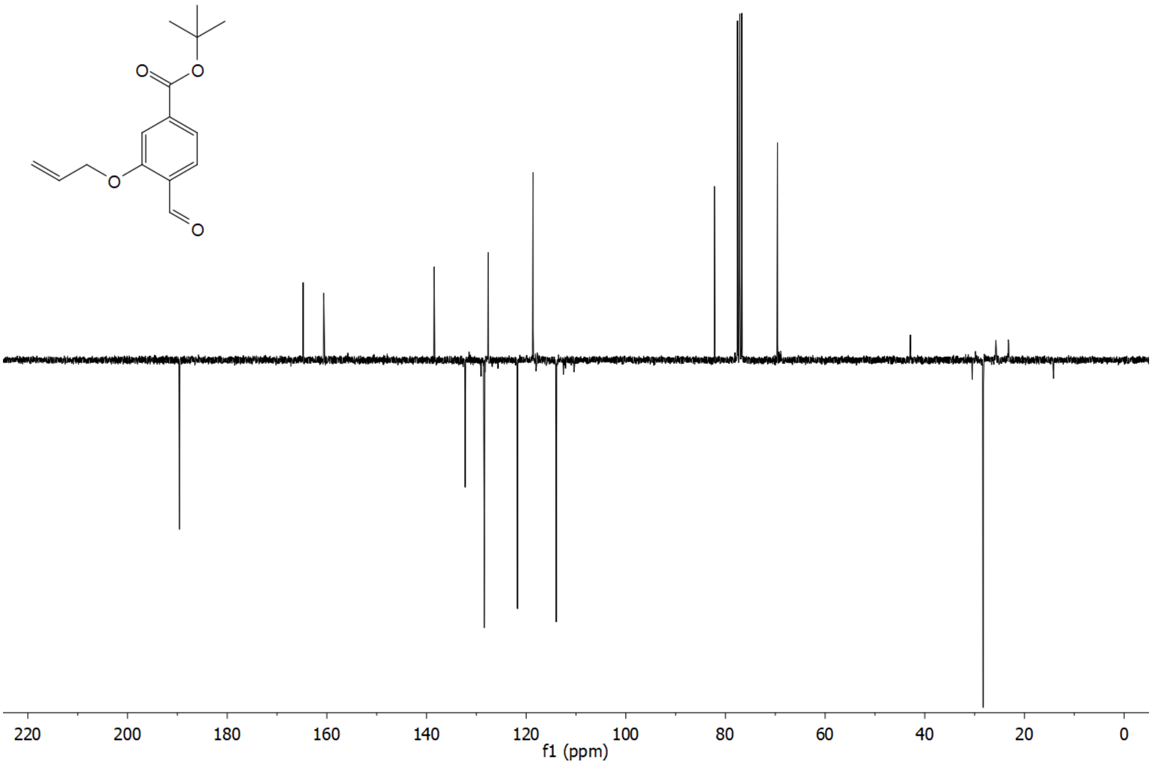

[1]H NMR of **10** ($CDCl_3$)

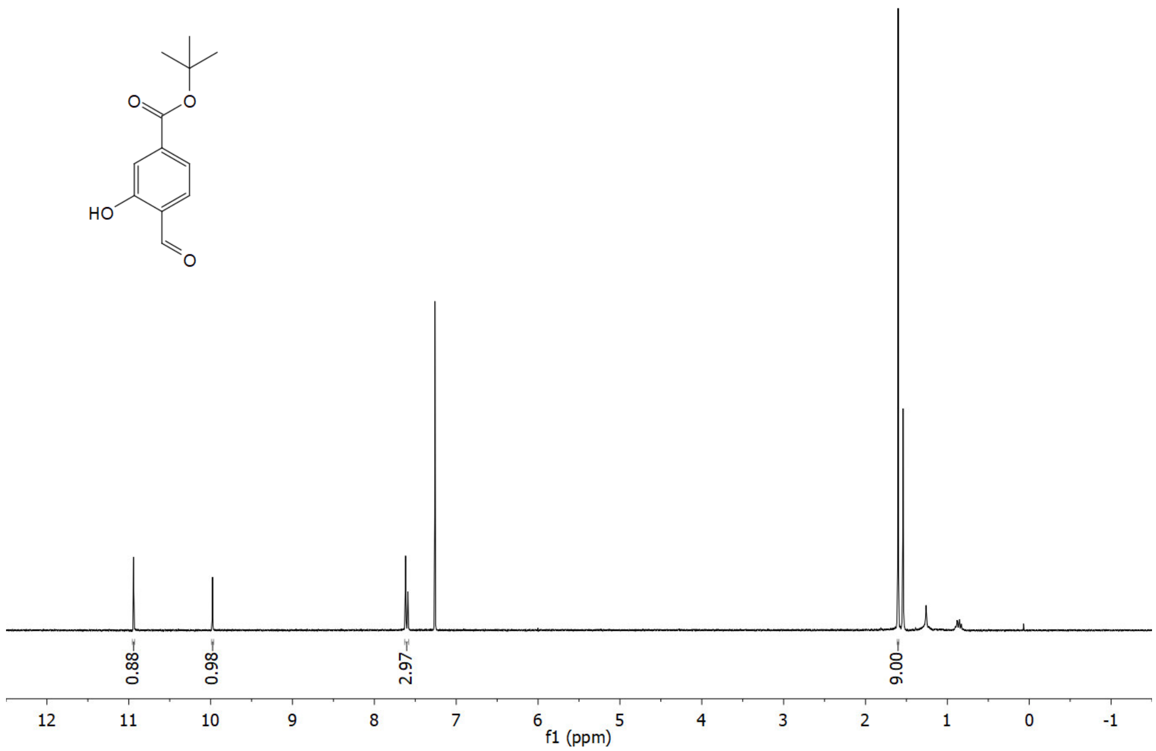

[13]C NMR (APT) of **10** ($CDCl_3$)

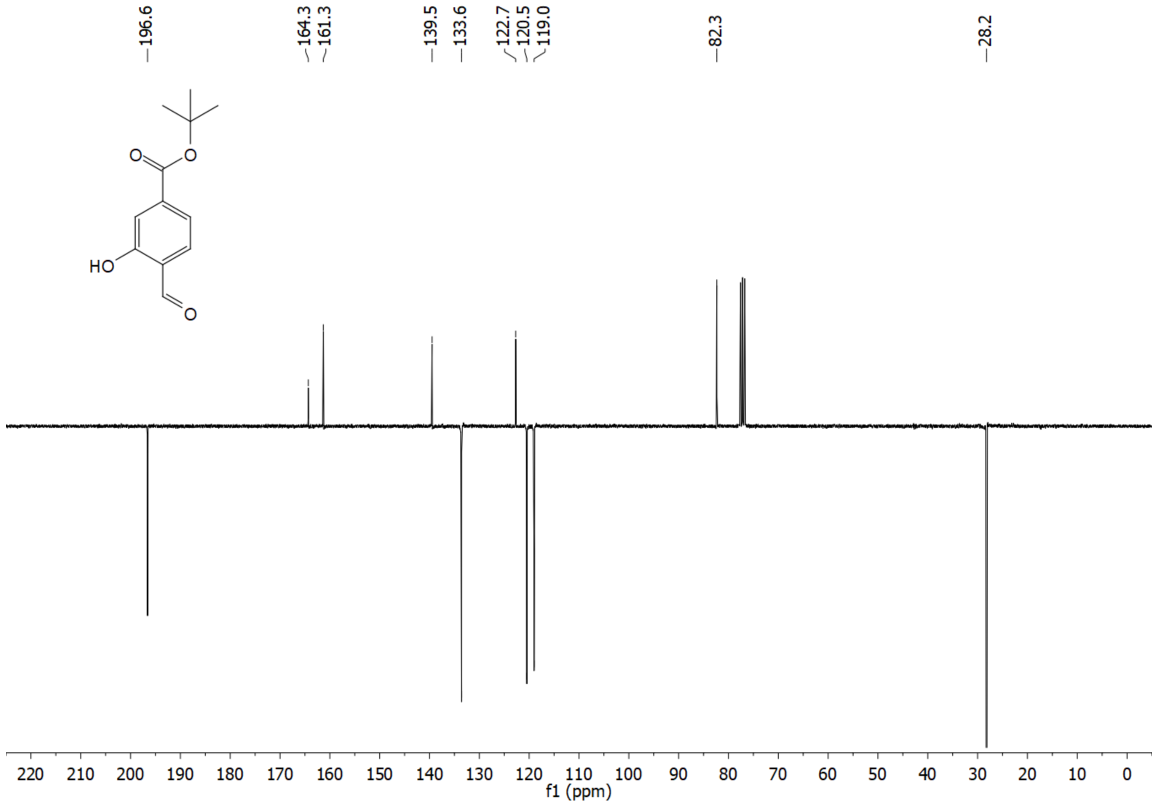

$^{1}H$ NMR of **11** ($CDCl_3$)

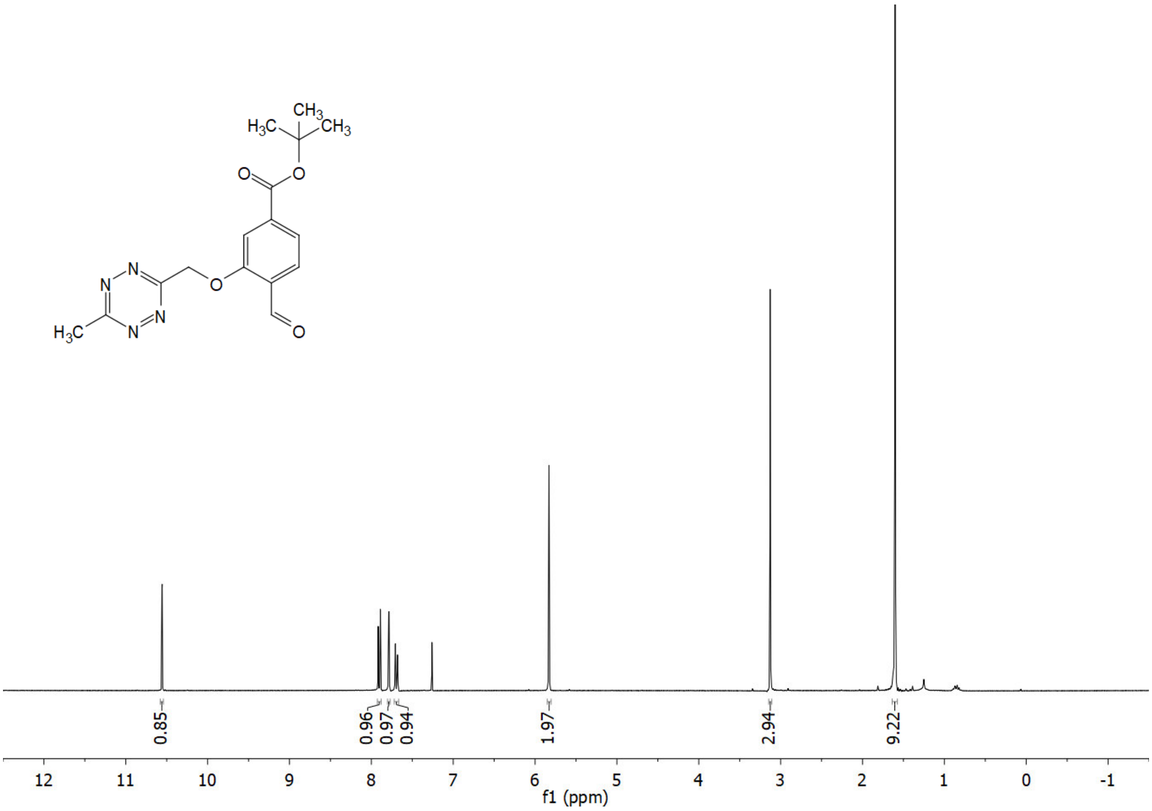

$^{13}C$ NMR (APT) of **11** ($CDCl_3$)

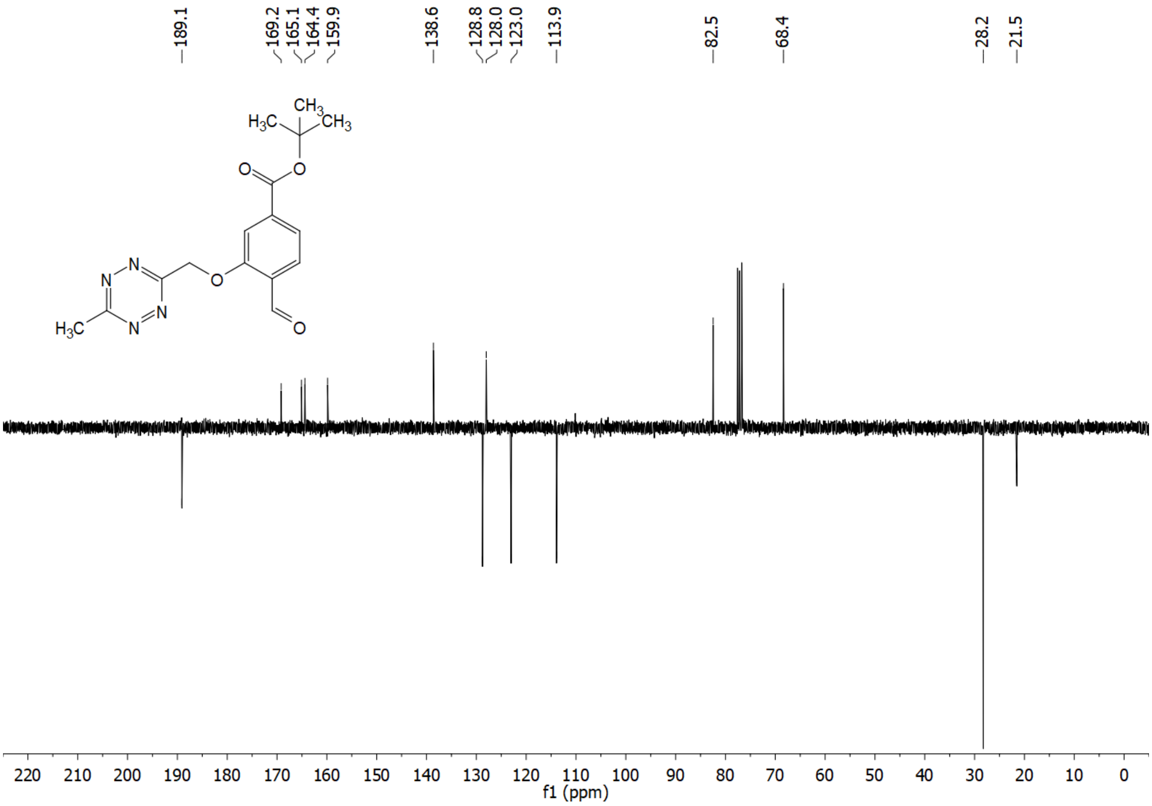

$^{1}H$ NMR of **12** ($CDCl_3$)

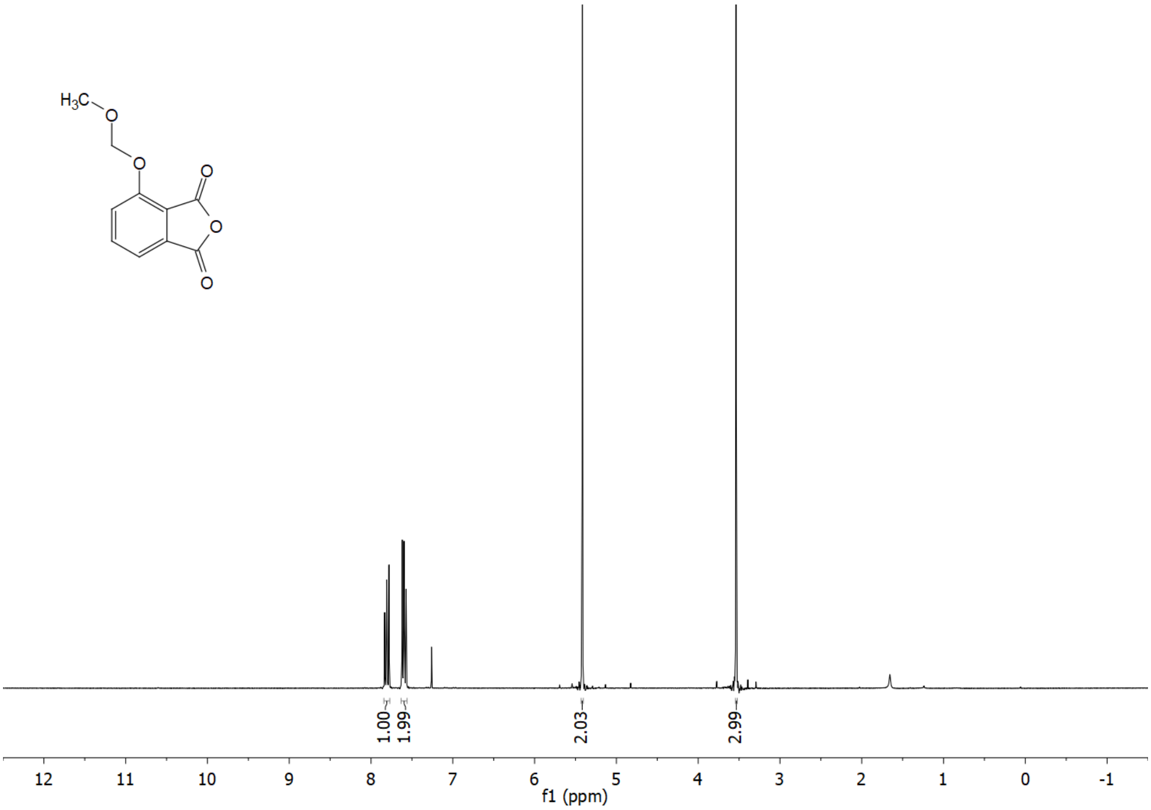

$^{13}C$ NMR (APT) of **12** ($CDCl_3$)

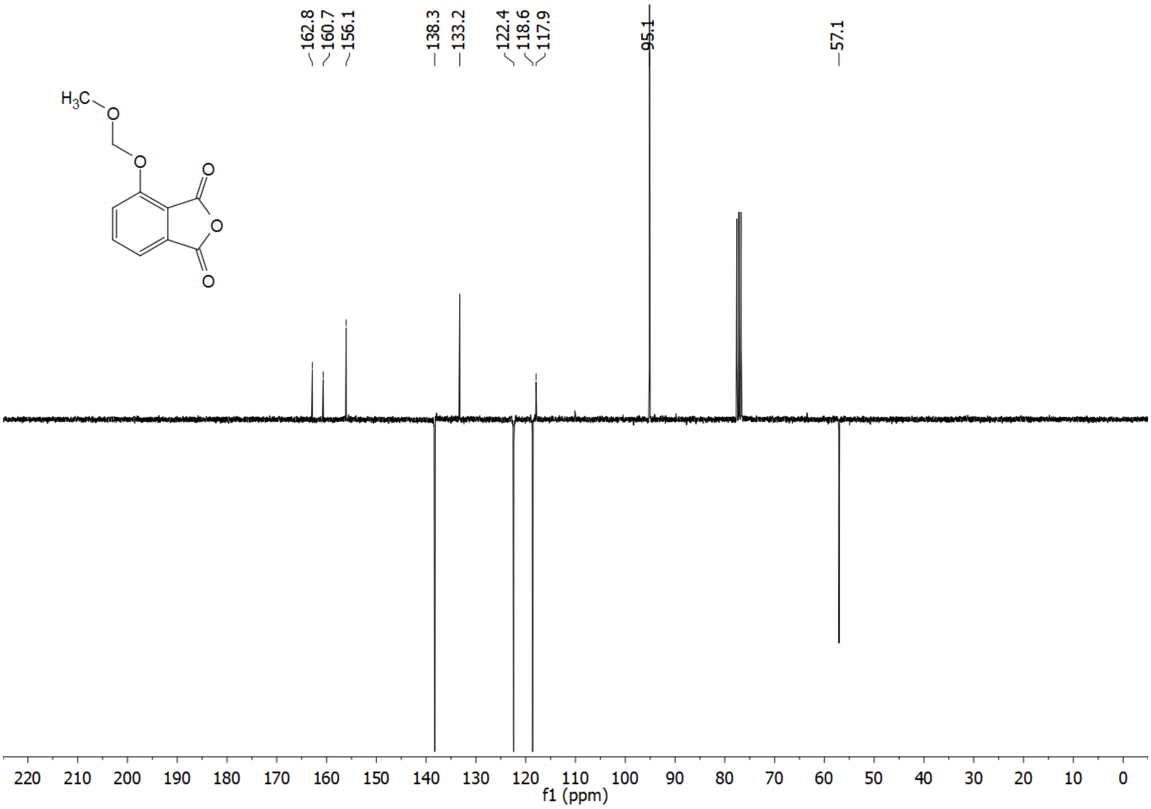

$^{1}$H NMR of **13** ($CDCl_3$)

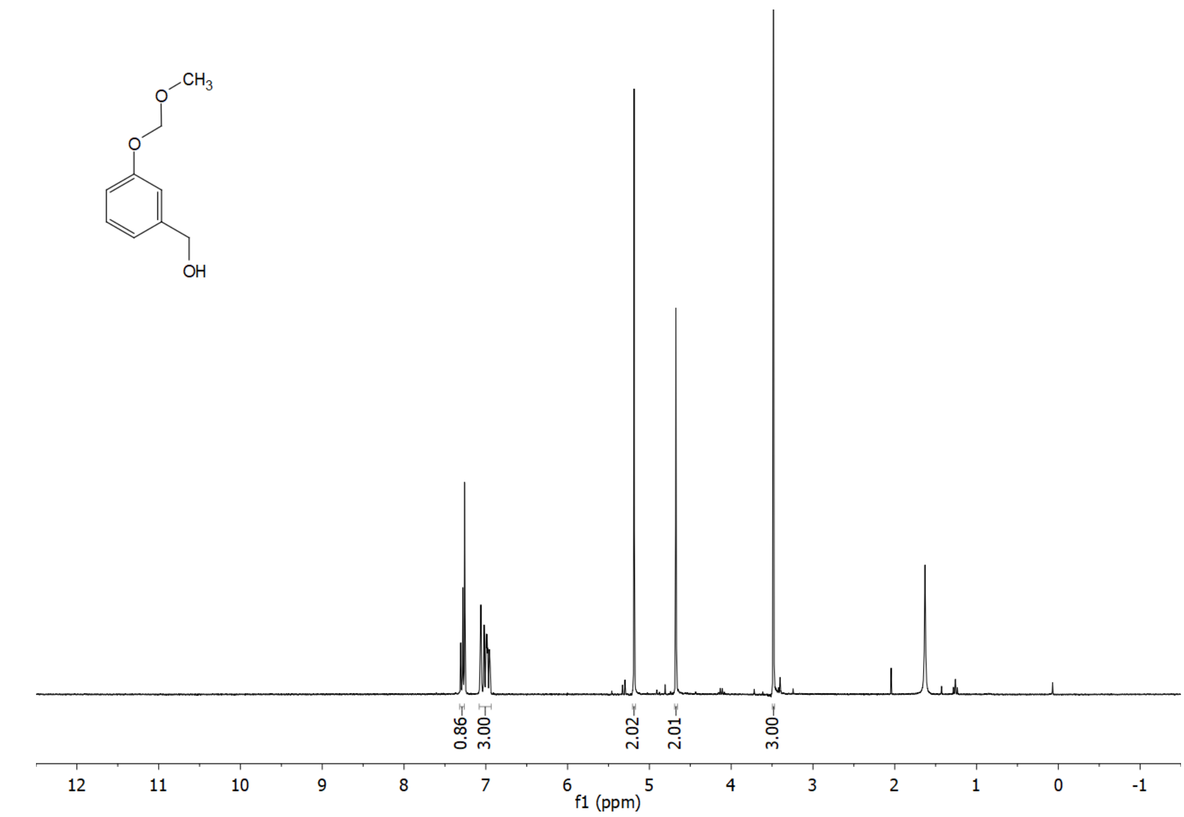

$^{13}$C NMR (APT) of **13** ($CDCl_3$)

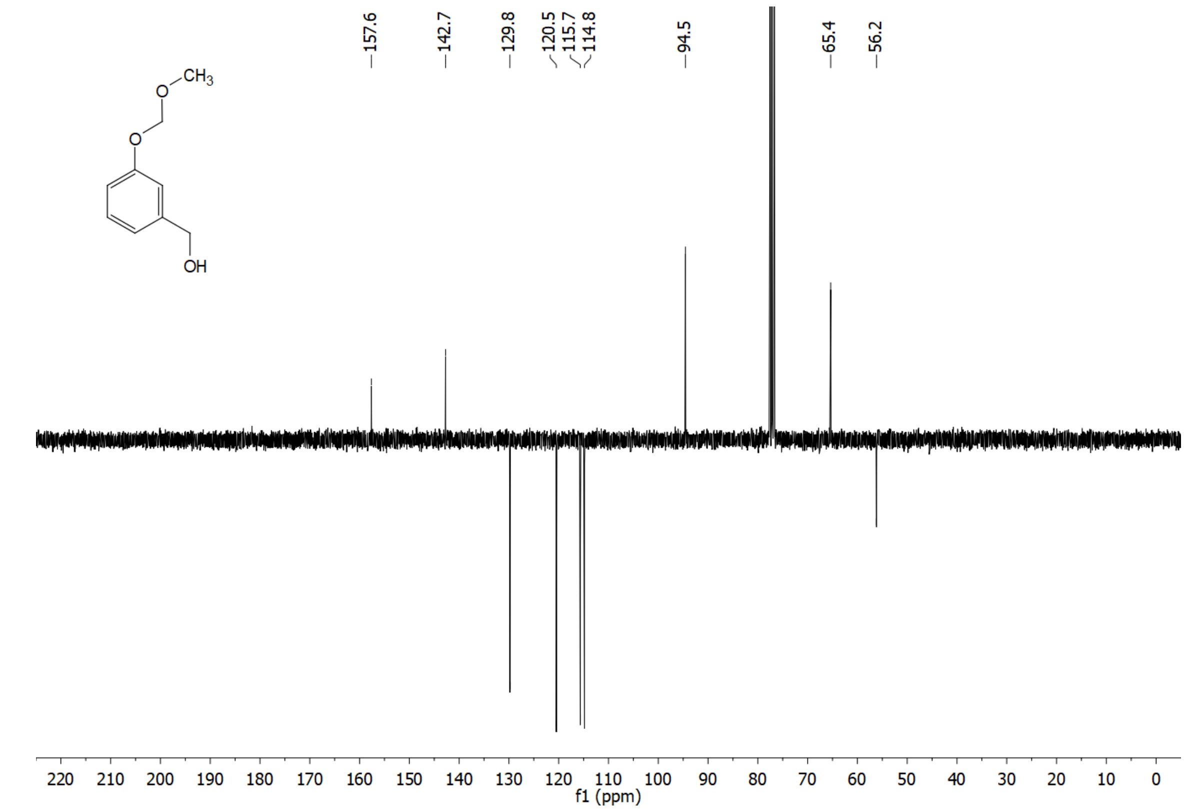

$^{1}H$ NMR of **14** ($CDCl_3$)

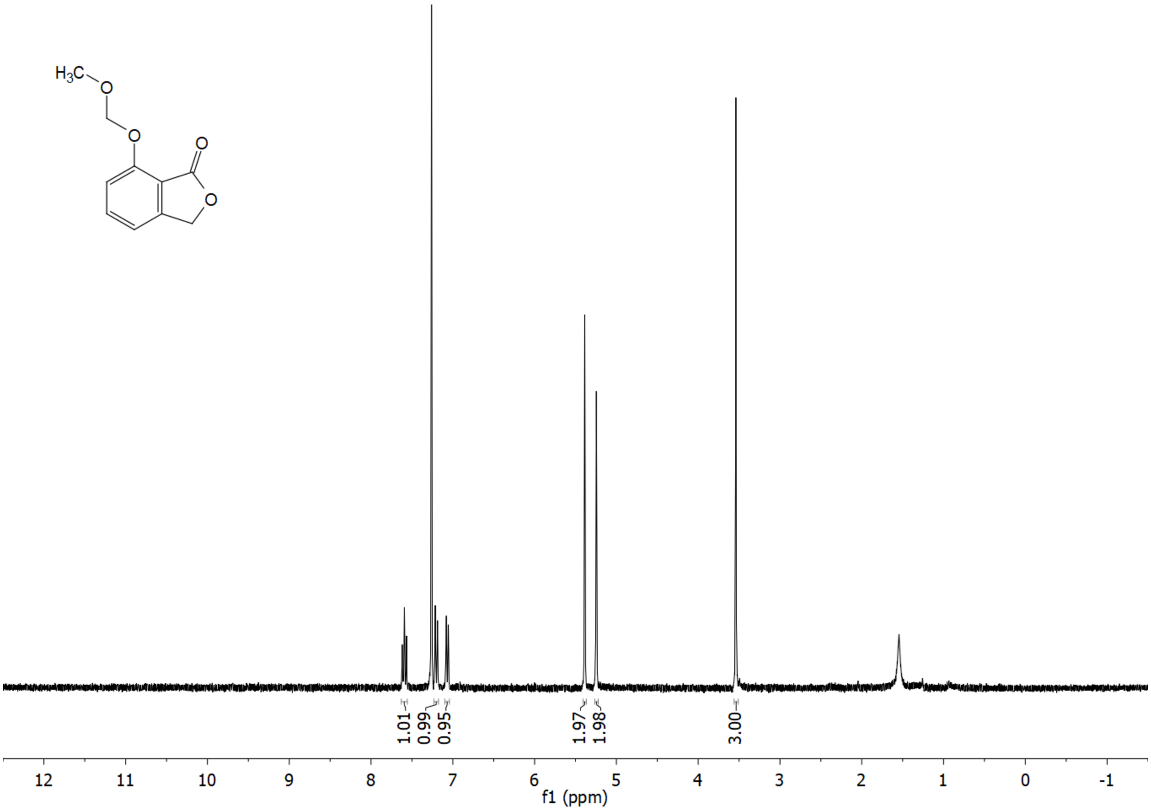

$^{13}C$ NMR (APT) of **14** ($CDCl_3$)

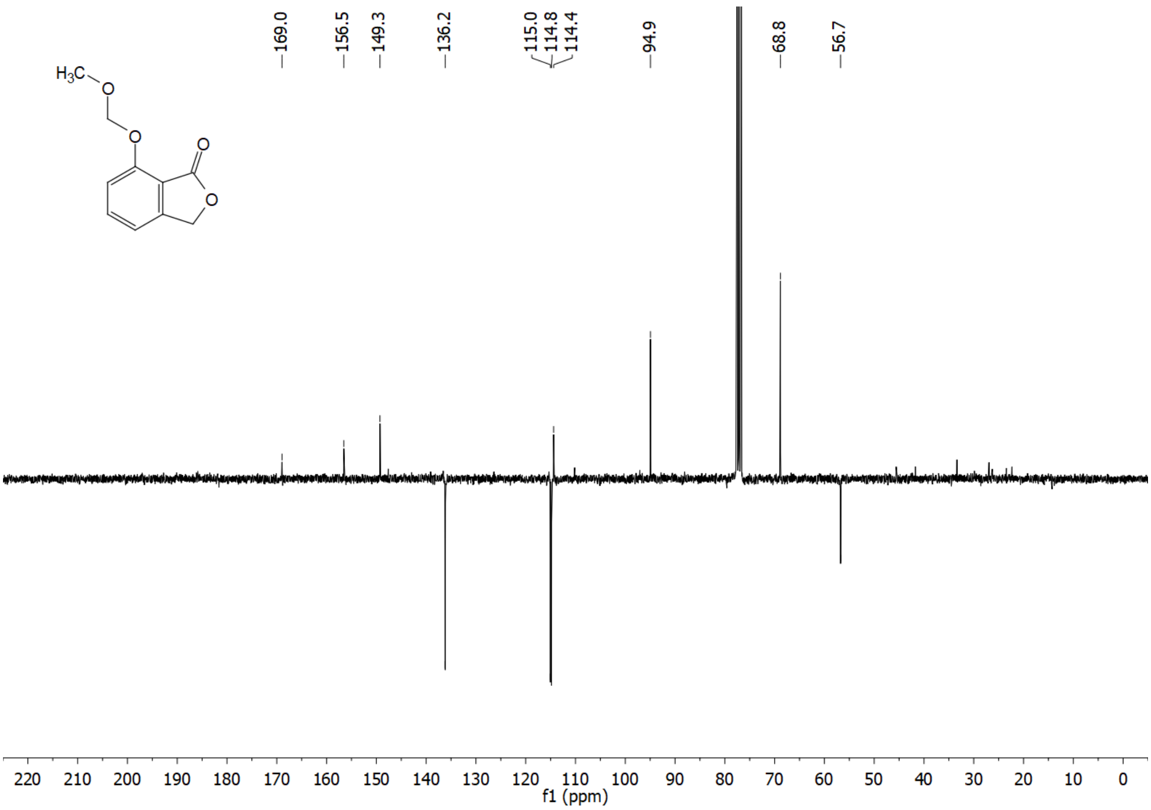

$^{1}H$ NMR of **19b** ($CDCl_3$)

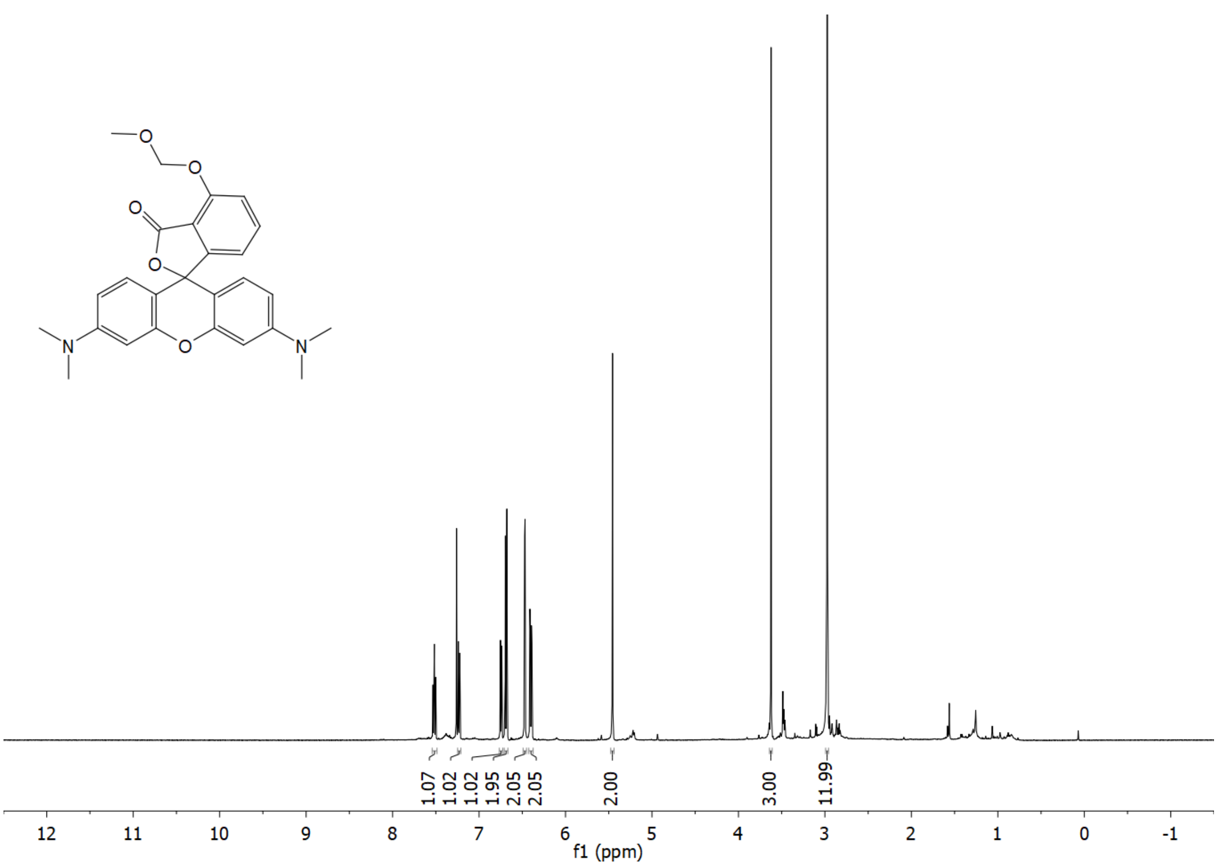

$^{13}C$ NMR of **19b** ($CDCl_3$)

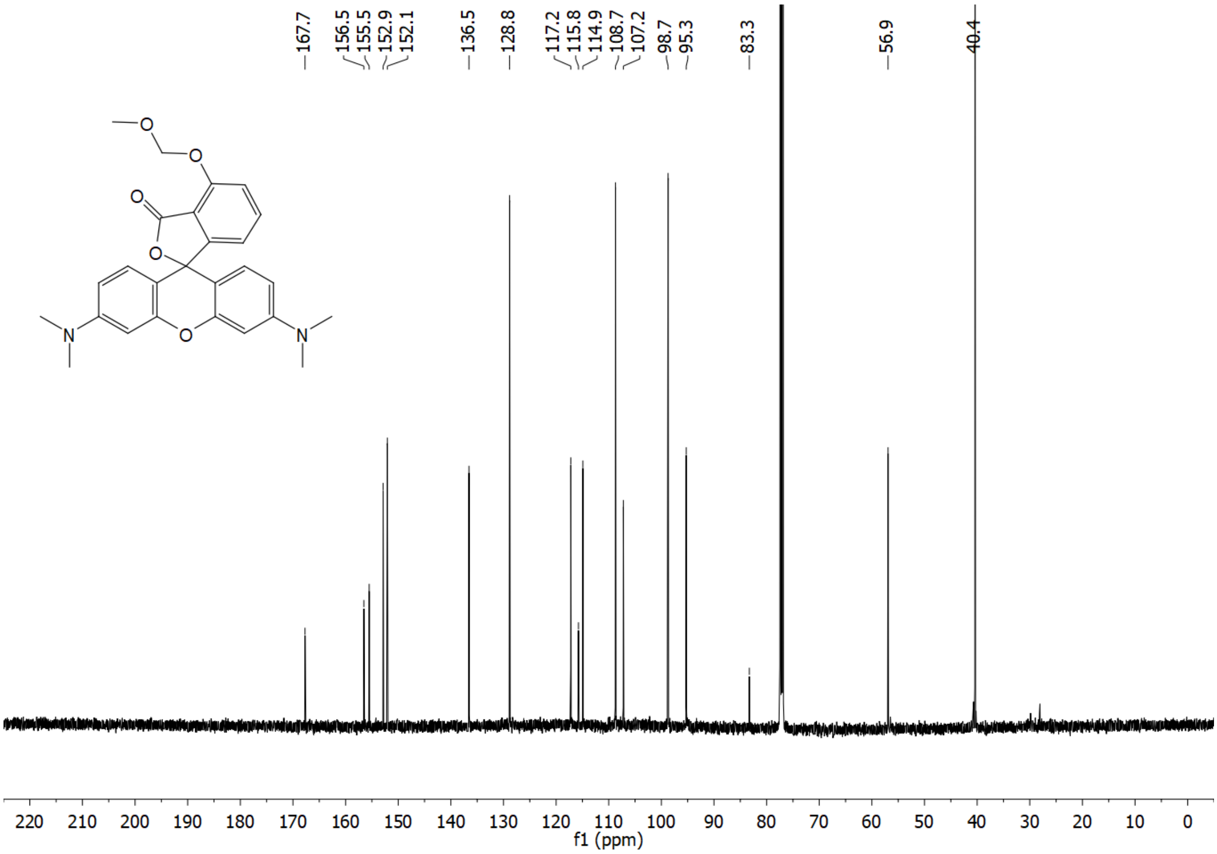

[1]H NMR of **19a** ($CDCl_3$)

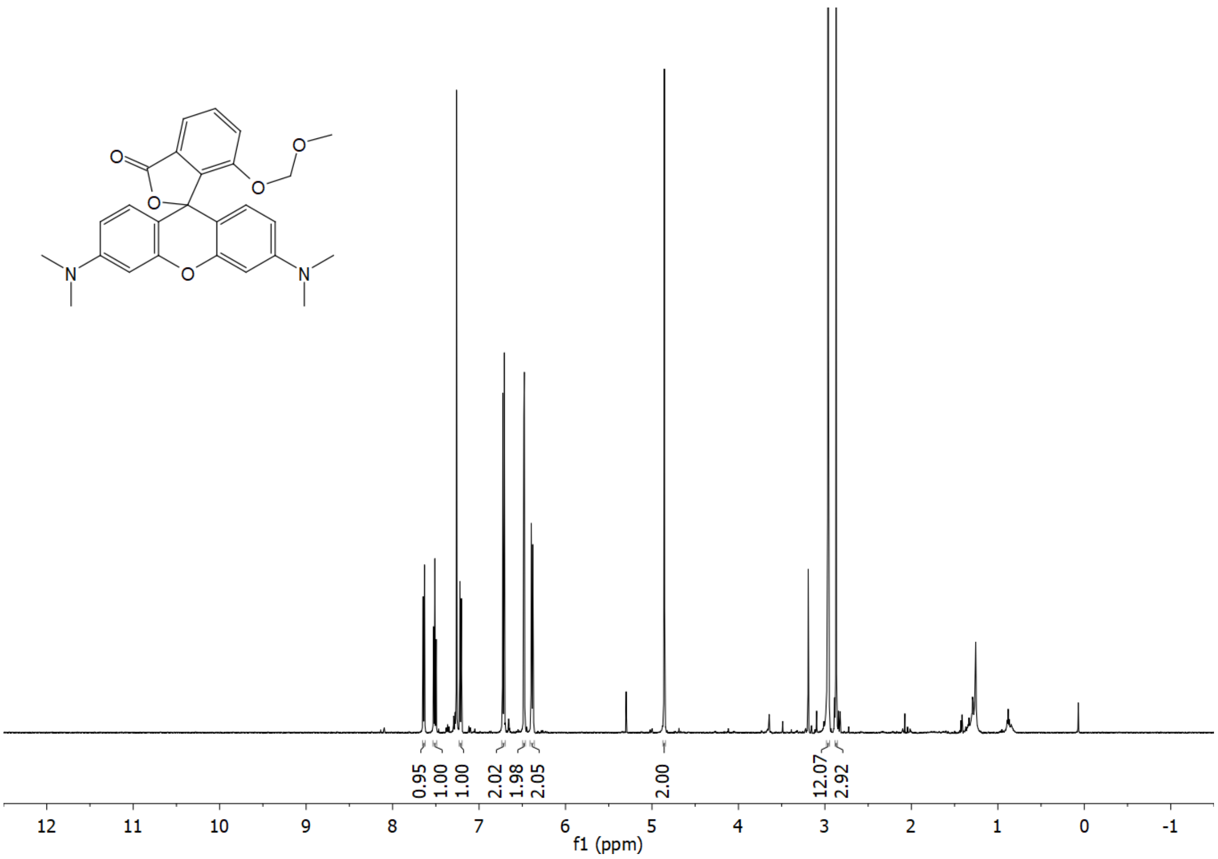

[13]C NMR of **19a** ($CDCl_3$)

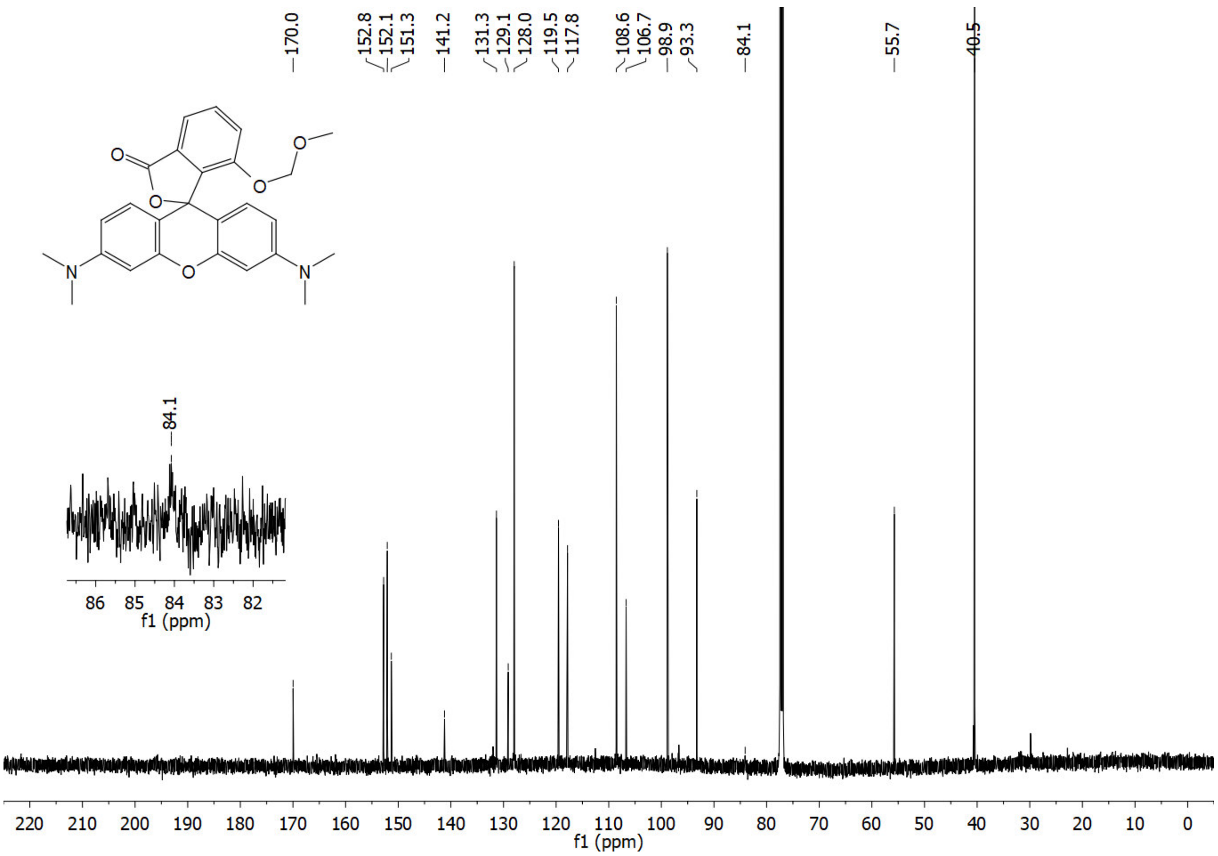

$^{1}H$ NMR of **22** ($CD_3OD$)

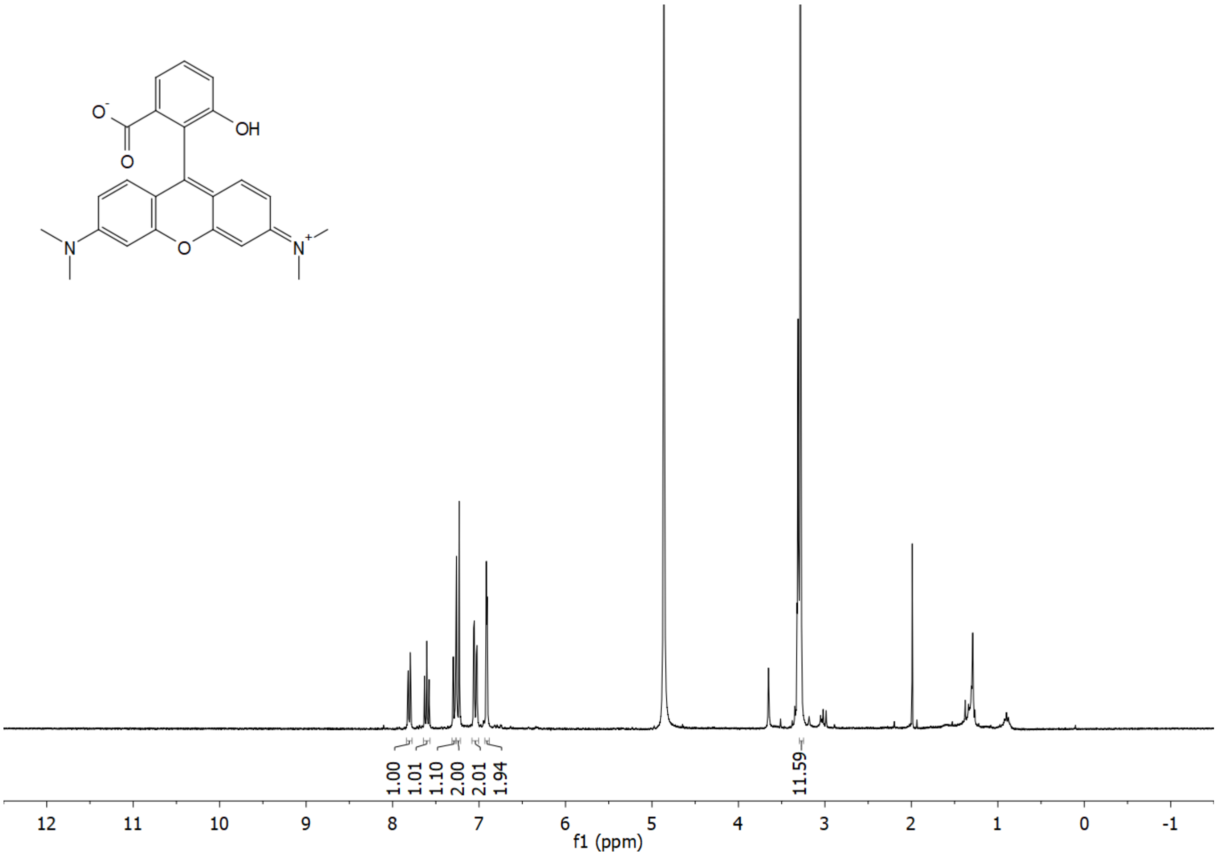

$^{13}C$ NMR of **22** ($CD_3OD$)

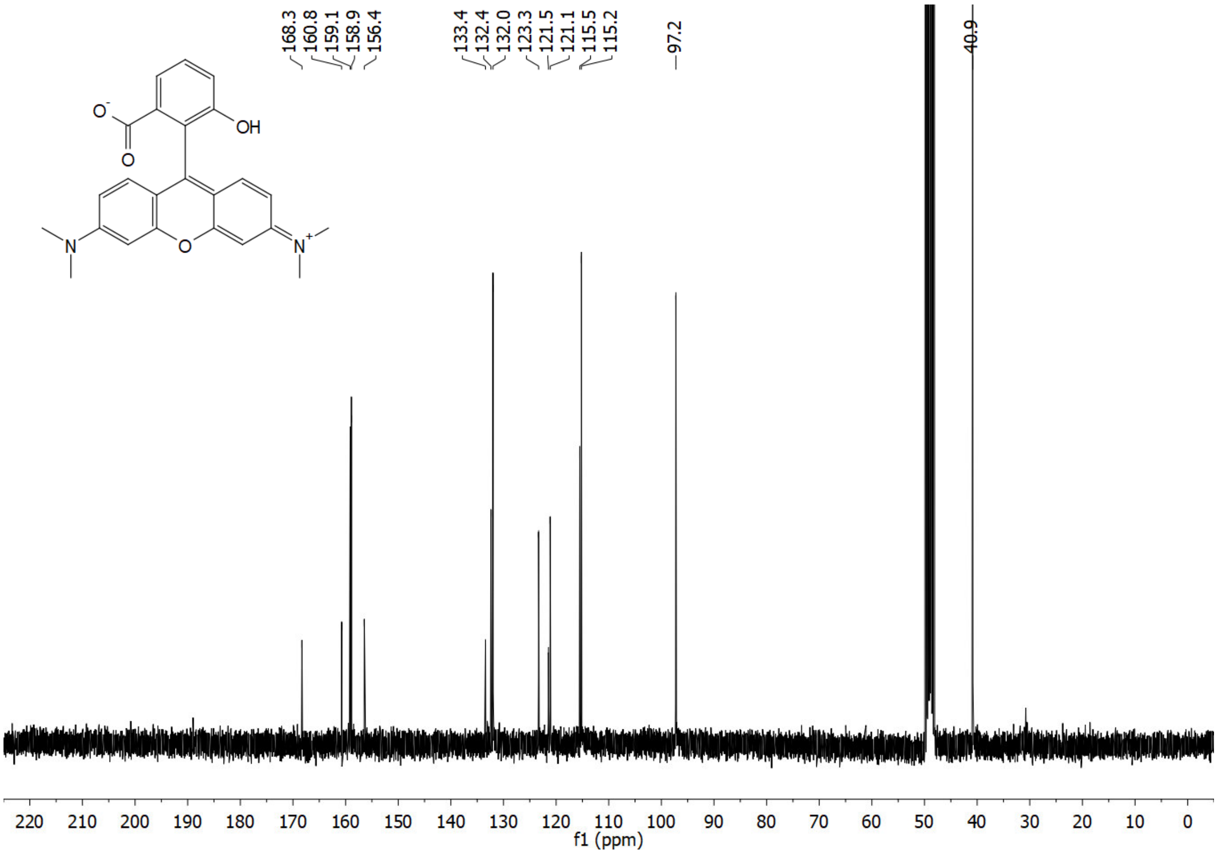

$^{1}H$ NMR of **HD555** ($CD_3OD$)

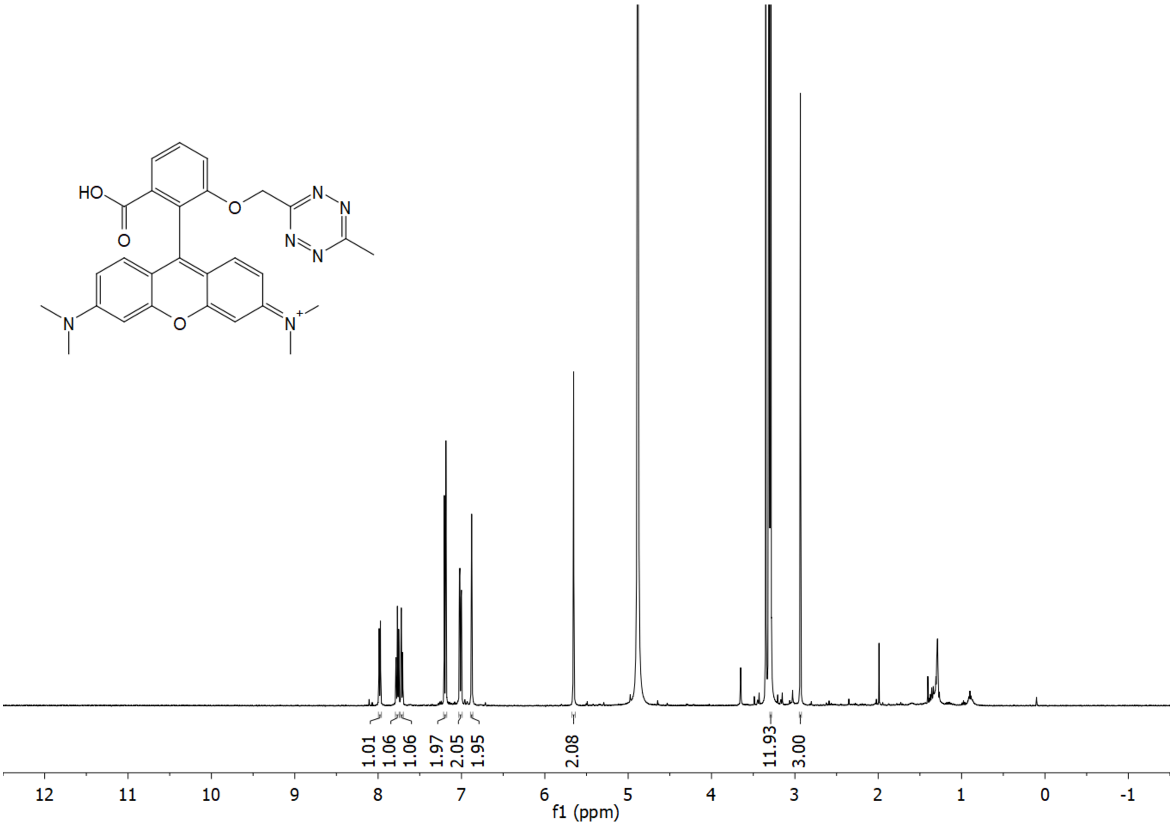

$^{13}C$ NMR of **HD555** ($CD_3OD$)

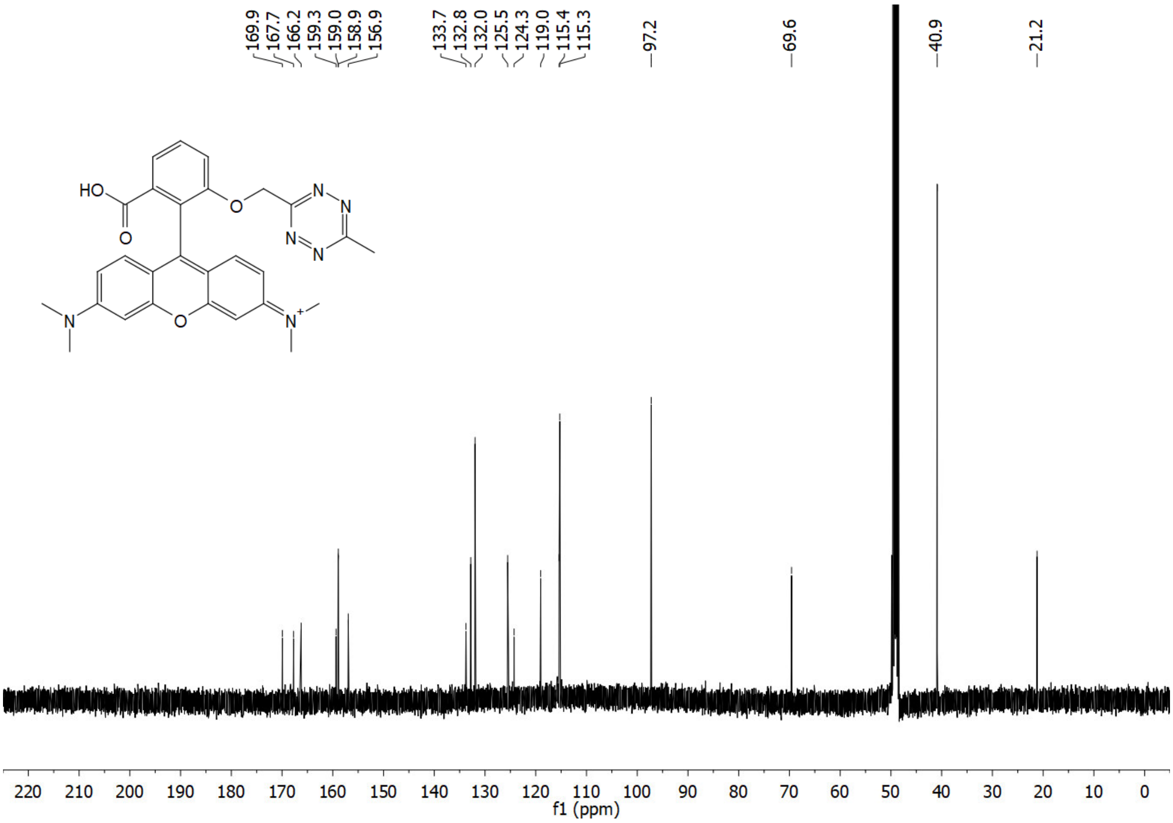

$^{1}$H NMR of **20b** ($CD_3CN$)

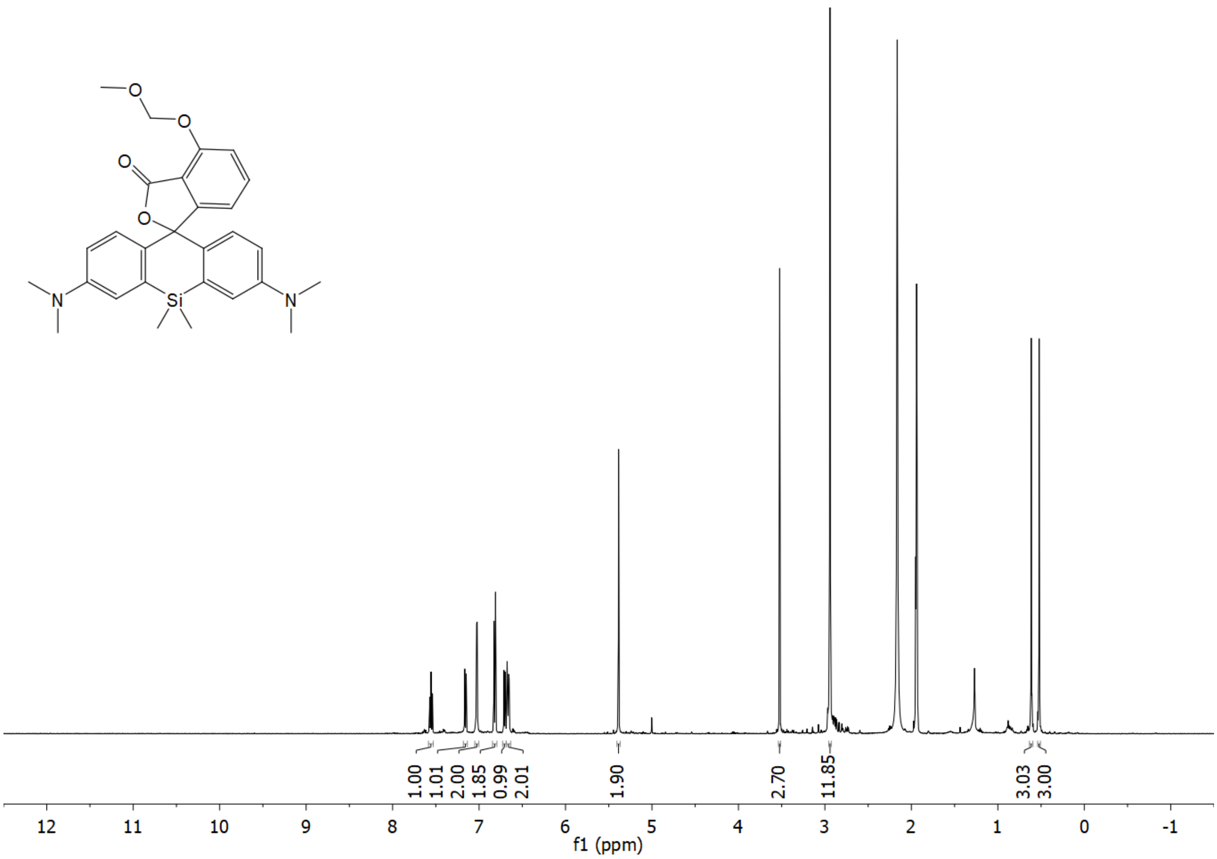

$^{13}$C NMR (APT) of **20b** ($CD_3CN$)

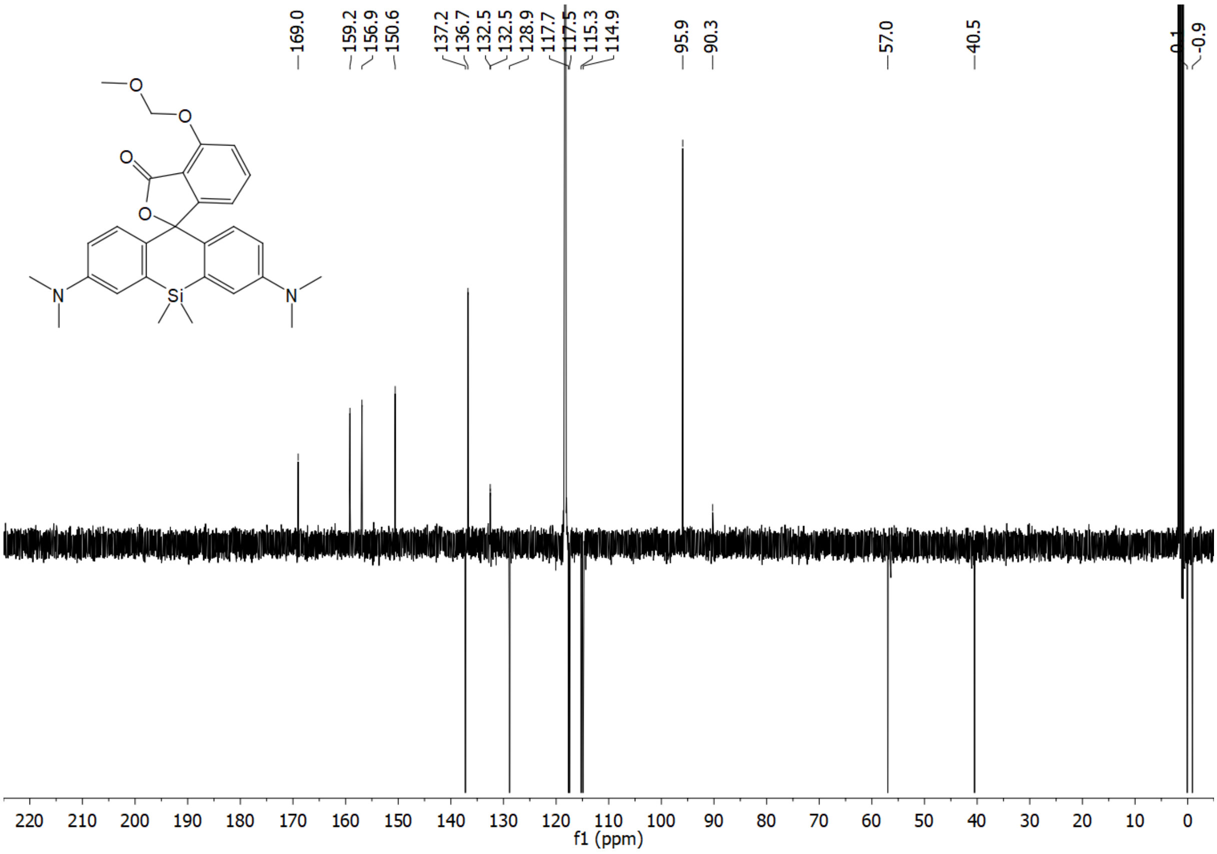

$^1$H NMR of **20a** ($CDCl_3$)

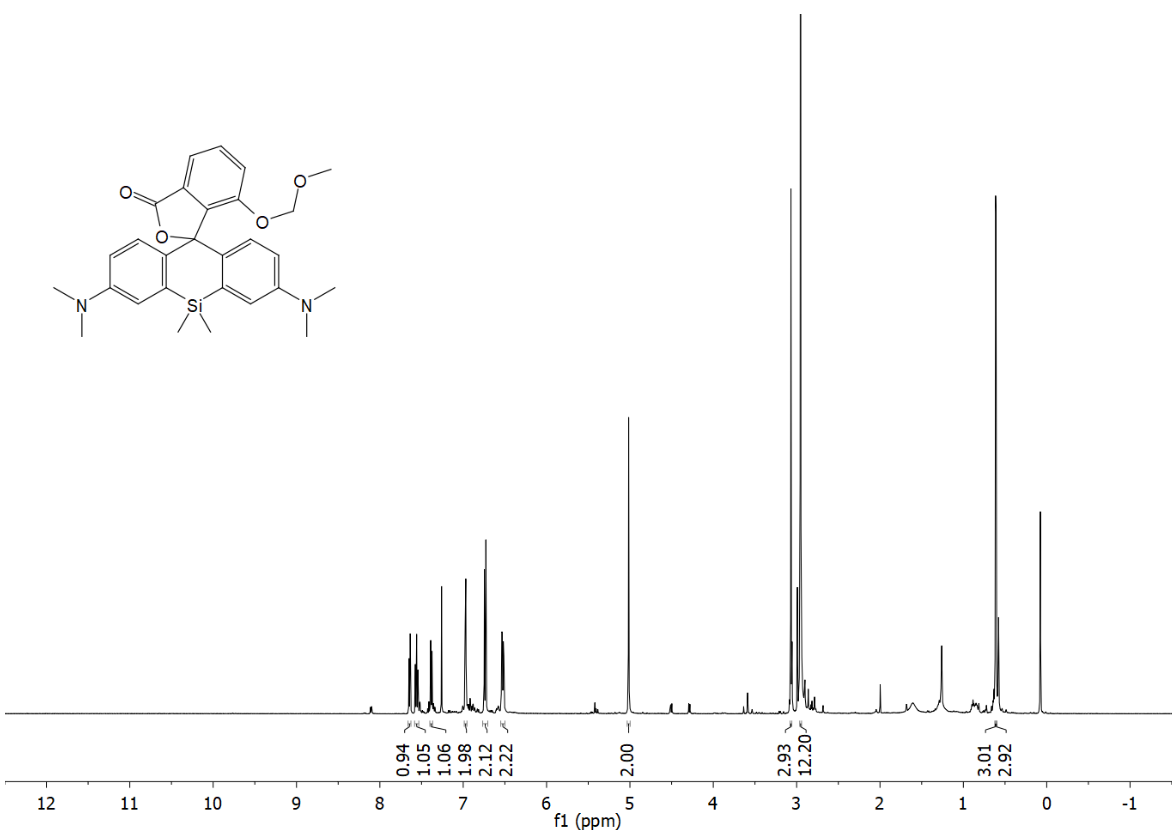

$^{13}$C NMR (APT) of **20a** ($CDCl_3$)

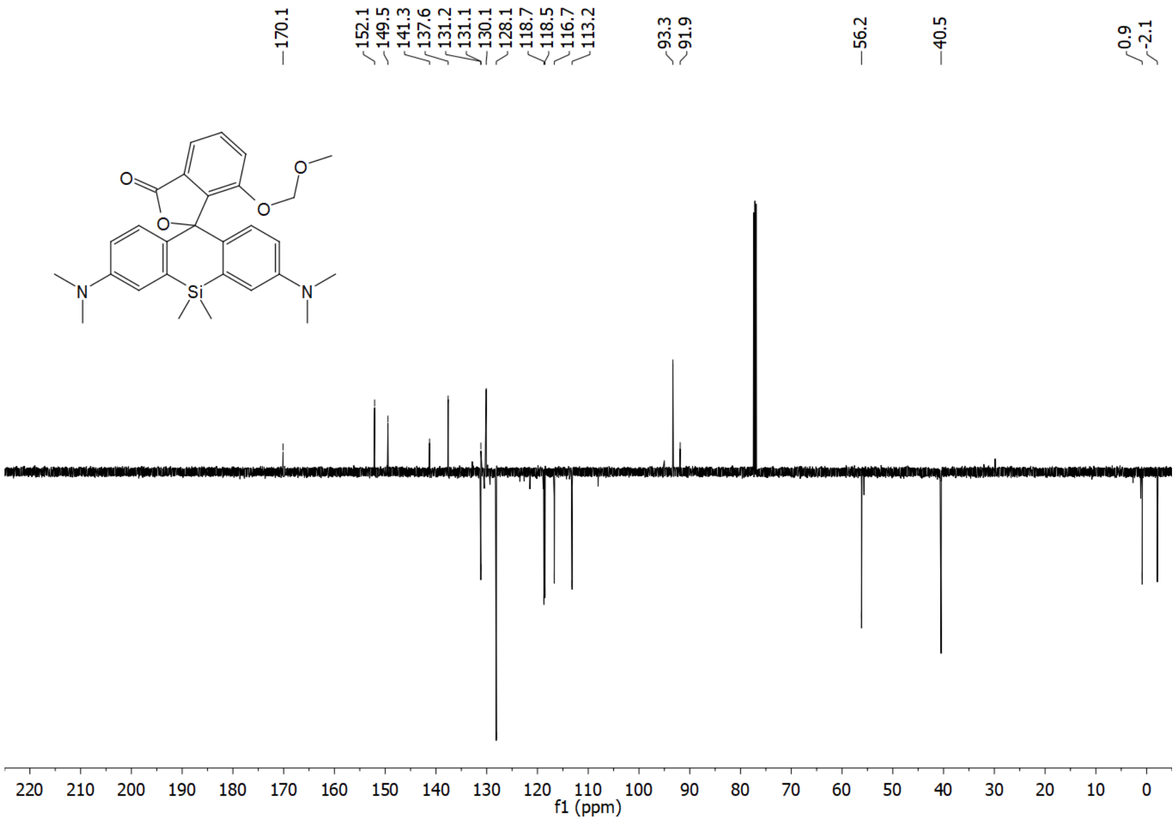

$^{1}H$ NMR of **23** ($CDCl_3$)

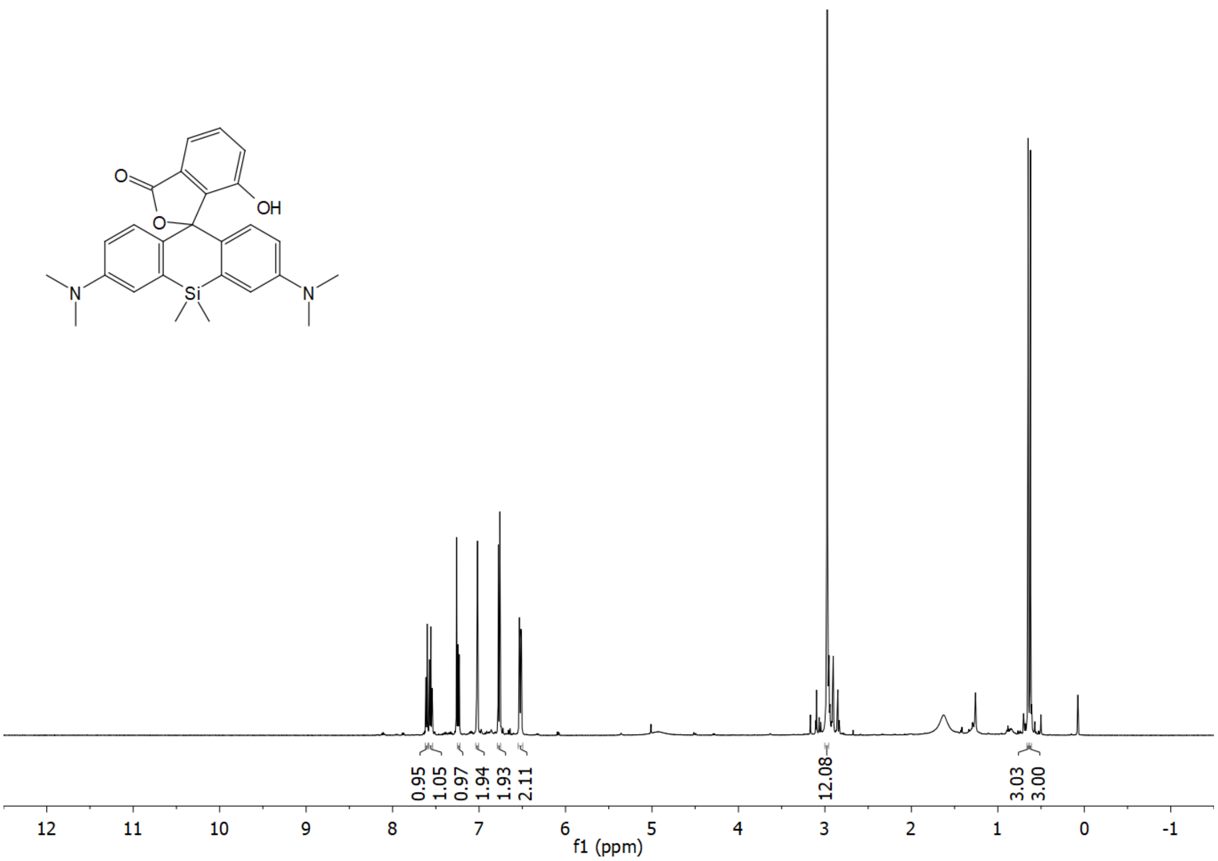

$^{13}C$ NMR (APT) of **23** ($CDCl_3$)

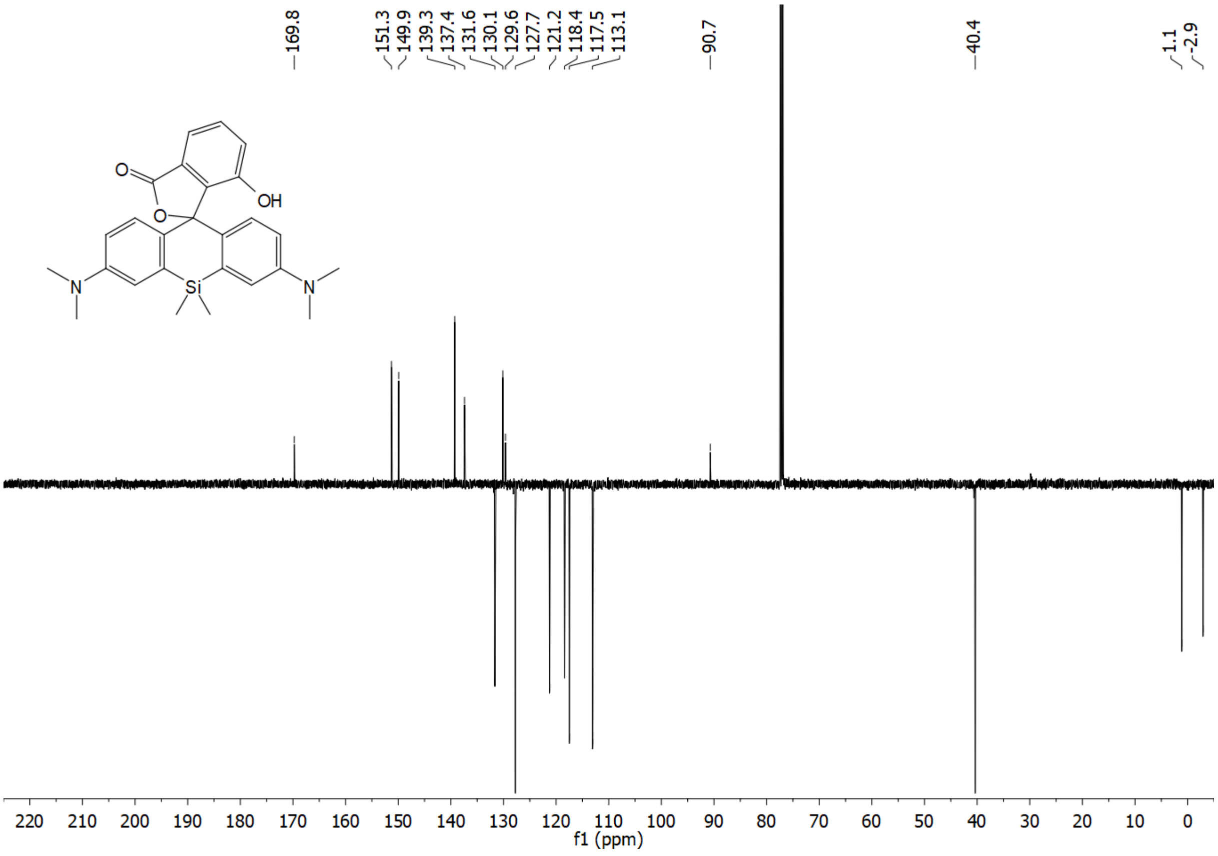

$^{1}H$ NMR of **HD653** ($C_6D_6$)

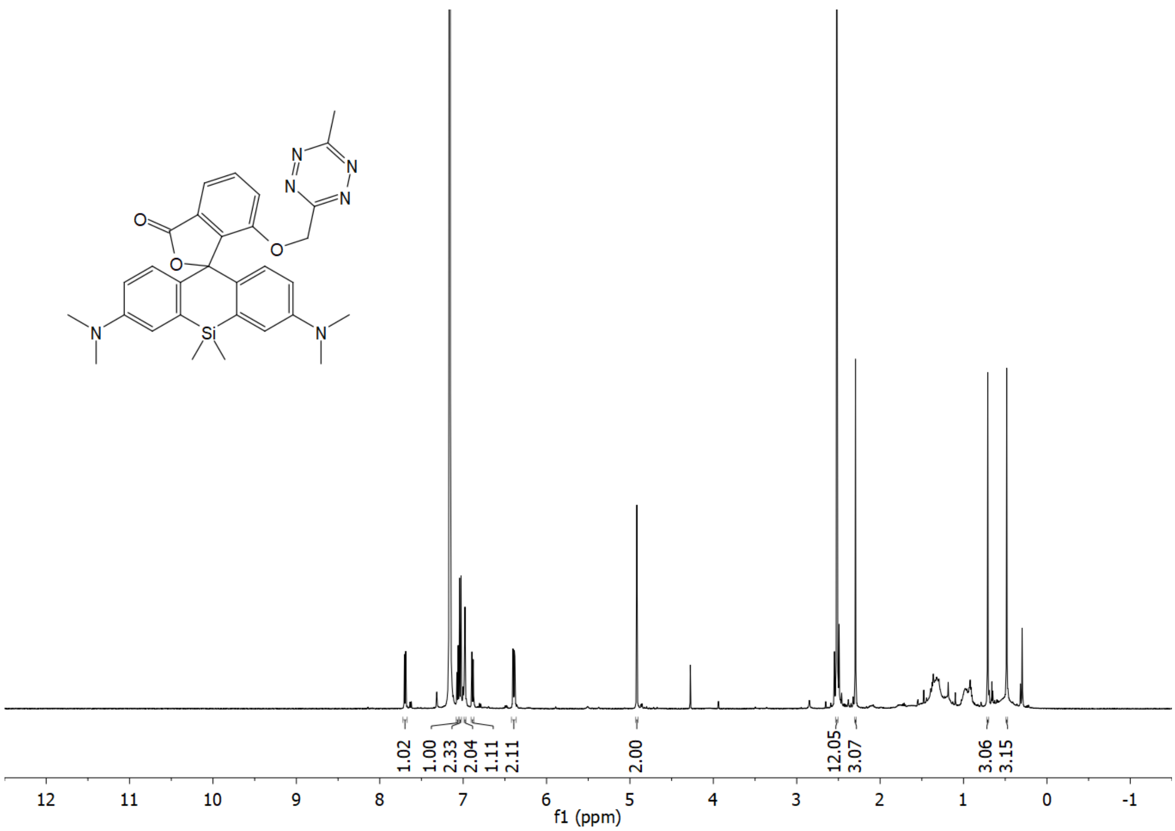

$^{13}C$ NMR of **HD653** ($C_6D_6$)

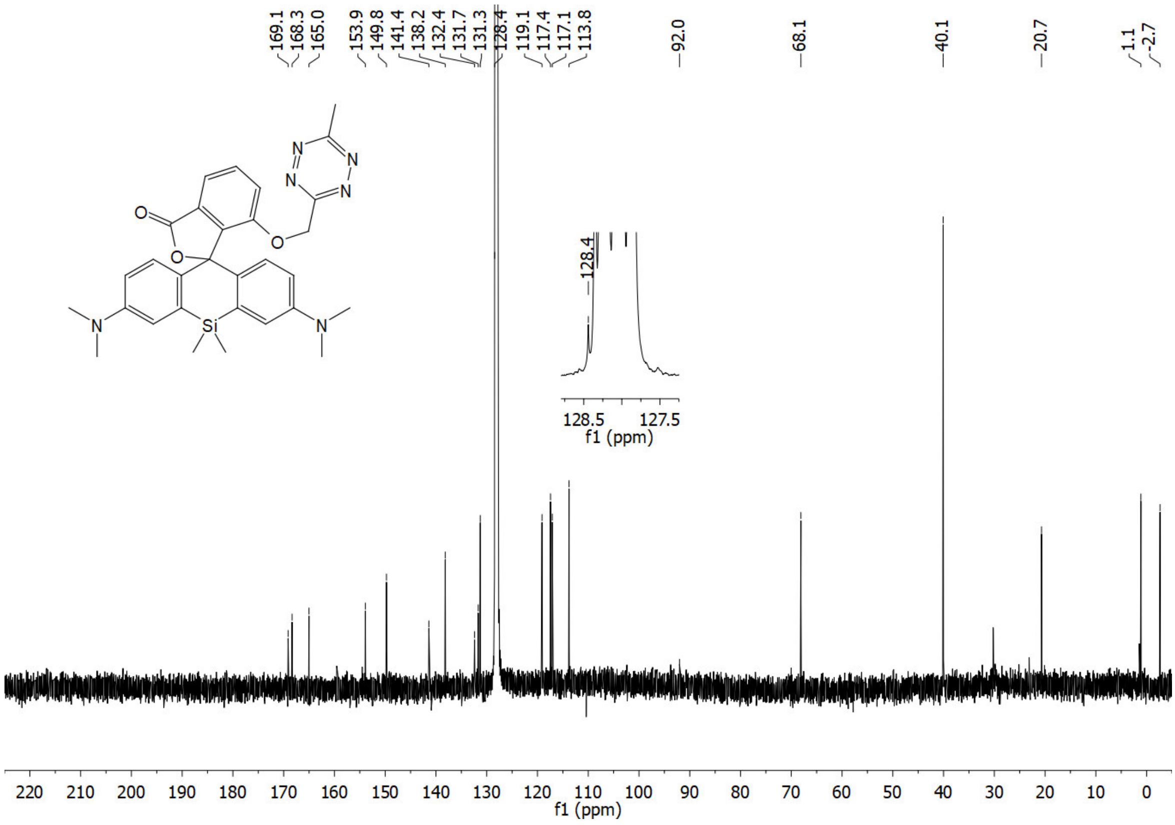

$^{1}$H NMR of **21** ($CDCl_3$)

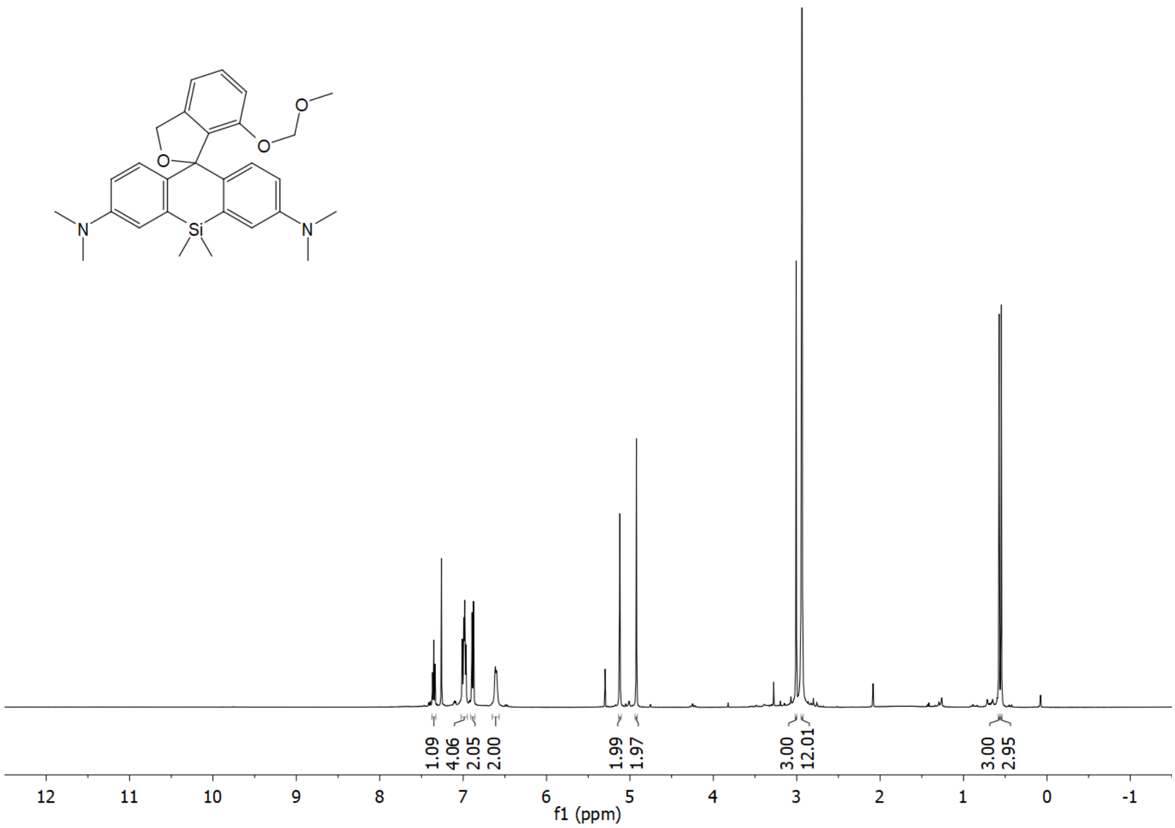

$^{13}$C NMR (APT) of **21** ($CDCl_3$)

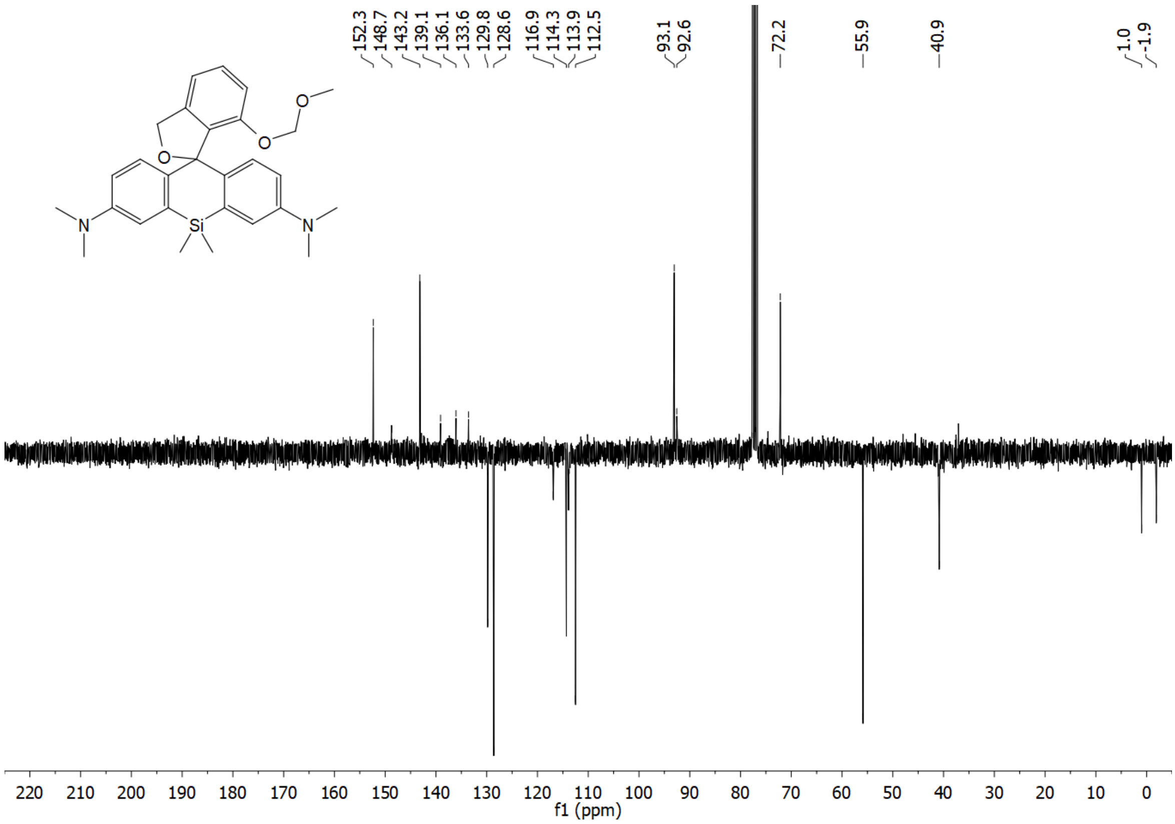

$^{1}H$ NMR of **24** ($CD_3OD$)

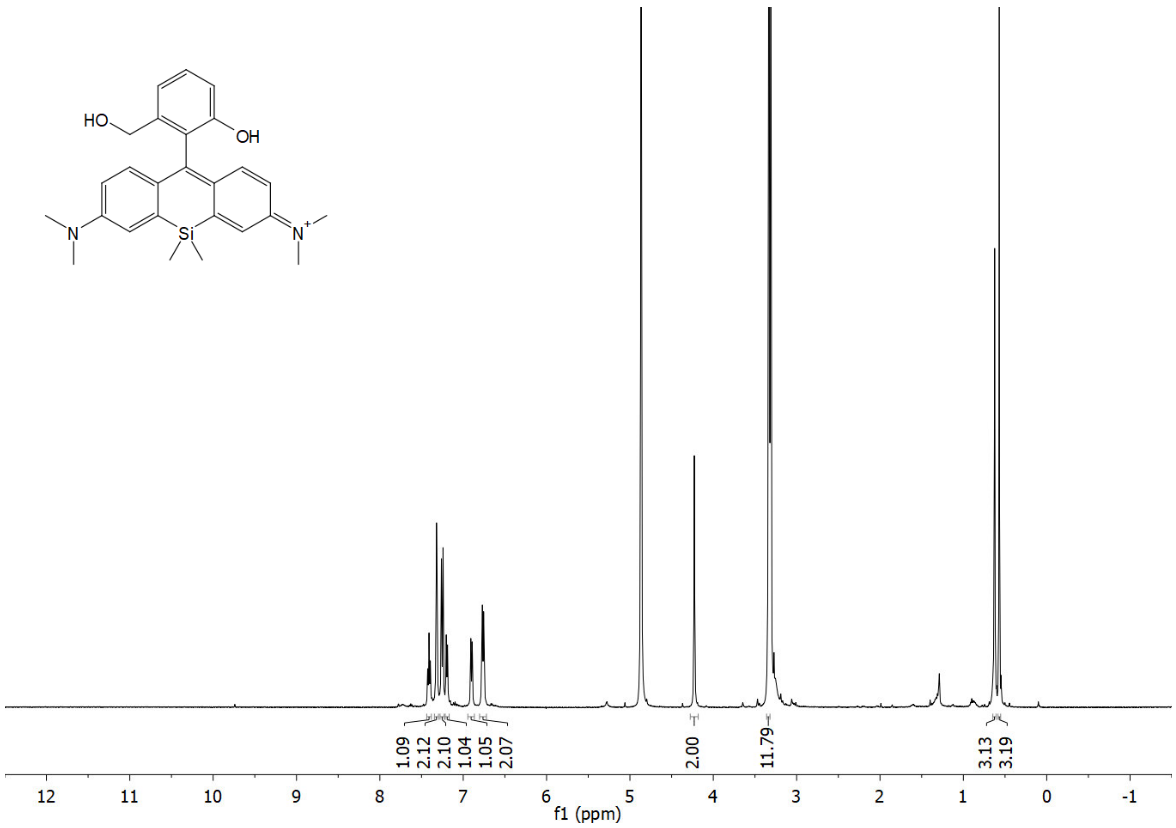

$^{13}C$ NMR (APT) of **24** ($CD_3OD$)

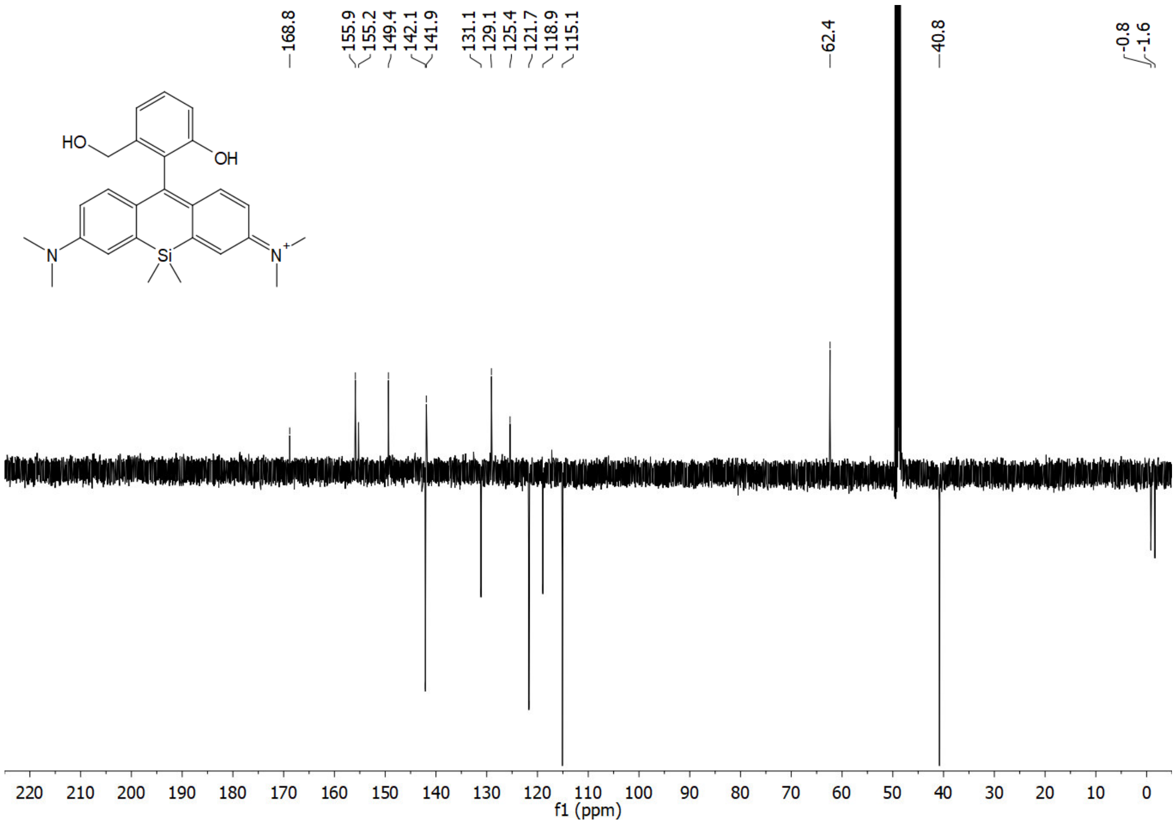

$^{1}H$ NMR of **sb-HD656** ($CD_3OD$)

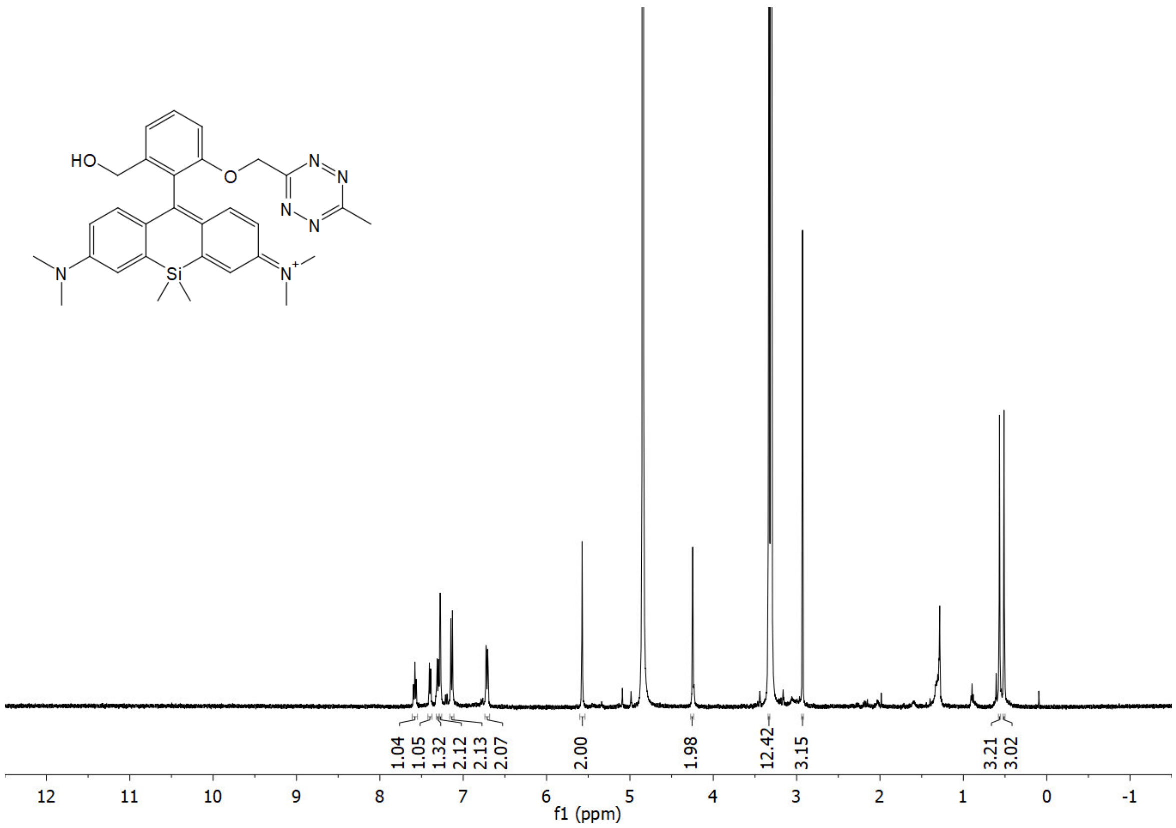

$^{13}C$ NMR (APT) of **sb-HD656** ($CD_3OD$)

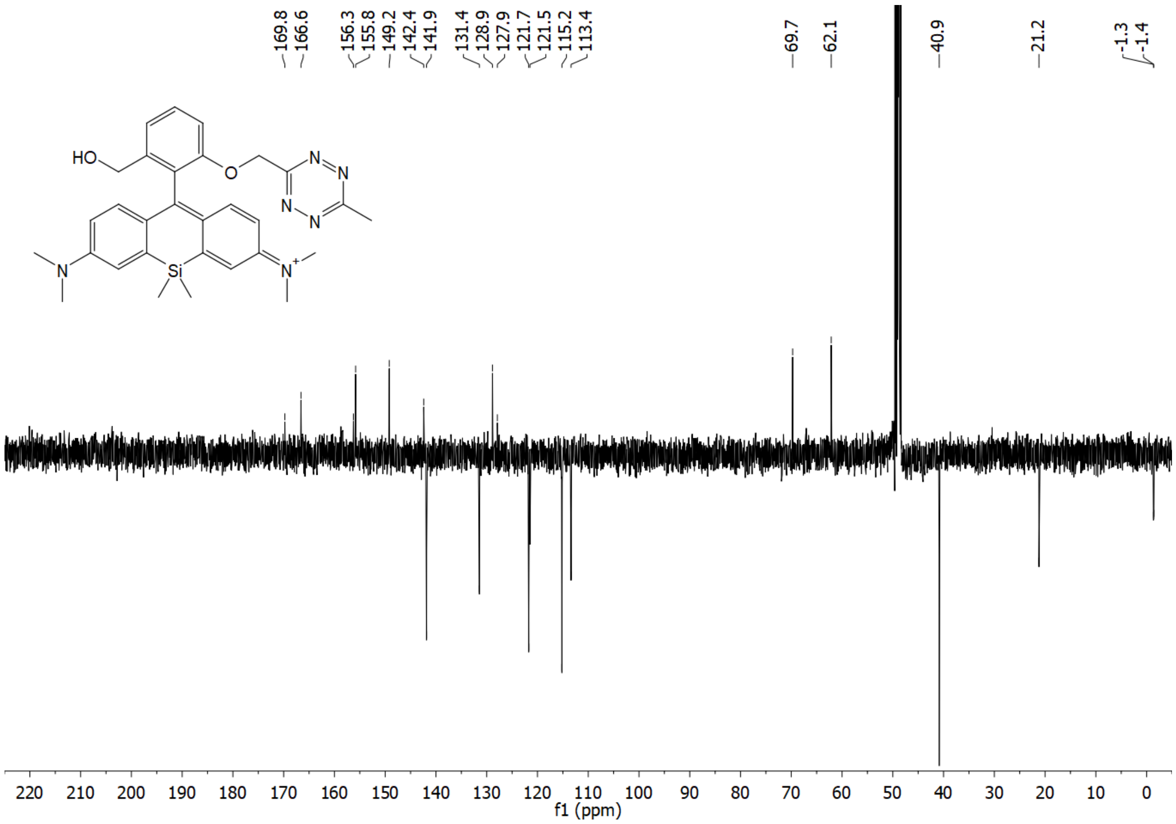

$^{1}H$ NMR of **HD561x** ($CD_3OD$)

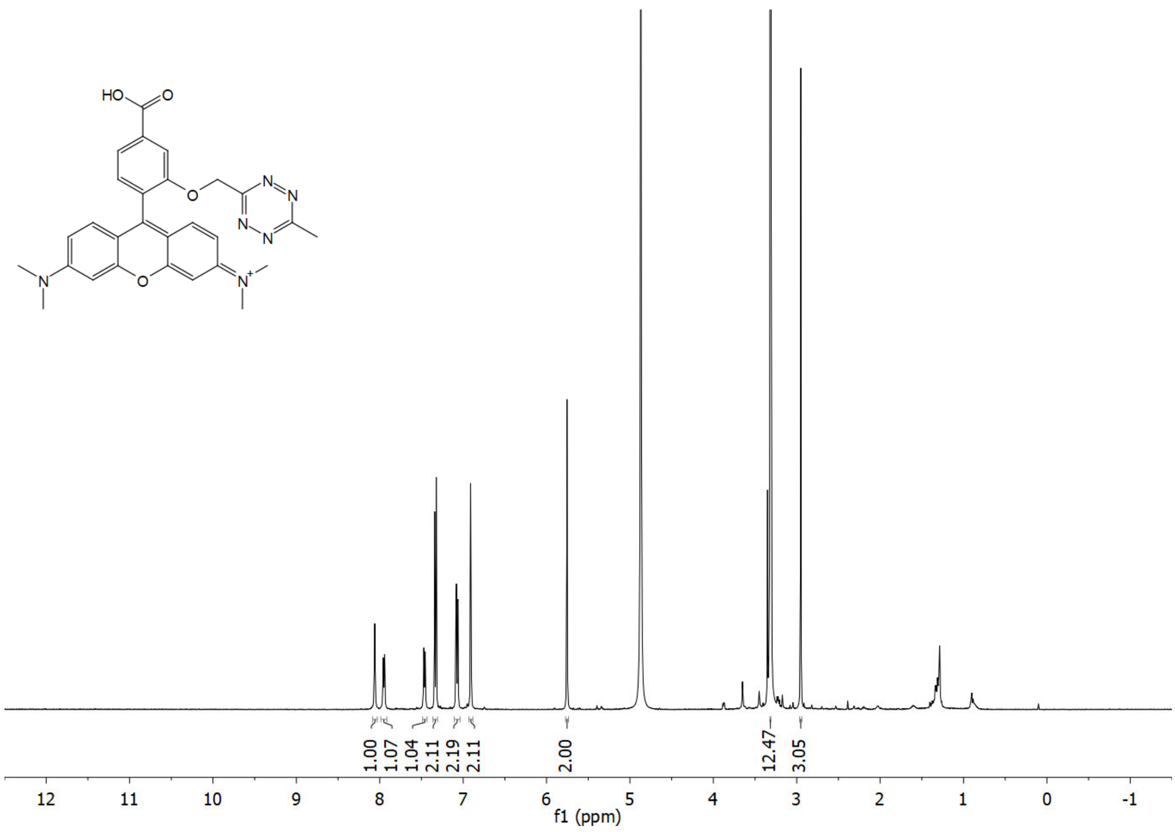

$^{13}C$ NMR (APT) of **HD561x** ($CD_3OD$)

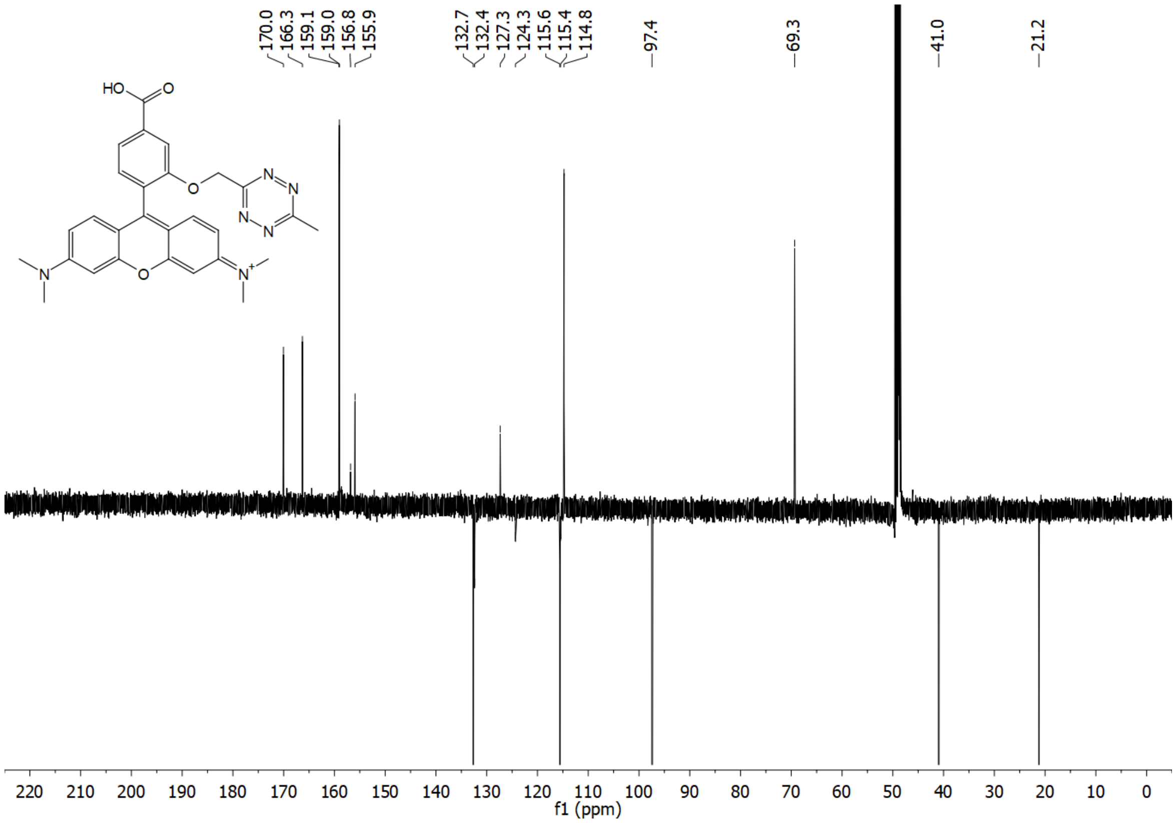

$^{1}H$ NMR of **HD654x** ($CD_3OD$)

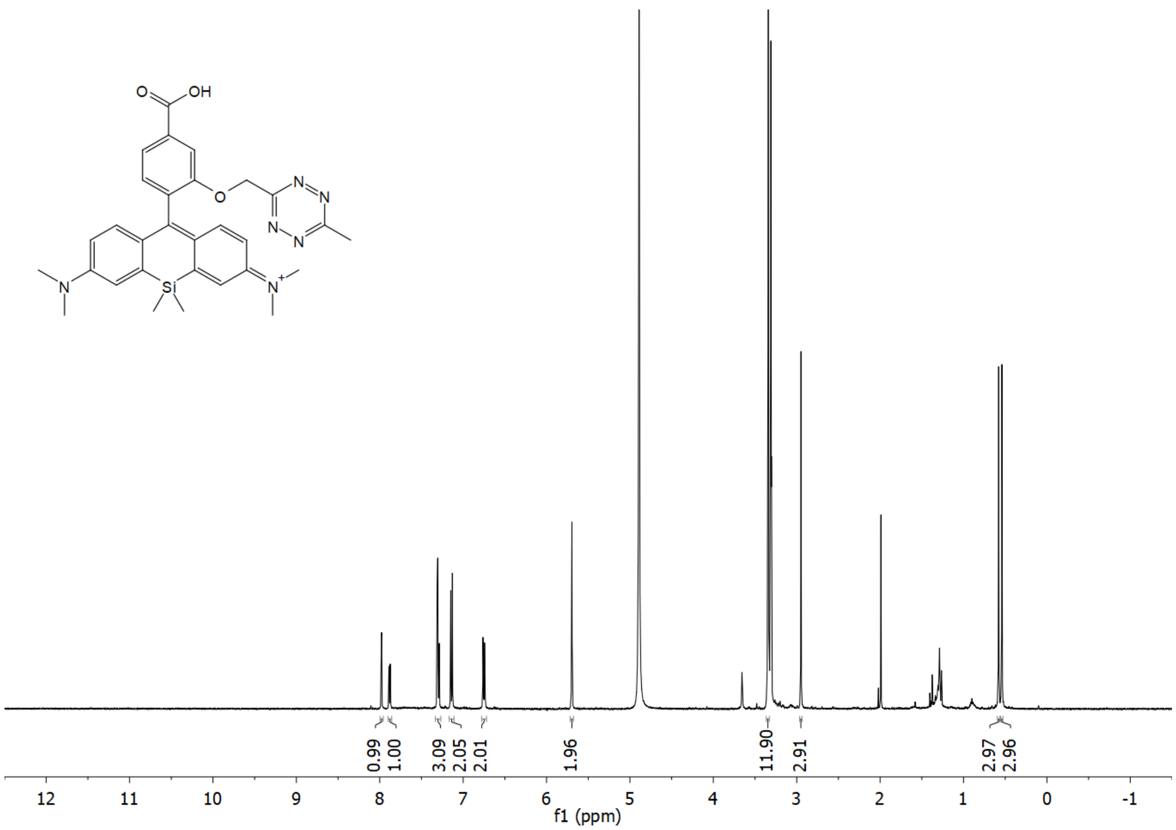

$^{13}C$ NMR (APT) of **HD654x** ($CD_3OD$)

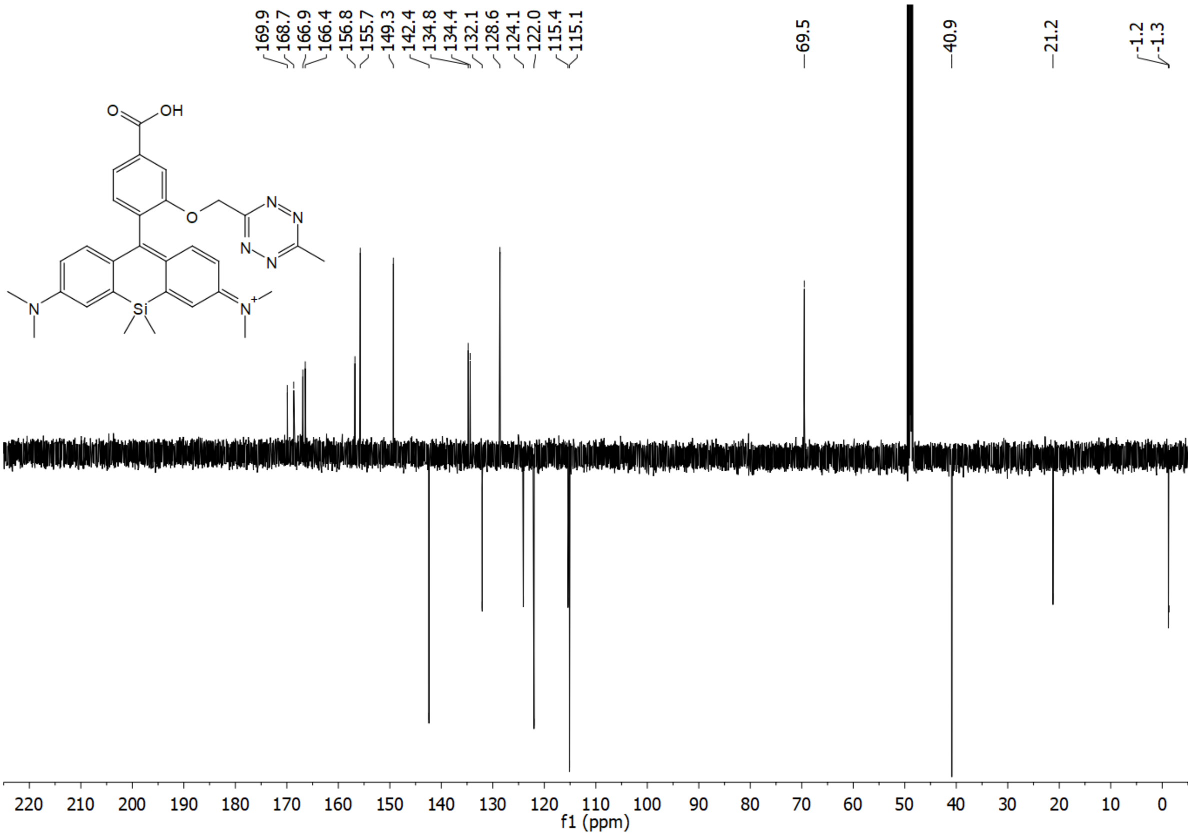

[1]H NMR of ***o*-TzR** ($CD_3CN$)

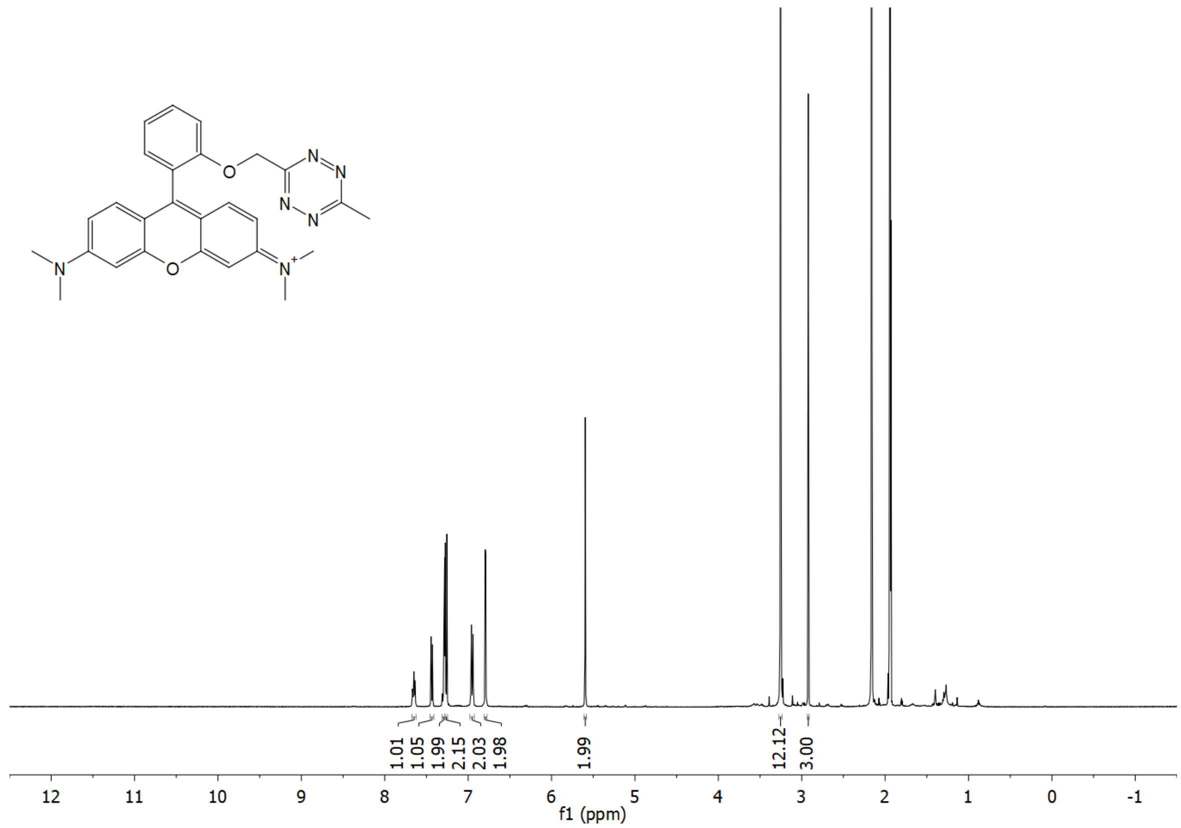

[13]C NMR (APT) of ***o*-TzR** ($CD_3CN$)

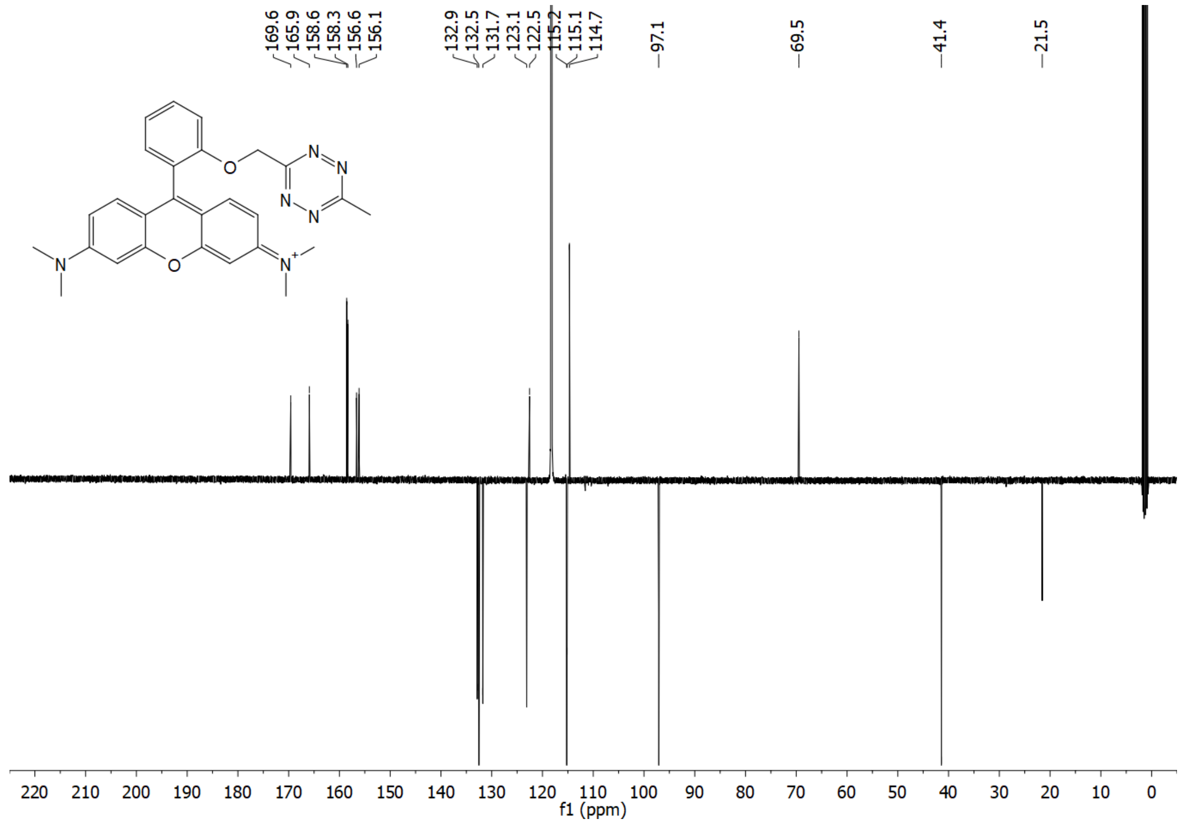

$^{1}H$ NMR of ***m*-TzR** ($CD_3CN$)

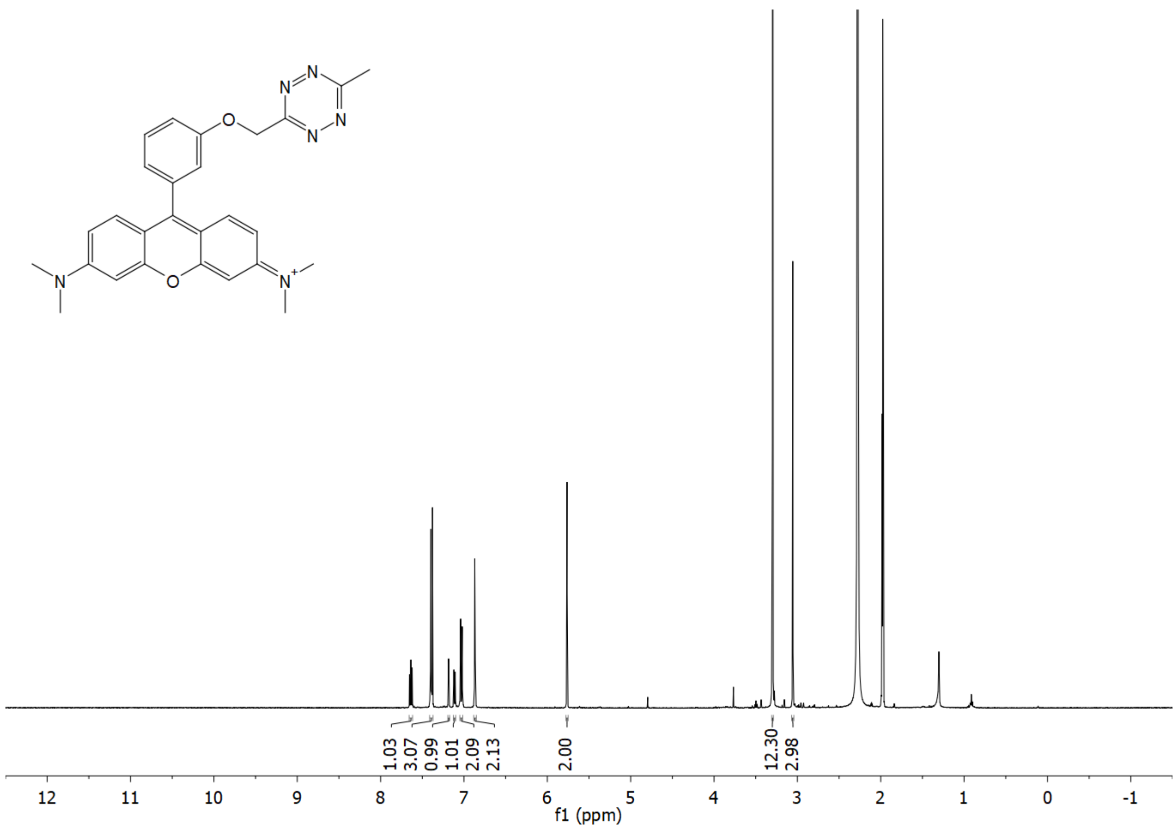

$^{13}C$ NMR (APT) of ***m*-TzR** ($CDCl_3$)

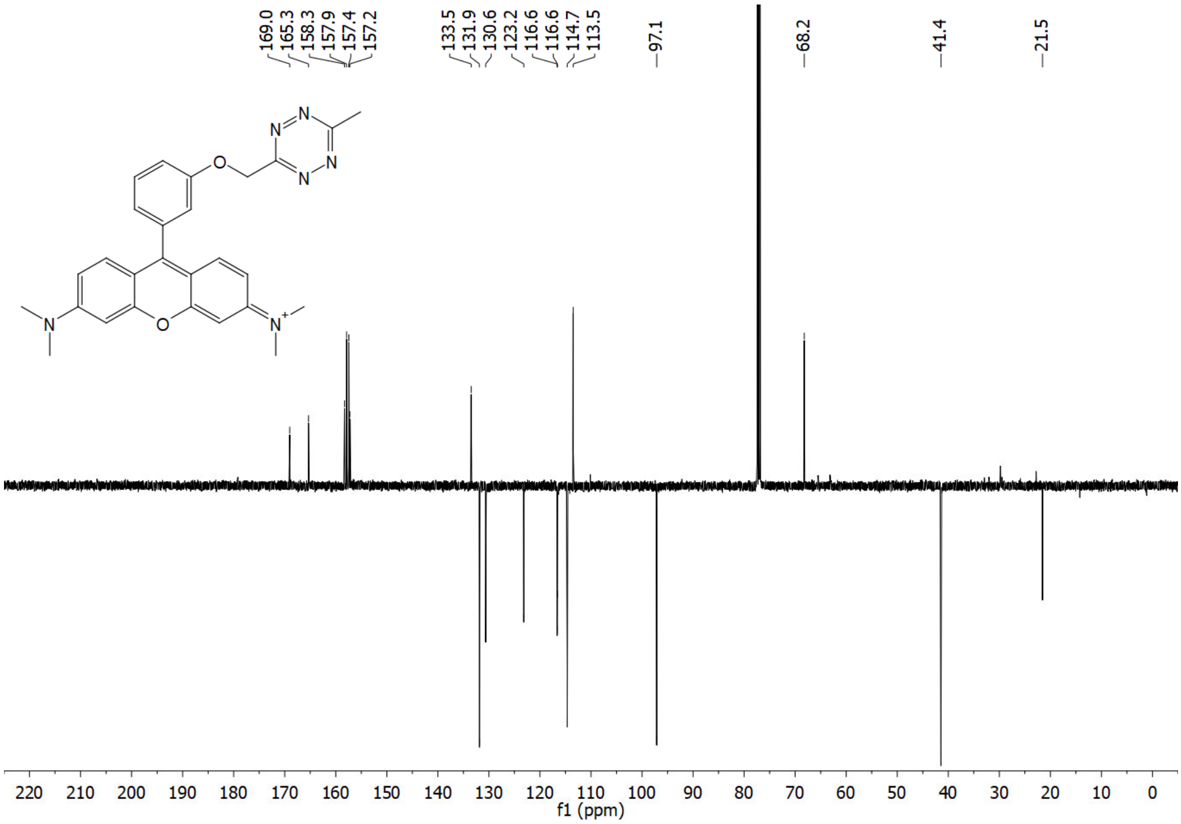

$^{1}H$ NMR of ***p*-TzR** ($CD_3CN$)

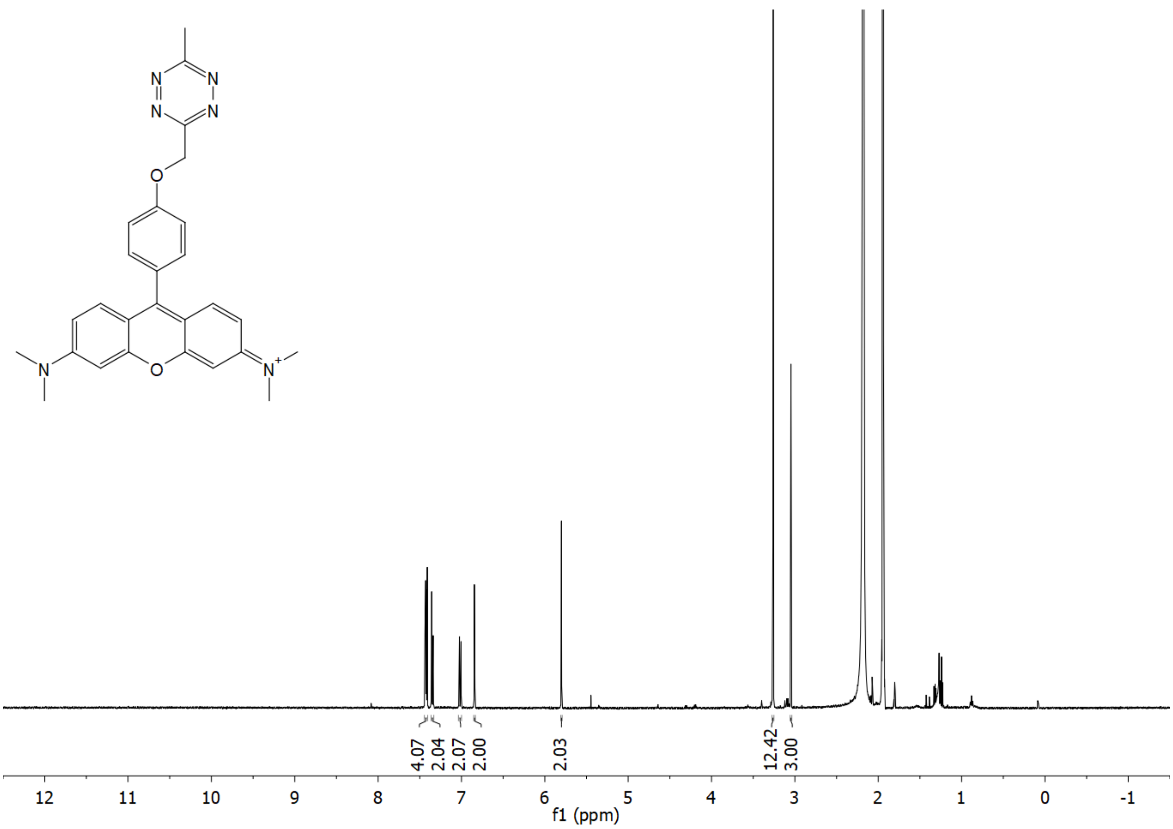

$^{13}C$ NMR (APT) of ***p*-TzR** ($CD_3CN$)

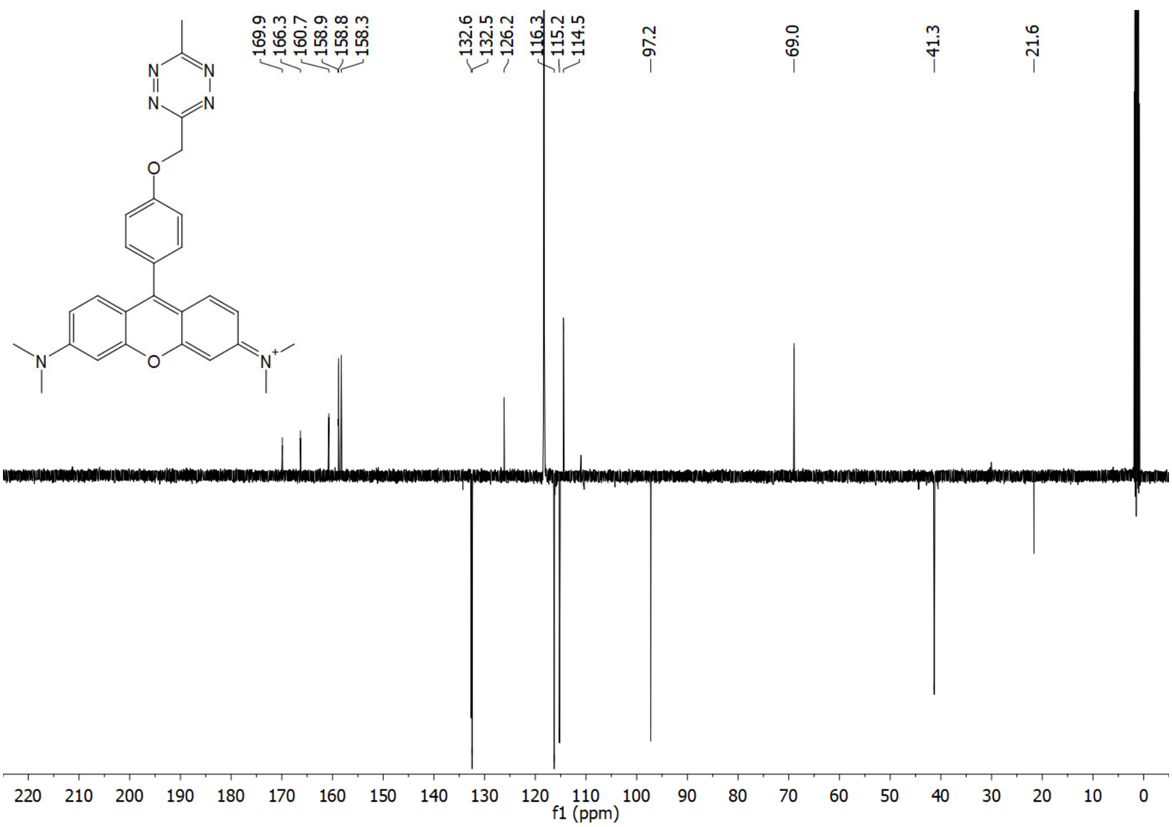

$^{1}H$ NMR of ***o*-TzSiR** ($CD_3CN$)

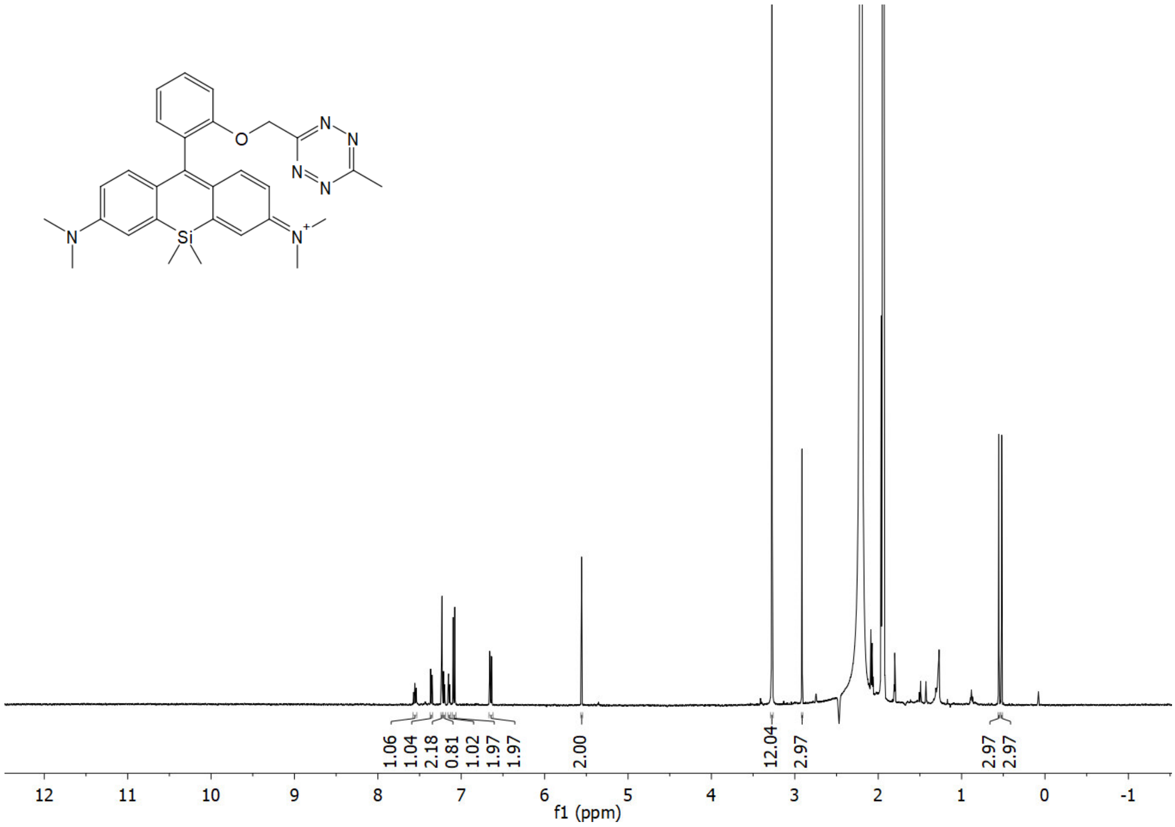

$^{13}C$ NMR (APT) of ***o*-TzSiR** ($CD_3OD$)

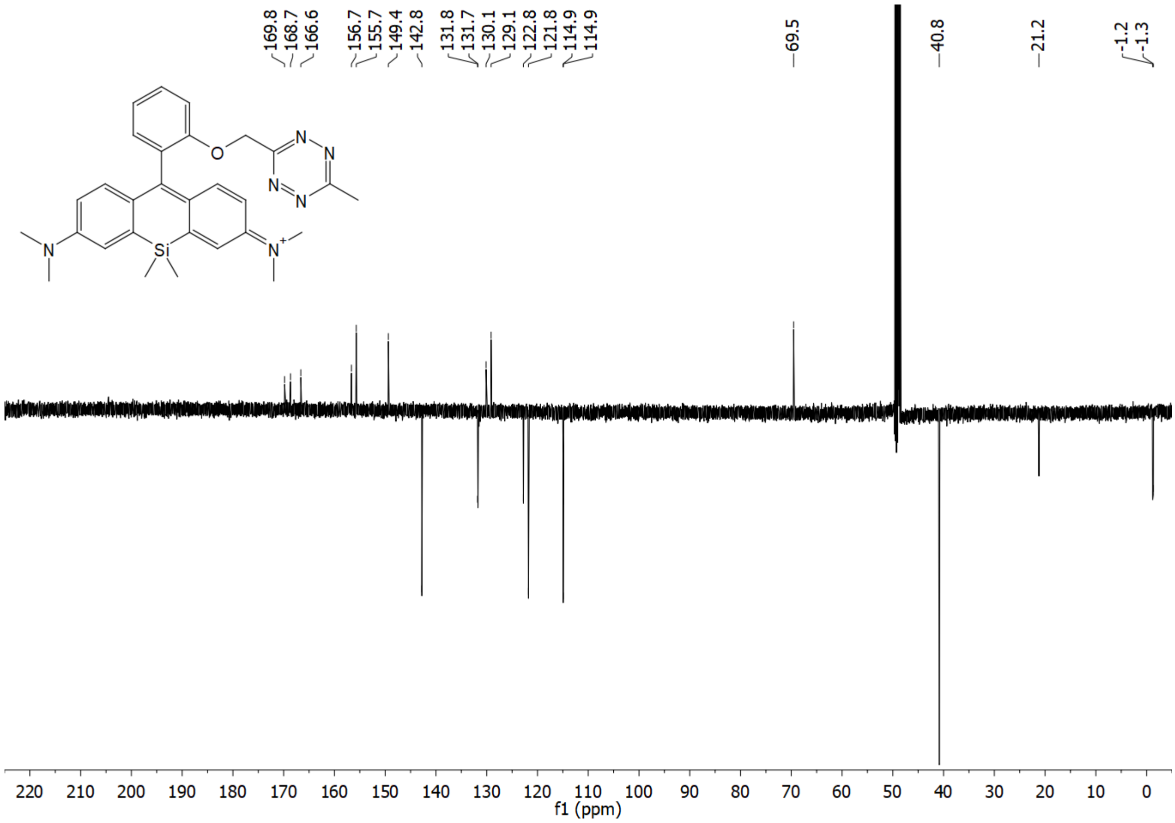

$^{1}H$ NMR of **Ref-2** ($CD_3OD$)

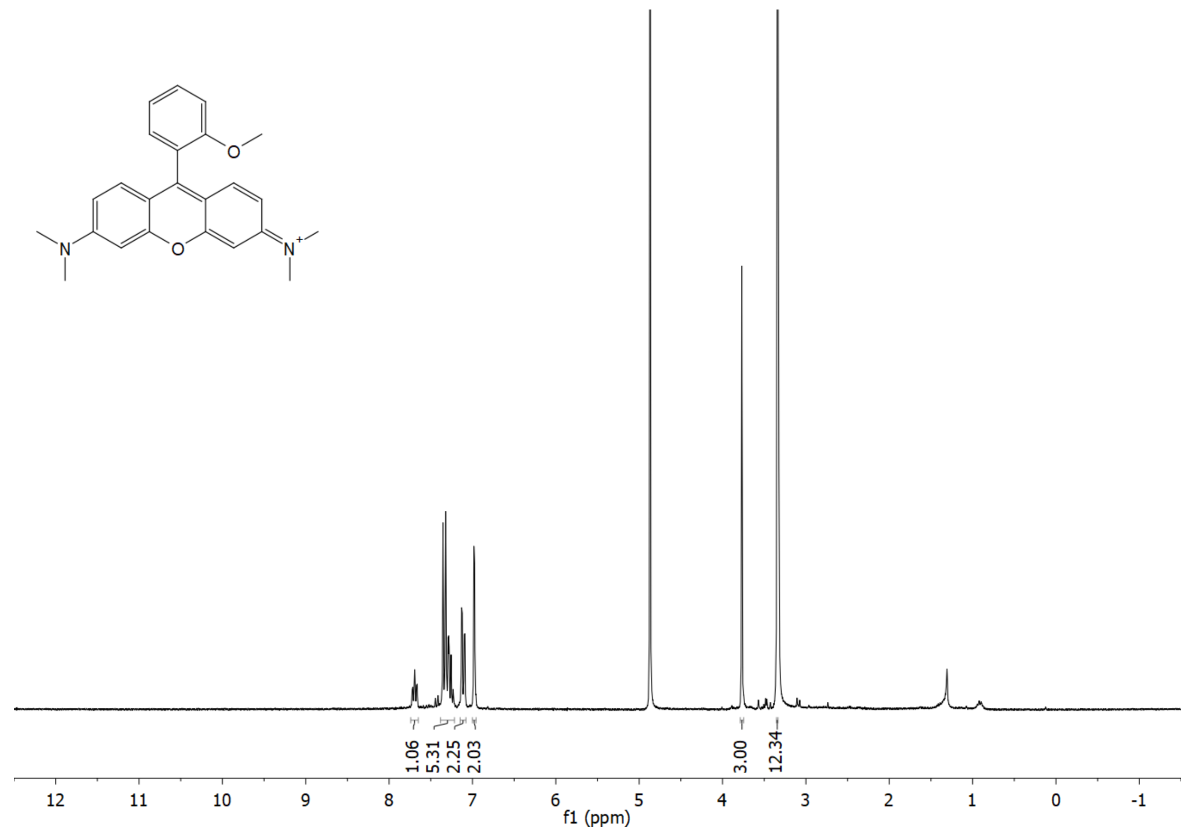

$^{13}C$ NMR (APT) of **Ref-2** ($CD_3OD$)

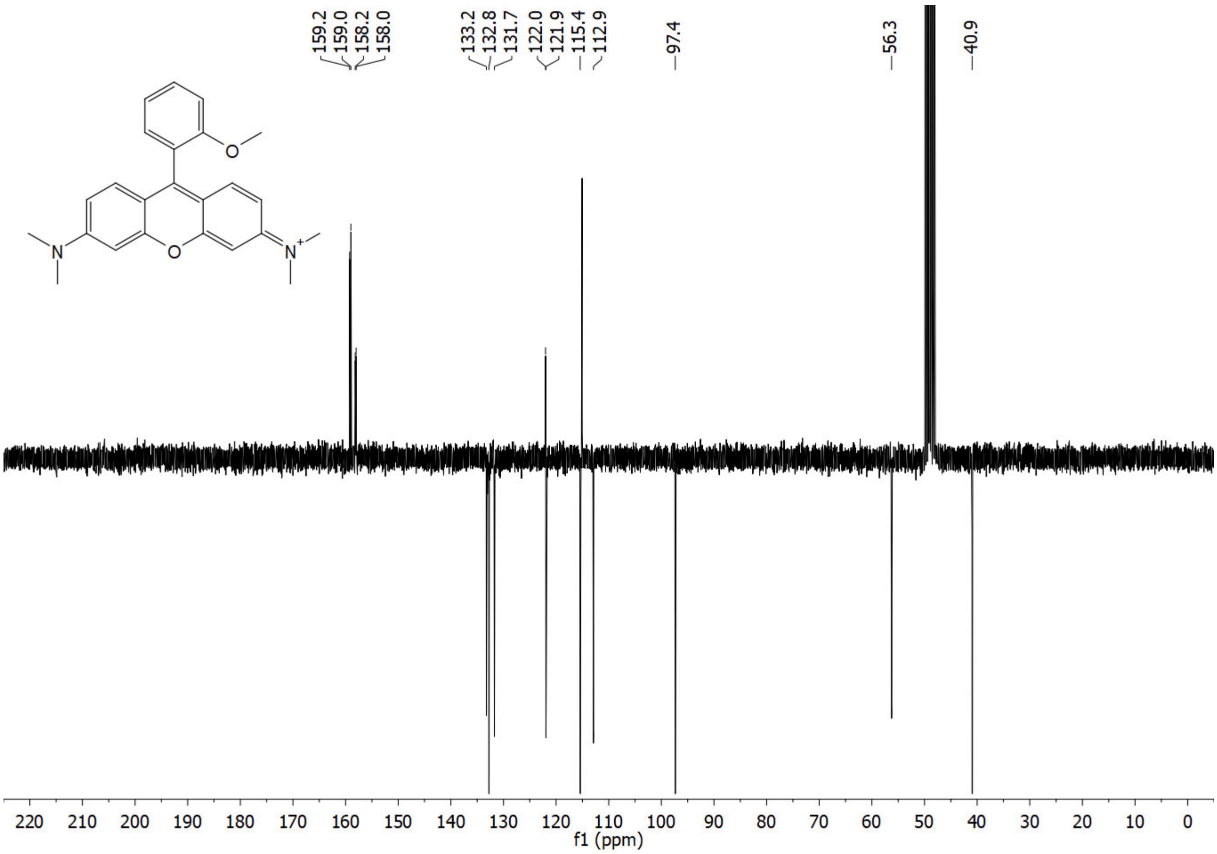
